# Stable population structure in Europe since the Iron Age, despite high mobility

**DOI:** 10.1101/2022.05.15.491973

**Authors:** Margaret L. Antonio, Clemens L. Weiß, Ziyue Gao, Susanna Sawyer, Victoria Oberreiter, Hannah M. Moots, Jeffrey P. Spence, Olivia Cheronet, Brina Zagorc, Elisa Praxmarer, Kadir Toykan Özdoğan, Lea Demetz, Pere Gelabert, Daniel Fernandes, Michaela Lucci, Timka Alihodžić, Selma Amrani, Pavel Avetisyan, Christèle Baillif-Ducros, Željka Bedić, Audrey Bertrand, Maja Bilić, Luca Bondioli, Paulina Borówka, Emmanuel Botte, Josip Burmaz, Domagoj Bužanić, Francesca Candilio, Mirna Cvetko, Daniela De Angelis, Ivan Drnić, Kristián Elschek, Mounir Fantar, Andrej Gaspari, Gabriella Gasperetti, Francesco Genchi, Snežana Golubović, Zuzana Hukeľová, Rimantas Jankauskas, Kristina Jelinčić Vučković, Gordana Jeremić, Iva Kaić, Kevin Kazek, Hamazasp Khachatryan, Anahit Khudaverdyan, Sylvia Kirchengast, Miomir Korać, Valérie Kozlowski, Mária Krošláková, Dora Kušan Špalj, Francesco La Pastina, Marie Laguardia, Sandra Legrand, Tino Leleković, Tamara Leskovar, Wiesław Lorkiewicz, Dženi Los, Ana Maria Silva, Rene Masaryk, Vinka Matijević, Yahia Mehdi Seddik Cherifi, Nicholas Meyer, Ilija Mikić, Nataša Miladinović-Radmilović, Branka Milošević Zakić, Lina Nacouzi, Magdalena Natuniewicz-Sekuła, Alessia Nava, Christine Neugebauer-Maresch, Jan Nováček, Anna Osterholtz, Julianne Paige, Lujana Paraman, Dominique Pieri, Karol Pieta, Stefan Pop-Lazić, Matej Ruttkay, Mirjana Sanader, Arkadiusz Sołtysiak, Alessandra Sperduti, Tijana Stankovic Pesterac, Maria Teschler-Nicola, Iwona Teul, Domagoj Tončinić, Julien Trapp, Dragana Vulović, Tomasz Waliszewski, Diethard Walter, Milos Zivanovic, Mohamed el Mostefa Filah, Morana Čaušević-Bully, Mario Šlaus, Dusan Boric, Mario Novak, Alfredo Coppa, Ron Pinhasi, Jonathan K. Pritchard

## Abstract

Ancient DNA research in the past decade has revealed that European population structure changed dramatically in the prehistoric period (14,000-3,000 years before present, YBP), reflecting the widespread introduction of Neolithic farmer and Bronze Age Steppe ancestries. However, little is known about how population structure changed from the historical period onward (3,000 YBP - present). To address this, we collected whole genomes from 204 individuals from Europe and the Mediterranean, many of which are the first historical period genomes from their region (e.g. Armenia and France). We found that most regions show remarkable inter-individual heterogeneity. At least 7% of historical individuals carry ancestry uncommon in the region where they were sampled, some indicating cross-Mediterranean contacts. Despite this high level of mobility, overall population structure across western Eurasia is relatively stable through the historical period up to the present, mirroring geography. We show that, under standard population genetics models with local panmixia, the observed level of dispersal would lead to a collapse of population structure. Persistent population structure thus suggests a lower effective migration rate than indicated by the observed dispersal. We hypothesize that this phenomenon can be explained by extensive transient dispersal arising from drastically improved transportation networks and the Roman Empire’s mobilization of people for trade, labor, and military. This work highlights the utility of ancient DNA in elucidating finer scale human population dynamics in recent history.

## Introduction

Ancient DNA (aDNA) sequencing has provided immense insight into previously unanswered questions about human population history. Initially, sequencing efforts were focused on identifying the main ancestry groups and transitions during prehistoric times, for which there is no written record. Recently, aDNA sampling has expanded to more recent times, allowing the study of movements of people using genetic data alongside the well-studied historical record. However, we lack a comprehensive assessment of historical genetic structure, including characterizing genetic heterogeneity and interactions across regions. Integrating historical period genetics will be instrumental to better understanding the development of European and Mediterranean population structure from prehistoric to present-day.

Prehistoric ancient genomes have allowed disentangling the movements of people and technologies across two major demographic transitions in prehistoric western Eurasia: first the farming transition ∼7,500 BCE (Lazaridis et al., 2014; Skoglund et al., 2012), and later the Bronze Age Steppe migrations ∼3,500 BCE (Haak et al., 2015). Over the course of generations, genetically differentiated peoples across western Eurasia came together and admixed. As a result, most present-day European genomes can be modeled as a three-way mixture of these prehistoric groups: Western Hunter-Gatherers, Neolithic farmers, and Bronze Age Herders from the Steppe (Haak et al., 2015; Lazaridis et al., 2014) with minor contributions from other groups (Antonio et al., 2019; Fernandes et al., 2020; Lazaridis et al., 2016; Marcus et al., 2020; Mathieson et al., 2018). These ancestry components are present at different proportions across western Eurasia, leading to a pattern where the genetic structure of Europe mirrors its geography (Novembre et al., 2008).

Given that the major ancestry components of present-day west Eurasians were largely established by the end of the Bronze Age, it is unclear how and what types of demographic processes impacted the genetic make-up of western Eurasia over the last ∼3000 years, from the end of the Bronze Age to present-day. Recent studies of historical period genomes from individual regions shed light on this question; they paint a picture of heterogeneity and mobility, rather than of stable population structure. In the city of Rome alone, the population was dynamic and harbored a large diversity of ancestries from across Europe and the Mediterranean from the Iron Age (∼1000 BCE) through the Imperial Roman period (27 BCE-300 CE) (Antonio et al., 2019). Historical genomes from the Iberian Peninsula also highlight gene flow from across the Mediterranean (Olalde et al., 2019).

These regional reports fit well with archaeological and historical records. By the Iron Age, sea travel was already common, enabling peoples from across the Mediterranean to come into contact for trade (Abulafia, 2011; Broodbank, 2013). Subsequently, the Roman Empire leveraged its organization, labor force, and military prowess to build upon existing waterways and roads throughout Europe and create a united Mediterranean for the only time in history (Beard, 2015; Harper, 2017; Symonds, 2017). Not only did the Empire provide a means for movement, it also provided a reason for individuals to move. Empire building activities, broadly categorized into military, labor, and trade, pulled in people and resources from inside and outside the Empire (Scheidel, 2019).

We sequenced 204 new historical period genomes from across Europe and the Mediterranean to more comprehensively investigate the Roman Empire’s impact on the genetic landscape suggested by these regional reports of heterogeneous, mobile populations. By analyzing genetic similarities between individuals across historical Eurasia, we were able to quantify individual movements during this time. Based on population genetic simulations, we explore potential explanations of how population structure may be maintained in the face of frequent individual dispersal.

## Results

### 204 new historical genomes from Europe and the Mediterranean

We collected whole genomes from 204 individuals across 53 archaeological sites in 18 countries spanning Europe and the Mediterranean (Figure 1 - figure supplement 1), 26 of these individuals were recently reported (Moots et al., 2022). This collection includes the first historical genomes (Iron Age and later, i.e. after 1000 BCE) from present-day Armenia, Algeria, Austria, and France. Dates for 126 samples were directly determined through radiocarbon dating, and were used alongside archaeological contexts to infer dates for the remaining samples.

DNA was extracted from either the powdered cochlear portion of the petrous bone (n = 203) or from teeth (n = 1). Libraries were partially treated with uracil-DNA glycosylase (UDG) and screened for ancient DNA damage patterns, high endogenous DNA content, and low contamination. We performed whole genome sequencing to a median depth of 0.92x (0.16x to 2.38x).

For downstream integration with published data, pseudohaploid genotypes were called for the 1240k SNP panel (Mathieson et al., 2015), resulting in a median of 685,058 SNPs (167,000 to 1,029,345) per sample. We analyzed newly reported genomes in conjunction with 2,033 present-day genomes, 1,998 prehistoric genomes, and 764 published historical period genomes ((Clemente et al., 2021; Kovacevic et al., 2014; Mallick et al., 2023; Pagani et al., 2016; Saupe et al., 2021; Žegarac et al., 2021), primary AADR sources cited in Methods). Genomes were grouped by regions and time periods (Figure 1) and analyzed using principal component analysis (PCA) and *qpAdm* modeling (Haak et al., 2015).

**Figure 1.**
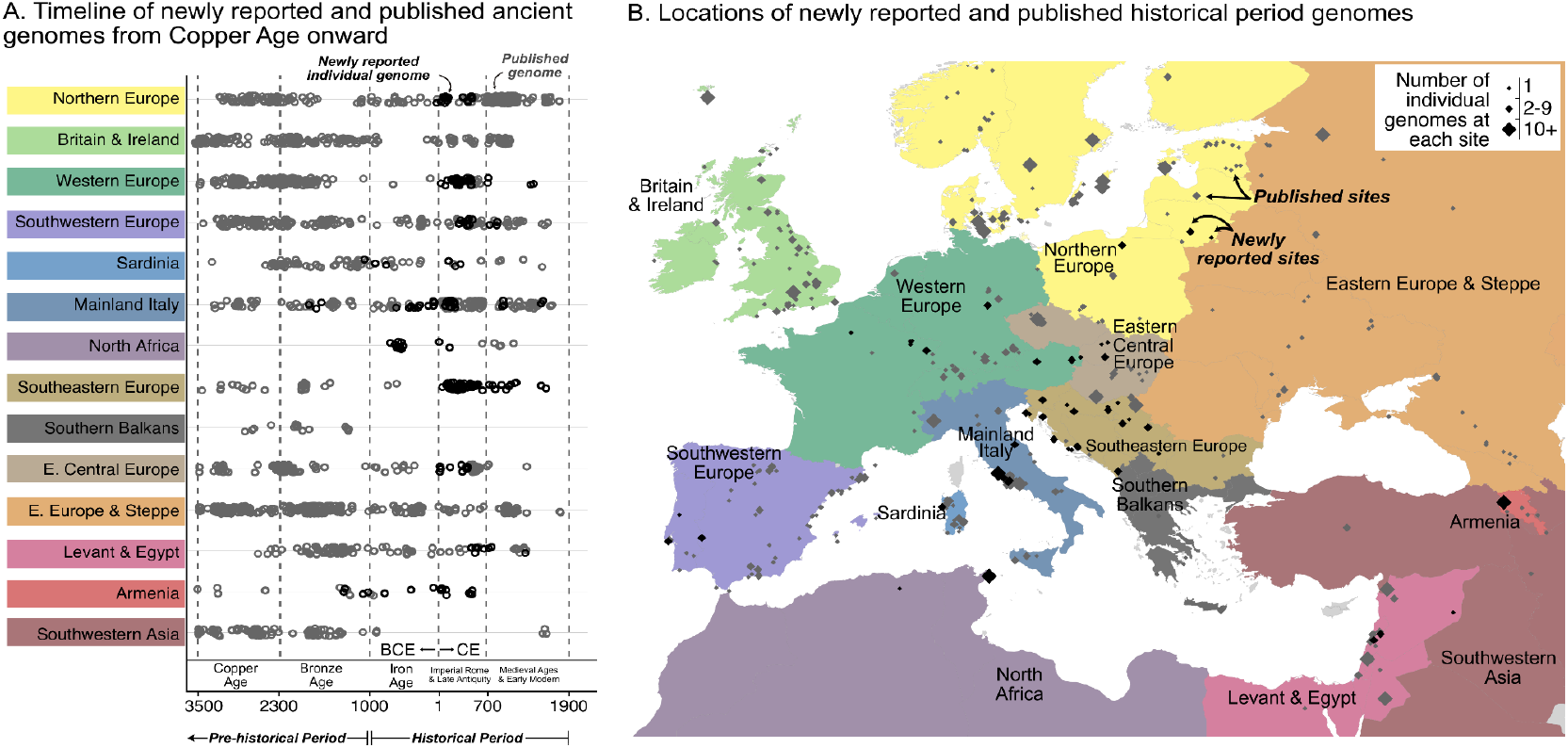
Timeline of new and published genomes. (A) 204 newly reported genomes (black circles) are shown alongside published genomes (gray circles), ordered by time and region (colored the same way as in B). (B) Sampling locations of newly reported (black) and published (gray) genomes are indicated by diamonds, sized according to the number of genomes at each location.

### Local historical population structure varies across regions

To investigate historical population structure, we categorized the data into 14 geographical regions, split into three sub-periods of the historical period: Iron Age (1000-1 BCE), Imperial Rome & Late Antiquity (1-700 CE), and Medieval Ages & Early Modern (700-1950 CE). We then characterized inter-individual heterogeneity within these spatio-temporal groups by examining (1) variation of projections onto a PCA space of present-day genomes (Figure 2 - figure supplement 1), (2) genetic groups identified by *qpAdm* and clustering across time within a region, and (3) admixture modeling of genetic groups.

A majority of regions have highly heterogeneous populations in at least one historical time period (Figure 2 - figure supplement 2). This is illustrated by both the visual spread in PCA and the genetically distinct clusters of individuals based on pairwise modeling with *qpAdm* (Haak et al., 2015; Harney et al., 2021). On average, we identified 10 genetic clusters within each region present during the historical period, with a minimum of two and a maximum of 23. With genetically similar samples grouped together, we have more power relative to individual-level analyses when performing admixture modeling on clusters of interest using *qpAdm*.

Regional vignettes reveal various patterns of historical population structure. In Armenia, for example, the population is highly homogeneous at any given time (Figure 2). After the Copper Age, there are two distinct genetic clusters, separated by a temporal split around 772-403 BCE (Figure 2BC). The earlier cluster (C1) includes newly reported samples (n=5) from Beniamin and published ones (n=6) from five other sites. This cluster cannot be modeled by any single source of ancestry using existing data. The later cluster (C3), which contains newly reported samples (n=12) from Beniamin dating between 403 BCE-500 CE, is genetically similar to present-day Armenians (excluding two Kurdish individuals; Figure 2C). Despite the split, there is evidence of partial continuity between the earlier and later clusters: the later (C3) can be modeled using around 50% of the earlier cluster (C1) and an additional source of Steppe ancestry. Historical genomes from Northern Europe, particularly newly reported genomes from Lithuania and Poland, exhibit a similar level of homogeneity (Figure 2 - figure supplement 2).

**Figure 2.**
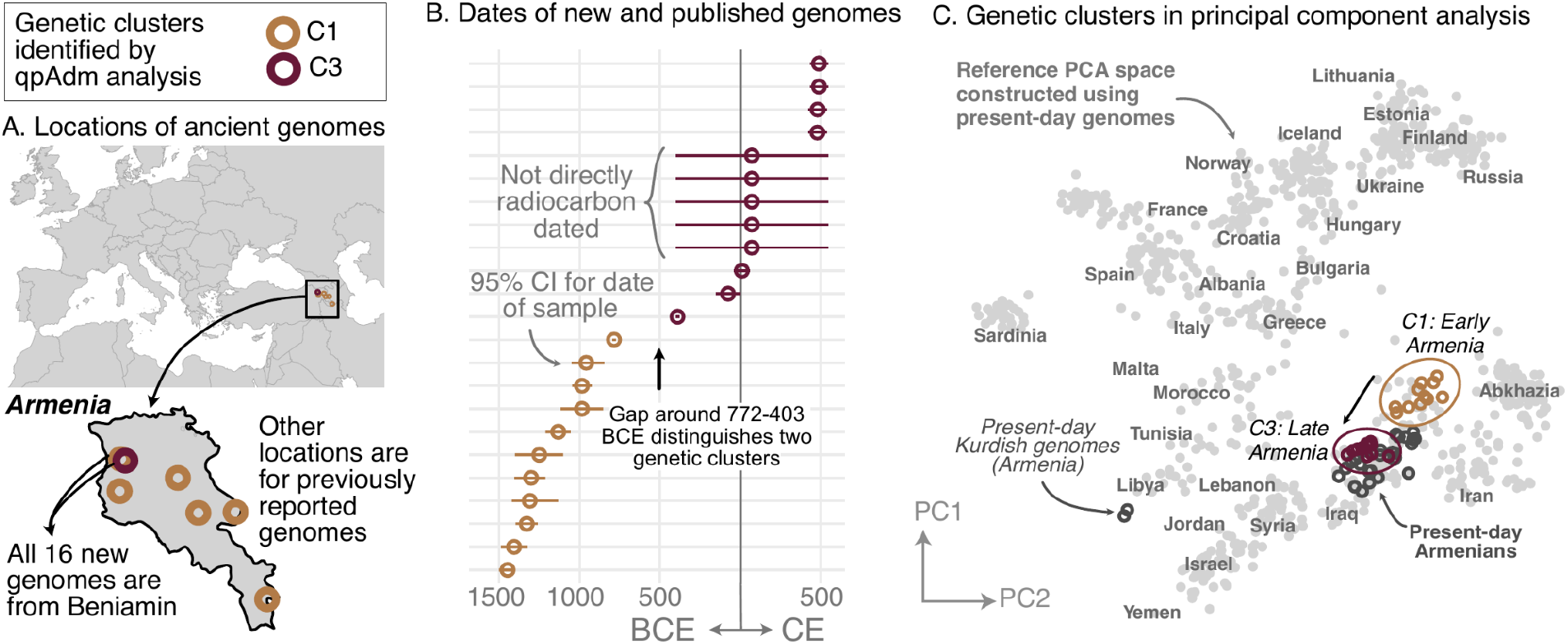
Armenia: two homogeneous genetic clusters distinguished by a temporal shift. (A) Sampling locations of ancient genomes (open circles) colored by their genetic cluster identified using qpAdm modeling. (B) Date ranges for the genomes: each line represents the 95% confidence interval for the radiocarbon date or the upper and lower limit of the inferred date, and the point represents the midpoint of that range. (C) Projections of the genomes onto a PCA of present-day genomes (gray points labeled by their population). Present-day genomes from Armenia are shown with dark gray open circles.

In contrast to the homogeneity of the Armenian population, most of the regions, including Italy, Southeastern Europe, and Western Europe, had strikingly heterogeneous populations. Newly collected samples reinforce previous findings of high heterogeneity in Rome, including a large portion of the population having affinities for present-day Near Eastern populations (Antonio et al., 2019; Posth et al., 2021) (Figure 3 - figure supplement 1). Interestingly, Southeastern European and Western European individuals during the Imperial Roman & Late Antiquity period also exhibit high heterogeneity, on par with that of contemporaneous Italy (Figures 3 and 4).

**Figure 3.**
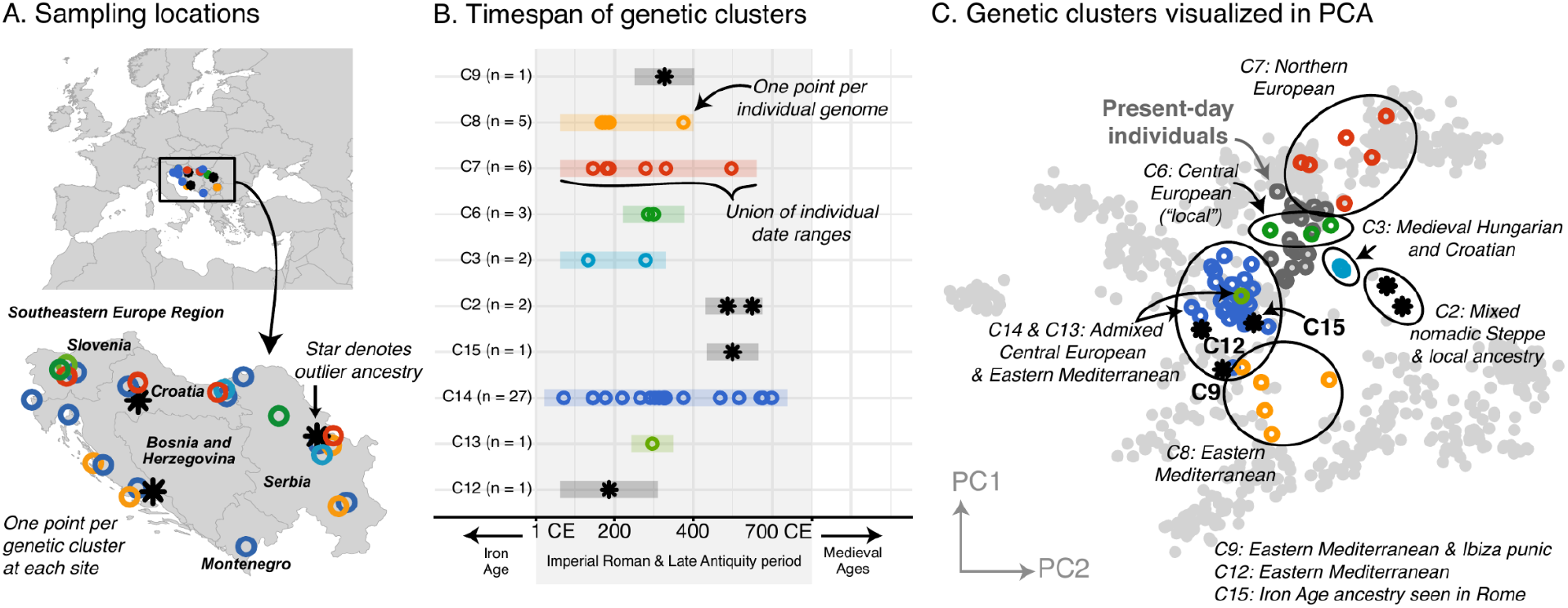
Southeastern Europe: highly heterogeneous Imperial Roman and Late Antiquity period population. (A) Sampling locations of genetic clusters are represented by a single point per location. Outlier ancestries are black stars, all others are open circles colored by genetic cluster. (B) Colored bars span the minimum and maximum of the date ranges of samples (95% confidence interval from radiocarbon dating or archaeological range). Points are the mean of an individual’s date range. (C) Projections of the ancient genomes onto a PCA of present-day genomes (gray points). Population labels for the PCA reference space are shown in Figure 2C. Present-day genomes from Southeastern Europe are shown with dark gray open circles.

**Figure 4.**
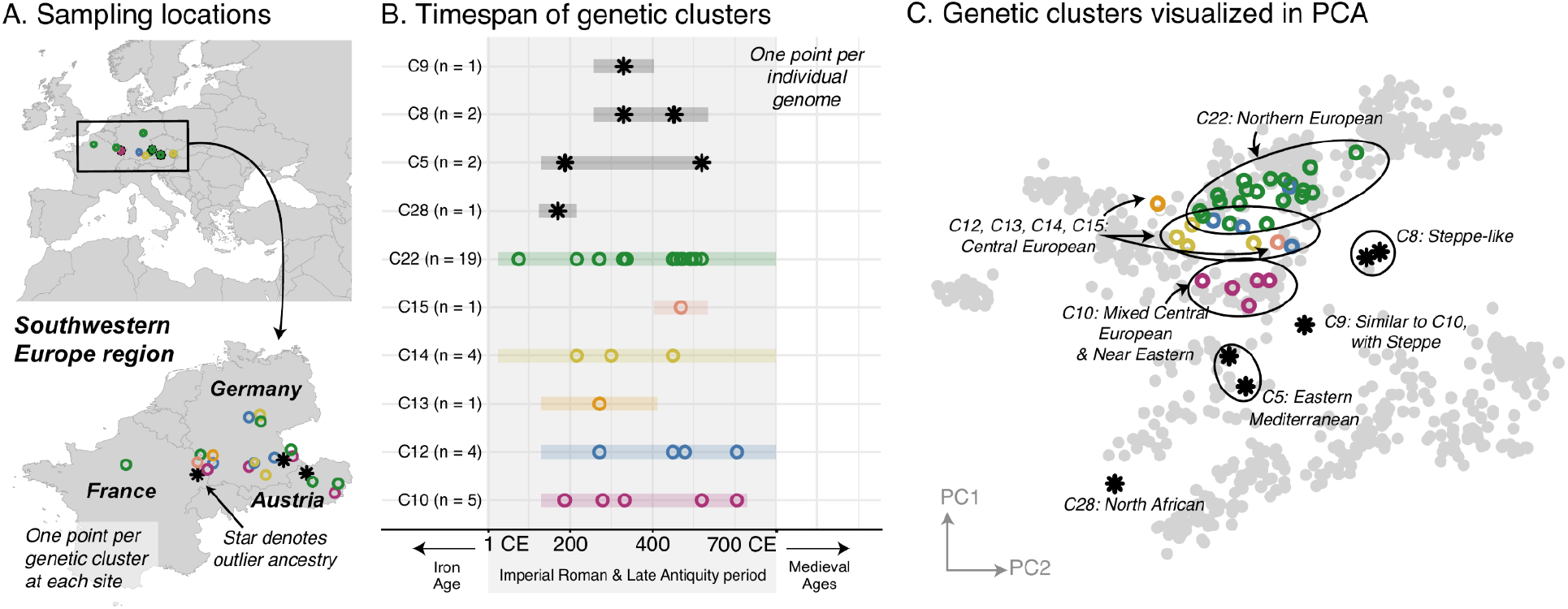
Western Europe: heterogeneous Imperial Roman and Late Antiquity period population. (A) Sampling locations of genetic clusters are represented by a single point per location. Outlier ancestries are black stars, all others are open circles colored by genetic cluster. (B) Colored bars span the minimum and maximum of the date ranges of samples (95% confidence interval from radiocarbon dating or archaeological range). Points are the mean of an individual’s date range. (C) Projections of the ancient genomes onto a PCA of present-day genomes (gray points). Population labels for the PCA reference space are shown in Figure 2C. Present-day genomes from Southeastern Europe are shown with dark gray open circles.

Furthermore, these ancestries are often shared across regions. In Southeastern Europe, a core group of individuals have ancestry similar to that of present-day and contemporaneous Central Europeans (C6), while other clusters have ancestry similar to that of Northern Europeans (C7) and Eastern Mediterraneans (C8) (Figure 3C). These ancestry groups are found in contemporaneous Italy and Western Europe as well (Figure 4C, Figure 3 - figure supplement 1). We also observe individuals of eastern nomadic ancestry, similar to that of Sarmatian individuals previously reported, in both Western Europe (C8, n=2) and Southeastern Europe (C2, n=2).

Overall, we see remarkable local genetic heterogeneity as well as cross-regional similarities which point to common ancestry sources and, on a broader scale, demographic events affecting different regions in similar ways.

### At least 7-11% of historical individuals are ancestry outliers

The high regional genetic heterogeneity with long range, cross-regional similarities suggests historical populations were highly mobile. We therefore sought to quantify the amount of movement during the historical period by estimating the proportion of individuals who are ancestry outliers with respect to all individuals found in the same region. We considered an individual an outlier if they belonged to an ancestry cluster that is underrepresented (consisting of fewer than 5% of individuals in a region or at most two individuals) within their sampling region from the Bronze Age up to present-day. To focus on first-generation migrants as well as long-range movements, we further identified outlier individuals who can be modeled as 100% (i.e. “one-component model”) of a majority ancestry cluster found in a different region.

In total, we identified 11% of individuals as outliers, and could connect 7% of individuals to a putative source in a different region (Figure 5A). Based on the regions where these outliers and their sources originated, we created a network to illustrate their movements (Figure 5B). This network reveals the interconnectedness of Europe and the Mediterranean during the historical period. For example, as discussed above, the Armenian population is quite homogeneous (Figure 2). Unsurprisingly, no outliers were found within Armenia; however, we found outlier individuals in the Levant and Italy who can be putatively traced back to Armenia according to their ancestry (Figure 5C; blue outgoing arrows from Armenia). In contrast, the heterogeneous population in Italy connects it to many other regions, with bi-directional movement in most cases. In North Africa, outliers found in Iron Age Tunisia (Moots et al., 2022) indicate movements from many regions in Europe, and North African-like outliers were found in Italy and Austria (Western Europe). North African ancestry in Italy is supported by a single previously reported individual from the Imperial Roman period (R132) (Antonio et al., 2019).

**Figure 5.**
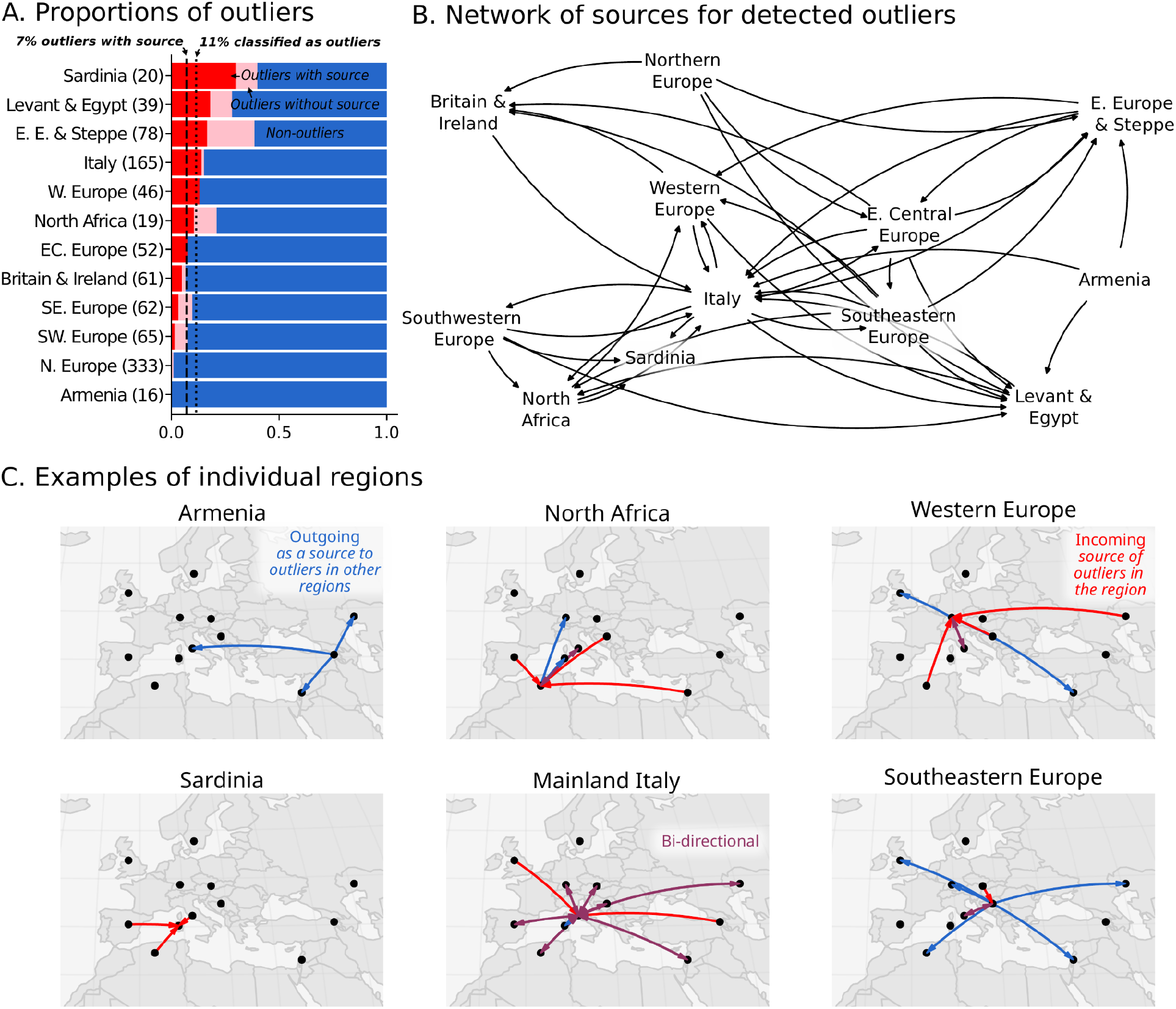
Ancestry outliers and their potential sources. (A) The proportions of outliers in each region were determined by individual pairwise qpAdm modeling followed by clustering. (B) Sources were inferred by one component qpAdm modeling of resulting clusters with all genetic clusters in the dataset. In the network visualizations, nodes are regions and directed edges are drawn from sources to outliers (i.e. potential migrants). The full network of source to outlier is shown. (C) Examples of individual regions are shown in greater detail.

Similar North African ancestry in Western Europe is supported by a single individual, R10667, from Wels, Austria, a site located on the frontier of the Roman Empire (C28 in Figure 4). This individual from Austria can be modeled using Canary Islander individuals from the Medieval Ages or an Iron Age outlier (distinguished by having more sub-Saharan ancestry) from Kerkouane, a Punic city near Carthage in modern-day Tunisia.

The 7% estimate for outliers with source should be considered conservative for the proportion of “non-local” individuals. There are several cases where a cluster comprises more than 5% of the individuals in the region, but are clearly of a different ancestry than the majority and seem to be transient (only found in a single sub-period of the historical period). For example, in Southeastern Europe (Figure 3B), Imperial Roman & Late Antiquity individuals in C8 are (1) of distant ancestry (Near Eastern) and (2) not found in previous or subsequent time periods. However, since there are five individuals in this cluster, it does not meet our strict criteria for outlier consideration. Additionally, many clusters of underrepresented ancestry cannot be modeled as one-component models because they are recently admixed (i.e. require two or more ancestry components) or of ancestry not sampled elsewhere. Thus, we expect the actual proportion of individuals involved in long distance movements to be higher than reported here.

### Spatial population structure is relatively stable in the last 3,000 years

The remarkable amount of heterogeneity and mobility in the historical period leads to the question of what impact this might have had on population structure over time. To investigate this, we sought to quantify the overall change in population structure across time, from prehistoric to present-day. To assess the spatial structure of population differentiation, we calculated F_ST_ across groups of individuals on a sliding spatial grid in each time period and related it to their mean geographic distance. In each time period, we recovered the classical pattern of isolation-by-distance (Figure 6A), where individuals closer in geographic space are also more similar genetically. Across time periods, we see a large decrease in overall F_ST_ from the Mesolithic & Neolithic periods to the Bronze Age (approximately 10,000-2300 BCE), coinciding with the major prehistoric migrations (Haak et al., 2015; Lazaridis et al., 2014). From the Bronze Age onward, however, F_ST_ does not decrease further with time, indicating that the level of genetic differentiation across space is relatively stable from the Bronze Age to present-day.

**Figure 6.**
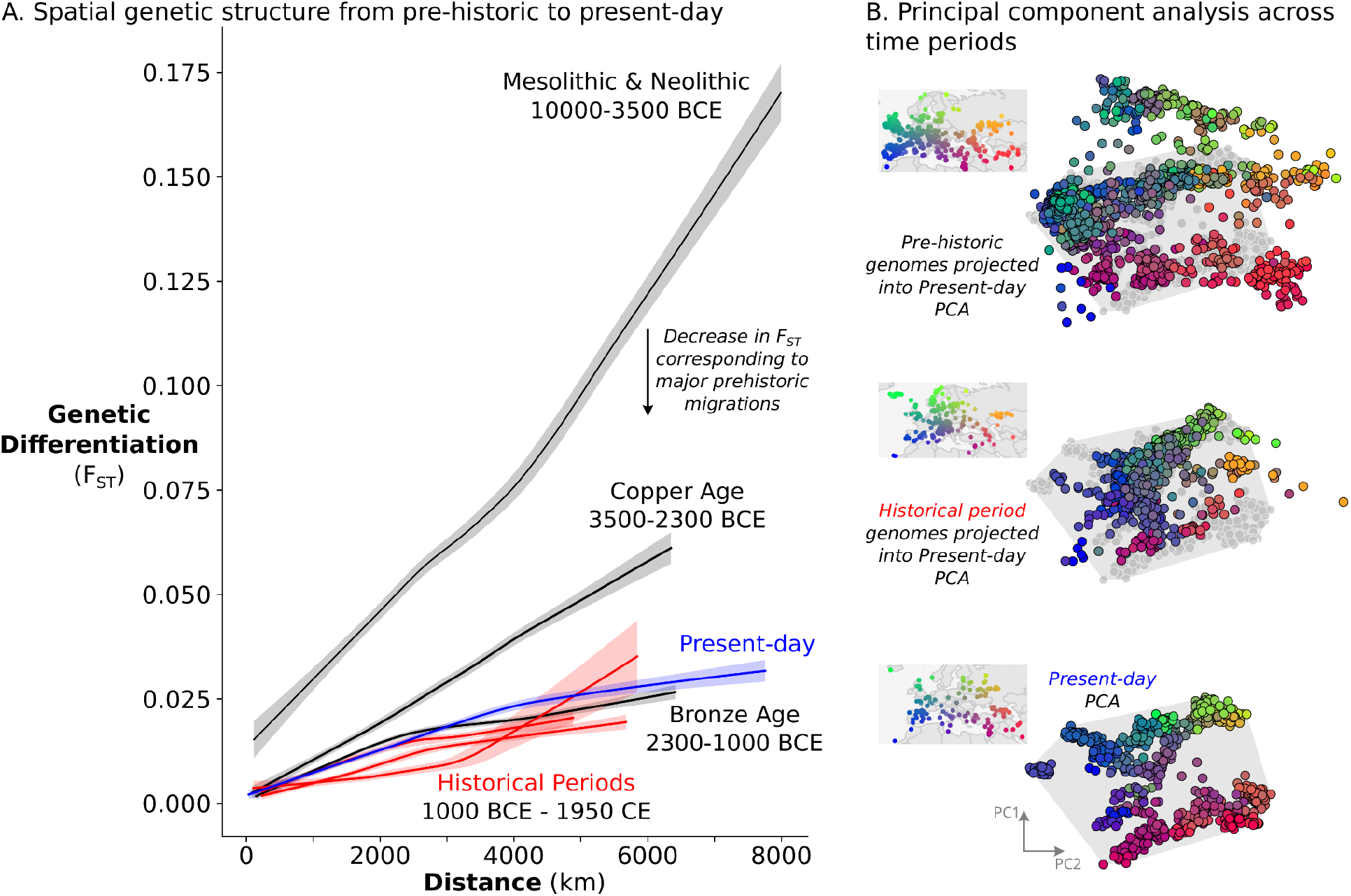
Relatively stable population structure from Bronze Age to present-day. (A) Overall genetic differentiation between populations (measured by F_ST_) and its relationship to geographical distance (spatial structure) is similar from Bronze Age onward. Confidence intervals were calculated through a bootstrap procedure, using 200 bootstrap replicates. (B) In PC space, each genome is represented by a point, colored based on their origin (for present-day individuals) or sampling location (for historical samples). The PC space is established by present-day samples (bottom), onto which either historical period (middle) or prehistoric genomes (top) were projected. For projections, the present-day samples are shown in gray, and their extent is visualized by a gray polygon.

To assess not only the amount, but also the structure of geographic population differentiation, we compared the “genetic maps” of historical period and present-day genomes. To construct these “maps”, we performed principal component analysis on 829 present-day European and Mediterranean genomes sampled across geographical space (Figure 6B, bottom) and projected historical period genomes onto the same PC space. Echoing close correspondence between genetic structure and geographic space in present-day Europeans (Novembre et al., 2008), we recovered similar spatial structure for historical samples as well, although noisier due to a narrower sampling distribution and higher local genetic heterogeneity (Figure 6B, middle). The similarity in structure between present-day and historical period is especially striking in comparison to a projection of prehistoric genomes, which shows much weaker correspondence to the present-day PCA as well as to geographic space (Figure 6B, top). Together, our analyses indicate that European and Mediterranean population structure has been relatively stable over the last 3,000 years.

This raises the question: is it surprising for stable population structure to be maintained in the presence of ∼7-11% long-range migration? To address this, we simulated Wright-Fisher populations evolving neutrally in continuous space. In these simulations, spatial population structure is established through local mate choice and limited dispersal, which we calibrated to approximately match the spatial differentiation observed in historical-period Europe (Figure 6A, Figure 7A and Figure 7 - figure supplement 1, maximum F_ST_ of ∼0.03). We then allowed a proportion of the population to disperse longer distances, empirically matching the migration distances we observed in the data during the historical period (Figure 7 - figure supplement 2). Even with long-range dispersal as low as 4%, we observe decreasing F_ST_ over 120 generations (∼3000 years with a generation time of 25 years) as individuals become less differentiated genetically across space (Figure 7B). At 8%, F_ST_ decreases dramatically within 120 generations as spatial structure collapses to the point that it is hardly detectable in the first two principal components (Figure 7C). These simulations indicate that under a basic spatial population genetics model we would expect structure to collapse by present-day given the levels of movement we observe.

**Figure 7.**
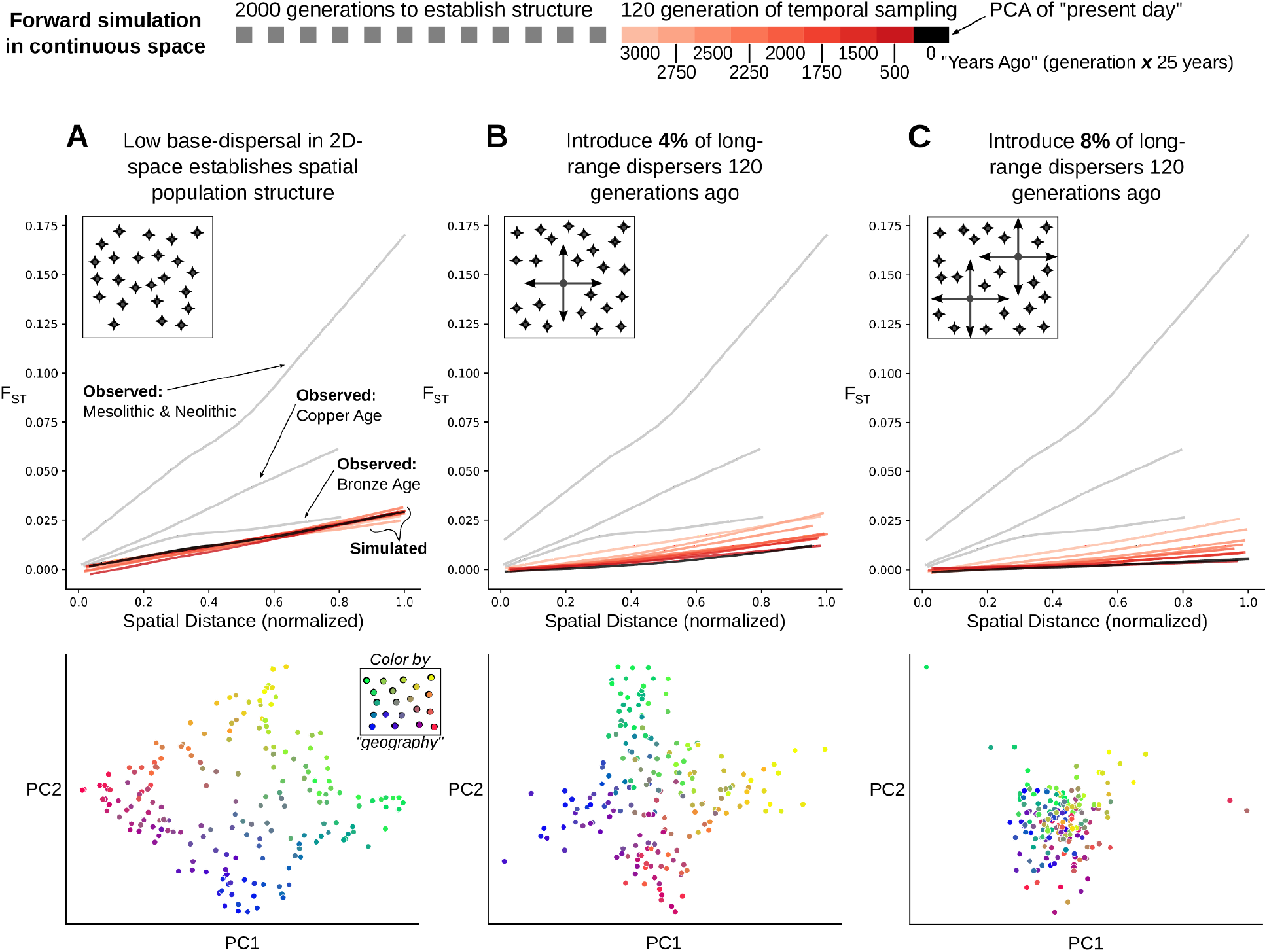
Simulation of population structure with and without long-range dispersal. (A) A base model of spatial structure is established by calibrating per-generation dispersal rate to generate a maximum F_ST_ of ∼0.03 across the maximal spatial distance, and visualized using PCA. In addition to this base dispersal, either 4% (B) or 8% (C) of individuals disperse longer distances, and the effect is tracked by analyzing spatial F_ST_ through time, as well as PCA after 120 generations of long-range dispersal.

## Discussion

In summary, we observed largely stable spatial population structure across western Eurasia and high mobility of people evidenced by local genetic heterogeneity and cross-regional connections. These two observations are seemingly incompatible with each other under standard population genetics assumptions.

A possible explanation for this apparent paradox is that our simulations did not capture some key features of human behavior and population dynamics. In the simulated populations, migration implies both movement and reproduction with local random mate choice. However, in real human populations migration can be more complex: people do not necessarily reproduce where they migrate, and reproduction is not necessarily random. We hypothesize that in the historical period there was an increasing decoupling of movement and reproduction, compared to prehistoric times. For the spread of Farmer and Steppe ancestry, we know that these prehistoric migrations would take hundreds of years to traverse the continent (Allentoft et al., 2015; Haak et al., 2015; Lazaridis et al., 2016). In contrast, in the historical period, there were dense travel networks of roads and waterways as well as clear incentives for cross-Mediterranean and cross-continental movement (Abulafia, 2011; Beard, 2015; Broodbank, 2013; Symonds, 2017). This enabled people to travel cross-continental distances on the order of weeks or months, well within their lifetimes (Figure 5 - figure supplement 2, 3) (Scheidel, 2015).

The Roman Empire is particularly important in understanding how transient mobility could become a unique hallmark of this period. During the expansion of the Empire, existing and new cities quickly expanded as hubs for trade and labor. Urban-military complexes emerged along the frontier as military forces established themselves and drew in local communities which sought protection or economic benefit (Séguy, 2019). To support these rapidly growing economic city-centers, human capital beyond the local population was necessary, thus drawing in people from far away places either freely or forcibly (e.g. slavery, military). According to a longstanding historical hypothesis, the Urban Graveyard Effect, the influx of migrants in city-centers disproportionately contributed to the death rate over the birth rate; a process which would contribute to observing individuals as “transient” migrants (de Ligt and Tacoma, 2016). Long-range, transient migration, combined with the Roman Empire’s highly efficient travel networks (Cherry, 2007; Oleson, 2008; Scheidel, 2015) may explain the genetically heterogeneous populations, especially along the frontier regions (e.g. Serbia, Croatia, and Austria).

With transient mobility as the main contributor to the observed heterogeneity, it remains unclear what additional demographic processes contributed to the maintenance of spatial genetic structure. The collapse of the Empire involved a loss of urban-military complexes and depopulation of cities, followed by ruralization (Burgess, 2007; Dey, 2015; Roymans et al., 2020). Without the Empire incentivising trade and movement, there may have been little motive for individuals to remain in these suddenly remote regions.

If this hypothesis is true, we would expect a reduction in local genetic heterogeneity after the collapse of the Empire. Unfortunately, we do not have this period sampled densely enough to assess this comprehensively. The lack of samples is further amplified by the fact that ancient DNA comes from archaeological excavations, which tend to be enriched in urban areas; a stone mausoleum in the city center, for example, will produce more surface scatter than a wood farmhouse, making urban areas more likely for excavation (Bowes, 2011). This makes it difficult to comprehensively address differences in rural versus urban demography. Collecting more genetic data from both urban and rural contexts across the historical period will be a valuable future step in understanding how spatial population structure was maintained. Furthermore, it could elucidate the role of other historical events and peoples, such as the Franks, Lombards, Visigoths, and Huns, during the Migration Period.

Based on genetic analyses and the rich historical record, we hypothesize that both the loss of transient migrants which contributed to population heterogeneity, as well as repopulation by less heterogeneous, but temporally stable, local populations could have helped maintain overall stability of genetic structure from the Iron Age to present-day. This work highlights the utility of ancient DNA in revealing complex population dynamics through direct genetic observations through time and the importance of integrating historical contexts to understand these complexities.

## Materials and Methods

### Sample collection and archaeological sites

The archaeological context for ancient individuals reported in this study is detailed in Appendix 1. Site descriptions were written by the contributing archaeologists.

Descriptions of individual-level burials are included where possible.

Sampling was performed to maximize coverage across Europe and the Mediterranean, as opposed to detailed site level sampling. We particularly focused on regions where there was no published genomic data for the Imperial Roman and Late Antiquity Period at the time of sampling (e.g. France, Austria, the Balkans, Armenia, and North Africa).

Although we aimed to collect samples in the Imperial Roman and Late Antiquity period (approximately 1 CE-700 CE), some samples fall outside of this period due to limited sample availability and/or lack of date specificity at the time of sampling (prior to radiocarbon dating).

### Date determination for individuals and time periods

Determination of time periods and boundaries were influenced by 1) the wide geographic range represented in the newly reported and published data (including historical changes in those regions), 2) the amount of data available in the proposed time periods, 3) the amount of genetic change observed during a time interval, and 4) the types of temporal comparisons made in the analysis.

● Mesolithic and Neolithic 10000 BCE - 3500 BCE
● Copper Age 3500 BCE - 2300 BCE
● Bronze Age 2300 BCE - 1000 BCE
● Iron Age 1000 BCE - 1 CE
● Imperial Rome & Late Antiquity 1 CE - 700 CE
● Medieval Ages & Early Modern 700 CE - 1900 CE
● Present-day 1900 CE - onward

Dates for newly reported study individuals were determined through a combination of radiocarbon dating and archaeologically inferred dates. Sample groups were created based on the finest grouping using a composite of criteria: site, projection in PCA, and archaeologically inferred date or period. We aimed to radiocarbon date at least one sample from each of these groups, summing to a total of 126 samples. For each sample, one gram of petrous bone was sent for radiocarbon dating by Accelerator Mass Spectrometry (AMS) at the Keck Carbon Cycle AMS Facility at the University of California, Irvine. Resulting dates were calibrated using https://c14.arch.ox.ac.uk/oxcal.html.

For samples not directly dated, we assigned the range of directly dated samples within their sample group (n = 49), or the archaeological date where the former was not available (n = 29). Note that one individual (R3477, R3476) had two radiocarbon dates since two samples (a tooth and petrous) were dated and sequenced before they were determined to be from the same individual based on genetic information. Another individual (R9818, R9823) had two radiocarbon dates because both left and right petrosals were sampled, but the sequence data for one turned out to be contaminated. In both cases of identical samples the radiocarbon date ranges were almost entirely overlapping.

In the study we use a single date estimate (for both newly reported and published samples) which is the midpoint of the 95% confidence interval when using AMS dates, and the average of the lower and upper bound inference dates when using archaeological context for dating. The dating approach used for each sample is included in files Appendix 1 (archaeological context) and Appendix 2 (sample metadata). The full AMS and calibration results are reported in Appendix 3.

### DNA extraction, library preparation and sequencing

The 204 ancient genomes reported in this study, 26 of which were recently reported in (Moots et al., 2022), represent a subset of samples screened from 53 archaeological sites across 18 countries.

We isolated and finely ground the cochlear regions of the petrous bones in dedicated clean room facilities at the University of Vienna following the protocols described in (Pinhasi et al., 2019, 2015). Using 50mg of bone powder, DNA was extracted by 18-hour incubation of the powder in a solution of Proteinase-K and EDTA. DNA was eluted in 50 μl 10 mM Tris-HCl, 1 mM EDTA, 0.05% Tween-20, pH 8.0 as in (Dabney et al., 2013; Rohland and Hofreiter, 2007). 12.5-25 uL of DNA extract was used to prepare partial uracil–DNA–glycosylase (UDG) double stranded libraries as described in (Rohland et al., 2015). After a partial (30 minute) UDG treatment, library preparation followed a modified version of the Meyer and Kircher 2010 protocol (Meyer and Kircher, 2010): the initial DNA fragmentation step was not required and MinElute PCR purification kits (Qiagen) were used for all library clean-up steps. Libraries were double-indexed with Accuprime Pfx Supermix. The PCR cycling conditions were as follows: an initial denaturation at 95°C for 5 min followed by 12 cycles of 95°C for 15 seconds, 60°C for 30 seconds and 68°C for 30 seconds with a final elongation at 68°C for 5 min. After indexing, the libraries were purified using the MinElute system (Qiagen) and eluted in 25uL of 1 mM EDTA, 0.05% Tween-20.

Libraries were screened based on Qubit concentration and visual validation of Bioanalyzer peaks for an initial low coverage (MiSeq or NextSeq) screening run.

### Processing sequence data and sample screening

Newly reported samples were initially sequenced to low coverage on MiSeq or NextSeq for screening. Following demultiplexing of the sequencing libraries, reads were trimmed, aligned, filtered for quality, and deduplicated. The following sequence data processing pipeline was applied to both screening and full sequencing runs for new data.

Adapters were removed from sequence reads using Cutadapt (v1.14) (Martin, 2011). Then, for each sample, reads were processed further (a) with the 2 base pairs at either end of the reads trimmed off and (b) without trimming. Since partial UDG treatment was performed on the libraries, a damage signature consisting of elevated C>T transitions on the 5’ end and G>A transitions on the 3’ end should remain at the ends of reads. Therefore, analyzing untrimmed, aligned reads would allow us to assess the amount of the ancient DNA damage signature present in a sample, and to use this as a criteria for authenticating that the sampled DNA is ancient. Other than the variable trimming parameter for the ends of the reads, all other parameters remained the same for both screening and high coverage sequencing data.

Following variable trimming, reads were filtered for minimum length of 30, then aligned to hg19 using bwa (0.7.15-r1140) (Li and Durbin, 2009), with seed length disabled (−l 350). For each sample, aligned reads were sorted by coordinate using Picard’s SortSam (version 2.9.0-1-gf5b9f50-SNAPSHOT) and read groups were added using Picard’s AddOrReplaceReadGroups (version 2.9.0-1-gf5b9f50-SNAPSHOT) (http://broadinstitute.github.io/picard/). Reads with mapping quality < 25 (including unaligned reads) were filtered out. For higher coverage sequencing runs, this process was parallelized by splitting raw fastq files and merging after alignment, sorting, and quality filtering. Duplicates were removed using samtools rmdup (http://www.htslib.org/doc/samtools.html). Genome-wide and chromosomal coverage were assessed using depth-cover (version 1.0.3, https://github.com/jalvz/depth-cover).

Samples were screened and selected using the following criteria: 1) >20% reads aligned to the hg19 build of the human genome; 2) a C>T mismatch rate at the 5’-end and G>A at the 3’-end of the sequencing read of 4% or above (characterized with mapDamage v2.0.8) (Jónsson et al., 2013); 3) library complexity estimates indicating that a minimum coverage of 0.5x would be achievable with further sequencing, 4) with a contamination level <= 5%.

Contamination rates were estimated with three methods: 1) damage pattern and polymorphism in mitochondrial DNA with Schmutzi (Renaud et al., 2015), 2) atypical ratios of coverages of X and Y chromosomes to autosomes calculated with ANGSD (Korneliussen et al., 2014), and 3) for male samples, high heterozygosity on non-pseudo-autosomal region of the X chromosome (chrX:5000000-154900000 in hg19) with the “contamination” tool in ANGSD (Korneliussen et al., 2014). If the contamination estimate for any of these three methods was above 5%, we considered the sample contaminated and excluded it from downstream analysis (n = 9).

For passing samples, processed data from all sequencing runs were merged into a single BAM file. The 204 new samples that passed quality filters have a median genome-wide depth of 0.92x (0.16x to 2.38x)

### Calling pseudohaploid genotypes

Pseudohaploid genotypes for study samples were called by randomly choosing one allele from each site where there was read coverage, following the approach and software provided by Stephan Schiffels (https://github.com/stschiff/sequenceTools). Variants were called for the 1240k SNP panel, which is commonly used for capture-based sequencing of ancient samples (Mathieson et al., 2015). For the newly reported samples, a median of 685,058 SNPs (167,000 to 1,029,345) were covered per sample. Data was output in eigenstrat format. This pipeline was also used to call genotypes for two published ancient DNA datasets which at the time were only available in BAM (sequence read) format (Clemente et al., 2021; Žegarac et al., 2021).

### Combining new genotypes with ancient and present-day published data

Newly processed pseudohaploid data was merged with several datasets. Most of the published data was retrieved from the Allen Ancient Data Resource (AADR) v44.3 (January 2021) (Allen Ancient DNA Resource, 2021; Mallick et al., 2023): a compilation of pseudohaploid and diploid genotypes for 5,225 ancient and 3,720 present-day individuals (1000 Genomes Project Consortium et al., 2015; Agranat-Tamir et al., 2020; Allentoft et al., 2015; Amorim et al., 2018; Antonio et al., 2019; Bergström et al., 2020; Biagini et al., 2019; Brace et al., 2019; Broushaki et al., 2016; Brunel et al., 2020; Cassidy et al., 2020, 2016; Damgaard et al., 2018; de Barros Damgaard et al., 2018; Ebenesersdóttir et al., 2018; Feldman et al., 2019a, 2019b; Fernandes et al., 2020, 2018; Fregel et al., 2018; Fu et al., 2016; Furtwängler et al., 2020; Gamba et al., 2014; Gokhman et al., 2020; González-Fortes et al., 2019, 2017; Günther et al., 2018, 2015; Haber et al., 2020, 2019, 2017; Harney et al., 2018; Hofmanová et al., 2016; Järve et al., 2019; Jeong et al., 2019; Jones et al., 2017, 2015; Keller et al., 2012; Kılınç et al., 2016; Krzewińska et al., 2018a, 2018b; Lamnidis et al., 2018; Lazaridis et al., 2017, 2016, 2014; Linderholm et al., 2020; Lipson et al., 2017; Mallick et al., 2016; Malmström et al., 2019; Marcus et al., 2020; Margaryan et al., 2020; Martiniano et al., 2017, 2016; Mathieson et al., 2018, 2015; Mittnik et al., 2019, 2018; Narasimhan et al., 2019; Nikitin et al., 2019; Olalde et al., 2019, 2018, 2015, 2014; Omrak et al., 2016; O’Sullivan et al., 2018; Patterson et al., 2012; Prüfer et al., 2017; Rivollat et al., 2020; Rodríguez-Varela et al., 2017; Saag et al., 2019, 2017; Sánchez-Quinto et al., 2019; Schiffels et al., 2016; Schroeder et al., 2019; Schuenemann et al., 2017; Sikora et al., 2017; Skoglund et al., 2014; Skourtanioti et al., 2020; Unterländer et al., 2017; Valdiosera et al., 2018; van den Brink et al., 2017; Veeramah et al., 2018; Villalba-Mouco et al., 2019; Wang et al., 2019; Zalloua et al., 2018). We also included relevant genetic data made available by authors that were not in the AADR: present-day genomes from the Balkans (Kovacevic et al., 2014), present-day genomes from 4 Poles, 3 Germans, and 2 Moldavians (Pagani et al., 2016), and Bronze Age Italian genomes (Saupe et al., 2021). Pseudohaploid genotypes for published Bronze Age Aegean genomes (Clemente et al., 2021) and Bronze Age Serbian genomes (Žegarac et al., 2021) were generated from BAM files using our pipeline. All published genomes were filtered for contamination based on reported contamination levels in the original study and SNP coverage based on the genomic data. All published samples that contributed to this study are listed in Appendix 5. To ensure maximum overlap with present-day and ancient samples in analyses, the merged dataset was subset to SNPs in the Human Origin Panel array, resulting in a total of 481,259 SNPs. For PCA and *qpAdm* modeling, SNPs that are transitions at CpG sites (n = 76,678) were excluded since they may have arisen from DNA damage as opposed to true genetic variation.

### Principal Component Analysis (PCA)

#### Setting up the principal component analysis

Principal component analysis was performed on genotypes from present-day and Mediterranean individuals using smartpca v16000 (https://github.com/chrchang/eigensoft/blob/master/POPGEN/README). The following parameters were used: 5 outlier iterations (numoutlieriter), 10 principal components along which to remove outliers (numoutlierevec), altnormstyle set to NO, with least squares projection turned on (lsqproject set to YES). To calculate principal components only using present-day individuals, a file (poplistname) was provided with the population names of present-day individuals, randomly subsampled per population. After outlier removal (which removed 55 samples), 829 individuals and 480,712 SNPs were used in the initial analysis. All individuals (non “reference” present-day genomes, and all of the ancient individuals) whose population was not listed in the poplistname file were projected onto the calculated principal components. In the paper, we refer to the individuals used in the calculation of principal components as belonging to the “reference PCA space”. These “reference” genomes were used to calculate the PCs because 1) they represent a wide range of present-day variation and 2) the genotypes tend to be of high quality.

#### Visual representation of PCA

In the figures, present-day genomes used in the reference space are generally colored gray in order to illustrate the background space of genetic variation. To reduce visual clutter and emphasize the ancient genomes, these present-day “reference” genomes are typically unlabeled. Labels for these populations are shown in Figure 2 - figure supplement 1.

### Calculation of F_ST_

To assess the extent of genetic differentiation across geographic space within a time period, we calculated the Fixation index (F_ST_) between groups of individuals on a sliding spatial grid. Each grid cell measured ten degrees longitude by ten degrees latitude, and was slid by one degree in both directions (north and east) nine times to build a total of ten spatial grids. For each of these grids, pairwise F_ST_ was calculated between all populated 10-by-10 grid cells using Hudson’s estimator, correcting for unequal sample size (Bhatia et al., 2013). In addition to F_ST_, we also calculated the average geographic distance (in kilometers) between all individuals across pairs of grid cells to assess how spatial distance relates to genetic differentiation. To visualize this relationship, we used lowess smoothing as implemented in python’s *statsmodels* package (*statsmodels.api.nonparametric.lowess*, v. 0.12.2). To infer confidence intervals for the lowess smoothing estimates, we devised a spatial bootstrapping procedure. Our bootstrap approach samples pairs of grid cells in a way that always samples all overlapping cells or none of them, so individuals are either fully included in a bootstrap replicate or not at all. This prevents double-counting individuals since they contribute to several comparisons across space due to the sliding grid.

### Modeling ancestry and identifying outliers using *qpAdm*

We used the *qpAdm* tool of *admixtools 2.0* to build a workflow that:

1. Identifies similar individuals within a region
2. Groups these into regional clusters
3. Compares these clusters both within and across regions

In this workflow, we heavily utilize what we call one-component models, where we test whether two individuals or clusters of individuals form a clade relative to a chosen set of reference populations (which we define below). To clarify what we mean by one-component model, assume that we have two focal individuals *i1* and *i2*, as well as a set of reference populations *ref*. Using the terminology of *admixtools 2* as well as Harney et al. (Harney et al., 2021), the following four tests are equivalent with respect to the resulting p-value:

- *qpWave(left = c(i1, i2), right = ref)*
- *qpAdm(left = c(i1, i2), target = NULL, right = ref)*
- *qpAdm(left = i1, target = i2, right = ref)*
- *qpAdm(left = i2, target = i1, right = ref)*

where we use the *R-style* notation of *c(i1, i2)* to denote a vector consisting of *i1* and *i2*. In all four cases, we are testing against the null hypothesis that the two individuals *do* form a clade, with low p-values indicating a rejection of that hypothesis. We will call any implementation of this test a “one-component *qpAdm* model”, but the equivalence stated above shows that our one-component models are equivalent to what was called a “qpWave analysis” by Fernandes et al. (Fernandes et al., 2020).

For our reference populations, we chose a set similar to those previously used to model Eurasian historical genomes (Fernandes et al., 2020). However, we added two Asian populations (Laos_Hoabinhian and Onge) based on evidence of gene flow from further east in a subset of our data. The purpose of a set of reference populations in the *qpAdm* modeling setting is to represent components of ancestry which are differentially related to the focal individuals being tested (i.e. “left”, or “sources” and “target”) in order to resolve differences in ancestry, but distally related enough to minimize the chance of recent gene flow between “left” and “right”. Our final set of references is: *Mbuti.DG (n = 4), WHG (n = 8), Russia_Ust_Ishim.DG (n = 1), CHG (n = 2), EHG (n = 3), Iran_GanjDareh_N (n = 8), Israel_Natufian_published (n = 3), Jordan_PPNB (n = 6), Laos_Hoabinhian (n = 1), Russia_EBA_Yamnaya_Samara (n = 9), Onge (n = 6), Spain_ElMiron (n = 1), Turkey_N_published (n = 8), Russia_MA1_HG (n = 1), Morocco_Iberomaurusian (n = 6), Czech_Vestonice16 (n = 1)*

#### Individual-based one-component models within regions

In the first step of our workflow, we perform one-component *qpAdm* tests between all pairs of individuals from the same region, from the Copper Age (inclusive) up to present-day (exclusive). As mentioned above, this approach tests against the null hypothesis that the two individuals *do* form a clade, with low p-values indicating a rejection of that hypothesis. Low *qpAdm* p-values thus suggest that the test individuals are not more closely related to each other than to one or more populations in the reference set. To convert the *qpAdm* p-value into a measure of dissimilarity (*d*), we calculate *d = −log_10_(p-value)*, where a large value of *d* indicates a low p-value, and thus a rejection of the null hypothesis of the two individuals forming a clade.

To cluster individuals into groups of genetically similar individuals, we performed hierarchical clustering on the dissimilarity matrix constructed from all pairwise values of *d* within a region. Hierarchical clustering was performed using the UPGMA algorithm as implemented in python’s *scipy.cluster.hierarchy* (v. 1.6.1). The hierarchical clustering was then split into flat clusters using a dissimilarity cutoff of *1.3*, which corresponds to a nominal p-value cutoff of *0.05*. Intermediate results from the pairwise analysis and clustering are shown for each region in Appendix 4.

#### Identifying ancestry outliers and their potential sources

Once clusters within regions were identified as described above, we classified them into two groups based on size. Clusters consisting of less than 5% of the total population or no more than two individuals in the region (across time) were classified as outlier candidates, whose ancestry is underrepresented in the region they were sampled. All other clusters were classified as majority clusters. Following this classification of clusters, we then split each cluster by time period (Copper Age, Bronze Age, Iron Age, Imperial Rome & Late Antiquity, Middle Ages & Early Modern), to end up with a *region_period_cluster* sub-classification. All downstream analyses were done using these *region_period* clusters.

For each cluster identified as an outlier candidate, we then tested all one-component models involving the candidate and a majority cluster **within** the same region. This was done to ensure that outlier candidates were truly distinct from majority ancestries identified in the region. If an outlier candidate could be connected to a majority cluster within its region through a valid one-component model (above a p-value threshold of *0.01*), it was removed from the outlier candidate list, as it represented a majority ancestry within the region.

For all outlier candidates that remained, we aimed to identify potential source ancestries that were majority ancestries in other regions. To do so, we tested all one-component models for a given outlier candidate and each of the majority clusters **across** all other regions. A majority cluster from a different region that had a valid one-component model with an outlier candidate was considered a potential source. To find the most likely source(s), we then subjected these potential source populations to model competition.

Model competition involves re-testing the model fit of each potential source after adding another potential source to the right group, first described in (Lazaridis et al., 2016), and more recently detailed in (Harney et al., 2021). The idea is that if a population in the right group has significantly more allele sharing with the target than the source, then the model will be rejected. (This is why right group populations are chosen to be distal, yet relevant, to target and source). We use this property to our advantage by rotating all n-1valid sources for the target through the right group set, one at a time, for the same target and a source *x* (the one source that is not included in the rotation). If the previously valid source *x* is rejected when including another valid source *y* in the right group then we remove source *x* from the list of potential sources. Note that this does not make source *y* the best source, only a better one than *x.* Thus, this scheme only eliminates sub-optimal sources, rather than selecting a best source.

If there were still multiple valid sources following the model competition scheme described above, we prioritized a candidate source that is from the same time period as the target cluster, or from a previous period going backwards in time. If there were still multiple candidate sources, they were considered equivalent and all kept in downstream analyses.

Among the outliers identified, we did not find a significant sex bias compared to non-outliers. Overall, there are more males than females in the dataset. However, the proportions of males in non-outliers, outliers with source, and outliers without source do not differ significantly by a Chi-squared test (p-value = 0.4117, df = 2) (Figure 5 - figure supplement 1). When outliers (with and without source) are treated as one group, there is still no significant association with outlier status and sex (p-value = 0.633, df = 1).

#### Admixture modeling with *qpAdm*

For targeted analyses of other clusters beyond just outlier candidates, for example to annotate Figures 2, 3 and 4, we also used cluster-based *qpAdm*. In addition to one-component models as described above, we also used two-component models of admixture. These models test the hypothesis that a focal target cluster can be modeled as a two-way admixture of two sources (or “left” populations). As above, a p-value below the threshold rejects this hypothesis, i.e. the proposed admixture model is not a good fit and a different model needs to be considered to disentangle the admixture scenario in question.

### Simulations

To assess how spatial population structure would be impacted by different modes of dispersal, we set up forward simulations in continuous 2D space using SLiM v. 3.6 (Haller and Messer, 2019). The aim of these simulations was to approximate the extent of spatial population structure we observe by the beginning of the Iron Age in Western Eurasia, after the major prehistoric migrations had taken place. To achieve that, we decided not to attempt simulating the precise ancestry composition of populations in different regions at that time, but rather to simulate simply the extent of spatial structure as measured by the relationship of population differentiation (F_ST_) and geographic distance. We chose the SLiM simulation framework to make use of its extensive feature set to simulate individuals in continuous space. We simulated diploid genomes made up of a single, 10^8^ bp long chromosome, with recombination rate and mutation rate set to 10^-8^. We used the default Wright-Fisher simulation mode, where a single population of constant size *N* is simulated with non-overlapping populations, i.e. each generation is made up of offspring generated from the previous generation. Spatial structure is established by associating each individual with a continuous 2D coordinate (i.e. latitude and longitude), and by using these coordinates to govern three demographic processes: mate choice, competition, and dispersal. An overview of how these processes can be set up to interact in SLiM can be found in *Recipe 15.4* of SLiM v. 3.6 (see e.g. here: https://github.com/MesserLab/SLiM/tree/v3.6/SLiMgui/Recipes). Briefly, for mate choice, a Gaussian interaction function with *maxDistance = 0.1, maxStrength = 1.0, sigma = 0.02* is used to govern a *mateChoice* callback using the *strength* of that interaction function. For competition, another Gaussian interaction function with *maxDistance = 0.3, maxStrength = 3.0, sigma = 0.1* is used to calculate competition using the *totalNeighborStrength* vector of that interaction function to scale an individual’s relative fitness as *1.1 - competition / N*. Finally, we establish local dispersal through a *modifyChild* callback, where a newly generated offspring’s position is drawn from a Gaussian centered at the location of the maternal individual with standard deviation *sigmaDisp*.

We let this population evolve forward in time for 2000 + 120 generations during which the processes outlined above lead to spatial population structure, where individuals sampled closely together in 2D space are also more closely related genetically. We do not simulate mutations in SLiM, as this poses a major computational burden. Instead, we use tree sequence recording (Haller et al., 2019; Kelleher et al., 2018) to track the full genealogy of all individuals in the simulation which are either alive at the end of the simulation, or explicitly sampled through time using the *treeSeqRememberIndividuals* function of SLiM. While 2,000 generations are enough to establish spatial structure under the parameters we consider, it is by far insufficient for all sampled individuals to fully coalesce. To accurately assess neutral variation however, we need all sampled individuals to have a common ancestor at some point in the past, as mutations may have arisen at any point leading back to this ultimate coalescence event. Therefore, we approximate the deep history of our population with a panmictic population simulated backwards in time using the coalescent with recombination as implemented in msprime (Kelleher et al., 2016). This process has been referred to as “recapitation” (Haller et al., 2019), where an incomplete genealogy with multiple roots (from SLiM) is “recapitated” using coalescent simulation backwards in time. This is made possible by using the tree sequence data structure to record and simulate genealogies in both SLiM and msprime. Since our simulation is only concerned with how processes such as dispersal affect neutral variation across space and through time, we can use the “recapitated” tree sequence to overlay mutations onto the full genealogy of all sampled individuals, also using msprime. The rationale here is that under neutrality, mutations will not affect the structure of the genealogy, so we can simulate the genealogy without mutations first, and overlay neutral mutations second, thereby greatly reducing computational burden. We then extracted the resulting genotypes of all individuals from the tree sequence for downstream analysis. We only kept sites segregating in individuals at the end of the simulation (“present day”), and filtered for minor allele frequency of at least 0.01 across the entire dataset, to make downstream analysis of simulated genomes comparable to how the empirical data was ascertained and analyzed.

To assess the relationship between F_ST_ and spatial distance, we split geographic space into a 10-by-10 grid and calculated all pairwise F_ST_ between inhabited grid cells using Hudson’s estimator with unequal sample size correction (Bhatia et al., 2013), as well as the average geographic (euclidean) distance between individuals across grid cells. We used this F_ST_ analysis to calibrate the base dispersal *sigmaDisp* as well as the population size *N*, so that F_ST_ at maximum distance (F_ST_max) would approximately match the F_ST_max we observed at the start of the historical period (∼0.03). We used grid search with a range of *sigmaDisp* and *N* values, and found the parameter pair *N = 50,000* & *sigmaDisp = 0.02* to qualitatively produce the closest match (Figure 7 - figure supplement 1). We use this parameter set as our base model of population structure without long-range dispersal, where we allow spatial structure to establish over 2,000 generations, and then observe the F_ST_ - Distance relationship over the following 120 generations for a total of 2,120 generations simulated in SLiM (Figure 7A).

Given this base model of spatial population structure, we can now start to introduce long-range dispersing individuals. We do this by allowing a specified fraction of individuals to use a higher *sigmaDispLR* than the *sigmaDisp* used by the rest of the population for the final 120 generations of the simulation, approximately matching the 3,000 years since the beginning of the historical period assuming a generation time of 25 years. Since the long-range dispersal is also drawn from a Gaussian, the distribution of dispersal distances will have substantial overlap with the distances produced by base dispersal. To make the fraction of long-range dispersal accurately represent the fraction of individuals that actually disperse longer distances, we thus require a long-range dispersing individual to disperse to a location outside of the 99th percentile of density covered by the short-range base dispersal.

We aimed to choose a *sigmaDispLR* that approximately matches the empirical distribution of long-range dispersing individuals we observe in the analysis displayed in Figure 4. Since the euclidean distances in the simulation are on a different scale than the geodesic distances observed in the data, we aimed to match qualitatively the relationship of long-range dispersal distances to random distances that could be observed if two populated locations were drawn at random. We visually analyzed this relationship from the data, and then performed a search across a range of possible *sigmaDistLR* to find a qualitative match. This led us to choose a value of *sigmaDistLR =*

*0.20* (Figure 7 - figure supplement 2).

Finally, we analyzed simulated “present-day” genomes (i.e. after 2120 generations of SLiM) using PCA. We used the *sklearn.decomposition.PCA* module (scikit-learn v. 0.24.2) with the *svd_solver == ‘arpack’* option to run non-probabilistic PCA to calculate the first 10 principal components. Similarly to how the empirical data was analyzed with *smartpca*, we also did 5 rounds of iterative outlier removal, removing individuals from the PCA that deviated by more than six standard deviations along any of the 10 principal components. The number of variants contributing to these PCA were 624,617, 625,669 and 626,052 for Figure 7 A, B and C respectively, and thus comparable to the number of variants contributing to our data analysis.

### Appendix Files

Appendix 1 (PDF) Archaeological context for sampling locations Appendix 2 (Table) Metadata for all newly reported individuals Appendix 3 (Table) AMS and calibration results Appendix 4 (PDF) Visualization of per-region qpAdm clustering Appendix 5 (Table) Published samples that contributed to this study

## Data Availability

All sequence data newly generated for this study are available at the European Nucleotide Archive (ENA) database. Raw sequencing data is available under the accession number PRJEB53565. Sequences mapped to the human reference genome are available under accession number PRJEB53564. Sequences previously reported by (Moots et al., 2022) are available under accession number PRJEB49419. All published data used in this study is listed in Appendix 5, and can be retrieved from primary sources (see Methods, “Combining new genotypes with ancient and present-day published data”) or from the Allen Ancient Data Resource (Mallick et al., 2023).

## Ethics

This study follows ethics guidelines adopted by the ancient DNA field (Alpaslan-Roodenberg et al., 2021). A clear plan of research was laid out before the collection of samples, leading us to focus on sampling from under-sampled historical regions in Eurasia and therefore minimize unnecessary destruction of human remains. The research intent for these samples was clearly communicated to caretakers of the samples prior to collection. Local anthropologists, archaeologists, and museum directors from each geographic region were involved in the sample acquisition, extraction from skeletal material, and interpretation of genetic results. The genetic findings regarding individual samples from each region were communicated to local collaborators, all of whom were included as co-authors on the paper and were supportive of the final results. The involvement of our local collaborators was essential for the interpretation of the genetic results through their input on the historical and archaeological characterization of the specimens. We have supported our local collaborators with immediate access to the raw genetic data, and by communicating results in written and oral forums. Authorities responsible for all archaeological sites provided written documentation for their specimens to be included in this study through collaboration with the Pinasi Lab (Vienna, Austria).

## Supporting information

Appendix 1 - Archaeological site information

Appendix 2 - Metadata of newly reported genomes

Appendix 3 - AMS dating results

Appendix 4 - Clustering results

Appendix 5 - Samples included in this study

## Acknowledgements

We would like to thank Professor Walter Scheidel for helpful discussions and feedback on the historical context, and all members of the Pritchard and Pinhasi labs for their valuable input. We thank Pieter W. Faber and the University of Chicago Genomics Facility for sequencing the samples reported here. We thank Benjamin Peter and an anonymous reviewer for their insightful and constructive reviews.

This project was partially supported by a National Science Foundation Graduate Research Fellowship (M.L.A.), a grant from the National Institutes of Health RO1 HG011432 (C.L.W.), the Austrian Science Fund (FWF) M3108-G (S.S.), and the Howard Hughes Medical Institute (J.K.P.).

---

**Figure 1 - figure supplement 1.**
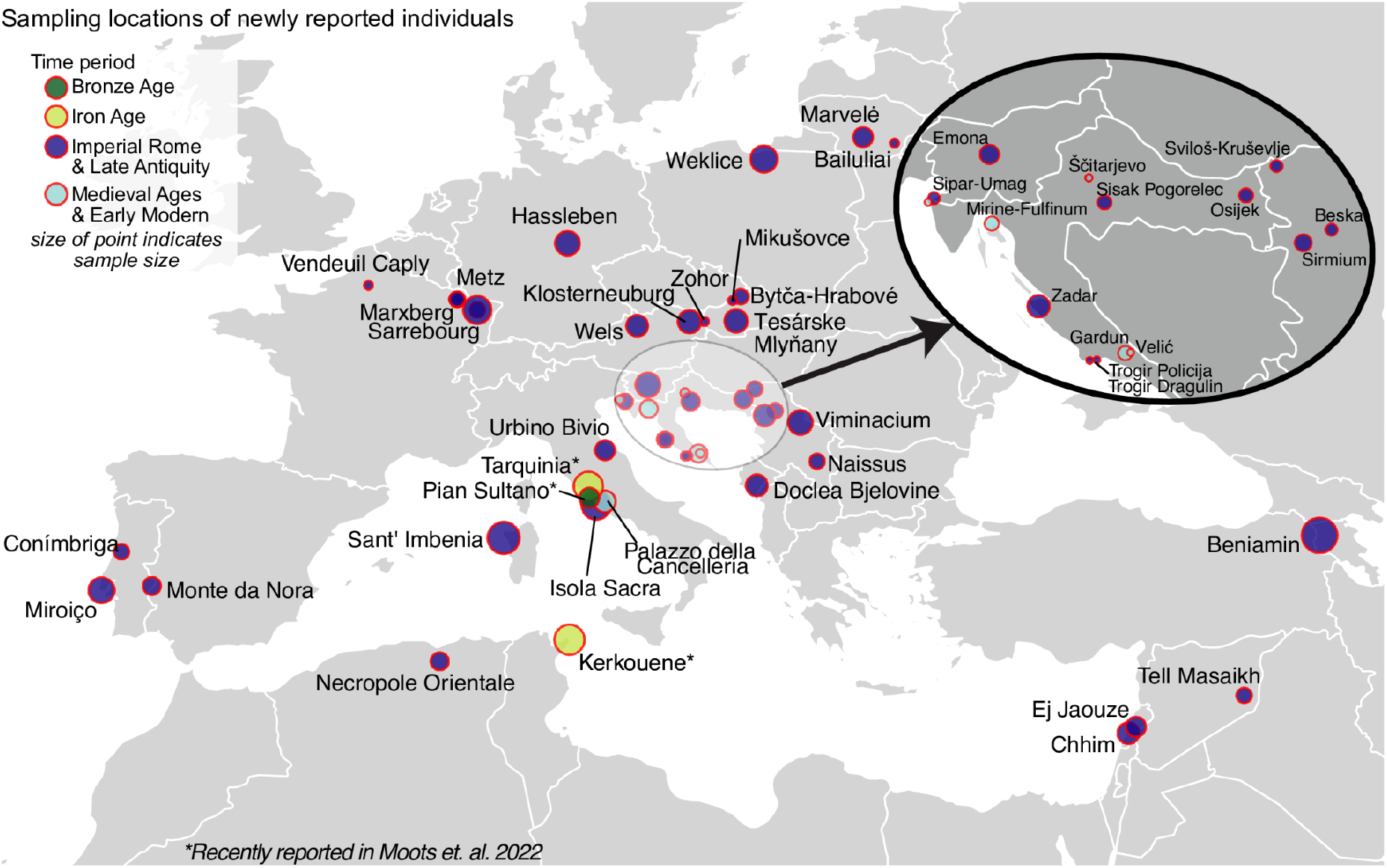
Detailed map of locations for newly reported samples. Each circle represents a location, the size of the circle corresponds to the number of individuals sampled from that location. Circles are colored by their time period: Bronze Age is green (Pian Sultano), Iron Age is yellow (two recently reported sites Tarquinia and Kerkouane), Imperial Rome and Late Antiquity is dark blue, Medieval Ages and Early Modern are light blue (Palazzo della Cancelleria, Velić, Gardun, Mirine-Fulfinum). Note that the Bronze Age and Iron Age sites were recently reported in (Moots et al., 2022).

**Figure 2 - figure supplement 1.**
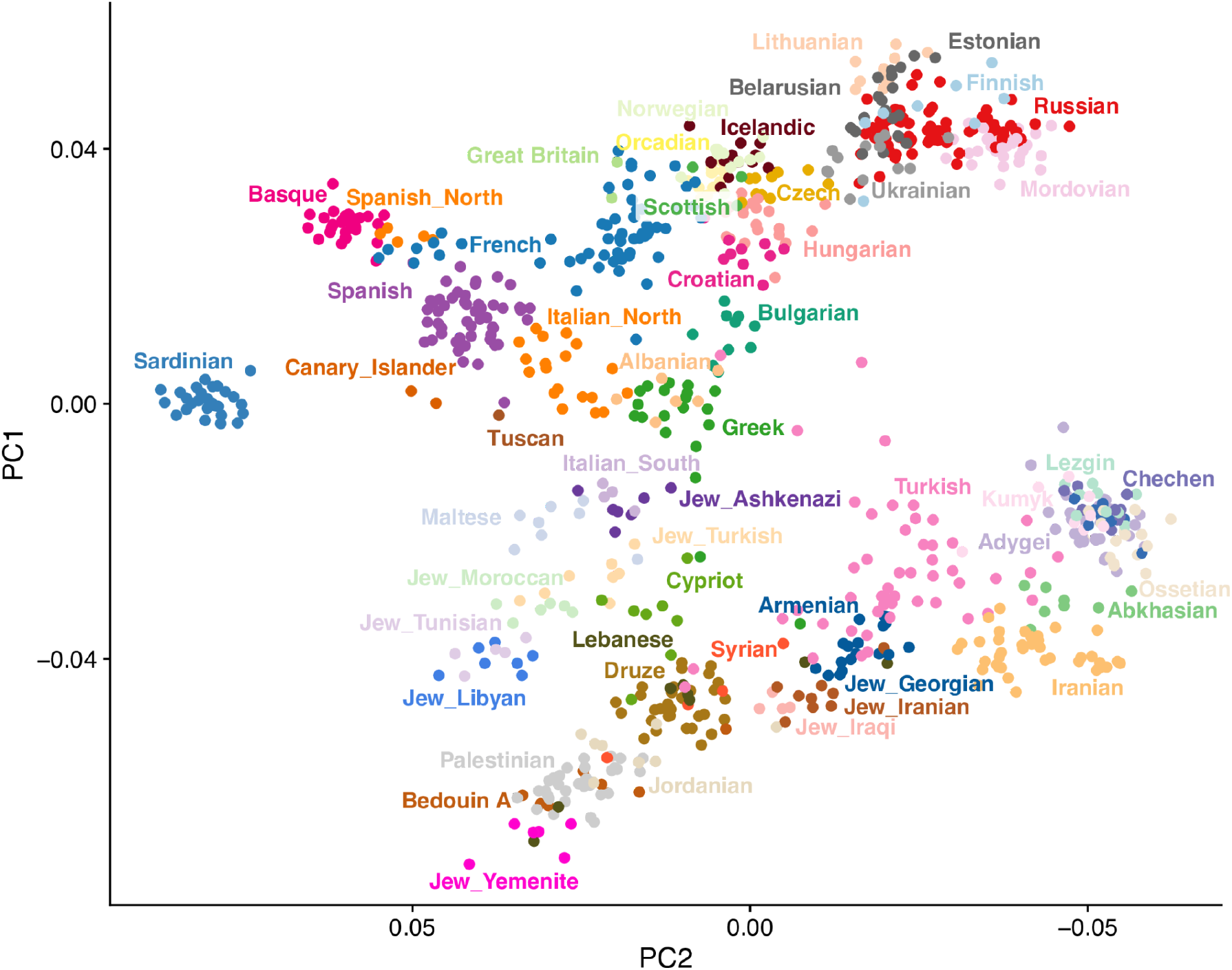
Principal component analysis of present-day genomes from Europe and the Mediterranean. PCA was performed on 829 individuals (480,712 snps) using *smartpca v1600*. The following parameters were used: 5 outlier iterations (numoutlieriter), 10 principal components along which to remove outliers (numoutlierevec), altnormstyle set to NO, with least squares projection turned on (lsqproject set to YES).

**Figure 2 - figure supplement 2.**
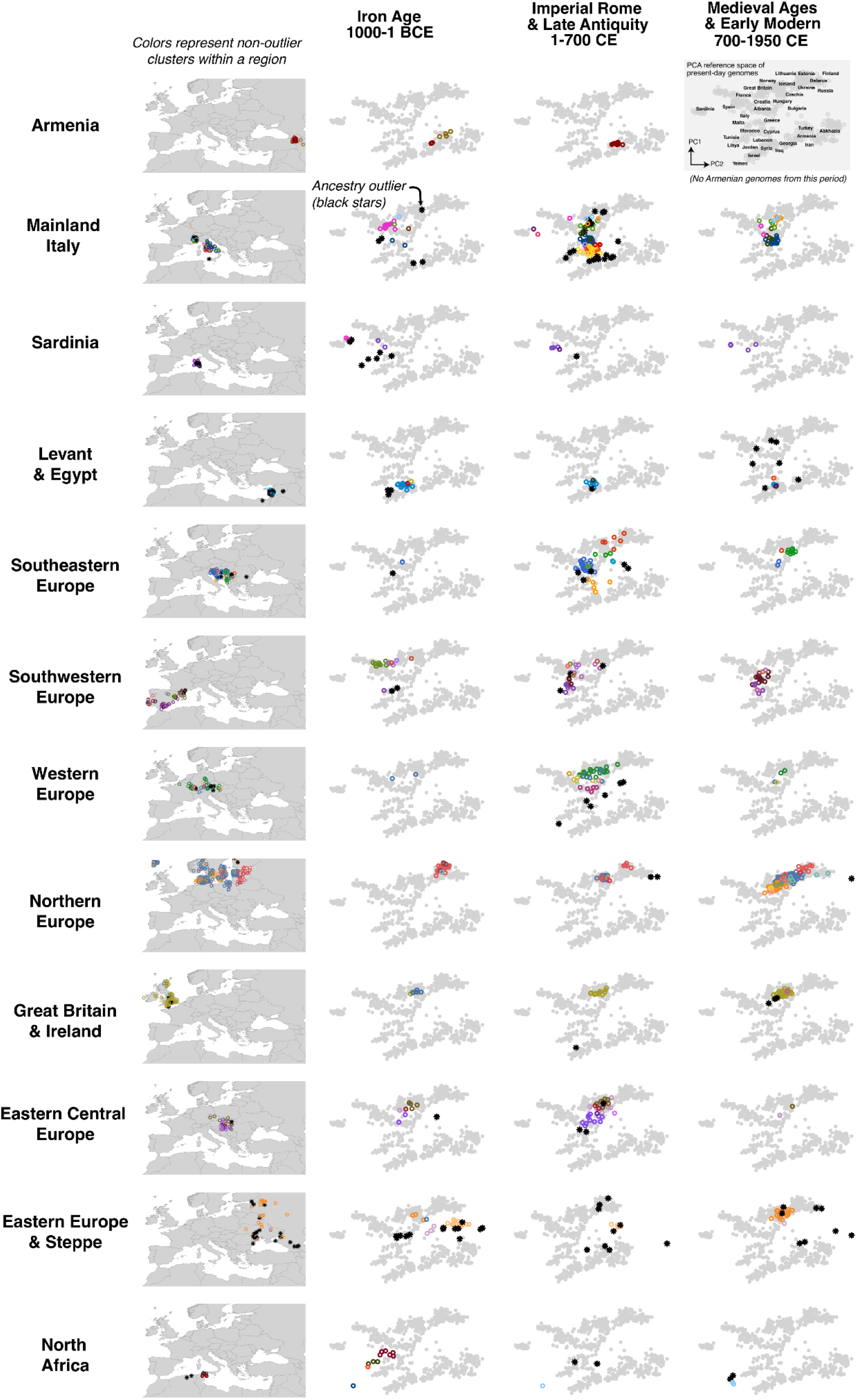
Ancestry clusters identified within regions. Each row displays data from a single study region. The first column shows a map with the sampling locations for the individuals, while columns two through four show the individuals projected onto a PCA space of present-day genomes (gray points) (populations are labeled in the far right panel in row 1 and in Figure 2 - figure supplement 1). Individual ancient genomes in the map and PCA panels are colored by ancestry clusters identified using qpAdm. Colors are not matched across regions. Star points are putative outliers, i.e. individuals with ancestry that is underrepresented in the region. They are not colored by ancestry clusters so as to reduce visual clutter.

**Figure 2 - figure supplement 3.**
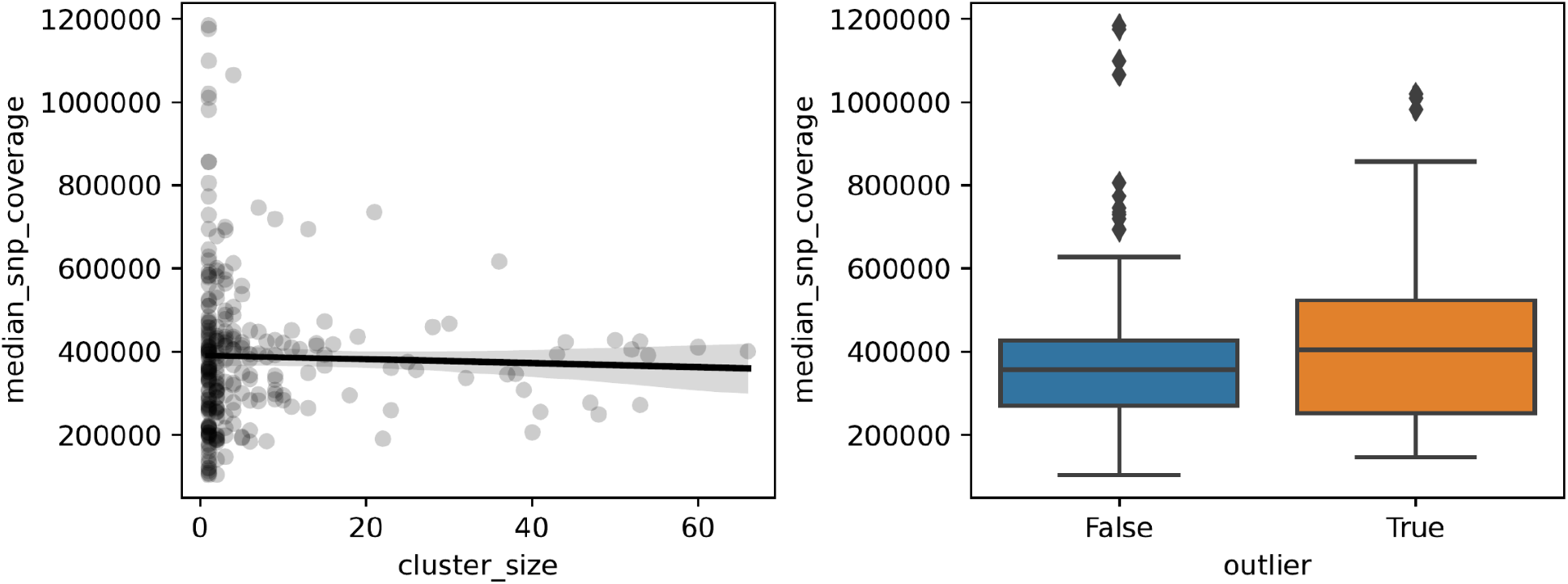
SNP coverage comparison across cluster sizes and downstream outlier status. (left) No significant correlation was detected between the median number of SNPs covered across the individuals in a cluster and cluster size. (right) There also was no significant difference in the number of SNPs covered between outlier and non-outlier clusters.

**Figure 3 - figure supplement 1.**
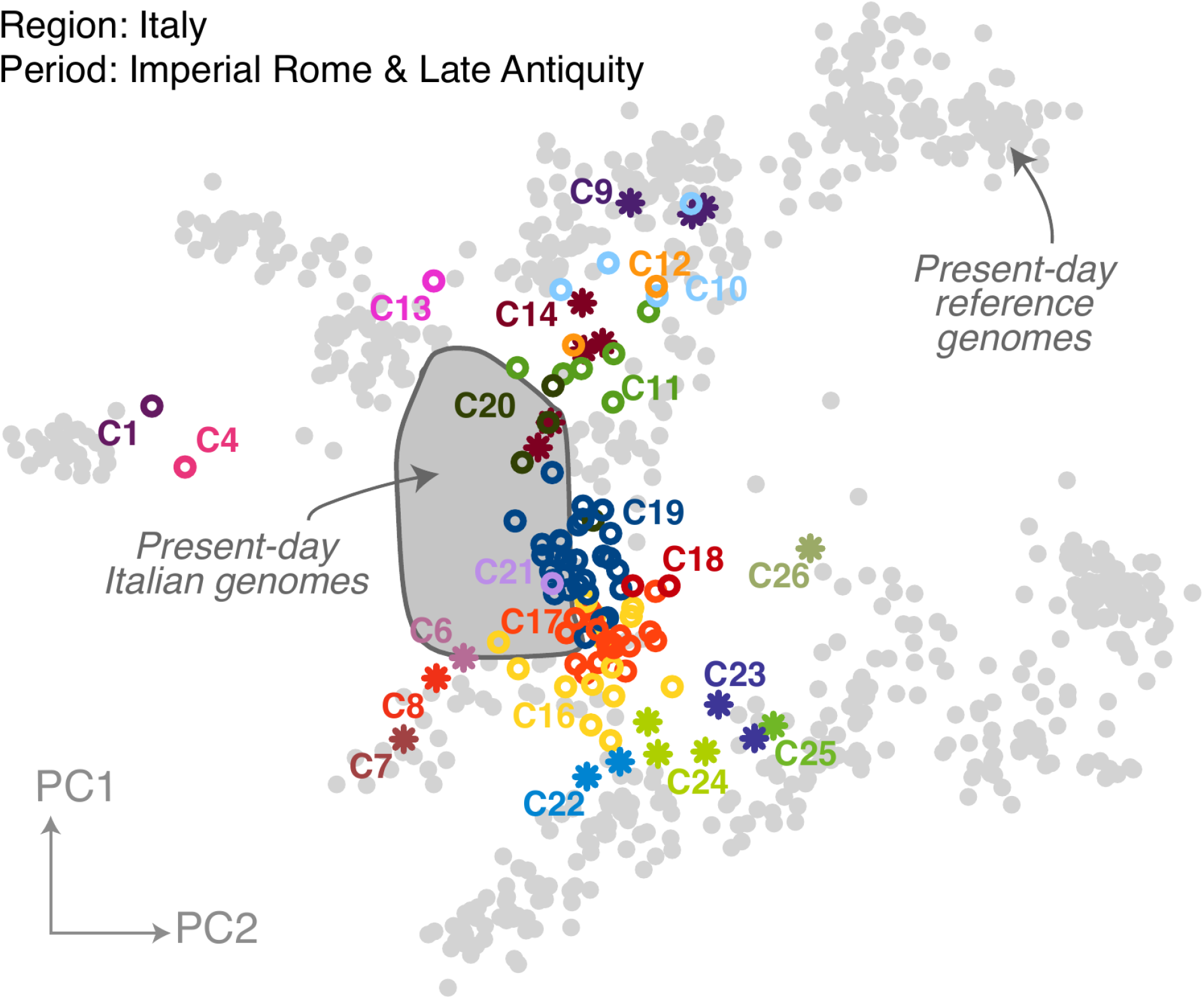
Population structure of Italy during the Imperial Roman and Late Antiquity period. Ancient Italian genomes (colored points) from the Imperial Roman and Late Antiquity period were projected onto principal components of present-day genomes (gray points, populations labeled in Figure 2 - figure supplement 1). Present-day Italian genomes are highlighted by a gray filled ellipse. Star points are outliers and circle points are non-outliers. Outlier clusters that can be modeled using contemporaneous populations are labeled with the potential source region.

**Figure 5 - figure supplement 1.**
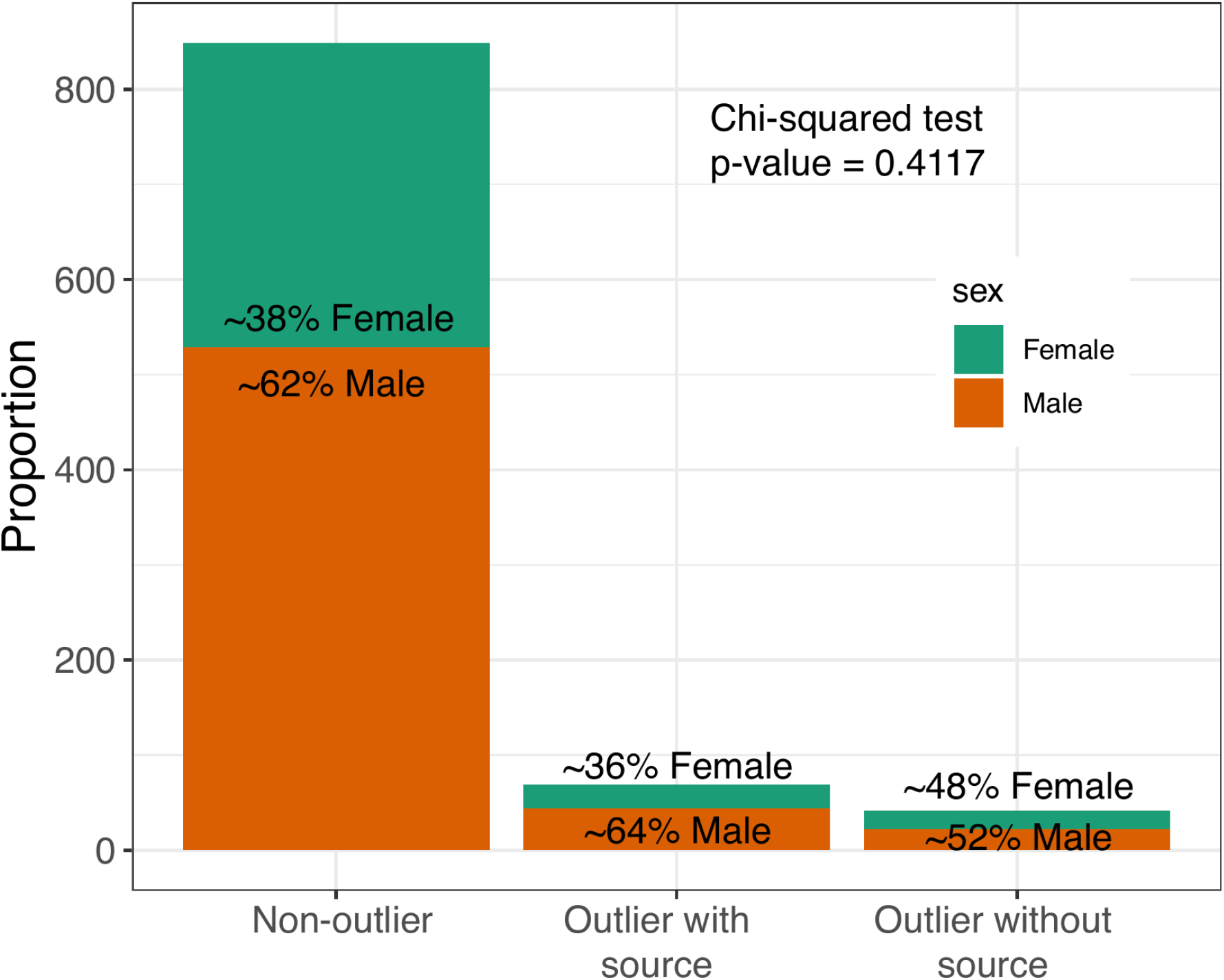
Lack of sex-bias amongst outliers with valid qpAdm sources. The proportions of males and females do not differ significantly between outlier and non-outlier groups (p = 0.4117). When outliers (with and without source) are treated as one group, there is still no significant association with outlier status and sex (p = 0.633).

**Figure 5 - figure supplement 2.**
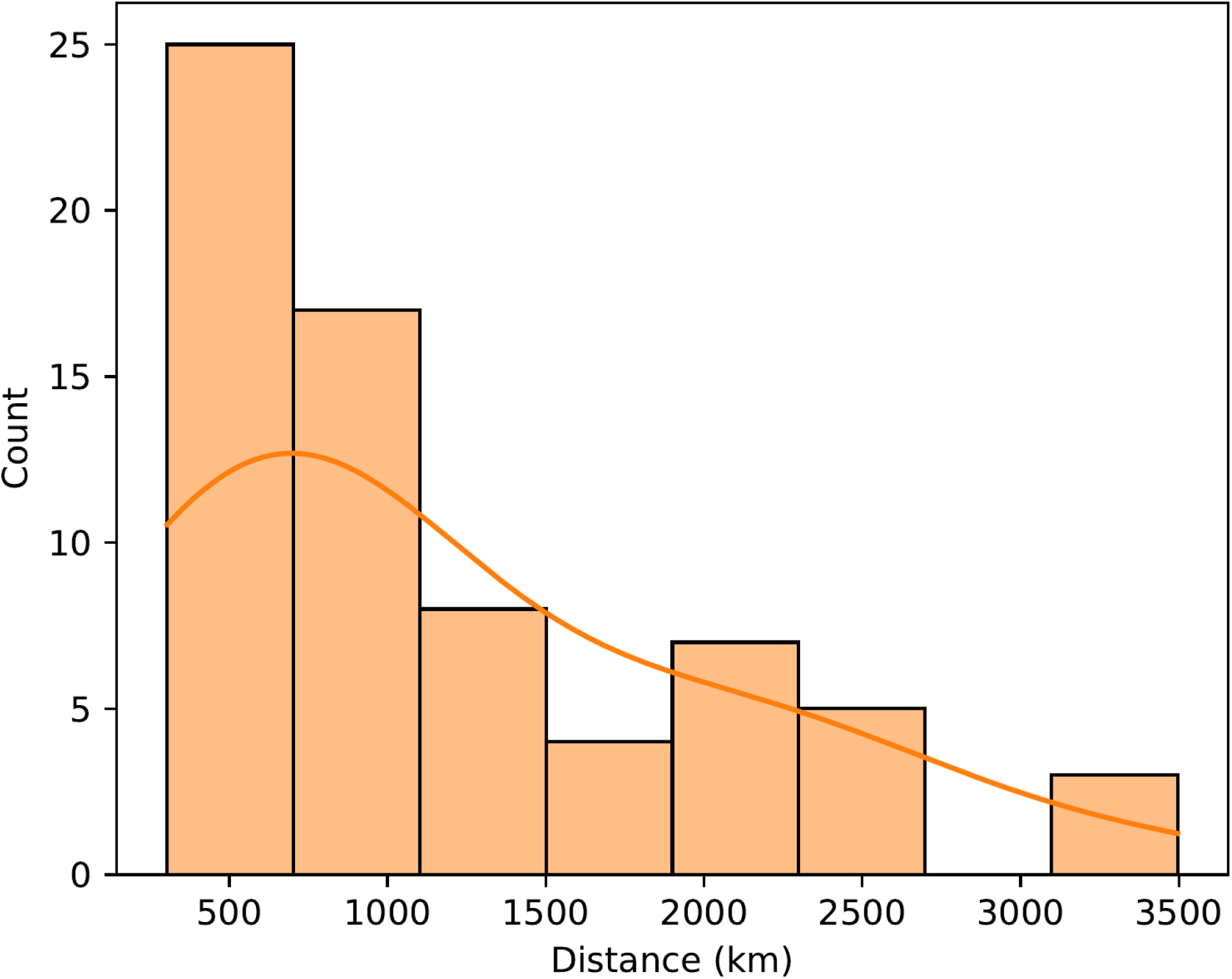
Distances of outliers to their candidate sources. Geographic distance between the sampling locations of “outlier with source” and the location of their putative source was calculated for each outlier. The mean distance was calculated if there were multiple putative sources.

**Figure 5 - figure supplement 3.**
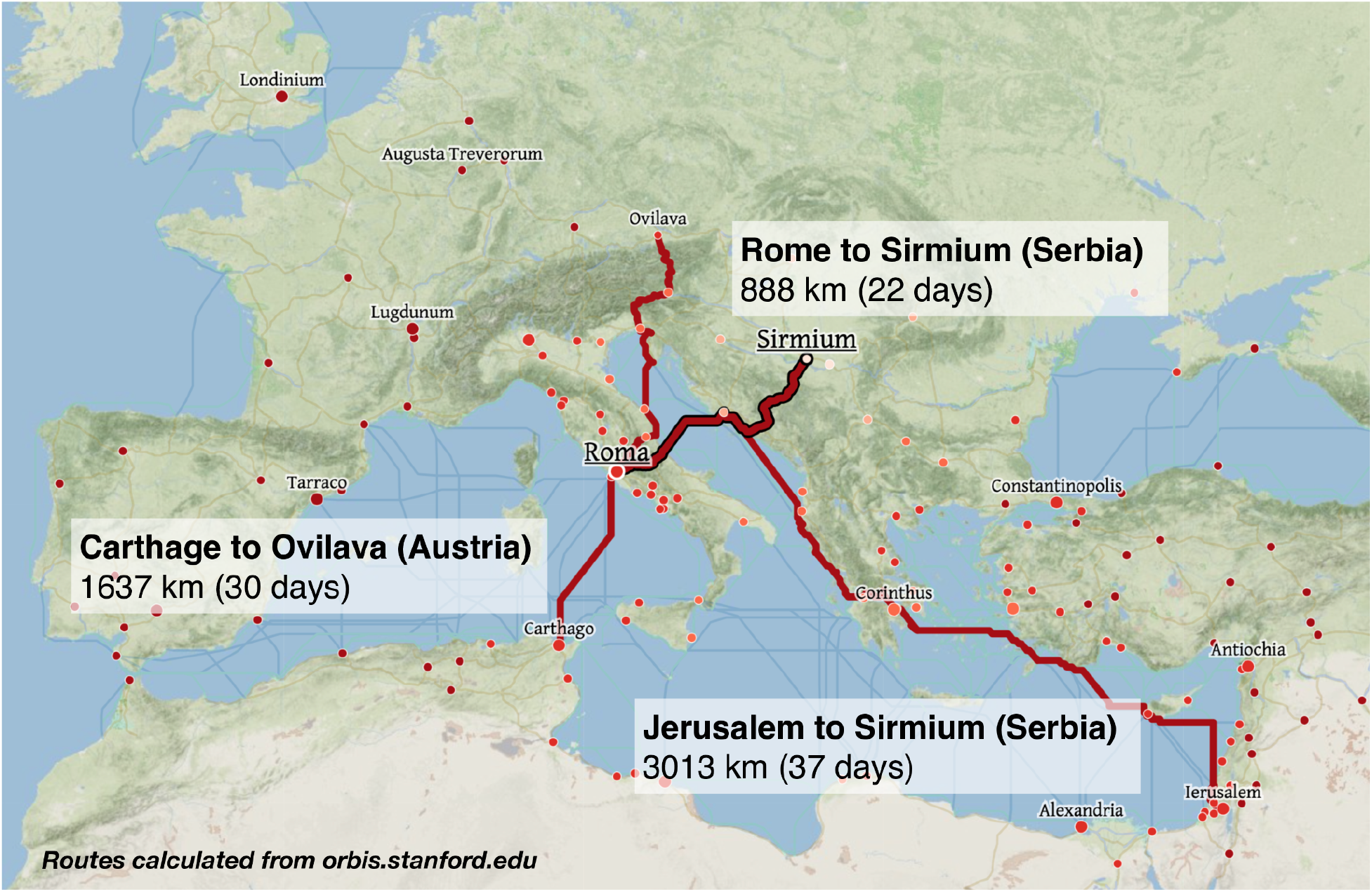
Example routes and travel times across the Roman Empire. Routes and travel times were approximated using orbis.stanford.edu, a geospatial network model of the Roman Empire. Routes shown are the fastest routes during Summer for civilians, utilizing road, river, coastal sea, and open sea, and by foot if on road. Routes for military individuals (not shown) are marginally faster.

**Figure 7 - figure supplement 1.**
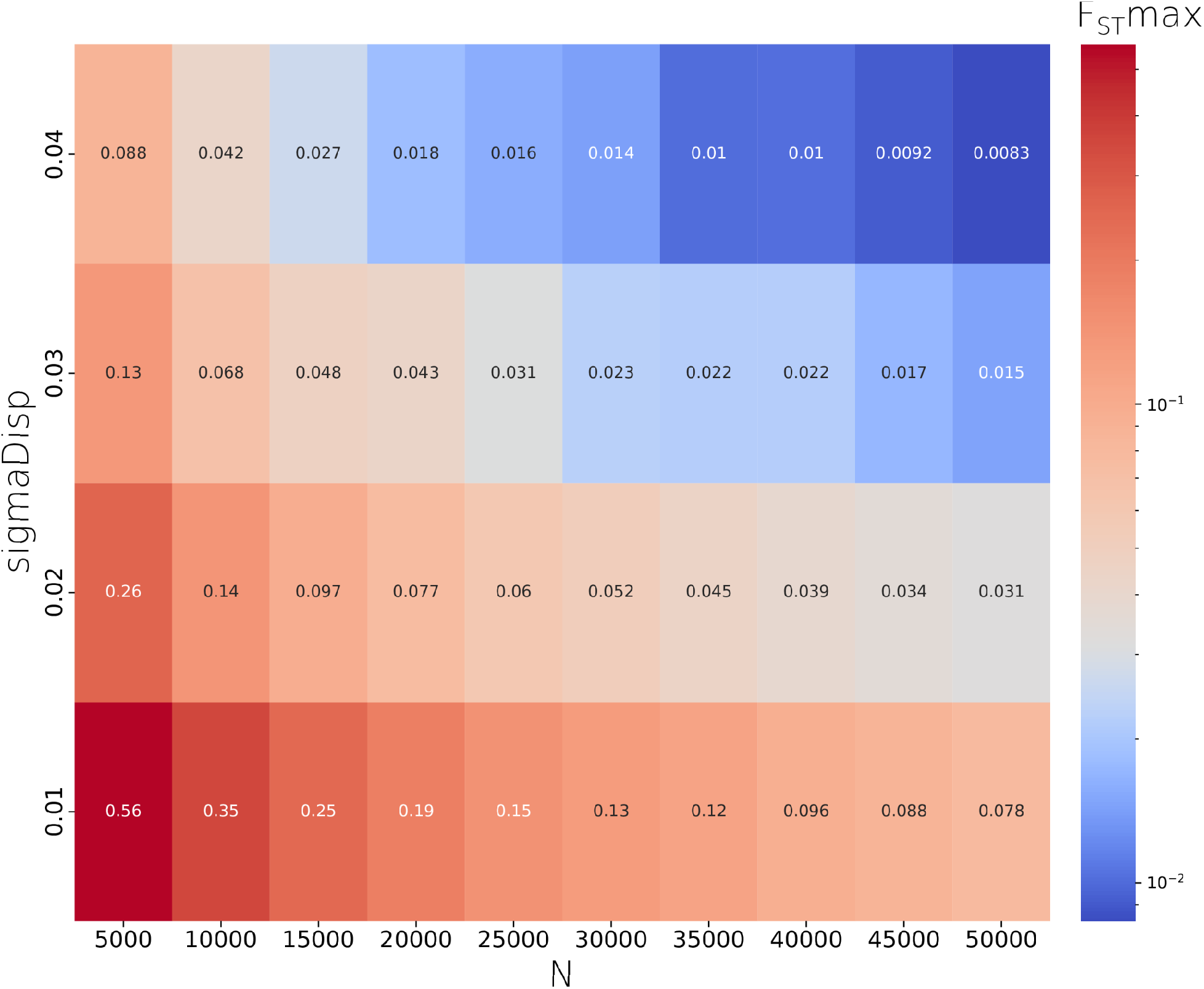
A *sigmaDisp - N* parameter pair was chosen to closely approximate the observed *F_ST_max* of ∼0.03 using grid search across a range of parameter pairs. We used the pair *N = 50,000* & *sigmaDisp = 0.02* for all other simulations we report.

**Figure 7 - figure supplement 2.**
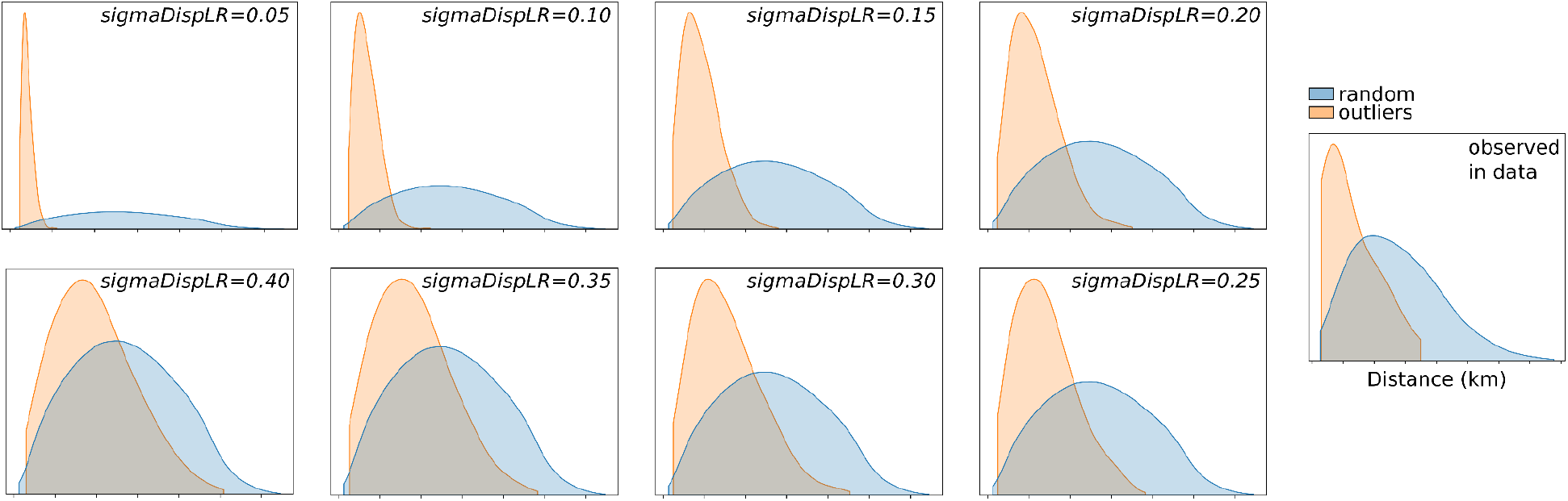
A *sigmaDispLR* parameter was chosen to qualitatively resemble long-range dispersal distances observed in the data, by comparing the distribution of distances under long-range dispersal (*outliers*) to randomly chosen distances given the spatial distribution of samples. We used a value of *0.20* for all other simulations we report.

## References

1. 1000 Genomes Project Consortium, Auton A, Brooks LD, Durbin RM, Garrison EP, Kang HM, Korbel JO, Marchini JL, McCarthy S, McVean GA, Abecasis GR. 2015. A global reference for human genetic variation. Nature 526:68–74.

2. Abulafia D. 2011. The Great Sea: A Human History of the Mediterranean. Oxford University Press.

3. Agranat-Tamir L, Waldman S, Martin MAS, Gokhman D, Mishol N, Eshel T, Cheronet O, Rohland N, Mallick S, Adamski N, Lawson AM, Mah M, Michel M, Oppenheimer J, Stewardson K, Candilio F, Keating D, Gamarra B, Tzur S, Novak M, Kalisher R, Bechar S, Eshed V, Kennett DJ, Faerman M, Yahalom-Mack N, Monge JM, Govrin Y, Erel Y, Yakir B, Pinhasi R, Carmi S, Finkelstein I, Carmel L, Reich D. 2020. The Genomic History of the Bronze Age Southern Levant. Cell 181:1146–1157.e11.

4. Allen Ancient DNA Resource. 2021. Allen Ancient DNA Resource (AADR): Downloadable genotypes of present-day and ancient DNA data. https://reich.hms.harvard.edu/allen-ancient-dna-resource-aadr-downloadable-genotypes-present-day-and-ancient-dna-data

5. Allentoft ME, Sikora M, Sjogren KG, Rasmussen S, Rasmussen M, Stenderup J, Damgaard PB, Schroeder H, Ahlstrom T, Vinner L, Malaspinas AS, Margaryan A, Higham T, Chivall D, Lynnerup N, Harvig L, Baron J, Della Casa P, Dabrowski P, Duffy PR, Ebel AV, Epimakhov A, Frei K, Furmanek M, Gralak T, Gromov A, Gronkiewicz S, Grupe G, Hajdu T, Jarysz R, Khartanovich V, Khokhlov A, Kiss V, Kolar J, Kriiska A, Lasak I, Longhi C, McGlynn G, Merkevicius A, Merkyte I, Metspalu M, Mkrtchyan R, Moiseyev V, Paja L, Palfi G, Pokutta D, Pospieszny L, Price TD, Saag L, Sablin M, Shishlina N, Smrcka V, Soenov VI, Szeverenyi V, Toth G, Trifanova SV, Varul L, Vicze M, Yepiskoposyan L, Zhitenev V, Orlando L, Sicheritz-Ponten T, Brunak S, Nielsen R, Kristiansen K, Willerslev E. 2015. Population genomics of Bronze Age Eurasia. Nature 522:167–172.

6. Alpaslan-Roodenberg S, Anthony D, Babiker H, Bánffy E, Booth T, Capone P, Deshpande-Mukherjee A, Eisenmann S, Fehren-Schmitz L, Frachetti M, Fujita R, Frieman CJ, Fu Q, Gibbon V, Haak W, Hajdinjak M, Hofmann KP, Holguin B, Inomata T, Kanzawa-Kiriyama H, Keegan W, Kelso J, Krause J, Kumaresan G, Kusimba C, Kusimba S, Lalueza-Fox C, Llamas B, MacEachern S, Mallick S, Matsumura H, Morales-Arce AY, Matuzeviciute GM, Mushrif-Tripathy V, Nakatsuka N, Nores R, Ogola C, Okumura M, Patterson N, Pinhasi R, Prasad SPR, Prendergast ME, Punzo JL, Reich D, Sawafuji R, Sawchuk E, Schiffels S, Sedig J, Shnaider S, Sirak K, Skoglund P, Slon V, Snow M, Soressi M, Spriggs M, Stockhammer PW, Szécsényi-Nagy A, Thangaraj K, Tiesler V, Tobler R, Wang C-C, Warinner C, Yasawardene S, Zahir M. 2021. Ethics of DNA research on human remains: five globally applicable guidelines. Nature 599:41–46.

7. Amorim CEG, Vai S, Posth C, Modi A, Koncz I, Hakenbeck S, La Rocca MC, Mende B, Bobo D, Pohl W, Baricco LP, Bedini E, Francalacci P, Giostra C, Vida T, Winger D, von Freeden U, Ghirotto S, Lari M, Barbujani G, Krause J, Caramelli D, Geary PJ, Veeramah KR. 2018. Understanding 6th-century barbarian social organization and migration through paleogenomics. NatCommun 9:354–354.

8. Antonio ML, Gao Z, Moots HM, Lucci M, Candilio F, Sawyer S, Oberreiter V, Calderon D, Devitofranceschi K, Aikens RC, Aneli S, Bartoli F, Bedini A, Cheronet O, Cotter DJ, Fernandes DM, Gasperetti G, Grifoni R, Guidi A, La Pastina F, Loreti E, Manacorda D, Matullo G, Morretta S, Nava A, Fiocchi Nicolai V, Nomi F, Pavolini C, Pentiricci M, Pergola P, Piranomonte M, Schmidt R, Spinola G, Sperduti A, Rubini M, Bondioli L, Coppa A, Pinhasi R, Pritchard JK. 2019. Ancient Rome: A genetic crossroads of Europe and the Mediterranean. Science 366:708–714.

9. Beard M. 2015. SPQR: a history of ancient Rome, First. ed. New York: Liveright Publishing Corporation, a Division of W.W. Norton & Company.

10. Bergström A, McCarthy SA, Hui R, Almarri MA, Ayub Q, Danecek P, Chen Y, Felkel S, Hallast P, Kamm J, Blanché H, Deleuze J-F, Cann H, Mallick S, Reich D, Sandhu MS, Skoglund P, Scally A, Xue Y, Durbin R, Tyler-Smith C. 2020. Insights into human genetic variation and population history from 929 diverse genomes. Science 367. doi:10.1126/science.aay5012

11. Bhatia G, Patterson N, Sankararaman S, Price AL. 2013. Estimating and interpreting FST: the impact of rare variants. Genome Res 23:1514–1521.

12. Biagini SA, Solé-Morata N, Matisoo-Smith E, Zalloua P, Comas D, Calafell F. 2019. People from Ibiza: an unexpected isolate in the Western Mediterranean. Eur J Hum Genet 27:941–951.

13. Bowes K. 2011. Rural Poverty in the Roman Empire. Accessed February 22:2021.

13a. Brace S, Diekmann Y, Booth TJ, van Dorp L, Faltyskova Z, Rohland N, Mallick S, Olalde I, Ferry M, Michel M, Oppenheimer J, Broomandkhoshbacht N, Stewardson K, Martiniano R, Walsh S, Kayser M, Charlton S, Hellenthal G, Armit I, Schulting R, Craig OE, Sheridan A, Parker Pearson M, Stringer C, Reich D, Thomas MG, Barnes I. 2019. Ancient genomes indicate population replacement in Early Neolithic Britain. Nature Ecology & Evolution 3:765–771.

14. Broodbank C. 2013. The making of the Middle Sea: a history of the Mediterranean from the beginning to the emergence of the classical world. Oxford; New York: Oxford University Press.

15. Broushaki F, Thomas MG, Link V, López S, van Dorp L, Kirsanow K, Hofmanová Z, Diekmann Y, Cassidy LM, Díez-Del-Molino D, Kousathanas A, Sell C, Robson HK, Martiniano R, Blöcher J, Scheu A, Kreutzer S, Bollongino R, Bobo D, Davudi H, Munoz O, Currat M, Abdi K, Biglari F, Craig OE, Bradley DG, Shennan S, Veeramah K, Mashkour M, Wegmann D, Hellenthal G, Burger J. 2016. Early Neolithic genomes from the eastern Fertile Crescent. Science 353:499–503.

16. Brunel S, Bennett EA, Cardin L, Garraud D, Barrand Emam H, Beylier A, Boulestin B, Chenal F, Ciesielski E, Convertini F, Dedet B, Desbrosse-Degobertiere S, Desenne S, Dubouloz J, Duday H, Escalon G, Fabre V, Gailledrat E, Gandelin M, Gleize Y, Goepfert S, Guilaine J, Hachem L, Ilett M, Lambach F, Maziere F, Perrin B, Plouin S, Pinard E, Praud I, Richard I, Riquier V, Roure R, Sendra B, Thevenet C, Thiol S, Vauquelin E, Vergnaud L, Grange T, Geigl E-M, Pruvost M. 2020. Ancient genomes from present-day France unveil 7,000 years of its demographic history. Proc Natl Acad Sci U S A 117:12791–12798.

17. Burgess RW. 2007. The Fall of Rome and the End of Civilization, by Bryan Ward-Perkins. Canadian Journal of History. doi:10.3138/cjh.42.1.83

18. Cassidy LM, Maoldúin RÓ, Kador T, Lynch A, Jones C, Woodman PC, Murphy E, Ramsey G, Dowd M, Noonan A, Campbell C, Jones ER, Mattiangeli V, Bradley DG. 2020. A dynastic elite in monumental Neolithic society. Nature 582:384–388.

19. Cassidy LM, Martiniano R, Murphy EM, Teasdale MD, Mallory J, Hartwell B, Bradley DG. 2016. Neolithic and Bronze Age migration to Ireland and establishment of the insular Atlantic genome. Proc Natl Acad Sci U S A 113:368–373.

20. Cherry D. 2007. The Frontier Zones. The Cambridge Economic History of the Greco-Roman World. doi:10.1017/chol9780521780537.028

21. Clemente F, Unterländer M, Dolgova O, Amorim CEG, Coroado-Santos F, Neuenschwander S, Ganiatsou E, Cruz Dávalos DI, Anchieri L, Michaud F, Winkelbach L, Blöcher J, Arizmendi Cárdenas YO, Sousa da Mota B, Kalliga E, Souleles A, Kontopoulos I, Karamitrou-Mentessidi G, Philaniotou O, Sampson A, Theodorou D, Tsipopoulou M, Akamatis I, Halstead P, Kotsakis K, Urem-Kotsou D, Panagiotopoulos D, Ziota C, Triantaphyllou S, Delaneau O, Jensen JD, Moreno-Mayar JV, Burger J, Sousa VC, Lao O, Malaspinas A-S, Papageorgopoulou C. 2021. The genomic history of the Aegean palatial civilizations. Cell 184:2565–2586.e21.

22. Dabney J, Knapp M, Glocke I, Gansauge M-T, Weihmann A, Nickel B, Valdiosera C, García N, Pääbo S, Arsuaga J-L, Meyer M. 2013. Complete mitochondrial genome sequence of a Middle Pleistocene cave bear reconstructed from ultrashort DNA fragments. Proc Natl Acad Sci U S A 110:15758–15763.

23. Damgaard P de B, Marchi N, Rasmussen S, Peyrot M, Renaud G, Korneliussen T, Moreno-Mayar JV, Pedersen MW, Goldberg A, Usmanova E, Baimukhanov N, Loman V, Hedeager L, Pedersen AG, Nielsen K, Afanasiev G, Akmatov K, Aldashev A, Alpaslan A, Baimbetov G, Bazaliiskii VI, Beisenov A, Boldbaatar B, Boldgiv B, Dorzhu C, Ellingvag S, Erdenebaatar D, Dajani R, Dmitriev E, Evdokimov V, Frei KM, Gromov A, Goryachev A, Hakonarson H, Hegay T, Khachatryan Z, Khaskhanov R, Kitov E, Kolbina A, Kubatbek T, Kukushkin A, Kukushkin I, Lau N, Margaryan A, Merkyte I, Mertz IV, Mertz VK, Mijiddorj E, Moiyesev V, Mukhtarova G, Nurmukhanbetov B, Orozbekova Z, Panyushkina I, Pieta K, Smrčka V, Shevnina I, Logvin A, Sjögren K-G, Štolcová T, Taravella AM, Tashbaeva K, Tkachev A, Tulegenov T, Voyakin D, Yepiskoposyan L, Undrakhbold S, Varfolomeev V, Weber A, Wilson Sayres MA, Kradin N, Allentoft ME, Orlando L, Nielsen R, Sikora M, Heyer E, Kristiansen K, Willerslev E. 2018. 137 ancient human genomes from across the Eurasian steppes. Nature 557:369–374.

24. de Barros Damgaard P, Martiniano R, Kamm J, Moreno-Mayar JV, Kroonen G, Peyrot M, Barjamovic G, Rasmussen S, Zacho C, Baimukhanov N, Zaibert V, Merz V, Biddanda A, Merz I, Loman V, Evdokimov V, Usmanova E, Hemphill B, Seguin-Orlando A, Yediay FE, Ullah I, Sjögren K-G, Iversen KH, Choin J, de la Fuente C, Ilardo M, Schroeder H, Moiseyev V, Gromov A, Polyakov A, Omura S, Senyurt SY, Ahmad H, McKenzie C, Margaryan A, Hameed A, Samad A, Gul N, Khokhar MH, Goriunova OI, Bazaliiskii VI, Novembre J, Weber AW, Orlando L, Allentoft ME, Nielsen R, Kristiansen K, Sikora M, Outram AK, Durbin R, Willerslev E. 2018. The first horse herders and the impact of early Bronze Age steppe expansions into Asia. Science 360. doi:10.1126/science.aar7711

25. de Ligt L, Tacoma LE. 2016. Approaching migration in the early Roman EmpireMigration and Mobility in the Early Roman Empire. Brill. pp. 1–22.

26. Dey HW. 2015. Introduction: Urban Living and the “Fall” of the Roman Empire. The Afterlife of the Roman City. doi:10.1017/cbo9781107706538.002

27. Ebenesersdóttir SS, Sandoval-Velasco M, Gunnarsdóttir ED, Jagadeesan A, Guðmundsdóttir VB, Thordardóttir EL, Einarsdóttir MS, Moore KHS, Sigurðsson Á, Magnúsdóttir DN, Jónsson H, Snorradóttir S, Hovig E, Møller P, Kockum I, Olsson T, Alfredsson L, Hansen TF, Werge T, Cavalleri GL, Gilbert E, Lalueza-Fox C, Walser JW 3rd, Kristjánsdóttir S, Gopalakrishnan S, Árnadóttir L, Magnússon ÓÞ, Gilbert MTP, Stefánsson K, Helgason A. 2018. Ancient genomes from Iceland reveal the making of a human population. Science 360:1028–1032.

28. Feldman M, Fernández-Domínguez E, Reynolds L, Baird D, Pearson J, Hershkovitz I, May H, Goring-Morris N, Benz M, Gresky J, Bianco RA, Fairbairn A, Mustafaoğlu G, Stockhammer PW, Posth C, Haak W, Jeong C, Krause J. 2019a. Late Pleistocene human genome suggests a local origin for the first farmers of central Anatolia. Nat Commun 10:1218.

29. Feldman M, Master DM, Bianco RA, Burri M, Stockhammer PW, Mittnik A, Aja AJ, Jeong C, Krause J. 2019b. Ancient DNA sheds light on the genetic origins of early Iron Age Philistines. Sci Adv 5:eaax0061.

30. Fernandes DM, Mittnik A, Olalde I, Lazaridis I, Cheronet O, Rohland N, Mallick S, Bernardos R, Broomandkhoshbacht N, Carlsson J, Culleton BJ, Ferry M, Gamarra B, Lari M, Mah M, Michel M, Modi A, Novak M, Oppenheimer J, Sirak KA, Stewardson K, Mandl K, Schattke C, Özdoğan KT, Lucci M, Gasperetti G, Candilio F, Salis G, Vai S, Camarós E, Calò C, Catalano G, Cueto M, Forgia V, Lozano M, Marini E, Micheletti M, Miccichè RM, Palombo MR, Ramis D, Schimmenti V, Sureda P, Teira L, Teschler-Nicola M, Kennett DJ, Lalueza-Fox C, Patterson N, Sineo L, Coppa A, Caramelli D, Pinhasi R, Reich D. 2020. The spread of steppe and Iranian-related ancestry in the islands of the western Mediterranean. Nat Ecol Evol 4:334–345.

31. Fernandes DM, Strapagiel D, Borówka P, Marciniak B, Żądzińska E, Sirak K, Siska V, Grygiel R, Carlsson J, Manica A, Lorkiewicz W, Pinhasi R. 2018. A genomic Neolithic time transect of hunter-farmer admixture in central Poland. Sci Rep 8:14879–14811.

32. Fregel R, Méndez FL, Bokbot Y, Martín-Socas D, Camalich-Massieu MD, Santana J, Morales J, Ávila-Arcos MC, Underhill PA, Shapiro B, Wojcik G, Rasmussen M, Soares AER, Kapp J, Sockell A, Rodríguez-Santos FJ, Mikdad A, Trujillo-Mederos A, Bustamante CD. 2018. Ancient genomes from North Africa evidence prehistoric migrations to the Maghreb from both the Levant and Europe. Proc Natl Acad Sci U S A 115:6774–6779.

33. Fu Q, Posth C, Hajdinjak M, Petr M, Mallick S, Fernandes D, Furtwangler A, Haak W, Meyer M, Mittnik A, Nickel B, Peltzer A, Rohland N, Slon V, Talamo S, Lazaridis I, Lipson M, Mathieson I, Schiffels S, Skoglund P, Derevianko AP, Drozdov N, Slavinsky V, Tsybankov A, Cremonesi RG, Mallegni F, Gely B, Vacca E, Morales MR, Straus LG, Neugebauer-Maresch C, Teschler-Nicola M, Constantin S, Moldovan OT, Benazzi S, Peresani M, Coppola D, Lari M, Ricci S, Ronchitelli A, Valentin F, Thevenet C, Wehrberger K, Grigorescu D, Rougier H, Crevecoeur I, Flas D, Semal P, Mannino MA, Cupillard C, Bocherens H, Conard NJ, Harvati K, Moiseyev V, Drucker DG, Svoboda J, Richards MP, Caramelli D, Pinhasi R, Kelso J, Patterson N, Krause J, Paabo S, Reich D. 2016. The genetic history of Ice Age Europe. Nature 534:200–205.

34. Furtwängler A, Neukamm J, Böhme L, Reiter E, Vollstedt M, Arora N, Singh P, Cole ST, Knauf S, Calvignac-Spencer S, Krause-Kyora B, Krause J, Schuenemann VJ, Herbig A. 2020. Comparison of target enrichment strategies for ancient pathogen DNA. Biotechniques 69:455–459.

35. Gamba C, Jones ER, Teasdale MD, McLaughlin RL, Gonzalez-Fortes G, Mattiangeli V, Domboroczki L, Kovari I, Pap I, Anders A, Whittle A, Dani J, Raczky P, Higham TF, Hofreiter M, Bradley DG, Pinhasi R. 2014. Genome flux and stasis in a five millennium transect of European prehistory. NatCommun 5:5257.

36. Gokhman D, Nissim-Rafinia M, Agranat-Tamir L, Housman G, García-Pérez R, Lizano E, Cheronet O, Mallick S, Nieves-Colón MA, Li H, Alpaslan-Roodenberg S, Novak M, Gu H, Osinski JM, Ferrando-Bernal M, Gelabert P, Lipende I, Mjungu D, Kondova I, Bontrop R, Kullmer O, Weber G, Shahar T, Dvir-Ginzberg M, Faerman M, Quillen EE, Meissner A, Lahav Y, Kandel L, Liebergall M, Prada ME, Vidal JM, Gronostajski RM, Stone AC, Yakir B, Lalueza-Fox C, Pinhasi R, Reich D, Marques-Bonet T, Meshorer E, Carmel L. 2020. Differential DNA methylation of vocal and facial anatomy genes in modern humans. Nat Commun 11:1189.

37. González-Fortes G, Jones ER, Lightfoot E, Bonsall C, Lazar C, Grandal-d’Anglade A, Garralda MD, Drak L, Siska V, Simalcsik A, Boroneanţ A, Vidal Romaní JR, Vaqueiro Rodríguez M, Arias P, Pinhasi R, Manica A, Hofreiter M. 2017. Paleogenomic Evidence for Multi-generational Mixing between Neolithic Farmers and Mesolithic Hunter-Gatherers in the Lower Danube Basin. Curr Biol 27:1801–1810.e10.

38. González-Fortes G, Tassi F, Trucchi E, Henneberger K, Paijmans JLA, Díez-Del-Molino D, Schroeder H, Susca RR, Barroso-Ruíz C, Bermudez FJ, Barroso-Medina C, Bettencourt AMS, Sampaio HA, Grandal-d’Anglade A, Salas A, de Lombera-Hermida A, Fabregas Valcarce R, Vaquero M, Alonso S, Lozano M, Rodríguez-Alvarez XP, Fernández-Rodríguez C, Manica A, Hofreiter M, Barbujani G. 2019. A western route of prehistoric human migration from Africa into the Iberian Peninsula. Proc Biol Sci 286:20182288.

39. Günther T, Malmström H, Svensson EM, Omrak A, Sánchez-Quinto F, Kılınç GM, Krzewińska M, Eriksson G, Fraser M, Edlund H, Munters AR, Coutinho A, Simões LG, Vicente M, Sjölander A, Jansen Sellevold B, Jørgensen R, Claes P, Shriver MD, Valdiosera C, Netea MG, Apel J, Lidén K, Skar B, Storå J, Götherström A, Jakobsson M. 2018. Population genomics of Mesolithic Scandinavia: Investigating early postglacial migration routes and high-latitude adaptation. PLoS Biol 16:e2003703.

40. Günther T, Valdiosera C, Malmström H, Ureña I, Rodriguez-Varela R, Sverrisdóttir ÓO, Daskalaki EA, Skoglund P, Naidoo T, Svensson EM, Bermúdez de Castro JM, Carbonell E, Dunn M, Storå J, Iriarte E, Arsuaga JL, Carretero J-M, Götherström A, Jakobsson M. 2015. Ancient genomes link early farmers from Atapuerca in Spain to modern-day Basques. Proc Natl Acad Sci U S A 112:11917–11922.

41. Haak W, Lazaridis I, Patterson N, Rohland N, Mallick S, Llamas B, Brandt G, Nordenfelt S, Harney E, Stewardson K, Fu Q, Mittnik A, Banffy E, Economou C, Francken M, Friederich S, Pena RG, Hallgren F, Khartanovich V, Khokhlov A, Kunst M, Kuznetsov P, Meller H, Mochalov O, Moiseyev V, Nicklisch N, Pichler SL, Risch R, Rojo Guerra MA, Roth C, Szecsenyi-Nagy A, Wahl J, Meyer M, Krause J, Brown D, Anthony D, Cooper A, Alt KW, Reich D. 2015. Massive migration from the steppe was a source for Indo-European languages in Europe. Nature 522:207–211.

42. Haber M, Doumet-Serhal C, Scheib CL, Xue Y, Mikulski R, Martiniano R, Fischer-Genz B, Schutkowski H, Kivisild T, Tyler-Smith C. 2019. A Transient Pulse of Genetic Admixture from the Crusaders in the Near East Identified from Ancient Genome Sequences. Am J Hum Genet 104:977–984.

43. Haber M, Doumet-Serhal C, Scheib C, Xue Y, Danecek P, Mezzavilla M, Youhanna S, Martiniano R, Prado-Martinez J, Szpak M, Matisoo-Smith E, Schutkowski H, Mikulski R, Zalloua P, Kivisild T, Tyler-Smith C. 2017. Continuity and Admixture in the Last Five Millennia of Levantine History from Ancient Canaanite and Present-Day Lebanese Genome Sequences. Am J Hum Genet 101:274–282.

44. Haber M, Nassar J, Almarri MA, Saupe T, Saag L, Griffith SJ, Doumet-Serhal C, Chanteau J, Saghieh-Beydoun M, Xue Y, Scheib CL, Tyler-Smith C. 2020. A Genetic History of the Near East from an aDNA Time Course Sampling Eight Points in the Past 4,000 Years. Am J Hum Genet 107:149–157.

45. Haller BC, Galloway J, Kelleher J, Messer PW, Ralph PL. 2019. Tree-sequence recording in SLiM opens new horizons for forward-time simulation of whole genomes. Mol Ecol Resour 19:552–566.

46. Haller BC, Messer PW. 2019. SLiM 3: Forward Genetic Simulations Beyond the Wright-Fisher Model. Mol Biol Evol 36:632–637.

47. Harney É, May H, Shalem D, Rohland N, Mallick S, Lazaridis I, Sarig R, Stewardson K, Nordenfelt S, Patterson N, Hershkovitz I, Reich D. 2018. Ancient DNA from Chalcolithic Israel reveals the role of population mixture in cultural transformation. Nat Commun 9:3336.

48. Harney É, Patterson N, Reich D, Wakeley J. 2021. Assessing the performance of qpAdm: a statistical tool for studying population admixture. Genetics 217. doi:10.1093/genetics/iyaa045

49. Harper K. 2017. The fate of Rome: climate, disease, and the end of an empire. Princeton: Princeton University Press.

50. Hofmanová Z, Kreutzer S, Hellenthal G, Sell C, Diekmann Y, Díez-Del-Molino D, van Dorp L, López S, Kousathanas A, Link V, Kirsanow K, Cassidy LM, Martiniano R, Strobel M, Scheu A, Kotsakis K, Halstead P, Triantaphyllou S, Kyparissi-Apostolika N, Urem-Kotsou D, Ziota C, Adaktylou F, Gopalan S, Bobo DM, Winkelbach L, Blöcher J, Unterländer M, Leuenberger C, Çilingiroğlu Ç, Horejs B, Gerritsen F, Shennan SJ, Bradley DG, Currat M, Veeramah KR, Wegmann D, Thomas MG, Papageorgopoulou C, Burger J. 2016. Early farmers from across Europe directly descended from Neolithic Aegeans. Proc Natl Acad Sci U S A 113:6886–6891.

51. Järve M, Saag L, Scheib CL, Pathak AK, Montinaro F, Pagani L, Flores R, Guellil M, Saag L, Tambets K, Kushniarevich A, Solnik A, Varul L, Zadnikov S, Petrauskas O, Avramenko M, Magomedov B, Didenko S, Toshev G, Bruyako I, Grechko D, Okatenko V, Gorbenko K, Smyrnov O, Heiko A, Reida R, Sapiehin S, Sirotin S, Tairov A, Beisenov A, Starodubtsev M, Vasilev V, Nechvaloda A, Atabiev B, Litvinov S, Ekomasova N, Dzhaubermezov M, Voroniatov S, Utevska O, Shramko I, Khusnutdinova E, Metspalu M, Savelev N, Kriiska A, Kivisild T, Villems R. 2019. Shifts in the Genetic Landscape of the Western Eurasian Steppe Associated with the Beginning and End of the Scythian Dominance. Curr Biol 29:2430–2441.e10.

52. Jeong C, Balanovsky O, Lukianova E, Kahbatkyzy N, Flegontov P, Zaporozhchenko V, Immel A, Wang C-C, Ixan O, Khussainova E, Bekmanov B, Zaibert V, Lavryashina M, Pocheshkhova E, Yusupov Y, Agdzhoyan A, Koshel S, Bukin A, Nymadawa P, Turdikulova S, Dalimova D, Churnosov M, Skhalyakho R, Daragan D, Bogunov Y, Bogunova A, Shtrunov A, Dubova N, Zhabagin M, Yepiskoposyan L, Churakov V, Pislegin N, Damba L, Saroyants L, Dibirova K, Atramentova L, Utevska O, Idrisov E, Kamenshchikova E, Evseeva I, Metspalu M, Outram AK, Robbeets M, Djansugurova L, Balanovska E, Schiffels S, Haak W, Reich D, Krause J. 2019. The genetic history of admixture across inner Eurasia. Nat Ecol Evol 3:966–976.

53. Jones ER, Gonzalez-Fortes G, Connell S, Siska V, Eriksson A, Martiniano R, McLaughlin RL, Gallego Llorente M, Cassidy LM, Gamba C, Meshveliani T, Bar-Yosef O, Muller W, Belfer-Cohen A, Matskevich Z, Jakeli N, Higham TF, Currat M, Lordkipanidze D, Hofreiter M, Manica A, Pinhasi R, Bradley DG. 2015. Upper Palaeolithic genomes reveal deep roots of modern Eurasians. NatCommun 6:8912.

54. Jones ER, Zarina G, Moiseyev V, Lightfoot E, Nigst PR, Manica A, Pinhasi R, Bradley DG. 2017. The Neolithic Transition in the Baltic Was Not Driven by Admixture with Early European Farmers. Curr Biol 27:576–582.

55. Jónsson H, Ginolhac A, Schubert M, Johnson PLF, Orlando L. 2013. mapDamage2.0: fast approximate Bayesian estimates of ancient DNA damage parameters. Bioinformatics 29:1682–1684.

56. Kelleher J, Etheridge AM, McVean G. 2016. Efficient Coalescent Simulation and Genealogical Analysis for Large Sample Sizes. PLoS Comput Biol 12:e1004842.

57. Kelleher J, Thornton KR, Ashander J, Ralph PL. 2018. Efficient pedigree recording for fast population genetics simulation. PLoS Comput Biol 14:e1006581.

58. Keller A, Graefen A, Ball M, Matzas M, Boisguerin V, Maixner F, Leidinger P, Backes C, Khairat R, Forster M, Stade B, Franke A, Mayer J, Spangler J, McLaughlin S, Shah M, Lee C, Harkins TT, Sartori A, Moreno-Estrada A, Henn B, Sikora M, Semino O, Chiaroni J, Rootsi S, Myres NM, Cabrera VM, Underhill PA, Bustamante CD, Vigl EE, Samadelli M, Cipollini G, Haas J, Katus H, O’Connor BD, Carlson MRJ, Meder B, Blin N, Meese E, Pusch CM, Zink A. 2012. New insights into the Tyrolean Iceman’s origin and phenotype as inferred by whole-genome sequencing. Nat Commun 3:698.

59. Kılınç GM, Omrak A, Özer F, Günther T, Büyükkarakaya AM, Bıçakçı E, Baird D, Dönertaş HM, Ghalichi A, Yaka R, Koptekin D, Açan SC, Parvizi P, Krzewińska M, Daskalaki EA, Yüncü E, Dağtaş ND, Fairbairn A, Pearson J, Mustafaoğlu G, Erdal YS, Çakan YG, Togan İ, Somel M, Storå J, Jakobsson M, Götherström A. 2016. The Demographic Development of the First Farmers in Anatolia. Curr Biol 26:2659–2666.

60. Korneliussen TS, Albrechtsen A, Nielsen R. 2014. ANGSD: Analysis of Next Generation Sequencing Data. BMC Bioinformatics 15:356.

61. Kovacevic L, Tambets K, Ilumäe A-M, Kushniarevich A, Yunusbayev B, Solnik A, Bego T, Primorac D, Skaro V, Leskovac A, Jakovski Z, Drobnic K, Tolk H-V, Kovacevic S, Rudan P, Metspalu E, Marjanovic D. 2014. Standing at the gateway to Europe--the genetic structure of Western balkan populations based on autosomal and haploid markers. PLoS One 9:e105090.

62. Krzewińska M, Kılınç GM, Juras A, Koptekin D, Chyleński M, Nikitin AG, Shcherbakov N, Shuteleva I, Leonova T, Kraeva L, Sungatov FA, Sultanova AN, Potekhina I, Łukasik S, Krenz-Niedbała M, Dalén L, Sinika V, Jakobsson M, Storå J, Götherström A. 2018a. Ancient genomes suggest the eastern Pontic-Caspian steppe as the source of western Iron Age nomads. Sci Adv 4:eaat4457.

63. Krzewińska M, Kjellström A, Günther T, Hedenstierna-Jonson C, Zachrisson T, Omrak A, Yaka R, Kılınç GM, Somel M, Sobrado V, Evans J, Knipper C, Jakobsson M, Storå J, Götherström A. 2018b. Genomic and Strontium Isotope Variation Reveal Immigration Patterns in a Viking Age Town. Curr Biol 28:2730–2738.e10.

64. Lamnidis TC, Majander K, Jeong C, Salmela E, Wessman A, Moiseyev V, Khartanovich V, Balanovsky O, Ongyerth M, Weihmann A, Sajantila A, Kelso J, Pääbo S, Onkamo P, Haak W, Krause J, Schiffels S. 2018. Ancient Fennoscandian genomes reveal origin and spread of Siberian ancestry in Europe. Nat Commun 9:5018.

65. Lazaridis I, Mittnik A, Patterson N, Mallick S, Rohland N, Pfrengle S, Furtwangler A, Peltzer A, Posth C, Vasilakis A, McGeorge PJP, Konsolaki-Yannopoulou E, Korres G, Martlew H, Michalodimitrakis M, Ozsait M, Ozsait N, Papathanasiou A, Richards M, Roodenberg SA, Tzedakis Y, Arnott R, Fernandes DM, Hughey JR, Lotakis DM, Navas PA, Maniatis Y, Stamatoyannopoulos JA, Stewardson K, Stockhammer P, Pinhasi R, Reich D, Krause J, Stamatoyannopoulos G. 2017. Genetic origins of the Minoans and Mycenaeans. Nature 548:214–218.

66. Lazaridis I, Nadel D, Rollefson G, Merrett DC, Rohland N, Mallick S, Fernandes D, Novak M, Gamarra B, Sirak K, Connell S, Stewardson K, Harney E, Fu Q, Gonzalez-Fortes G, Jones ER, Roodenberg SA, Lengyel G, Bocquentin F, Gasparian B, Monge JM, Gregg M, Eshed V, Mizrahi A-S, Meiklejohn C, Gerritsen F, Bejenaru L, Blüher M, Campbell A, Cavalleri G, Comas D, Froguel P, Gilbert E, Kerr SM, Kovacs P, Krause J, McGettigan D, Merrigan M, Merriwether DA, O’Reilly S, Richards MB, Semino O, Shamoon-Pour M, Stefanescu G, Stumvoll M, Tönjes A, Torroni A, Wilson JF, Yengo L, Hovhannisyan NA, Patterson N, Pinhasi R, Reich D. 2016. Genomic insights into the origin of farming in the ancient Near East. Nature 536:419–424.

67. Lazaridis I, Patterson N, Mittnik A, Renaud G, Mallick S, Kirsanow K, Sudmant PH, Schraiber JG, Castellano S, Lipson M, Berger B, Economou C, Bollongino R, Fu Q, Bos KI, Nordenfelt S, Li H, de Filippo C, Prufer K, Sawyer S, Posth C, Haak W, Hallgren F, Fornander E, Rohland N, Delsate D, Francken M, Guinet JM, Wahl J, Ayodo G, Babiker HA, Bailliet G, Balanovska E, Balanovsky O, Barrantes R, Bedoya G, Ben-Ami H, Bene J, Berrada F, Bravi CM, Brisighelli F, Busby GB, Cali F, Churnosov M, Cole DE, Corach D, Damba L, van Driem G, Dryomov S, Dugoujon JM, Fedorova SA, Gallego Romero I, Gubina M, Hammer M, Henn BM, Hervig T, Hodoglugil U, Jha AR, Karachanak-Yankova S, Khusainova R, Khusnutdinova E, Kittles R, Kivisild T, Klitz W, Kucinskas V, Kushniarevich A, Laredj L, Litvinov S, Loukidis T, Mahley RW, Melegh B, Metspalu E, Molina J, Mountain J, Nakkalajarvi K, Nesheva D, Nyambo T, Osipova L, Parik J, Platonov F, Posukh O, Romano V, Rothhammer F, Rudan I, Ruizbakiev R, Sahakyan H, Sajantila A, Salas A, Starikovskaya EB, Tarekegn A, Toncheva D, Turdikulova S, Uktveryte I, Utevska O, Vasquez R, Villena M, Voevoda M, Winkler CA, Yepiskoposyan L, Zalloua P, Zemunik T, Cooper A, Capelli C, Thomas MG, Ruiz-Linares A, Tishkoff SA, Singh L, Thangaraj K, Villems R, Comas D, Sukernik R, Metspalu M, Meyer M, Eichler EE, Burger J, Slatkin M, Paabo S, Kelso J, Reich D, Krause J. 2014. Ancient human genomes suggest three ancestral populations for present-day Europeans. Nature 513:409–413.

68. Li H, Durbin R. 2009. Fast and accurate short read alignment with Burrows-Wheeler transform. Bioinformatics 25:1754–1760.

69. Linderholm A, Kılınç GM, Szczepanek A, Włodarczak P, Jarosz P, Belka Z, Dopieralska J, Werens K, Górski J, Mazurek M, Hozer M, Rybicka M, Ostrowski M, Bagińska J, Koman W, Rodríguez-Varela R, Storå J, Götherström A, Krzewińska M. 2020. Corded Ware cultural complexity uncovered using genomic and isotopic analysis from south-eastern Poland. Sci Rep 10:6885.

70. Lipson M, Szécsényi-Nagy A, Mallick S, Pósa A, Stégmár B, Keerl V, Rohland N, Stewardson K, Ferry M, Michel M, Oppenheimer J, Broomandkhoshbacht N, Harney E, Nordenfelt S, Llamas B, Gusztáv Mende B, Köhler K, Oross K, Bondár M, Marton T, Osztás A, Jakucs J, Paluch T, Horváth F, Csengeri P, Koós J, Sebők K, Anders A, Raczky P, Regenye J, Barna JP, Fábián S, Serlegi G, Toldi Z, Gyöngyvér Nagy E, Dani J, Molnár E, Pálfi G, Márk L, Melegh B, Bánfai Z, Domboróczki L, Fernández-Eraso J, Antonio Mujika-Alustiza J, Alonso Fernández C, Jiménez Echevarría J, Bollongino R, Orschiedt J, Schierhold K, Meller H, Cooper A, Burger J, Bánffy E, Alt KW, Lalueza-Fox C, Haak W, Reich D. 2017. Parallel palaeogenomic transects reveal complex genetic history of early European farmers. Nature 551:368–372.

71. Mallick S, Li H, Lipson M, Mathieson I, Gymrek M, Racimo F, Zhao M, Chennagiri N, Nordenfelt S, Tandon A, Skoglund P, Lazaridis I, Sankararaman S, Fu Q, Rohland N, Renaud G, Erlich Y, Willems T, Gallo C, Spence JP, Song YS, Poletti G, Balloux F, van Driem G, de Knijff P, Romero IG, Jha AR, Behar DM, Bravi CM, Capelli C, Hervig T, Moreno-Estrada A, Posukh OL, Balanovska E, Balanovsky O, Karachanak-Yankova S, Sahakyan H, Toncheva D, Yepiskoposyan L, Tyler-Smith C, Xue Y, Abdullah MS, Ruiz-Linares A, Beall CM, Di Rienzo A, Jeong C, Starikovskaya EB, Metspalu E, Parik J, Villems R, Henn BM, Hodoglugil U, Mahley R, Sajantila A, Stamatoyannopoulos G, Wee JTS, Khusainova R, Khusnutdinova E, Litvinov S, Ayodo G, Comas D, Hammer MF, Kivisild T, Klitz W, Winkler CA, Labuda D, Bamshad M, Jorde LB, Tishkoff SA, Watkins WS, Metspalu M, Dryomov S, Sukernik R, Singh L, Thangaraj K, Pääbo S, Kelso J, Patterson N, Reich D. 2016. The Simons Genome Diversity Project: 300 genomes from 142 diverse populations. Nature 538:201–206.

72. Mallick S, Micco A, Mah M, Ringbauer H, Lazaridis I, Olalde I, Patterson N, Reich D. 2023. The Allen Ancient DNA Resource (AADR): A curated compendium of ancient human genomes. bioRxiv. doi:10.1101/2023.04.06.535797

73. Malmström H, Günther T, Svensson EM, Juras A, Fraser M, Munters AR, Pospieszny Ł, Tõrv M, Lindström J, Götherström A, Storå J, Jakobsson M. 2019. The genomic ancestry of the Scandinavian Battle Axe Culture people and their relation to the broader Corded Ware horizon. Proc Biol Sci 286:20191528.

74. Marcus JH, Posth C, Ringbauer H, Lai L, Skeates R, Sidore C, Beckett J, Furtwängler A, Olivieri A, Chiang CWK, Al-Asadi H, Dey K, Joseph TA, Liu C-C, Der Sarkissian C, Radzevičiūtė R, Michel M, Gradoli MG, Marongiu P, Rubino S, Mazzarello V, Rovina D, La Fragola A, Serra RM, Bandiera P, Bianucci R, Pompianu E, Murgia C, Guirguis M, Orquin RP, Tuross N, van Dommelen P, Haak W, Reich D, Schlessinger D, Cucca F, Krause J, Novembre J. 2020. Genetic history from the Middle Neolithic to present on the Mediterranean island of Sardinia. Nat Commun 11:939.

75. Margaryan A, Lawson DJ, Sikora M, Racimo F, Rasmussen S, Moltke I, Cassidy LM, Jørsboe E, Ingason A, Pedersen MW, Korneliussen T, Wilhelmson H, Buś MM, de Barros Damgaard P, Martiniano R, Renaud G, Bhérer C, Moreno-Mayar JV, Fotakis AK, Allen M, Allmäe R, Molak M, Cappellini E, Scorrano G, McColl H, Buzhilova A, Fox A, Albrechtsen A, Schütz B, Skar B, Arcini C, Falys C, Jonson CH, Błaszczyk D, Pezhemsky D, Turner-Walker G, Gestsdóttir H, Lundstrøm I, Gustin I, Mainland I, Potekhina I, Muntoni IM, Cheng J, Stenderup J, Ma J, Gibson J, Peets J, Gustafsson J, Iversen KH, Simpson L, Strand L, Loe L, Sikora M, Florek M, Vretemark M, Redknap M, Bajka M, Pushkina T, Søvsø M, Grigoreva N, Christensen T, Kastholm O, Uldum O, Favia P, Holck P, Sten S, Arge SV, Ellingvåg S, Moiseyev V, Bogdanowicz W, Magnusson Y, Orlando L, Pentz P, Jessen MD, Pedersen A, Collard M, Bradley DG, Jørkov ML, Arneborg J, Lynnerup N, Price N, Gilbert MTP, Allentoft ME, Bill J, Sindbæk SM, Hedeager L, Kristiansen K, Nielsen R, Werge T, Willerslev E. 2020. Population genomics of the Viking world. Nature 585:390–396.

76. Martiniano R, Caffell A, Holst M, Hunter-Mann K, Montgomery J, Müldner G, McLaughlin RL, Teasdale MD, van Rheenen W, Veldink JH, van den Berg LH, Hardiman O, Carroll M, Roskams S, Oxley J, Morgan C, Thomas MG, Barnes I, McDonnell C, Collins MJ, Bradley DG. 2016. Genomic signals of migration and continuity in Britain before the Anglo-Saxons. Nat Commun 7:10326.

77. Martiniano R, Cassidy LM, Ó Maoldúin R, Mclaughlin R, M Silva N, Manco L, Fidalgo D, Pereira T, Coelho MJ, Serra M, Burger J, Parreira R, Moran E, Valera AC, Porfirio E, Boaventura R, M Silva A, G Bradley D. 2017. The population genomics of archaeological transition in west Iberia: Investigation of ancient substructure using imputation and haplotype-based methods. doi:10.1371/journal.pgen.1006852

78. Martin M. 2011. Cutadapt removes adapter sequences from high-throughput sequencing reads. EMBnet.journal 17:10.

79. Mathieson I, Alpaslan-Roodenberg S, Posth C, Szecsenyi-Nagy A, Rohland N, Mallick S, Olalde I, Broomandkhoshbacht N, Candilio F, Cheronet O, Fernandes D, Ferry M, Gamarra B, Fortes GG, Haak W, Harney E, Jones E, Keating D, Krause-Kyora B, Kucukkalipci I, Michel M, Mittnik A, Nagele K, Novak M, Oppenheimer J, Patterson N, Pfrengle S, Sirak K, Stewardson K, Vai S, Alexandrov S, Alt KW, Andreescu R, Antonovic D, Ash A, Atanassova N, Bacvarov K, Gusztav MB, Bocherens H, Bolus M, Boroneant A, Boyadzhiev Y, Budnik A, Burmaz J, Chohadzhiev S, Conard NJ, Cottiaux R, Cuka M, Cupillard C, Drucker DG, Elenski N, Francken M, Galabova B, Ganetsovski G, Gely B, Hajdu T, Handzhyiska V, Harvati K, Higham T, Iliev S, Jankovic I, Karavanic I, Kennett DJ, Komso D, Kozak A, Labuda D, Lari M, Lazar C, Leppek M, Leshtakov K, Vetro DL, Los D, Lozanov I, Malina M, Martini F, McSweeney K, Meller H, Mendusic M, Mirea P, Moiseyev V, Petrova V, Price TD, Simalcsik A, Sineo L, Slaus M, Slavchev V, Stanev P, Starovic A, Szeniczey T, Talamo S, Teschler-Nicola M, Thevenet C, Valchev I, Valentin F, Vasilyev S, Veljanovska F, Venelinova S, Veselovskaya E, Viola B, Virag C, Zaninovic J, Zauner S, Stockhammer PW, Catalano G, Krauss R, Caramelli D, Zarina G, Gaydarska B, Lillie M, Nikitin AG, Potekhina I, Papathanasiou A, Boric D, Bonsall C, Krause J, Pinhasi R, Reich D. 2018. The genomic history of southeastern Europe. Nature 555:197–203.

80. Mathieson I, Lazaridis I, Rohland N, Mallick S, Patterson N, Roodenberg SA, Harney E, Stewardson K, Fernandes D, Novak M, Sirak K, Gamba C, Jones ER, Llamas B, Dryomov S, Pickrell J, Arsuaga JL, de Castro JM, Carbonell E, Gerritsen F, Khokhlov A, Kuznetsov P, Lozano M, Meller H, Mochalov O, Moiseyev V, Guerra MA, Roodenberg J, Verges JM, Krause J, Cooper A, Alt KW, Brown D, Anthony D, Lalueza-Fox C, Haak W, Pinhasi R, Reich D. 2015. Genome-wide patterns of selection in 230 ancient Eurasians. Nature 528:499–503.

81. Meyer M, Kircher M. 2010. Illumina sequencing library preparation for highly multiplexed target capture and sequencing. Cold Spring Harb Protoc 2010. doi:10.1101/pdb.prot5448

82. Mittnik A, Massy K, Knipper C, Wittenborn F, Friedrich R, Pfrengle S, Burri M, Carlichi-Witjes N, Deeg H, Furtwängler A, Harbeck M, von Heyking K, Kociumaka C, Kucukkalipci I, Lindauer S, Metz S, Staskiewicz A, Thiel A, Wahl J, Haak W, Pernicka E, Schiffels S, Stockhammer PW, Krause J. 2019. Kinship-based social inequality in Bronze Age Europe. Science 366:731–734.

83. Mittnik A, Wang C-C, Pfrengle S, Daubaras M, Zariņa G, Hallgren F, Allmäe R, Khartanovich V, Moiseyev V, Tõrv M, Furtwängler A, Andrades Valtueña A, Feldman M, Economou C, Oinonen M, Vasks A, Balanovska E, Reich D, Jankauskas R, Haak W, Schiffels S, Krause J. 2018. The genetic prehistory of the Baltic Sea region. Nat Commun 9:442.

84. Moots HM, Antonio M, Sawyer S, Spence JP, Oberreiter V, Weiß CL, Lucci M, Cherifi YMS, La Pastina F, Genchi F, Praxmeier E, Zagorc B, Cheronot O, Özdoğan KT, Demetz L, Amrani S, Candilio F, De Angelis D, Gasperetti G, Fernandes D, Gao Z, Fantar M, Coppa A, Pritchard JK, Pinhasi R. 2022. A Genetic History of Continuity and Mobility in the Iron Age Central Mediterranean. bioRxiv. doi:10.1101/2022.03.13.483276

85. Narasimhan VM, Patterson N, Moorjani P, Rohland N, Bernardos R, Mallick S, Lazaridis I, Nakatsuka N, Olalde I, Lipson M, Kim AM, Olivieri LM, Coppa A, Vidale M, Mallory J, Moiseyev V, Kitov E, Monge J, Adamski N, Alex N, Broomandkhoshbacht N, Candilio F, Callan K, Cheronet O, Culleton BJ, Ferry M, Fernandes D, Freilich S, Gamarra B, Gaudio D, Hajdinjak M, Harney É, Harper TK, Keating D, Lawson AM, Mah M, Mandl K, Michel M, Novak M, Oppenheimer J, Rai N, Sirak K, Slon V, Stewardson K, Zalzala F, Zhang Z, Akhatov G, Bagashev AN, Bagnera A, Baitanayev B, Bendezu-Sarmiento J, Bissembaev AA, Bonora GL, Chargynov TT, Chikisheva T, Dashkovskiy PK, Derevianko A, Dobeš M, Douka K, Dubova N, Duisengali MN, Enshin D, Epimakhov A, Fribus AV, Fuller D, Goryachev A, Gromov A, Grushin SP, Hanks B, Judd M, Kazizov E, Khokhlov A, Krygin AP, Kupriyanova E, Kuznetsov P, Luiselli D, Maksudov F, Mamedov AM, Mamirov TB, Meiklejohn C, Merrett DC, Micheli R, Mochalov O, Mustafokulov S, Nayak A, Pettener D, Potts R, Razhev D, Rykun M, Sarno S, Savenkova TM, Sikhymbaeva K, Slepchenko SM, Soltobaev OA, Stepanova N, Svyatko S, Tabaldiev K, Teschler-Nicola M, Tishkin AA, Tkachev VV, Vasilyev S, Velemínský P, Voyakin D, Yermolayeva A, Zahir M, Zubkov VS, Zubova A, Shinde VS, Lalueza-Fox C, Meyer M, Anthony D, Boivin N, Thangaraj K, Kennett DJ, Frachetti M, Pinhasi R, Reich D. 2019. The formation of human populations in South and Central Asia. Science 365. doi:10.1126/science.aat7487

86. Nikitin AG, Stadler P, Kotova N, Teschler-Nicola M, Price TD, Hoover J, Kennett DJ, Lazaridis I, Rohland N, Lipson M, Reich D. 2019. Interactions between earliest Linearbandkeramik farmers and central European hunter gatherers at the dawn of European Neolithization. Sci Rep 9:19544.

87. Novembre J, Johnson T, Bryc K, Kutalik Z, Boyko AR, Auton A, Indap A, King KS, Bergmann S, Nelson MR, Stephens M, Bustamante CD. 2008. Genes mirror geography within Europe. Nature 456:98–101.

88. Olalde I, Allentoft ME, Sánchez-Quinto F, Santpere G, Chiang CWK, DeGiorgio M, Prado-Martinez J, Rodríguez JA, Rasmussen S, Quilez J, Ramírez O, Marigorta UM, Fernández-Callejo M, Prada ME, Encinas JMV, Nielsen R, Netea MG, Novembre J, Sturm RA, Sabeti P, Marquès-Bonet T, Navarro A, Willerslev E, Lalueza-Fox C. 2014. Derived immune and ancestral pigmentation alleles in a 7,000-year-old Mesolithic European. Nature 507:225–228.

89. Olalde I, Brace S, Allentoft ME, Armit I, Kristiansen K, Booth T, Rohland N, Mallick S, Szécsényi-Nagy A, Mittnik A, Altena E, Lipson M, Lazaridis I, Harper TK, Patterson N, Broomandkhoshbacht N, Diekmann Y, Faltyskova Z, Fernandes D, Ferry M, Harney E, de Knijff P, Michel M, Oppenheimer J, Stewardson K, Barclay A, Alt KW, Liesau C, Ríos P, Blasco C, Miguel JV, García RM, Fernández AA, Bánffy E, Bernabò-Brea M, Billoin D, Bonsall C, Bonsall L, Allen T, Büster L, Carver S, Navarro LC, Craig OE, Cook GT, Cunliffe B, Denaire A, Dinwiddy KE, Dodwell N, Ernée M, Evans C, Kuchařík M, Farré JF, Fowler C, Gazenbeek M, Pena RG, Haber-Uriarte M, Haduch E, Hey G, Jowett N, Knowles T, Massy K, Pfrengle S, Lefranc P, Lemercier O, Lefebvre A, Martínez CH, Olmo VG, Ramírez AB, Maurandi JL, Majó T, McKinley JI, McSweeney K, Mende BG, Modi A, Kulcsár G, Kiss V, Czene A, Patay R, Endrődi A, Köhler K, Hajdu T, Szeniczey T, Dani J, Bernert Z, Hoole M, Cheronet O, Keating D, Velemínský P, Dobeš M, Candilio F, Brown F, Fernández RF, Herrero-Corral A-M, Tusa S, Carnieri E, Lentini L, Valenti A, Zanini A, Waddington C, Delibes G, Guerra-Doce E, Neil B, Brittain M, Luke M, Mortimer R, Desideri J, Besse M, Brücken G, Furmanek M, Hałuszko A, Mackiewicz M, Rapiński A, Leach S, Soriano I, Lillios KT, Cardoso JL, Pearson MP, Włodarczak P, Price TD, Prieto P, Rey P-J, Risch R, Rojo Guerra MA, Schmitt A, Serralongue J, Silva AM, Smrčka V, Vergnaud L, Zilhão J, Caramelli D, Higham T, Thomas MG, Kennett DJ, Fokkens H, Heyd V, Sheridan A, Sjögren K-G, Stockhammer PW, Krause J, Pinhasi R, Haak W, Barnes I, Lalueza-Fox C, Reich D. 2018. The Beaker phenomenon and the genomic transformation of northwest Europe. Nature 555:190–196.

90. Olalde I, Mallick S, Patterson N, Rohland N, Villalba-Mouco V, Silva M, Dulias K, Edwards CJ, Gandini F, Pala M, Soares P, Ferrando-Bernal M, Adamski N, Broomandkhoshbacht N, Cheronet O, Culleton BJ, Fernandes D, Lawson AM, Mah M, Oppenheimer J, Stewardson K, Zhang Z, Jimenez Arenas JM, Toro Moyano IJ, Salazar-Garcia DC, Castanyer P, Santos M, Tremoleda J, Lozano M, Garcia Borja P, Fernandez-Eraso J, Mujika-Alustiza JA, Barroso C, Bermudez FJ, Viguera Minguez E, Burch J, Coromina N, Vivo D, Cebria A, Fullola JM, Garcia-Puchol O, Morales JI, Oms FX, Majo T, Verges JM, Diaz-Carvajal A, Ollich-Castanyer I, Lopez-Cachero FJ, Silva AM, Alonso-Fernandez C, Delibes de Castro G, Jimenez Echevarria J, Moreno-Marquez A, Pascual Berlanga G, Ramos-Garcia P, Ramos-Munoz J, Vijande Vila E, Aguilella Arzo G, Esparza Arroyo A, Lillios KT, Mack J, Velasco-Vazquez J, Waterman A, Benitez de Lugo Enrich L, Benito Sanchez M, Agusti B, Codina F, de Prado G, Estalrrich A, Fernandez Flores A, Finlayson C, Finlayson G, Finlayson S, Giles-Guzman F, Rosas A, Barciela Gonzalez V, Garcia Atienzar G, Hernandez Perez MS, Llanos A, Carrion Marco Y, Collado Beneyto I, Lopez-Serrano D, Sanz Tormo M, Valera AC, Blasco C, Liesau C, Rios P, Daura J, de Pedro Micho MJ, Diez-Castillo AA, Flores Fernandez R, Frances Farre J, Garrido-Pena R, Goncalves VS, Guerra-Doce E, Herrero-Corral AM, Juan-Cabanilles J, Lopez-Reyes D, McClure SB, Merino Perez M, Oliver Foix A, Sanz Borras M, Sousa AC, Vidal Encinas JM, Kennett DJ, Richards MB, Werner Alt K, Haak W, Pinhasi R, Lalueza-Fox C, Reich D. 2019. The genomic history of the Iberian Peninsula over the past 8000 years. Science 363:1230–1234.

91. Olalde I, Schroeder H, Sandoval-Velasco M, Vinner L, Lobon I, Ramirez O, Civit S, Garcia Borja P, Salazar-Garcia DC, Talamo S, Maria Fullola J, Xavier Oms F, Pedro M, Martinez P, Sanz M, Daura J, Zilhao J, Marques-Bonet T, Gilbert MT, Lalueza-Fox C. 2015. A Common Genetic Origin for Early Farmers from Mediterranean Cardial and Central European LBK Cultures. MolBiolEvol 32:3132–3142.

92. Oleson JP. 2008. The Oxford Handbook of Engineering and Technology in the Classical World. Oxford University Press.

93. Omrak A, Günther T, Valdiosera C, Svensson EM, Malmström H, Kiesewetter H, Aylward W, Storå J, Jakobsson M, Götherström A. 2016. Genomic Evidence Establishes Anatolia as the Source of the European Neolithic Gene Pool. Curr Biol 26:270–275.

94. O’Sullivan N, Posth C, Coia V, Schuenemann VJ, Price TD, Wahl J, Pinhasi R, Zink A, Krause J, Maixner F. 2018. Ancient genome-wide analyses infer kinship structure in an Early Medieval Alemannic graveyard. Sci Adv 4:eaao1262.

95. Pagani L, Lawson DJ, Jagoda E, Mörseburg A, Eriksson A, Mitt M, Clemente F, Hudjashov G, DeGiorgio M, Saag L, Wall JD, Cardona A, Mägi R, Wilson Sayres MA, Kaewert S, Inchley C, Scheib CL, Järve M, Karmin M, Jacobs GS, Antao T, Iliescu FM, Kushniarevich A, Ayub Q, Tyler-Smith C, Xue Y, Yunusbayev B, Tambets K, Mallick CB, Saag L, Pocheshkhova E, Andriadze G, Muller C, Westaway MC, Lambert DM, Zoraqi G, Turdikulova S, Dalimova D, Sabitov Z, Sultana GNN, Lachance J, Tishkoff S, Momynaliev K, Isakova J, Damba LD, Gubina M, Nymadawa P, Evseeva I, Atramentova L, Utevska O, Ricaut F-X, Brucato N, Sudoyo H, Letellier T, Cox MP, Barashkov NA, Skaro V, Mulahasanovic L, Primorac D, Sahakyan H, Mormina M, Eichstaedt CA, Lichman DV, Abdullah S, Chaubey G, Wee JTS, Mihailov E, Karunas A, Litvinov S, Khusainova R, Ekomasova N, Akhmetova V, Khidiyatova I, Marjanović D, Yepiskoposyan L, Behar DM, Balanovska E, Metspalu A, Derenko M, Malyarchuk B, Voevoda M, Fedorova SA, Osipova LP, Lahr MM, Gerbault P, Leavesley M, Migliano AB, Petraglia M, Balanovsky O, Khusnutdinova EK, Metspalu E, Thomas MG, Manica A, Nielsen R, Villems R, Willerslev E, Kivisild T, Metspalu M. 2016. Genomic analyses inform on migration events during the peopling of Eurasia. Nature 538:238–242.

96. Patterson N, Moorjani P, Luo Y, Mallick S, Rohland N, Zhan Y, Genschoreck T, Webster T, Reich D. 2012. Ancient admixture in human history. Genetics 192:1065–1093.

97. Pinhasi R, Fernandes DM, Sirak K, Cheronet O. 2019. Isolating the human cochlea to generate bone powder for ancient DNA analysis. NatProtoc 14:1194–1205.

98. Pinhasi R, Fernandes D, Sirak K, Novak M, Connell S, Gerritsen FA. 2015. Optimal Ancient DNA Yields from the Inner Ear Part of the Human Petrous Bone. PLoS One 10. doi:10.1371/journal.pone.0129102

99. Posth C, Zaro V, Spyrou MA, Vai S, Gnecchi-Ruscone GA, Modi A, Peltzer A, Mötsch A, Nägele K, Vågene ÅJ, Nelson EA, Radzevičiūtė R, Freund C, Bondioli LM, Cappuccini L, Frenzel H, Pacciani E, Boschin F, Capecchi G, Martini I, Moroni A, Ricci S, Sperduti A, Turchetti MA, Riga A, Zavattaro M, Zifferero A, Heyne HO, Fernández-Domínguez E, Kroonen GJ, McCormick M, Haak W, Lari M, Barbujani G, Bondioli L, Bos KI, Caramelli D, Krause J. 2021. The origin and legacy of the Etruscans through a 2000-year archeogenomic time transect. Sci Adv 7:eabi7673.

100. Prüfer K, de Filippo C, Grote S, Mafessoni F, Korlević P, Hajdinjak M, Vernot B, Skov L, Hsieh P, Peyrégne S, Reher D, Hopfe C, Nagel S, Maricic T, Fu Q, Theunert C, Rogers R, Skoglund P, Chintalapati M, Dannemann M, Nelson BJ, Key FM, Rudan P, Kućan Ž, Gušić I, Golovanova LV, Doronichev VB, Patterson N, Reich D, Eichler EE, Slatkin M, Schierup MH, Andrés AM, Kelso J, Meyer M, Pääbo S. 2017. A high-coverage Neandertal genome from Vindija Cave in Croatia. Science 358:655–658.

101. Renaud G, Slon V, Duggan AT, Kelso J. 2015. Schmutzi: estimation of contamination and endogenous mitochondrial consensus calling for ancient DNA. Genome Biol 16:224.

102. Rivollat M, Jeong C, Schiffels S, Küçükkalıpçı İ, Pemonge M-H, Rohrlach AB, Alt KW, Binder D, Friederich S, Ghesquière E, Gronenborn D, Laporte L, Lefranc P, Meller H, Réveillas H, Rosenstock E, Rottier S, Scarre C, Soler L, Wahl J, Krause J, Deguilloux M-F, Haak W. 2020. Ancient genome-wide DNA from France highlights the complexity of interactions between Mesolithic hunter-gatherers and Neolithic farmers. Sci Adv 6:eaaz5344.

103. Rodríguez-Varela R, Günther T, Krzewińska M, Storå J, Gillingwater TH, MacCallum M, Arsuaga JL, Dobney K, Valdiosera C, Jakobsson M, Götherström A, Girdland-Flink L. 2017. Genomic Analyses of Pre-European Conquest Human Remains from the Canary Islands Reveal Close Affinity to Modern North Africans. Curr Biol 27:3396–3402.e5.

104. Rohland N, Harney E, Mallick S, Nordenfelt S, Reich D. 2015. Partial uracil-DNA-glycosylase treatment for screening of ancient DNA. Philos Trans R Soc Lond B Biol Sci 370:20130624.

105. Rohland N, Hofreiter M. 2007. Ancient DNA extraction from bones and teeth. Nat Protoc 2:1756–1762.

106. Roymans N, Derks T, Heeren S. 2020. Roman Imperialism and the Transformation of Rural Society in a Frontier Province: Diversifying the Narrative. Britannia 51:265–294.

107. Saag L, Laneman M, Varul L, Malve M, Valk H, Razzak MA, Shirobokov IG, Khartanovich VI, Mikhaylova ER, Kushniarevich A, Scheib CL, Solnik A, Reisberg T, Parik J, Saag L, Metspalu E, Rootsi S, Montinaro F, Remm M, Mägi R, D’Atanasio E, Crema ER, Díez-Del-Molino D, Thomas MG, Kriiska A, Kivisild T, Villems R, Lang V, Metspalu M, Tambets K. 2019. The Arrival of Siberian Ancestry Connecting the Eastern Baltic to Uralic Speakers further East. Curr Biol 29:1701–1711.e16.

108. Saag L, Varul L, Scheib CL, Stenderup J, Allentoft ME, Saag L, Pagani L, Reidla M, Tambets K, Metspalu E, Kriiska A, Willerslev E, Kivisild T, Metspalu M. 2017. Extensive Farming in Estonia Started through a Sex-Biased Migration from the Steppe. Curr Biol 27:2185–2193.e6.

109. Sánchez-Quinto F, Malmström H, Fraser M, Girdland-Flink L, Svensson EM, Simões LG, George R, Hollfelder N, Burenhult G, Noble G, Britton K, Talamo S, Curtis N, Brzobohata H, Sumberova R, Götherström A, Storå J, Jakobsson M. 2019. Megalithic tombs in western and northern Neolithic Europe were linked to a kindred society. Proc Natl Acad Sci U S A 116:9469–9474.

110. Saupe T, Montinaro F, Scaggion C, Carrara N, Kivisild T, D’Atanasio E, Hui R, Solnik A, Lebrasseur O, Larson G, Alessandri L, Arienzo I, De Angelis F, Rolfo MF, Skeates R, Silvestri L, Beckett J, Talamo S, Dolfini A, Miari M, Metspalu M, Benazzi S, Capelli C, Pagani L, Scheib CL. 2021. Ancient genomes reveal structural shifts after the arrival of Steppe-related ancestry in the Italian Peninsula. Curr Biol 31:2576–2591.e12.

111. Scheidel W. 2019. Escape from Rome. Princeton University Press.

112. Scheidel W. 2015. Orbis: The Stanford Geospatial Network Model of the Roman World. doi:10.2139/ssrn.2609654

113. Schiffels S, Haak W, Paajanen P, Llamas B, Popescu E, Loe L, Clarke R, Lyons A, Mortimer R, Sayer D, Tyler-Smith C, Cooper A, Durbin R. 2016. Iron Age and Anglo-Saxon genomes from East England reveal British migration history. Nat Commun 7:10408.

114. Schroeder H, Margaryan A, Szmyt M, Theulot B, Włodarczak P, Rasmussen S, Gopalakrishnan S, Szczepanek A, Konopka T, Jensen TZT, Witkowska B, Wilk S, Przybyła MM, Pospieszny Ł, Sjögren K-G, Belka Z, Olsen J, Kristiansen K, Willerslev E, Frei KM, Sikora M, Johannsen NN, Allentoft ME. 2019. Unraveling ancestry, kinship, and violence in a Late Neolithic mass grave. Proc Natl Acad Sci U S A 116:10705–10710.

115. Schuenemann VJ, Peltzer A, Welte B, van Pelt WP, Molak M, Wang C-C, Furtwängler A, Urban C, Reiter E, Nieselt K, Teßmann B, Francken M, Harvati K, Haak W, Schiffels S, Krause J. 2017. Ancient Egyptian mummy genomes suggest an increase of Sub-Saharan African ancestry in post-Roman periods. Nat Commun 8:15694.

116. Séguy I. 2019. Current trends in Roman demography and empirical approaches to the dynamics of the Limes populationsFinding the Limits of the Limes. Springer, Cham. pp. 23–41.

117. Sikora M, Seguin-Orlando A, Sousa VC, Albrechtsen A, Korneliussen T, Ko A, Rasmussen S, Dupanloup I, Nigst PR, Bosch MD, Renaud G, Allentoft ME, Margaryan A, Vasilyev SV, Veselovskaya EV, Borutskaya SB, Deviese T, Comeskey D, Higham T, Manica A, Foley R, Meltzer DJ, Nielsen R, Excoffier L, Mirazon Lahr M, Orlando L, Willerslev E. 2017. Ancient genomes show social and reproductive behavior of early Upper Paleolithic foragers. Science 358:659–662.

118. Skoglund P, Malmström H, Omrak A, Raghavan M, Valdiosera C, Günther T, Hall P, Tambets K, Parik J, Sjögren K-G, Apel J, Willerslev E, Storå J, Götherström A, Jakobsson M. 2014. Genomic diversity and admixture differs for Stone-Age Scandinavian foragers and farmers. Science 344:747–750.

119. Skoglund P, Malmström H, Raghavan M, Storå J, Hall P, Willerslev E, Gilbert MTP, Götherström A, Jakobsson M. 2012. Origins and genetic legacy of Neolithic farmers and hunter-gatherers in Europe. Science 336:466–469.

120. Skourtanioti E, Erdal YS, Frangipane M, Balossi Restelli F, Yener KA, Pinnock F, Matthiae P, Özbal R, Schoop U-D, Guliyev F, Akhundov T, Lyonnet B, Hammer EL, Nugent SE, Burri M, Neumann GU, Penske S, Ingman T, Akar M, Shafiq R, Palumbi G, Eisenmann S, D’Andrea M, Rohrlach AB, Warinner C, Jeong C, Stockhammer PW, Haak W, Krause J. 2020. Genomic History of Neolithic to Bronze Age Anatolia, Northern Levant, and Southern Caucasus. Cell 181:1158–1175.e28.

121. Symonds M. 2017. Protecting the Roman Empire: Fortlets, Frontiers, and the Quest for Post-Conquest Security. Cambridge University Press.

122. Unterländer M, Palstra F, Lazaridis I, Pilipenko A, Hofmanová Z, Groß M, Sell C, Blöcher J, Kirsanow K, Rohland N, Rieger B, Kaiser E, Schier W, Pozdniakov D, Khokhlov A, Georges M, Wilde S, Powell A, Heyer E, Currat M, Reich D, Samashev Z, Parzinger H, Molodin VI, Burger J. 2017. Ancestry and demography and descendants of Iron Age nomads of the Eurasian Steppe. Nat Commun 8:14615.

123. Valdiosera C, Günther T, Vera-Rodríguez JC, Ureña I, Iriarte E, Rodríguez-Varela R, Simões LG, Martínez-Sánchez RM, Svensson EM, Malmström H, Rodríguez L, Bermúdez de Castro J-M, Carbonell E, Alday A, Hernández Vera JA, Götherström A, Carretero J-M, Arsuaga JL, Smith CI, Jakobsson M. 2018. Four millennia of Iberian biomolecular prehistory illustrate the impact of prehistoric migrations at the far end of Eurasia. Proc Natl Acad Sci U S A 115:3428–3433.

124. van den Brink ECM, Beeri R, Kirzner D, Bron E, Cohen-Weinberger A, Kamaisky E, Gonen T, Gershuny L, Nagar Y, Ben-Tor D, Sukenik N, Shamir O, Maher EF, Reich D. 2017. A Late Bronze Age II clay coffin from Tel Shaddud in the Central Jezreel Valley, Israel: context and historical implications. Levantina 49:105–135.

125. Veeramah KR, Rott A, Groß M, van Dorp L, López S, Kirsanow K, Sell C, Blöcher J, Wegmann D, Link V, Hofmanová Z, Peters J, Trautmann B, Gairhos A, Haberstroh J, Päffgen B, Hellenthal G, Haas-Gebhard B, Harbeck M, Burger J. 2018. Population genomic analysis of elongated skulls reveals extensive female-biased immigration in Early Medieval Bavaria. Proc Natl Acad Sci U S A 115:3494–3499.

126. Villalba-Mouco V, van de Loosdrecht MS, Posth C, Mora R, Martínez-Moreno J, Rojo-Guerra M, Salazar-García DC, Royo-Guillén JI, Kunst M, Rougier H, Crevecoeur I, Arcusa-Magallón H, Tejedor-Rodríguez C, García-Martínez de Lagrán I, Garrido-Pena R, Alt KW, Jeong C, Schiffels S, Utrilla P, Krause J, Haak W. 2019. Survival of Late Pleistocene Hunter-Gatherer Ancestry in the Iberian Peninsula. Curr Biol 29:1169–1177.e7.

127. Wang C-C, Reinhold S, Kalmykov A, Wissgott A, Brandt G, Jeong C, Cheronet O, Ferry M, Harney E, Keating D, Mallick S, Rohland N, Stewardson K, Kantorovich AR, Maslov VE, Petrenko VG, Erlikh VR, Atabiev BC, Magomedov RG, Kohl PL, Alt KW, Pichler SL, Gerling C, Meller H, Vardanyan B, Yeganyan L, Rezepkin AD, Mariaschk D, Berezina N, Gresky J, Fuchs K, Knipper C, Schiffels S, Balanovska E, Balanovsky O, Mathieson I, Higham T, Berezin YB, Buzhilova A, Trifonov V, Pinhasi R, Belinskij AB, Reich D, Hansen S, Krause J, Haak W. 2019. Ancient human genome-wide data from a 3000-year interval in the Caucasus corresponds with eco-geographic regions. Nat Commun 10:590.

128. Zalloua P, Collins CJ, Gosling A, Biagini SA, Costa B, Kardailsky O, Nigro L, Khalil W, Calafell F, Matisoo-Smith E. 2018. Ancient DNA of Phoenician remains indicates discontinuity in the settlement history of Ibiza. Sci Rep 8:17567–17515.

129. Žegarac A, Winkelbach L, Blöcher J, Diekmann Y, Krečković Gavrilović M, Porčić M, Stojković B, Milašinović L, Schreiber M, Wegmann D, Veeramah KR, Stefanović S, Burger J. 2021. Ancient genomes provide insights into family structure and the heredity of social status in the early Bronze Age of southeastern Europe. Sci Rep 11:10072.

