## Appendix 1 - Archaeological site information for "Stable population structure in Europe since the Iron Age, despite high mobility"

**Archaeological Sites** 3
Necropole Orientale, Algeria 4
Beniamin, Armenia 6
Klosterneuburg, Austria 11
Wels, Austria 15
Beli Manastir, Croatia 17
Gardun (Tilurium), Croatia 18
Novo Selo-Bunje, Croatia 22
Mirine-Fulfinum, Omišalj, Croatia 23
Osijek, Croatia 26
Ščitarjevo, Croatia 28
Sipar - Umag, Croatia 30
Sisak-Pogorelec, Croatia 33
Trogir-Dragulin, Croatia 36
Trogir-Policija, Croatia 38
Velić, Croatia 39
Zadar, Croatia 40
Metz, France 44
Sarrebourg, France 47
Vendeuil Caply Les Marmousets, France 49
Hassleben, Germany 50
Isola Sacra, Italy 52
Palazzo della Cancelleria, Italy 55
Urbino Bivio, Italy 57
Chhim, Lebanon 59
Ej-Jaouze, Lebanon 61
Bailuliai, Lithuania 63
Marvelė, Lithuania 64
Doclea Bjelovine, Montenegro 70
Weklice, Poland 72
Conímbriga, Portugal 75
Miroiço, Portugal 76
Monte da Nora, Portugal 78
Sant’ Imbenia, Sardinia 80

| | |
|---|---|
| Beška-Brest, Serbia | 83 |
| Naissus, Serbia | 85 |
| Sirmium, Serbia | 87 |
| Sviloš-Kruševlje, Serbia | 92 |
| Viminacium, Serbia | 94 |
| Bytča-Hrabové, Slovakia | 97 |
| Mikušovce, Slovakia | 99 |
| Tesárske Mlyňany, Slovakia | 101 |
| Zohor, Slovakia | 104 |
| Emona, Slovenia | 106 |
| Tell Masaikh, Syria | 109 |

### Archaeological Sites

#### Necropole Orientale, Algeria

**Collaborators**: Yahia Mehdi Seddik Cherifi, Selma Amrani, Mohamed el Mostefa Filah
**Coordinates**: 36.1898, 5.4108
**Individuals** (n = 3): R10760, R10766, R10770

<u>The Eastern Necropolis of Sitifis</u>

Sitifis, an inland town in north-eastern Algeria (North Africa; today the town of Sétif), was known to be founded in 97 A.D. by the Romans (Nerva's reign) as a colony for Roman veterans. It grew to become the capital of *Mauretania Sitifensis*, a newly erected province from the eastern part of Mauretania Caesariensis, under the Emperor Diocletian (293 A.D.).

The Eastern Necropolis of Sitifis, from the Numido-Roman period (1st-5th centuries C.E.), is located east of Sétif (Algeria), with 352 recognized and excavated graves (by R. Guéry; between 1959-1967). The site reported three occupational phases: burial (early graves, known to be older than the Sitifis colony), then cremation, and finally return to burial (late graves).

From the 214 burial graves, both early and late, 228 individuals were identified without any anthropological study. However, the approximate age was determined by Dr. Jean Leuret, a former professor of osteology at the Faculty of Medicine of Paris and at the time surgeon-head of the hospital of Sétif.
The current corpus comprises 132 individuals, the missing individuals since 1971 are somewhere between France and Algeria. In this collection, we clearly have distinguished 68 immatures and 64 adults, after applying different methods for skeletal age estimation.
The additional singularity of the population of this necropolis is to be categorized as native: the Numidians (Berbers).

The bioarchaeology of the dentoskeletal collection was recently investigated and these contributions have been communicated in the Safa conference 2018 (Canada) and publications are under review, as part of the PhD project of Selma Amrani.

**<u>Literature</u>**:

Nacéra Benseddik, Autels votifs de la région de Sétif: païens ou chrétiens?, Monuments funéraires, institutions autochtones en Afrique du Nord antique et médiévale, VIe Colloque International sur L'Histoire et l'Archéologie de l'Afrique du Nord, Pau, 1993, C.T.H.S. [1995], p. 179-186.

Serge Lancel et Omar Daoud. *L'Algérie antique : De Massinissa à saint Augustin*, Place des Victoires, 2008 (ISBN 9782844591913)

B. A. R. Guéry, La nécropole orientale de Sititis (Sétif, Algérie), fouilles de 1966-1967 ,1985, 1988.

P. A. Février, R. Guéry, Les rites funéraires de la nécropole orientale de Sétif, Antiquités 15 (1) (1980) 91–124.

| sample_id | sex (genetic) | date lower (95% CI) | date upper (95% CI) | date inference | burial_id | excluded |
|---|---|---|---|---|---|---|
| R10760 | F | 38 calBCE | 109 calCE | 14C | Necropole Orientale_Aos785 | |
| R10766 | M | 63 calCE | 204 calCE | 14C | Necropole Orientale_Aos703 | |
| R10770 | M | 81.5 calCE | 210 calCE | 14C | Necropole Orientale_Aos697 | Yes (Contam.) |

#### Beniamin, Armenia

**Collaborators:** Anahit Khudaverdyan, Hamazasp Khachatryan, Pavel Avetisyan
**Coordinates**: 40.6885, 43.8493
**Individuals** (n = 18): R11535, R11536, R11538, R11540, R11541, R11542, R11543, R11544, R11545, R11546, R11651, R11653, R11655, R11668, R11675, R11678, R11713, R11714

The Beniamin necropolis

Beniamin is located in Armenia in the Shirak province, and was investigated by archaeologists F.I. Ter-Martirosov, H.H. Khachatryan, L.G. Yeganyan, and by physical anthropologist A. Yu. Khudaverdyan. Between 1989 and 1999 the Institute of Archaeology and Ethnography, National Academy of Science, Republic of Armenia and the Shirak Museum conducted a joint excavation at Beniamin. From 1999-2007, excavations continued by the Armenian-French expedition. Four stages of occupation dating from the 5th to 1st centuries BC were identified, of which two – Periods Ia and Ib – date to the Achaemenid epoch. The site is composed of an Achaemenid-era palace on a hill and a Hellenistic lower settlement. Deposits from the first period of use, dated to early Achaemenid occupation, have yielded few remains since they were not extensively excavated and the pottery assemblage has not been published. The Beniamin anthropological collection is one of the few large collections in Armenia. Data from excavations suggest that a small farming community occupied the site from at least 1st century BC − 3rd century AD. A total of 350 burials were recovered from Beniamin during field seasons from 1989 to 2007. Grave associations were generally uncommon, and when present, usually consisted of small personal items like tupu pins, goods including metalwork, pottery, etc. Individuals were placed in extended positions. The sample consisted of 165 individuals: 63 from females, 44 from males, 57 from subadults. One adult individual was of undetermined sex (very fragmented skull, absent pelvic bones). The detailed analysis of the craniological, cranioscopic (non-metric traits) and odontological materials from the Beniamin allows to explain not only the complicated anthropological composition of the population but also to discover the reason for the anthropological and ethnic non-homogeneity in the population of the ancient age. Intragroup analysis revealed two groups within the population (Khudaverdyan, 2012a,b, 2018). Intergroup analysis showed the tribes of the Chernyakhov culture and the population of the Middle East to be closest to the Beniamin male population. The female population was closest to the Scythians of Ukraine and Crim (Khudaverdyan et al., 2020). This confirms the fact of sustainable, constant migration flow to the territory of the Armenian Highlands. Other contributions have been published, exploring demography and artificial deformation of skulls.

**Literature**:

Khudaverdyan, A.Yu. 2010. Pattern of disease in three 1st century BC – 3rd century AD burials from Beniamin, Vardbakh and the Black Fortress I, Shiraksky plateau (Armenia). Journal of Paleopathology (Italy), 22, 15–41.

Khudaverdyan, A.Yu. 2011. Artificial modification of skulls and teeth from ancient burials in Armenia. Anthropos, 106 (2), 602–609.

Khudaverdyan, A.Yu. 2012a. Armenia in the Eurasian ethnic context of late classical antiquity: craniometric evidence. Archaeology, Ethnography & Anthropology of Eurasia, 3 (51), 138-148.

Khudaverdyan, A.Yu. 2012b. Nonmetric cranial variation in human skeletal remains from Armenian Highland: microevolutionary relations and intergroup analysis. European Journal of Anatomy (Spanish), 16 (2), 134-149.

Khudaverdyan, A.Yu. 2014. Les inhumations de la cimetières de la plaine Chirak (Arménie), approche biologique et sociale. Etnoantropološki problem, 9 (1), 219-242.

Khudaverdyan, A.Yu. 2015a. Paleodemography of the Armenian population in the LateAantiquity (based on the materials of ancient monuments of the Shirak plain). Scientific Herald of the Volgograd branch of RANEPA, 4, 56-62.

Khudaverdyan, A.Yu. 2015b. Palaeopathology of human remains of the 1st century BC–3rd century AD from Armenia (Beniamin, Shirakavan I). Anthropological Review, 78 (2), 213-228.

Khudaverdyan, A.Yu. 2018. Illuminating the processes of microevolution: A bioarchaeological analysis of dental non-metric traits from Armenian Highland. HOMO - Journal of Comparative Human Biology, 69, 304–323

Khudaverdyan, A.Yu., Hovhanisyan, A.A., Yengibaryan, A.A., Matevosyan, R.Sh., Qocharyan, G.G., Palanjan, R.S., Eganyan, L.G., Khachatryan, H.H. 2020. Population of the Armenian Higlands in the age of Antiquity(according of anthropological materials of urban and rural settlements). Bulletin of archeology, anthropology and ethnography, 1 (48), 96-115

On the establishment and inhabitation of Beniamin settlement
Hamazasp Khachatryan

The classical period settlement of Beniamin has been founded as an administrative and religious center in the 5th century BC. This is evidenced by the discovered palace buildings belonging to the 5-2 and 1st centuries BC. The residence has no earlier layer than 5th century BC. About 300 tombs dating back to the 2nd century BC and 3, 4th centuries AD have been excavated in and around the upper layer of the settlement. The residence is based in a formerly uninhabitated area, which we believed to be populated by relocating people from other settlements.

About 3 km northeast of the Beniamin site, we have been studying the complex of Azatan for about ten years. The research was together with Martin Luther University of Halle, headed by H. Khachatryan, Prof. A. Furtwangler. The site consists of a castle, a settlement and 3 cemeteries surrounding them. The castle and the settlement are located on the heights. Life here begins in the Early Iron Age and continues until the Hellenistic period. Approximately in the 5th century BC a small part of the settlement goes down to the valley, then life stops in that part. This can be explained by the relocation of the population to the newly established Beniamin site. However, a certain population remains, and a small number of Hellenistic period structures dating to the 2nd BC - 1st centuries AD emerges around the castle. Numerous granaries of various sizes have been discovered in and around the castle. We believe that the area of Azatan castle is being turned into food warehouses of the Beniamin royal center and the remaining small part of the population was responsible for the upkeep and maintenance of these warehouses. About 30 tombs of 12th -1st centuries BC have been excavated in the cemeteries of Azatan. One Achaemenid and three Hellenistic period burials have been uncovered among the above mentioned 30 tombs.

In our opinion, the Beniamin population was formed and changed in the following 3 stages:

- 1st stage
  With the population relocated from Azatan settlement in the 5th century BC.

- 2nd stage
  When the palace complex of the Achaemenid period and the monuments of King Artashes (189-160 BC) were destroyed, the large torus and bell-shaped Achaemenid column bases were used as building materials. A plain building of a different layout appears on the site, with different construction equipment that

could have existed during the Armenian-Parthian war and the invasion of another ethnic group invading here at the end of the 2nd century BC.

- 3rd stage
  As a result of allocating Shirak as a feud to the Kamsarakan tribe and settling them here, who did not reconcile with the Sasanian dynasty during the second half of 2nd century AD. These events are evidenced by historical sources.

| sample_id | sex (genetic) | date lower (95% CI) | date upper (95% CI) | date inference | burial_id | excluded | notes |
|---|---|---|---|---|---|---|---|
| R11535 | M | 799 calBCE | 772 calBCE | 14C | Beniamin_bur. 11 | | |
| R11536 | M | 1045.5 calBCE | 924 calBCE | 14C | Beniamin_bur. 17 (right petrous) | | REL_1ST of 11655 |
| R11538 | F | 431 calCE | 545 calCE | 14C | Beniamin_bur. 203 | | |
| R11540 | F | 403 calBCE | 378.5 calBCE | 14C | Beniamin_bur. 36 | | |
| R11541 | M | 42.5 calBCE | 61 calCE | 14C | Beniamin_bur. 47 | | |
| R11542 | M | 431 calCE | 545 calCE | 14C | Beniamin_bur. 20/1 | | |
| R11543 | F | 421.5 calCE | 538.5 calCE | 14C | Beniamin_bur. 214 | | |
| R11544 | F | 419 calCE | 538 calCE | 14C | Beniamin_bur. 30 | | |
| R11545 | F | 1396 calBCE | 1259.5 calBCE | 14C | Beniamin_bur. 5/1 | | |
| R11546 | M | 154.5 calBCE | 0.5 calCE | 14C | Beniamin_bur. 60 | | |
| R11651 | M | 403 calBCE | 545 calCE | SITE | Beniamin_bur. 130 | | |
| R11653 | M | 403 calBCE | 545 calCE | SITE | Beniamin_bur. 15/1 | | |
| R11655 | F | 403 calBCE | 545 calCE | SITE | Beniamin_bur. 17 | Yes (Contam. / Related) | REL_1ST of 11536 |
| R11668 | M | 1211.5 calBCE | 1053 calBCE | 14C | Beniamin_bur. 21 | | |
| R11675 | M | 1490.5 calBCE | 1323.5 calBCE | 14C | Beniamin_bur. 7 | | |
| R11678 | F | 403 calBCE | 545 calCE | SITE | Beniamin_bur. 96 | | |
| R11713 | M | 403 calBCE | 545 calCE | SITE | Beniamin_bur. 1 | | |
| R11714 | F | 403 calBCE | 545 calCE | SITE | Beniamin_bur. ??/4 | | |

### Klosterneuburg, Austria

**Collaborators**: Maria Teschler-Nicola, Christine Neugebauer-Maresch
**Coordinates**: 48.3099, 16.3238
**Individuals** (n = 6): R10654, R10656, R10657, R10658, R10659, R10660

The necropolis of the Roman imperial time at Klosterneuburg (Lower Austria)

The most western situated auxiliary castle of the province of Pannonia was located at today's "Upper Town" of Klosterneuburg (Lower Austria), a city in close vicinity of Vienna (ca. 13 km NW of Vienna). The castle Klosterneuburg was a fortress within the Roman fortification line of the Donaulimes, a part of the Roman frontier system. In its early time, it was used as cohort-camp for auxiliary-forces; from the 2nd century on it was a cavalry base (Aelische cohort of archers: *Cohors I Aelia Sagittariorum [Severiana] [milliaria] [equitata*]). The fortification's function was the control of the Danube-passage and the Limes-road. It was in use until the 5th century A.D. In the periphery of the fort a vicus and necropolis were situated. Archaeological findings include, among others, parts of the dress, weapons, ceramics, and Terra sigillata, fragments of a scale armour (Neugebauer-Maresch & Neugebauer 1985/86, Neugebauer-Maresch & Neugebauer 1993, Neugebauer-Maresch, Neugebauer & Ubl 1998, sub-construction of a street (Neugebauer 2000), and a camp-bath (Ubl 1995). In the 4th and 5th century A.D. the Late Antique population of Klosterneuburg was not only composed of Romans but was already strongly influenced by Germanic immigrants. The burials did not reveal a clear separation between military and civilians.
The first graves of the Roman imperial time were already discovered at the end of 19th century; later, during construction works in the 1960s and 1970s further archaeological findings appeared and were secured and recovered by Ubl (Ubl 1976) and Johannes-Wolfgang Neugebauer (Neugebauer & Neugebauer 1985/86) from the Federal Monument Office. Interestingly, among the burials was a "soldier's grave" identified by particular items, e.g. belt fittings that were correlated with "soldier's graves in the area between Britain and North Africa" (Neugebauer-Maresch & Neugebauer 1986:320). The authors concluded further that this male, buried around 200 A.D. might have been a member of the *Cohors I Aelia sagittariorium* (Neugebauer-Maresch and Neugebauer 1986:320).
Because continuing construction and remodelling works within this archaeologically promising area were carried out, the Federal Monument Office started in 1983 with systematic rescue excavations. Up to 1992 several subranges of the necropolis were exposed, and more than 160 cremation and inhumation graves, ditches, enclosures, as well as segments of a road construction ("Römerstrasse") were uncovered. Most of the inhumation graves and objects date between 3rd and 5th century after Christ.

The burials include single and multiple, familial burials (with primary and secondary interments), the corpses were either enwrapped in linen or placed in a wooden coffin, in some cases elaborate constructions such as chambers made of stone or bricks were found. Although some graves were found to be wealthy, equipped with grave goods, jewellery, dress parts, weapons and tools, clay- and glass-jars, the majority was plundered in ancient time.

**Verf. 44, Fn. 2 (Sample ID R10654)**, Buchberggasse, excavation campaign 1992:
N-S oriented grave, fragments of bricks and Laterculi, S-N-oriented burial in extended supine position, right arm elongated and close to the body, left arm bent. - No grave goods.
Well preserved skeleton**:** hand and foot bones missing; male, age at death ca. 30-40 years; pathological alterations: small osteoma at the occipital bone, signs of chronic inflammation at the palate.

**Verf. Fn. 779/a (Sample ID R10659)**, Jahngasse, excavation campaign 1990:
NW-SO oriented oval grave pit, SO-oriented burial in a slightly crouched position, both arms in front of the body in crouched position. – No grave goods.
Incomplete preserved skeleton**:** Neonate.

**Verf. Fn. 780/a (Sample ID R10660)**, Jahngasse, excavation campaign 1990:
NS oriented oval grave pit (ca.58x38 cm), at the NW-sector overlapping with Verf. 806, burial in crouched position, corpse placed at its left side, left arm elongated in front of the body, forearm distorted, right arm at the side. – No grave goods.
Incomplete preserved skeleton**:** Neonate.

**Verf. Fn. 806/10 (Sample ID R10657)**, „Male below a horse“, Jahngasse, excavation campaign 1990:
S-N-oriented burial (grave pit 190x78 cm) in extended supine position, arms alongside the body. – No grave goods, the deceased was buried below a horse; a cremation burial was recovered above.
Complete and well-preserved skeleton**:** male, age at death between 25 and 35 years; pathological alterations: remodelled (old) fractures of left $5^{th}$ metacarpal bone; healed fracture of the inferior ramus of the left pubic bone; deformed right distal metatarsal I-phalangeal joint (trauma and/or severe arthrosis, hallux rigidus); left phalanges of the foot with ankylosis and arthritic alterations; right thumb with severe destructions, newly build bone and arthritic changes (archer?); remodelled subperiosteal bony appositions caused by chronic stress (“striae”) at the femora and tibiae; $5^{th}$ lumbar vertebra asymmetrical formed; partial spina bifida, habitual changes at the femoral neck caused by horseback riding (“Reiterfacette”).

**Verf. Fn. 831/1, (Sample ID R10658)**, Jahngasse, excavation campaign 1990:
SO-NW-oriented burial in a small grave pit (145x38 cm), burial in extended position with slightly bended legs, right arm alongside the body, left arm above the pelvis and bended. – Sherds recovered from the burial area.
Complete and well-preserved skeleton**:** age at death ca. 9-10 years (conspicuous discrepancy between age at death based on long bone dimension and age at death based on the mineralisation and development of the teeth; length of long bones would imply an age of death of ca. 6 years).

**Verf. Fn. 849/1 (Sample ID R10656)**, Jahngasse, excavation campaign 1990:
NO-SW-oriented grave (195x70 cm), burial in prone position, right arm bent and placed alongside the body, left arm and hand extended alongside the body (in pronation position), right leg in lateral position and slightly bent, left leg in extended position. – Only sherds collected from the grave shaft.
Well preserved skeleton**:** male, age at death ca. 25-35 years; pathological alterations: striae caused by chronic periosteal inflammation at both femora, tibiae, and fibulae; one fractured rib with signs of remodelling, inflammation, and “pseudo-joint” formation; other ribs reveal “erosions” of varying depth at the collum; conspicuous erosions anteriorly at the lumbar and thoracic vertebrae may be the result of aneurysms of the descending aorta which is closely attached to the vertebral column. These structural changes are most likely of arteriosclerotic etiology.

| sample_id | sex (genetic) | date lower (95% CI) | date upper (95% CI) | date inference | burial_id | excluded |
|---|---|---|---|---|---|---|
| R10654 | M | 258.5 calCE | 407 calCE | 14C | Klosterneuburg Verf. 44, Fn. 2a | |
| R10659 | M | 26 calCE | 126 calCE | 14C | Fn. 779/a | |
| R10660 | F | 26 calCE | 407 calCE | SITE | Fn. 780/a | |
| R10657 | M | 26 calCE | 126 calCE | 14C | Klosterneuburg-Jahnstr. Fn. 806/10 | |
| R10658 | F | 26 calCE | 407 calCE | SITE | Klosterneuburg Fn. 831/1 | Yes (Contam.) |
| R10656 | M | 26 calCE | 407 calCE | SITE | Klosterneuburg-Jahng. Fn. 849/1 | |

**Literature**:

Neugebauer, J.-W. 2000. Fundberichte aus Österreich 39, 19.

Neugebauer, J.-W., Neugebauer-Maresch, Ch. 1985/86. Fundberichte aus Österreich 24/25, 291-292.

Neugebauer-Maresch, Ch., Neugebauer, J.-W. 1986. Ein Friedhof der römischen Kaiserzeit in Klosterneuburg. Die Rettungsgrabungen des Bundesdenkmalamtes in den Jahren 1983-84 im Bereich des evangelischen Pfarramtes. ArchA 70, 317-384.

Neugebauer-Maresch, Ch., Neugebauer, J.-W. 1993. Das Gräberfeld der römischen Kaiserzeit und beginnenden Völkerwanderungszeit in Klosterneuburg-Buchberggasse, in: Klosterneuburg, Geschichte und Kultur, Band 1: Die Stadt. Stiftung Klosterneuburg Verlag, Klosterneuburg-Wien 1993, 97-120.

Neugebauer-Maresch, Ch., Neugebauer, J.-W., Ubl, H. 1998. Fundberichte aus Österreich 37, 22–25.

Ubl, H. 1976. Fundberichte aus Österreich 15, 266.

Ubl, H. 1995. Fundberichte aus Österreich 34, 21.

https://de.wikipedia.org/wiki/Kastell_Klosterneuburg#cite_ref-15, accessed on January 3rd, 2021.

#### Wels, Austria

**Collaborator**: Sylvia Kirchengast
**Coordinates**: 48.1654, 14.0366
**Individuals** (n = 5): R10665, R10666, R10667, R10668, R10670

The cemetery of the Roman town Ovilava
Sylvia Kirchengast, Renate Miglbauer, Michael Greisinger

The Roman town of *Ovilava* (*today* Wels in Upper Austria) was founded in the Roman Province *Noricum* at the *intersection* of two important roads in the last third of the 1st century AD. At the area of *Ovilava* the east-west running trunk road that connected the Roman towns *Lauriacum* (today Enns) and *Juvavum* (today Salzburg) and the road that led from *Aquilea* in northern Italy over the Alps to *Boiodurum* (today Passau) crossed each other. Consequently, *Ovilava* gained quickly in importance as commercial and administrative center. Around the years 120 AD *Ovilava* received Roman city rights by the Roman emperor Hadrian (76-138 AD) and *Ovilava* became the *municipium Aelium Ovilava*. During the second half of the 2$^{nd}$ century AD *Ovilava* remained untroubled by the Marcomannic wars and was finally raised to *Colonia Aurelia Antoniniana* by the Roman emperor Caracalla (211-217 AD). The civil population of *Ovilava* consisted mainly of traders, craftsmen and civil servants, which originated from Norcium but some of them may be migrated from other parts of the Roman empire to this area near the Danube limes. The civil inhabitants of *Ovilava* were buried in the necropolis outside the city along the road from *Lauriacum* to *Juvavum*. This cemetery was in use up to early medieval times. Today the area of the Roman cemeteries is located around the main train station and is therefore part of the densely built city of Wels. Consequently, excavations are only possible in the context of construction projects within the city. The construction of an underground car park near the main train station resulted in an emergency excavation of large parts of the Roman cemetery between the years 2004 and 2006. During this campaign, more than 220 burials (inhumations and cremations) could be excavated. Grave goods and tombs indicate that wealthy people but also commoners were buried in this area. According to the grave goods, first of all coins, the burials dated mainly in the 2$^{nd}$ and 3$^{rd}$ century AD. Only a few burials are attributed to Bavarian people dating in early medieval times. More than half of the burials consisted of cremations, the others were inhumations. The skeletal remains have been stored at the Archeological Department of the Stadtmuseum Wels. In 2019 the skeletons were transferred to the Department of Evolutionary Anthropology, University of Vienna, where the bioarcheological analysis is currently in progress.

**Literature:**

Miglbauer R. (2007) Die letzte Reise Ausgrabungen im östlichen Gräberfeld von Ovilava/Wels. Mitteilungen aus dem Stadtmuseum Wels 117.

| sample_id | sex (genetic) | date lower (95% CI) | date upper (95% CI) | date inference | burial_id |
|---|---|---|---|---|---|
| R10665 | M | 660 calCE | 774.5 calCE | 14C | Wels(Ovilava)_Gr. 34 |
| R10666 | F | 657 calCE | 774.5 calCE | 14C | Wels(Ovilava)_Gr. 54 |
| R10667 | M | 124 calCE | 217 calCE | 14C | Wels(Ovilava)_Gr. 59 |
| R10668 | M | 131 calCE | 245.5 calCE | 14C | Wels(Ovilava)_Gr. 13 |
| R10670 | M | 216.5 calCE | 325 calCE | 14C | Wels(Ovilava)_Gr. 65 |

### Beli Manastir, Croatia

**Collaborators**: Mario Novak, Dženi Los, Josip Burmaz
**Coordinates**: 45.7729, 18.6108
**Individuals** (n = 1): R3542

Beli Manastir - Popova zemlja, Croatia

The site is located in the eastern part of Croatia in the Baranja region. Large-scale salvage excavations conducted in 2014/2015 revealed structures and a very rich concentration of artifacts belonging to multiple periods from the Early Neolithic to the Roman period. Most of the Neolithic site consists of large pit houses, fireplaces and stoves, and many channels, as well as almost 40 inhumation burials. Beside the prehistoric finds, a Roman period layer with two rectangular brick furnaces and two inhumation burials was also identified and excavated.
Burial 22 was constructed of roof tiles (*tegulae*) and contained a completely preserved skeleton of a subadult 9-12 months old. The individual way lying on the back with the arms and legs outstretched along the body. A small ceramic vessel was located by the legs in the left lower corner of the burial. The only recorded pathology was the presence of active cribra orbitalia in both orbits.

**Literature**:

Los Dž. 2020. Results of the archaeological excavations of the site AN 2 Beli Manastir – Popova zemlja. Annales Instituti Archaeologici, 15: 90-102.

| sample_id | sex (genetic) | date lower (95% CI) | date upper (95% CI) | date inference | burial_id |
|---|---|---|---|---|---|
| R3542 | M | 255.5 calCE | 405 calCE | 14C | BELI MANASTIR - POPOVA ZEMLJA; G 22; left temporal |

#### Gardun (Tilurium), Croatia

**Collaborators**: Mario Novak, Iva Kaić, Mirna Cvetko, Mirjana Sanader, Domagoj Tončinić, Mario Šlaus
**Coordinates**: 43.6076, 16.7143
**Individuals** (n = 3): R3543, R3544, R3545

Archeological site Gardun (*Tilurium*) near Trilj

The legionary fortress Tilurium was situated on the territory of the present-day village Gardun, on a dominating and strategic position on the right bank of the Cetina river, controlling the surrounding fields and plateaus, as well as the crossing over the Cetina in the area of the present-day town of Trilj. Tilurium and its immediate surroundings have been attracting attention for more than 150 years due to the archaeological finds that were purchased or donated to several different museums and collections (Sanader and Tončinić, 2010, p. 33, 2013, p. 411; Tončinić and Vukov, 2019, p. 39; Tončinić and Cvetko, 2021a, p. 7). The most prominent finds are tombstones and other inscriptions bearing testimony to the presence of various Roman military units at Tilurium (Sanader and Tončinić, 2010, pp. 34–37, 2013, pp. 412–413; Tončinić and Vukov, 2019, p. 39; Tončinić and Cvetko, 2021a, p. 7, 2021b, p. 202; Tončinić and Pavlović, 2021, p. 56). The most represented unit is *Legio VII*, or *Legio VII Claudia pia fidelis*. The oldest reliably dated stone monuments of the *Legio VII* in Dalmatia are the so-called *Tabulae Dolabellae*, dating to the year 16/17 A.D. According to paleographic parallels, some tombstones of the *Legio VII* from Tilurium can be dated between the 1st and 3rd decade of the 1st cent. A.D.[1] Most authors agree that the latest plausible date for the arrival of the VII legion in Tilurium was already during the Dalmatian-Pannonian revolt of A.D. 6-9, or immediately after.[2] In the meantime, some finds of Roman military equipment from Tilurium can be dated even to the 4th decade of the 1st cent. B.C.[3] After the departure of *Legio VII Claudia pia fidelis* to Viminacium in the province of Moesia, in the beginning of the second half of the 1st cent. A.D., according to the inscriptions, auxiliary units and perhaps divisions of other legionary units were located in Tilurium. The last auxiliary unit left Tilurium soon after 245 A.D.[4] Following the departure of the last military unit from Tilurium, archaeological evidence of drastic changes regarding the organization of space have been documented. These manifest in the appearance of graves and tombs within the former fortress and, in some cases, in the transformation of individual structures and / or their complete destruction.
Among the aforementioned features, grave “Gardun 2005, SU 20 of 21” from Trench B should be highlighted. It was located in close proximity to the earlier mosaic. The grave was dug into the wall SU 5 and constructed of finely shaped stone fragments used secondarily for the burial of the deceased.[5] The grave did not include any artefacts; therefore it lacks a basis for the precise dating of the architectural remains which were

reused as the bottom and the cover of the grave. According to the result of the anthropological analysis, skeletal remains were of a child who died at the age of 8.5-9.5 years.[6]

During archeological surveillance of the water supply system in the village of Gardun, a grave without a cover (grave “Gardun 1999; VOD 1161; 81”) was documented in the western part of the legionary fortress. The tomb had east-west orientation, with a bottom constructed of tegulae which also formed the bottom of the ancient canal. This canal served as a tomb. At the left of the head of the deceased, a belt buckle was found. It was dated between the 2nd half of the 4th and the 5th cent.[7] Anthropological analysis has shown that it was also a skeleton of a child who died at the age of 3.5-4.5 years.[8]

During the archaeological excavations of the *intervallum*, in the southwestern part of the legionary fortress, right next to the south rampart, grave 1 was documented („Gardun 1999, SU 19“). The grave was built of a cut stone block and can be dated to the Late Antiquity.[9] Anthropological analysis has shown that it was also a skeleton of a child who died at the age of 3.5-4.5 years.[10]

| sample_id | sex (genetic) | date lower (95% CI) | date upper (95% CI) | date inference | burial_id |
|---|---|---|---|---|---|
| R3543 | M | 431 calCE | 600.5 calCE | SITE | Gardun; 1999; SJ 19; G 1 |
| R3544 | M | 549.5 calCE | 600.5 calCE | 14C | Gardun; 1999; VOD 1161; 81 |
| R3545 | F | 431 calCE | 542 calCE | 14C | Gardun; 2005; SJ 20/21 |

**Footnotes**:

[1] Mirjana Sanader, Dino Demicheli, and Marina Milićević Bradač, ‘A “Poet” in the Military Camp at Tilurium’, in *Rimska Vojna Oprema u Pogrebnom Kontekstu*, ed. by Mirjana Sanader and others (Zagreb: Filozofski fakultet Sveučilišta u Zagrebu/ Arheološki muzej u Zagrebu, 2013), pp. 483–91 (p. 489).

[2] Emil Ritterling, ‘Legio. Bestand, Verteilung Und Kriegerische Betätigung Der Legionen Des Stehenden Heeres von Augustus Bis Diocletian’, *Paulys Realencyclopädie Der Classischen Altertumswissenschaft, Band XII, 2* (Stuttgart, 1925), 1329–1829 (col. 1616); J. J. Wilkes, *Dalmatia* (Harvard University Press, 1969), pp. 92–94; Marin Zaninović, ‘Vojni Značaj Tilurija u Antici’, in *Od Helena Do Hrvata* (Zagreb, 1996), pp. 280–90 (p. 284); Stephen Mitchell, ‘Legio VII and the Garrison of Augustan Galatia.’, *The Classical Quarterly*, N.S. 26 (1976), 298-308. (p. 303); Karl Strobel, ‘Zur Geschichte der Legiones V (Macedonica) und VII (Claudia pia felix) in der frühen Kaiserzeit und zur Stellung der Provinz Galatia in der augusteischen Heeresgeschichte’, in *Les légions de Rome sous le Haut-Empire : actes du congrès de*

*Lyon (17 - 19 septembre 1998)*, ed. by Yann Le Bohec (Paris ua, 2000), pp. 515–28 (p. 528).

[3] S. Ivčević u Mirjana Sanader, Domagoj Tončinić, Zrinka Šimić-Kanaet, and others, *Tilurium IV: Arheološka Istraživanja 2007. - 2010. Godine*, ed. by Vinka Matijević, Dissertationes et Monographiae / Arheološki Zavod Filozofskog Fakulteta u Zagrebu, 7 (Zagreb: Filozofski fakultet, Zavod za arheologiju, FF press, 2017), pp. 246–50.

[4] Marin Zaninović, 'Beneficiarii Consularis Na Području Delmata = Beneficiarii Consularis in the Territory of the Delmatae', *Prilozi Instituta Za Arheologiju u Zagrebu*, 24 (2008), 181–84 (pp. 182–83).

[5] Sanader and Tončinić in Mirjana Sanader, Domagoj Tončinić, Zrinka Buljević, and others, *Tilurium III: Istraživanja 2002. - 2006. Godine*, Dissertationes et Monographiae / Arheološki Zavod Filozofskog Fakulteta u Zagrebu, 6 (Zagreb: Filozofski fakultet u Zagrebu, Zavod za arheologiju, 2014), pp. 85–88 Fig. 95–99.

[6] Mario Šlaus and Mario Novak, 'Antropološka Analiza Ljudskog Osteološkog Materijala', in *M. Sanader et Alii, Tilurium III: Istraživanja 2002. - 2006. Godine*, Dissertationes et Monographiae / Arheološki Zavod Filozofskog Fakulteta u Zagrebu, 6 (Zagreb: Filozofski fakultet u Zagrebu, Zavod za arheologiju, 2014), p. 123.

[7] Tomislav Šeparović, 'Metalni Nalazi', in *Tilurium I: Istraživanja = Forschungen: 1997.-2001*, Dissertationes et Monographiae / Arheološki Zavod Filozofskog Fakulteta u Zagrebu, 4 (Zagreb: Filozofski fakultet u Zagrebu, Zavod za arheologiju, 2003), pp. 219–56 (pp. 221, 233) kat. br. 18, T. 2. 7.

[8] Šlaus and Novak, p. 120.

[9] Mirjana Sanader, *Tilurium I: Istraživanja = Forschungen: 1997.-2001*, trans. by Domagoj Tončinić, Disertacije i Monografije 4 (Zagreb: Arheološki zavod Filozofskog fakulteta : Golden marketing, 2003), p. 79.

[10] Šlaus and Novak, p. 121 vidi Grob SJ 235 od 235.

**<u>Literature</u>**:


Mitchell, Stephen, 'Legio VII and the Garrison of Augustan Galatia.', *The Classical Quarterly*, N.S. 26 (1976), 298-308.

Ritterling, Emil, 'Legio. Bestand, Verteilung Und Kriegerische Betätigung Der Legionen Des Stehenden Heeres von Augustus Bis Diocletian', *Paulys Realencyclopädie Der Classischen Altertumswissenschaft, Band XII, 2* (Stuttgart, 1925), 1329–1829

Sanader, Mirjana, *Tilurium I: Istraživanja = Forschungen: 1997.-2001*, trans. by Domagoj Tončinić, Disertacije i Monografije 4 (Zagreb: Arheološki zavod Filozofskog fakulteta : Golden marketing, 2003)

Sanader, Mirjana, Dino Demicheli, and Marina Milićević Bradač, 'A "Poet" in the Military Camp at Tilurium', in *Rimska Vojna Oprema u Pogrebnom Kontekstu*, ed. by Mirjana Sanader, Ante Rendić-Miočević, Domagoj Tončinić, and Ivan Radman-Livaja (Zagreb:

Filozofski fakultet Sveučilišta u Zagrebu/ Arheološki muzej u Zagrebu, 2013), pp. 483–91
Sanader, Mirjana, and Domagoj Tončinić, ‘Das Projekt Tilurium’, in *Rimska Vojna Oprema u Pogrebnom Kontekstu: Radovi XVII. ROMEC-a = Weapons and Military Equipment in a Funerary Context: Proceedings of the XVIIth Roman Military Equipment Conference = Militaria Als Grabbeilage: Akten Der 17. Roman Military Equipment Conference*, ed. by Mirjana Sanader, Ante Rendić-Miočević, Domagoj Tončinić, and Ivan Radman-Livaja (Zagreb, 2013), pp. 411–33
———, ‘Gardun - antički TILURIUM = Gardun – The Ancient TILURIUM’, in *Nalazi rimske vojne opreme u Hrvatskoj - The Finds of the Roman Military Equipment in Croatia*, ed. by Ivan Radman-Livaja (Zagreb: Arheološki muzej u Zagrebu, 2010), pp. 33–53
Sanader, Mirjana, Domagoj Tončinić, Zrinka Buljević, Sanja Ivčević, and Tomislav Šeparović, *Tilurium III: Istraživanja 2002. - 2006. Godine*, Dissertationes et Monographiae / Arheološki Zavod Filozofskog Fakulteta u Zagrebu, 6 (Zagreb: Filozofski fakultet u Zagrebu, Zavod za arheologiju, 2014)
Sanader, Mirjana, Domagoj Tončinić, Zrinka Šimić-Kanaet, Sanja Ivčević, Zrinka Buljević, Tomislav Šeparović, and others, *Tilurium IV: Arheološka Istraživanja 2007. - 2010. Godine*, ed. by Vinka Matijević, Dissertationes et Monographiae / Arheološki Zavod Filozofskog Fakulteta u Zagrebu, 7 (Zagreb: Filozofski fakultet, Zavod za arheologiju, FF press, 2017)
Šeparović, Tomislav, ‘Metalni Nalazi’, in *Tilurium I: Istraživanja = Forschungen: 1997.-2001*, Dissertationes et Monographiae / Arheološki Zavod Filozofskog Fakulteta u Zagrebu, 4 (Zagreb: Filozofski fakultet u Zagrebu, Zavod za arheologiju, 2003), pp. 219–56
Šlaus, Mario, and Mario Novak, ‘Antropološka Analiza Ljudskog Osteološkog Materijala’, in *M. Sanader et Alii, Tilurium III: Istraživanja 2002. - 2006. Godine*, Dissertationes et Monographiae / Arheološki Zavod Filozofskog Fakulteta u Zagrebu, 6 (Zagreb: Filozofski fakultet u Zagrebu, Zavod za arheologiju, 2014)
Strobel, Karl, ‘Zur Geschichte der Legiones V (Macedonica) und VII (Claudia pia felix) in der frühen Kaiserzeit und zur Stellung der Provinz Galatia in der augusteischen Heeresgeschichte’, in *Les légions de Rome sous le Haut-Empire : actes du congrès de Lyon (17 - 19 septembre 1998)*, ed. by Yann Le Bohec (Paris ua, 2000), pp. 515–28
Wilkes, J. J., *Dalmatia* (Harvard University Press, 1969)
Zaninović, Marin, ‘Beneficiarii Consularis Na Području Delmata = Beneficiarii Consularis in the Territory of the Delmatae’, *Prilozi Instituta Za Arheologiju u Zagrebu*, 24 (2008), 181–84
———, ‘Vojni Značaj Tilurija u Antici’, in *Od Helena Do Hrvata* (Zagreb, 1996), pp. 280–90

#### Novo Selo-Bunje, Croatia

**Collaborators**: Mario Novak, Emmanuel Botte, Kristina Jelinčić Vučković, Audrey Bertrand
**Coordinates**: 45.41, 13.66194
**Individuals** (n = 1): R3547

The Novo Selo Bunje site is situated in central Dalmatia, on the island of Brač. A team of experts from Croatia and France has been working on the site since 2015, under the project organized by CNRS (Centre Camille Jullian, Aix-en Provence) and Institute of Archaeology, Zagreb [1]. In the 1st century AD a small Roman villa was constructed there, followed by construction of the large villa rustica a century later. Latest data suggest that life was organized on the site until the 7th century but it is unclear if there were any interruptions. Several construction phases are documented on the site as well. Olive oil and wine production were the main activity on the villa with several production elements from different periods such as the mill, presses, tanks with hydraulic mortar and mosaics.
Three graves were found on the site so far, and the analysed remains were recovered from Grave 1. The grave is dated to the Late Roman period and was located in the western part of the villa. It was cut into the small apse that belonged previously to the thermae of the Pars Urbana of the villa. The archaeological material, except for a late Roman bronze buckle, is not providing enough information for a more precise datation of the grave [2]. The grave itself contained the commingled remains of at least four subadults (6 months, 2 x 1.5-2y, 3-4 y) – the MNI is based on the presence of four right femora. Mild healed new periosteal bone growth (periostitis) was registered on the left tibia of an individual between 1.5 and 2 years of age.

| sample_id | sex (genetic) | date lower (95% CI) | date upper (95% CI) | date inference | burial_id |
|---|---|---|---|---|---|
| R3547 | F | 545.5 calCE | 597 calCE | 14C | NOVO SELO - BUNJE, 2017; Gr 1; SP 5084; SQ 5096; (A) |

**Literature**:
[1] Jelinčić Vučković, Kristina; Botte, Emmanuel, Arheološko istraživanje na lokalitetu Novo Selo Bunje na otoku Braču, 2017. godine,Annales Instituti rchaeologici, 14 (2018), 1; 127-135
[2] Botte, Emmanuel; Bertrand, Audrey; Jelinčić Vučković, Kristina; Leys, Nicolas; Boisson, Antoine, Bunje (Novo Selo, Croatie), campagnes 2017-2018 // Chronique des activités archéologiques de l’École française de Rome, Balkans, 2018, Balkans (2019), 1; 2419, 29 doi:10.4000/cefr.2419

#### Mirine-Fulfinum, Omišalj, Croatia

**Collaborators**: Mario Novak, Morana Čaušević-Bully
**Coordinates**: 45.2039, 14.5437
**Individuals** (n = 3): R2050, R2051, R2053

Archaeological site of Mirine-Fulfinum

The archaeological site of Mirine – *Fulfinum* is a complex archaeological zone situated in the Bay of Sepen (Omišalj County) on the island of Krk. It is composed of several well defined entities:
***Fulfinum*** – an early imperial Roman municipium constructed ex nihilo through the 1st half of the 1st c. AD. The urban center was in use until at least 5th – 7th c. (Čaušević-Bully, Valent, 2015) as the latest date proposed on the basis of the analysis made on ceramic finds (Konestra, 2015). Previous researchers claim that the city was founded exclusively for and by Roman veterans, and so the population would have not originated from the island. On the basis of our recent research, we have a tendency to believe that this was not the case, and that the city had a "local" foundation.
***Mirine*** – is an early Christian basilica built in the first half of the 5th century. The church is preserved up to the roof level; it has, amongst other features, an atrium and a vestibule, where the masonry tombs were discovered.
***Sector 2*** – corresponds to the Late Antique necropolis and mausoleums (3). The typology of the tombs in this sector is very rich and varied from simple grave-pits to the masonry tombs or graves in amphoras for the smallest children. The anthropological analysis made by Mario Novak proves that we have indeed a mixed population, including men, women and children, maybe belonging to the same family? This necropolis seems to be in use for a short period (4th – 5 th c.), after which the church itself, with its surroundings, is used for that particular funerary purpose. The atrium is the last to be abandoned by the funerary function – it was in use at least until the end of the 8th century (knowing that the Adriatic islands can be, by their development, classified as one of those regions where the "very Late Antiquity" is effective, and can be defined chronologically from the 4th to the 8th c.). Other parts of the sites are notable, like for example a late antique villa, situated to the west of the church. But in any case, the whole zone is abandoned definitely around the 9th c.

***Detailed context for the analysed individuals :***

| Sector | Tomb | N° of Individual | C14 datations | sex | age | Type of the funerary architecture |
|---|---|---|---|---|---|---|
| II.1B | T. 2.119 B | IND 7 | 255-300 AD; 315-405 AD 95% | M ? | 50+ | Tomb dug directly in the soil, rudimentary architecture composed of uncut stone blocs disposed against the inner limits of the pit. Two individuals, IND 7 in situ, the other reduced. |
| II.1A | T. 2.120 | IND 9 | | F | 30-38 | Tomb dug directly in the soil, managed directly on the bed-rock. This tomb takes place inside the mausoleum (1st half of the 5th c.). |
| II.2 | T. 2.202 | IND 6 | 390-540 AD 95% | sub-adult | 12-14 | Special type of the grave architecture, grave-pit dug directly in the soil, but arranged with vertical lime slabs on the sides + lime stone slabs in horizontal position for the cover. |

| sample_id | sex (genetic) | date lower (95% CI) | date upper (95% CI) | date inference | burial_id |
|---|---|---|---|---|---|
| R2051 | F | 402.5 calCE | 533.5 calCE | SITE | Omišalj-Mirine 2014, G2.119, IND7, R. petrous 30/11/2015 |
| R2053 | F | 402.5 calCE | 533.5 calCE | 14C | Omišalj-Mirine 2014, G2.120, IND9, petrous, 2/12/2015 |
| R2050 | F | 402.5 calCE | 533.5 calCE | SITE | Omišalj-Mirine 2014, G2.202, IND6, petrous, 4/12/2015 |

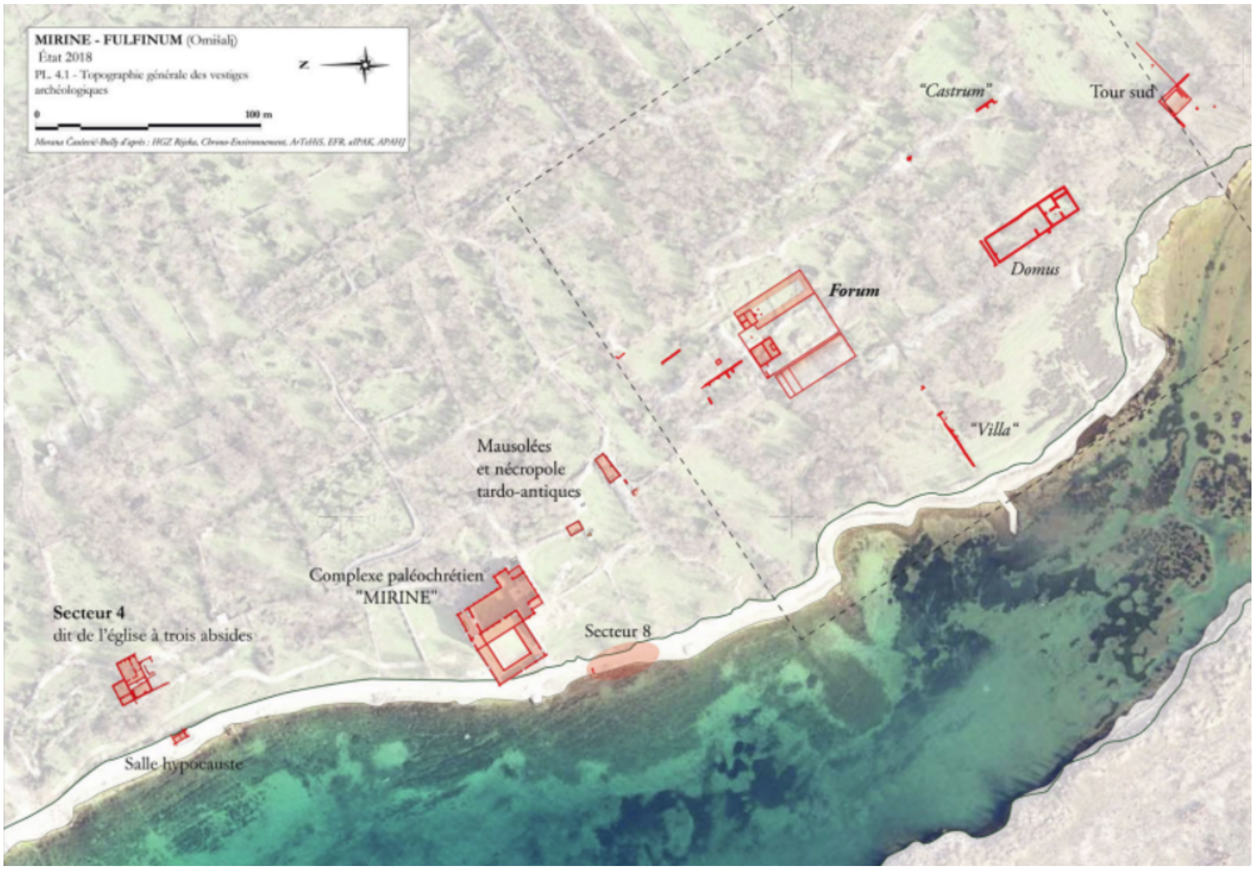

General topography of the site Mirine-Fulfinum (2019)

**Literature**:
KONESTRA, Ana, Keramika s Foruma Municipia Flavia Fulfinuma (otok Krk, Hrvatska) istraživanja od 2007. do 2013. godine (Pottery from the Forum of Municipium Flavium Fulfinum (Krk Island, Croatia) – research between 2007 and 2013), Prilozi Instituta za arheologiju, Zagreb, 32, 2015, p. 147-214.
ČAUŠEVIĆ-BULLY, Morana, VALENT, Ivan, "Municipium Flavium Fulfinum. Dijakronijska studija gradske strukture s posebnim osvrtom na forumski prostor (Municipium Flavium Fulfinum. Diachronic study of the city structure with a special attention to the forum)", Prilozi Instituta za arheologiju, Zagreb, 32, 2015, p. 111-146.
ČAUŠEVIĆ-BULLY, Morana, "La ville de Fulfinum (île de Krk). Nouveau regard sur le sort d'une ville antique entre l'Antiquité tardive et le haut Moyen âge", Hortus Artium Medievalium, 20, Zagreb-Motovun, 2014, p. 157-169.

#### Osijek, Croatia

**Collaborators**: Mario Novak, Tino Leleković
**Coordinates**: 45.550532, 18.678046
**Individuals** (n = 3): R3655, R3656, R3657

The analysed burials were excavated in the eastern cemetery of the Roman period city Mursa (city of Osijek in eastern Croatia). Mursa was a veteran colony (*colonia Aelia Mursa*) founded by Emperor Hadrian in 133 CE as the civilian and administrative center of this part of the border (limes) zone; Mursa is one of largest settlement in the eastern part of *Pannonia* (Roman province *Pannonia Inferior*). The graves were excavated during the preventive excavation in 2008 conducted at the Ban Jelačić Square and led by Tino Leleković from the Croatian Academy of Sciences and Arts in Zagreb. During this occasion, a space outside of the city walls some 100m from the town gate was excavated during which five different development stages were recorded. This excavation, covering an area of about 3000 $m^2$, revealed 101 cinerary graves from late 1st and early 2nd century, preceding the foundation of the colony, and 311 graves that were apparently buried within a very short time period, in time of crisis between 260 and 270 CE. The appearance of family graves suggests that the cemetery was used for people who died of an epidemic or war.

Burials 43, 167 and 196, Ban Jelačić Square
Inhumation burials 43, 167 and 196 belong to the eastern cemetery of the colony; they are part of the cemetery dated narrowly to the 260's and 270's CE.

**Grave 43**
The skeleton from burial 43 was found buried in the brick tomb 159 cm in length – based on the tomb length it may be presumed that the grave belonged to a child or a gracile woman. The bones were found scattered without any artefacts suggesting the tomb was looted.

**Grave 167**
The skeleton was placed directly in the soil, slightly on its left side with the legs in flexed position, the left arm on the chest and the right arm under the head. No artefacts were found in the grave. According to the burial rite that was common for this part of the province of Pannonia, the burial can be dated between the 2nd and the 5th century CE.

**Grave 196**
The deceased was placed directly in the soil, on its back in an extended position with the left arm on the chest and the right arm under the head. No artefacts were found in

the grave. According to the burial rite that was common for this part of the province of Pannonia the burial can be dated between the 2nd and the 5th century CE.

| sample_id | sex (genetic) | date lower (95% CI) | date upper (95% CI) | date inference | burial_id |
|---|---|---|---|---|---|
| R3655 | M | 133.5 calCE | 306 calCE | 14C | Osijek - Trg B. Josipa Jelačića 2008, G 43 |
| R3656 | F | 229.5 calCE | 329.5 calCE | 14C | Osijek - Trg B. Josipa Jelačića 2008, G 167 |
| R3657 | F | 229.5 calCE | 329.5 calCE | 14C | Osijek - Trg B. Josipa Jelačića 2008, G 196 |

### Ščitarjevo, Croatia

**Collaborators**: Mario Novak, Dora Kušan Špalj
**Coordinates**: 45.767834, 16.123606
**Individuals** (n = 2): R3659, R3660

Ščitarjevo (Roman period *Andautonia*) is a small settlement in continental Croatia, some 10 km south of Zagreb. The settlement was located in the Roman province of Pannonia, on the road connecting Poetovia (modern Ptuj) and Siscia (Sisak). The settlement was municipium and one of the major centers in this part of the province of Pannonia. The settlement existed between the 1st and the 4th century, after which it is believed to have been destroyed during the Great Migration. Systematic archaeological excavations conducted continuously since 1981 uncovered the remains of a thermal complex with water pipes and sewers, the remains of numerous buildings and roads, and possibly an amphitheatre. Today, this is a large open-air archaeological park open to the public.
In 2007 a rescue excavation led by Dorica Nemeth-Erlich and Dora Kušalj Špan from the Archaeological Museum in Zagreb was conducted due to the construction of a new sports hall and upgrade of the local elementary school. Beside numerous architectural remains (buildings, roads) and small finds, especially important is a find of a single inhumation burial in the SE part of the schoolyard near the remains of Roman buildings.
In January 2009, the company Geoarheo (Goran Skelac) carried out a small-scale rescue excavation at the site Ščitarjevo – School. During the excavation a late Roman period double inhumation burial was uncovered.

**Ščitarjevo, 2007, grave 1**
The burial in plain soil contained the skeleton of a younger male. The skeleton is about 185 cm long, complete and nicely preserved, with orientation SW - NE. Roman bricks were put around the head of the deceased. The burial contained an iron spear tip, a shield boss, two silvered and gilded belt-buckles in the pelvic region, a long sword (spatha) under the left arm and knife scabbard chape (Ortband) under the left scapula. Based on the recovered artefacts the burial can be dated to the beginning of the 6th century CE (Langobard?).

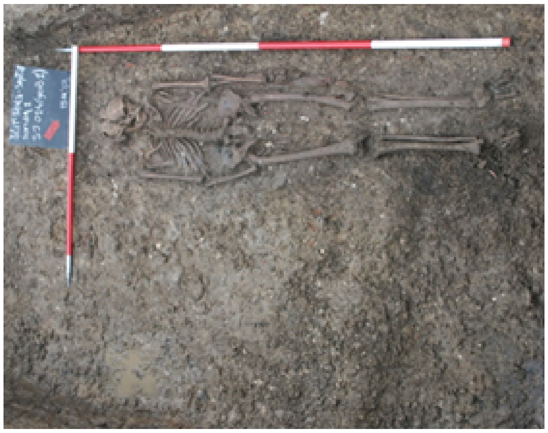

**Ščitarjevo, 2009, SU 3**
Double inhumation burial in plain soil, no grave architecture; the skeletons were oriented west (head) – east (legs). Southern skeleton (younger male), much better preserved, was laid on its back with the arms extended along the body, while a very badly preserved skeleton (probably an adult woman) was buried on its side in a flexed position. The only find in the burial was a silvered coin of Constantine II. The burial can be dated to the middle of the 4$^{th}$ century CE.

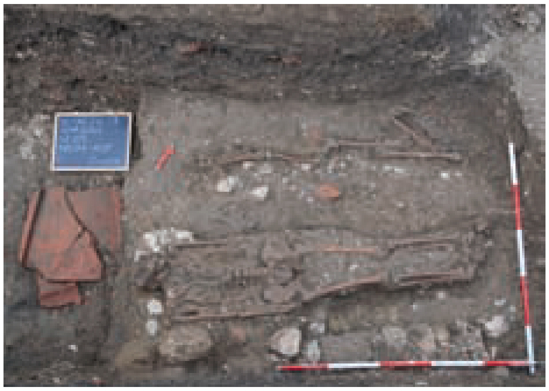

| sample_id | sex (genetic) | date lower (95% CI) | date upper (95% CI) | date inference | burial_id |
|---|---|---|---|---|---|
| R3660 | M | 433 calCE | 560.5 calCE | 14C | Ščitarjevo, 2007, G 1 |
| R3659 | M | 248.5 calCE | 378 calCE | 14C | Ščitarjevo, 2009, SJ 03 |


**Literature**:
Kušan Špalj D., Nemeth-Erlich D. 2008. Šćitarjevo – dvorište osnovne škole. Hrvatski arheološki godišnjak, 4/2007, 198-201.
Nodilo H. 2010. Šćitarjevo – škola. Hrvatski arheološki godišnjak, 6/2009, 254-255.

#### Sipar - Umag, Croatia

**Collaborators**: Mario Novak, Branka Milošević Zakić
**Coordinates**: 45.467293, 13.506565
**Individuals** (n = 4): R2045, R3662, R3663, R3664

Sipar is located in the north-western part of Croatian Istria, about 4 kilometers north of Umag, on a small peninsula (about 0.5 ha) that connects the 80 meter long slope with the mainland. In the written documents, Sipar is mentioned, along with other Istrian coastal towns (*civitates*), as *Sapparis* and *Sipparis* in the Cosmography of the Anonymous from Ravenna. The archaeological excavations that started in 2013 led by Branka Milošević Zakić from the Museum of Croatian Archaeological Monuments revealed several developmental cultural stages of the site. Based on the archaeological finds, the existence of Sipar can be dated between the 1$^{st}$ century BCE and the end of the 9$^{th}$ century CE. The earliest material finds show that Sipar was inhabited at least from the middle of the 1$^{st}$ century BCE, possibly as a temporary military camp for the Roman army located on the trade route leading from northern Italy to the eastern Adriatic coast. The fortification in the eastern part and the settlement in the western part of the peninsula are dated to the 6th century and were destroyed probably at the end of the 9$^{th}$ century. By the end of 2018, a total of 11 children's graves were found in the area of the settlement: late antique graves were buried below the walkway of the first phase of the settlement, while early medieval graves were buried in the collapsing layer. Archaeological excavations of Sipar are still ongoing.

**Sipar 2016, grave 1**
Room: 4/5
Grave construction: amphora LR4 (Gaza type) (SU 36)
Skeleton: SU 36A, subadult, under 1 year
Discovered: 12/10/2016
Child's grave in amphora was found in room 4/5, oriented east-west. The amphora was buried in the leveling stratum SU 41 (early Roman material: tegulae, amphorae, fine and coarse pottery, amphorae lids, spinning whorls, fibulae, nails). The amphora is a late Roman product (so called Gaza type amphora produced in Palestine) that was often used to bury children. No other artefacts were found in the grave. The burial can be dated to the 6th century CE (probably the second half).

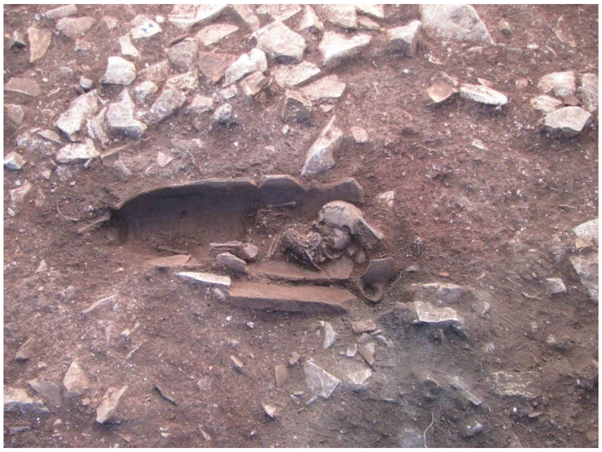

**Sipar 2017, grave 4**
Room: 15
Grave construction: stone grave construction (SU 110), only a few stone slabs are preserved
Skeleton: incomplete (SU 110A), subadult, under 1 year
Discovered: 3/10/2017
Child’s grave 4 was found after the removal of the burned structure SU 105; it was dug in the red soil SU 106. The stone construction was preserved along its southern side, and the grave is of east-west orientation. No artefacts were found in the grave. The burial can be dated to the second half of the 6th century CE.

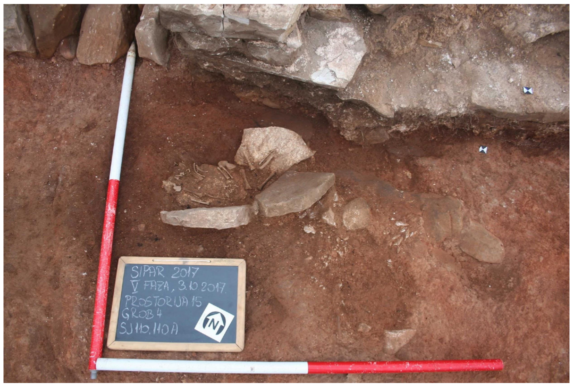


**Sipar 2018, grave 5**
Room: 17
Grave construction: no grave construction
Skeleton: SU 127, subadult, under 1 year
Found: 18/9/2018
Subadult bones were found on the structure of the grave 6 (SU 128); everything was found within the layer SU 114 (fine dusty soil, at the same stratum one so called “Buzet type earring” was also found), and outside of the bricked structure SU 113 along the

western wall PR 17. The grave is partially destroyed, the head of the deceased was located on the northern side. No artefacts were found in the grave. The burial can be dated to the 7th/8th century CE.

**Sipar 2018, grave 6**
Room: 17
Grave construction: stone grave construction (SU 128)
Skeleton: SU 128, subadult, under 1 year
Discovered: 18/9/2018
Grave 6 was discovered after the removal of the skeleton from grave 5. Grave construction is nicely preserved, it was found within the layer of fine dusty soil SU 114 (see grave 5). No artefacts were found in the grave. The burial can be dated to the 7th/8th century CE.

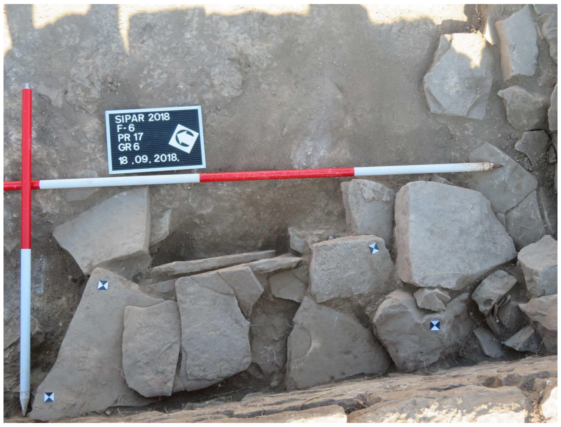


| sample_id | sex (genetic) | date lower (95% CI) | date upper (95% CI) | date inference | burial_id |
|---|---|---|---|---|---|
| R2045 | F | 550.5 calCE | 603.5 calCE | 14C | Umag-Sipar 2016, G1 petrous, 21/3/2017 |
| R3662 | F | 558.5 calCE | 639 calCE | 14C | Sipar; 2017; G 4; L temporal; Roman period |
| R3663 | F | 686.5 calCE | 876 calCE | 14C | SIPAR 2018; GR 5; LEFT TEMPORAL |
| R3664 | M | 679.5 calCE | 823 calCE | 14C | SIPAR 2018; GR 6; LEFT TEMPORAL |

**Literature**:

B. Milošević Zakić, Sipar,arheološka istraživanja i zaštita lokaliteta od 2013. do 2015. Godine, SHP Vol. III, 46, Split 2019, 205-211.

### Sisak-Pogorelec, Croatia

**Collaborators**: Mario Novak, Ivan Drnić, Željka Bedić
**Coordinates**: 45.464666, 16.377654
**Individuals** (n = 3): R2040, R2041, R2042

**Sisak – Pogorelec (western necropolis)**
Sisak is a modern city in continental Croatia, located some 50 km SE of Zagreb. During the 4th century BCE it was inhabited by both the indigenous people Breuci and members of the Celtic tribe Segestanoi (hence the name of the settlement Segestica). The Romans conquered Segestica in 35 CE and founded a new settlement on the opposite bank of the river Kupa (Siscia). It became a Roman colony during the reign of the emperor Vespasianus (Colonia Flavia Siscia). It was one of the largest settlements in the Roman province of Dalmatia with a river harbour, fortifications, theatre and amphitheatre, termae, aqueducts, several cemeteries and mint. During Diocletian's rule it became the capital of the province Pannonia Savia.
The western necropolis of the city was located in the north-eastern corner of the Pogorelec peninsula, on the right bank of the river Kupa. Continuous archaeological excavations at Pogorelec started in 2012 and are led by Ivan Drnić from the Archaeological Museum in Zagreb. So far, a total of 23 skeletal burials have been discovered at this site. The whole western necropolis of Siscia can be dated to the 3rd/4th century CE.

**Skeletal burial 11**
Burial of an adult male. The skeleton was laid on its back with arms extended along the body, oriented east-west with the head on east. One iron object (tweezers?) was found by the left arm.

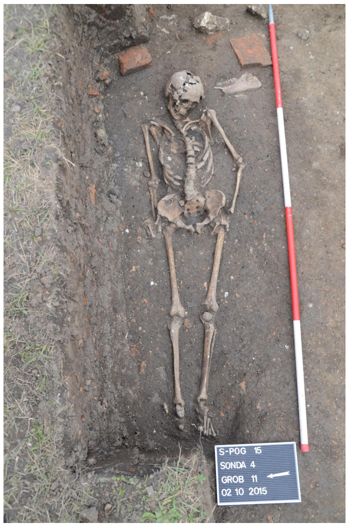

**Skeletal burial 12**

Burial in a wooden coffin (iron nails). The skeleton of a younger male was found laid on its back with bent arms put on the chest, oriented east-west with the head on west. Ceramic cup with folds (Faltenbecher) was found between the lower legs.

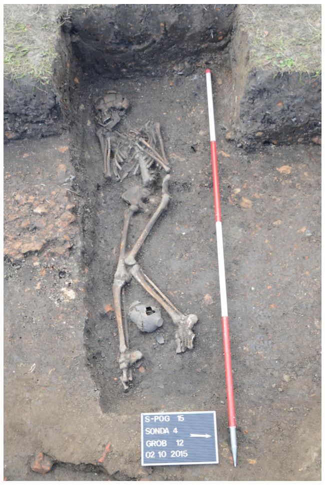


**Skeletal burial 13**

Burial in plain soil. The skeleton of a middle-aged woman was laid on its back with arms extended along the body, oriented NE-SW with the head on NE. One bronze and one bone bracelets were found on the left arm, along with one bronze earring, one bone and four glass beads.

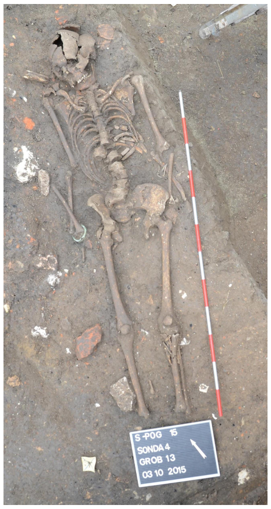

| sample_id | sex (genetic) | date lower (95% CI) | date upper (95% CI) | date inference | burial_id |
|---|---|---|---|---|---|
| R2040 | M | 245 calCE | 402 calCE | SITE | Sisak-Pogorelec 2015, Grave 11, male, 40-50 |
| R2041 | M | 245 calCE | 361 calCE | 14C | Sisak-Pogorelec 2015, Grave 12, male, 25-30 |
| R2042 | F | 252 calCE | 402 calCE | 14C | Sisak-Pogorelec 2015, Grave 13, female, 35-45 |

**Literature**:

Baćani I., Škrgulja R., Tomaš-Barišić T. 2018. Nekropole Siscije. Sisak: Gradski muzej.

#### Trogir-Dragulin, Croatia

**Collaborators**: Mario Novak, Anna Osterholtz, Lujana Paraman, Julianne Paige
**Coordinates**: 43.524984, 16.252825
**Individuals** (n = 1): R3665

Excavation type: rescue excavation
Excavation director: Lujana Paraman, Trogir Town Museum
Excavation time: 16. 5. - 7. 6. 2016.
Anthropological analysis: Anna Osterholtz, Julianne Paige, Mississippi State University, USA

The site Dragulin is located in the western part of Trogir's Malo polje plain, about 400 m west of the main cemetery of Roman Tragurium in the Dobrić area. Several cremation burials found in the 1960s and 1990s pointed to the existence of an Early Roman cemetery in this area, next to the modern Put Dragulina street which follows the route of one of the main Roman roads leading from Tragurium towards the hinterland. It is still unclear whether this is a public cemetery or a cemetery connected to one of the unidentified villa rusticas in the western part of the Malo polje plain.
In 2016 a small segment of the cemetery was excavated after the construction work on a private parking lot revealed several stone urns. The earthwork destroyed the upper layers of the cemetery, leaving untouched only the graves buried in the soil between the bedrock. During the rescue excavation led by the Trogir Town Museum, 42 graves were documented, dating from the beginning of the 1st c. AD to the 4th c. AD, but most of them dated to the 1st and 2nd c. AD. The anthropological analysis on recorded remains of 37 individuals was carried out by Anna Osterholtz and Julianne Paige from the Mississippi State University during the 2017 and 2018.
About 60% of graves were cremated individuals buried in different containers – various ceramic pots (71.43 %) and stone urns (28.57 %). The individuals were mostly adults (43.24 % of the analysed sample, adolescents included), few of them distinguished as young females (3) and a male (1). The smaller number of cremated burials belonged to young children (10.81 %) aged between 1,5 and 5 years and perinates (5.40 %). Grave goods consisted mostly of ceramic, glass and one bronze unguentaria, followed by ceramic lamps and ceramic vessels (cups, bowls, a juglet). Jewellery, costume pieces and cosmetic accessories were also represented, as well as gaming pieces (astragali) found in two child's burials. Along with grave goods, the remains of special rituals such as covering the urn with large stone slabs and funeral pyre remains were also recorded. The remaining 40% of graves were inhumations belonging to the perinates and infants (40.54 % of the analysed sample), who were buried in a small pit (50%) or in an amphora coffin (43.75%), with the exception of one burial in a miniature stone urn (Grave 5). The individuals were mostly E-W oriented, although the W-E, N-S and S-N

orientation were also recorded among pit burials. Grave goods were present in four pit burials and one amphora burial and are represented mostly by symbolic objects such as gaming pieces (animal phalanx bone, Grave 13), led pendant? (Grave 17), glass bottle (Grave 31) or chert fragments (Grave 24). The grave goods and burial rite of the infant burial Grave 5 are consistent with the adult cremation burials.
The findings and the stratigraphy point out to dating of the cremated burials and most of the perinate and infant burials from the beginning of the 1st to the end of 2nd c. AD, with two amphora burials belonging to the period of 3rd and 4th c. AD (Grave 35 and 41).

**Grave 17** - R3665

- Infant pit burial (b - 2 months), found above adolescent cremation burial Grave 27 (1st half of 1st c. AD). Good preservation of skeleton.
- Grave goods: folded lead sheet metal (pendant?)
- Description: The infant was set in a shallow oval burial pit, NW-SE oriented (with the head on the NW side). It was laid slightly on the left side, facing east. The folded lead sheet metal was placed between the head and the left shoulder.

| sample_id | sex (genetic) | date lower (95% CI) | date upper (95% CI) | date inference | burial_id |
|---|---|---|---|---|---|
| R3665 | M | 124 calCE | 217 calCE | 14C | TROGIR; DRAGULIN; 2016; G 17; pars petrose |

#### Trogir-Policija, Croatia

**Collaborators**: Mario Novak, Lujana Paraman, Julianne Paige, Maja Bilić
**Coordinates**: 43.51859, 16.13651
**Individuals** (n = 1): R3670

The site of Dobrić is considered to be a central cemetery of Roman Tragurium, modern-day Trogir (central Dalmatia). Located on the mainland just opposite of the historic town, the area is marked by an intersection of ancient roads and pathways leading from the northern city gate to its ager, and further towards the hinterland and the Kaštela bay.
The available data suggest that the cemetery was in use from the Early Roman to the end of the Late Antique Period with documented cremation urn burials of 1st and 2nd c. AD and different types of inhumation burials (pit, amphora, tile and cist graves, sarcophagi, vaulted tombs,) of 1st to 6th c. AD.
In 2011, during the construction work on the sewage and water supply system Kaštela - Trogir, a small segment of the cemetery was excavated at the beginning of Put Dragulina Street, just south of the modern police station (PP Trogir). During the rescue excavation led by the firm Palisada Ltd., 28 graves were documented in several layers, dating from the end of 2nd or 3rd c. probably to the 5th or 6th c. AD. Buried individuals were mostly perinates and infants, buried in an amphora coffin. Grave goods and other findings were recorded only in three graves:  jewelry and costume pieces in the adult tile burial Grave 5, pottery and iron nails in the infant amphora burial Grave 22 and adult pit burial Grave 17.

**Grave 18** - R3670
Perinate pit burial (38 - 40 weeks). Skeleton partly preserved, fragmented.
Grave goods: /
Description: The perinate was laid on a right side (facing east) in a plain burial pit, S-N oriented (with the head on the S side). A bronze chain was placed on the left side of its chest.

| sample_id | sex (genetic) | date lower (95% CI) | date upper (95% CI) | date inference | burial_id |
|---|---|---|---|---|---|
| R3670 | F | 120.5 calCE | 215 calCE | 14C | TROGIR; POLICIJA; 2011; G 18; pars petrose |

#### Velić, Croatia

**Collaborators**: Mario Novak, Mirjana Sanader, Domagoj Tončinić, Mario Šlaus, Vinka Matijević, Domagoj Bužanić
**Coordinates**: 43.616374, 16.801661
**Individuals** (n = 1): R3685

Velić is a small settlement near the town of Trilj in southern Croatia. The late Antique site has been known since 1921 due to a find of a Roman milestone dated to the 3rd century CE. In 2011, experts from various institutions conducted a field survey of the area. Based on the results of this survey, in 2013, archaeologists from the Department of Archaeology, Faculty of Humanities and Arts Archaeological of the University of Zagreb (Domagoj Tončinić, Mirjana Sanader) conducted archaeological excavations of the remains of the rectangular building with a semicircular apse. At that point, the whole structure was dated to the period of late antiquity and interpreted as an early Christian memoria. The *memoria* consists of three rooms with a vaulted tomb with an entrance of the *a pozzetto* type, which was originally enclosed by a stone slab.

**Tomb 1, 2013** - R3685
The tomb has a rectangular ground plan of east-west orientation. Along the inner side walls of the tomb two stone-built and plastered beds divided by a canal were documented. On the bed along the south wall a skeleton with a skull oriented to the east was documented. Anthropological analysis revealed that it was a skeleton of a man who died at the age of 50-60 years. Human bones, along with several fragments of animal bones, were also documented in the canal between beds, and at the entrance to the tomb. This, among other things, confirms that the tomb was devastated prior to the excavation. Very poorly preserved coin, dated in the mid-4th century, was found in the north room, in front of the tomb entrance.

| sample_id | sex (genetic) | date lower (95% CI) | date upper (95% CI) | date inference | burial_id |
|---|---|---|---|---|---|
| R3685 | M | 434.5 calCE | 565 calCE | 14C | Velić; 2013; GR 1 |

**Literature**:
Bubić V. 2015. Terenski pregled lokaliteta kod Velića. Izdanja Hrvatskog arheološkog društva, 29: 91-101.

#### Zadar, Croatia

**Collaborators**: Mario Novak, Timka Alihodžić
**Coordinates**: 44.114352, 15.227943
**Individuals** (n=6): R3745, R3746, R3747, R3742, R3743, R3744

Zadar is a city situated on the eastern Adriatic coast in modern-day Croatia. Iulius Caesar founded Zadar as a Roman colony in 48 BC (Colonia Iulia Iader) although the city was inhabited at least from the early Iron Age by the local population called Liburni. Iader was part of the Roman province Dalmatia and it was primarily settled by veterans from Italy. The city was situated on the peninsula and organized on principles of Roman urbanism with major streets intersected at right angles surrounded by massive stone walls. Zadar was primarily an agricultural colony with a large seaport, thermae, a public square (forum) and an elevated capitolium with a temple. A 40 km long aqueduct brought drinking water to the city. In 481 Dalmatia became part of the Ostrogothic kingdom, which is the period of decline for the city when many civic buildings were ruined due to the advanced age and negligence. At the beginning of the 6$^{th}$ century it was hit by an earthquake, which destroyed entire complexes in the city.
Due to urban construction, rescue excavations of the Roman necropolis in Relja city district were carried out between 2005 and 2014 by the Archaeological Museum Zadar (Timka Alihodžić). The necropolis was located about 0.5 km meters from the city walls along the road leading to the southeast (province capital of Salona). Together with previous excavations the total number of recovered burials is around 2500 (inhumations and cremations) thus making this site one of the largest Roman period cemeteries on the Adriatic. Inhumation burials from this site are dated between the 1$^{st}$ and 5$^{th}$/6$^{th}$ century CE.

**Ulica Polačišće, 2008, grave 6**
Rite: inhumation, adult individual
Grave description: the deceased was buried in the plain soil; the orientation is east (head) - west (legs) with amorphous stones laid around the skeleton. From the waist down the skeleton was destroyed by the recent construction of water pipes. The arms are bent, the left was placed on the pelvic girdle while the right was put on the chest.
Associated artefacts: none.
Chronology: $1^{st}$-$4^{th}$ century CE.

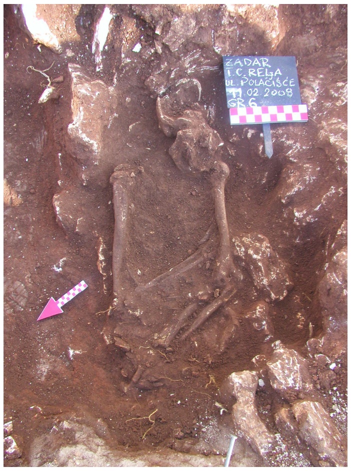


**Ulica Polačišće, 2008, grave 35**
Rite: inhumation, adult individual
Grave description: the deceased was buried in the plain soil along the eastern side of the grave; the orientation is NE (legs) - SW (head). The skeleton was laid on its back with the arms on the pelvic girdle. Amorphous stones were laid above the deceased on the left side.
Associated artefacts: oyster shell.
Chronology: $1^{st}$-$4^{th}$ century CE.

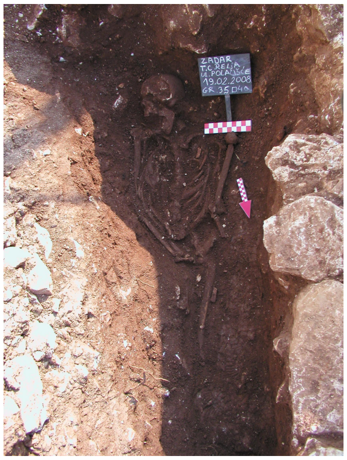

**Hypo banka, 2006, grave 56**
Rite: inhumation, subadult
Grave description: the deceased was laid on its back with the orientation east (head) - west (legs), along the wall of the grave plot. Amorphous stones were laid above and around the deceased. The skeleton is not complete.
Associated artefacts: ceramic bowl, four glass beads, glass balsamarium, bronze chain with a bottle (aribalos).
Chronology: beginning of the $3^{rd}$ century CE.

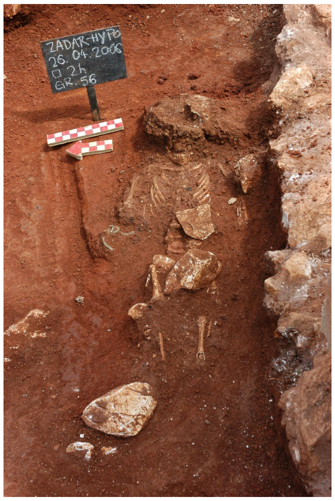


**Ulica Katarine Zrinski / Poliklinika, 2013, grave 8**
Rite: inhumation, adult individual
Grave description: the deceased was laid on its back in the plain soil parallel to the remains of the wall (probably the wall of the grave plot). Amorphous stones were laid above the deceased. This burial destroyed the previous cremation burial dated to the $1^{st}/2^{nd}$ century CE (the remains of ceramic and glass vessels, lucernae, burnt bones and charcoal).
Associated artefacts: none.
Chronology: $2^{nd}$-$4^{th}$ century CE.

**Poliklinika, 2014, grave 38**
Rite: inhumation, adult individual
Grave description: the deceased was laid on its back on bedrock with the orientation west (head) - east (legs). Amorphous stones were laid above and around the deceased. The skeleton is badly preserved and incomplete.
Associated artefacts: ceramic glass and jug, bronze coin (unreadable).
Chronology: $2^{nd}$-$4^{th}$ century CE.

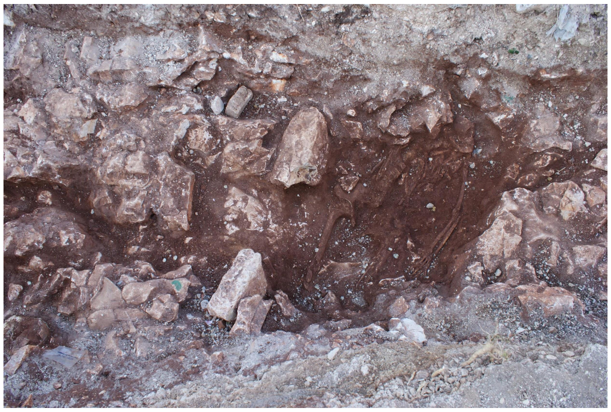

**TC Relja, 2005, grave 173**
Rite: inhumation, subadult
Grave description: incomplete subadult skeleton positioned between bedrock on eastern and western sides, and smaller stones on southern and northern sides. The grave was probably covered by tegulae. The deceased was laid on its back with the orientation east (head) - west (legs).
Associated artefacts: two bronze coins, three silver styli.
Chronology: 1st/2nd century CE.

| sample_id | sex (genetic) | date lower (95% CI) | date upper (95% CI) | date inference | locality | burial_id |
|---|---|---|---|---|---|---|
| R3743 | M | 81.5 calCE | 210 calCE | 14C | Zadar Ulica | ZADAR; Ul. Polačišće; 2008; GR 6; temp |
| R3744 | M | 81.5 calCE | 210 calCE | SITE | Zadar Ulica | ZADAR; Ul. Polačišće; 2008; GR 35 |
| R3745 | M | 22.5 calCE | 121 calCE | 14C | Zadar Hypo_banka | ZADAR; HYPO BANKA; 2006; GR 56; R. temp |
| R3747 | M | 127 calCE | 227.5 calCE | SITE | Zadar Poliklinika | ZADAR; POLIKNIKA; 2014; GR 8; R temp |
| R3746 | F | 127 calCE | 227.5 calCE | 14C | Zadar Poliklinika | ZADAR; POLIKNIKA; 2014; GR 38; R temp |
| R3742 | M | 127 calCE | 227.5 calCE | 14C | Zadar Relja | ZADAR; TC RELJA; 2005; GR 17; temporal |

### Metz, France

**Collaborators**: K. Kazek, J. Trapp, Julien Trapp, Olivia Cheronet
**Coordinates**: 49.119677, 6.174949
**Individuals** (n = 5): R2055, R2057, R2058, R2065, R2066

Metz, Sablon (K. Kazek, J. Trapp)

The Sablon district, which is located in the southern part of the city of Metz, was, during the Gallo-Roman period, a huge necropolis where both inhumations and cremations are found. Towards the end of the 19th century, the exploitation of the sandpits enabled the uncovering of sarcophagi (stone), cists (brick and tile), coffins (wood) and vats (lead). These characterise the new burial practices developed during late Antiquity.
The largest funerary space spans almost a kilometre, on either side of the *via Scarponensis* (portion of the Reims/Metz road). Another site has been located further to the West, and must be linked to the Chemin de la Moselle (presently Avenue de Nancy). Other burials have been found close to the Voie de la Meurthe (presently Avenue André-Malraux) and much further to the South.
The first important finds occurred and were progressively exhumed from the start of the development of the train station around 1840-1850. The building works undertaken by the army at the Citadelle and in the ditches of the Lunette d'Arçon (end of 19th century) have significantly enriched the body of finds. With the urban expansion of the city during the German period, ballast and sand pits appeared over the whole territory of the Sablon and quickly became a golden opportunity for archaeological research.
The workers made numerous and often spectacular finds: the discovery of the sévir *C. Livius Castor* in the Distler sand pit in 1904, an onyx cinerary urn from Egypt found in a stone box close to the Place du Roi Georges in 1908, or an ivory ring case in the shape of a temple placed in a burial in the area of the Lunette d'Arçon in 1848.
The finds that particularly struck their discoverers at that time were the sarcophagi and coffins. Notably, a small lead coffin of a newborn was found in the zone demarcated by the rue Monseigneur-Heintz and the Jean-XXIII school.
Close to the Mey sandpit two limestone Victory statues were other noteworthy finds. They could belong to important mausoleums.
It is also to be noted that the human remains found in the Distler sandpit were also buried in stone coffins without many grave goods. This suggests that these were the graves of a relatively wealthy group of people, with the means to commission graves of high quality for themselves.
Nevertheless, for this huge burial zone which was used from the 1st century BC to the early Middle Ages ("Haut Moyen Age"), it remains difficult to identify distinct groupings making it challenging to identify a social hierarchy of the graves, although the presence

of burials of a familial type (cremation burials 350 m north of the Horgne farm) has made archaeologists reconsider some of their approaches to this problem.
The Sablon area can be compared to the Collatina necropolis close to Rome by its chaotic organisation, although at a different scale.

| sample_id | sex (genetic) | date lower (95% CI) | date upper (95% CI) | date inference | burial_id |
|---|---|---|---|---|---|
| R2055 | M | 432.5 calCE | 551 calCE | 14C | Metz: Sablon, Gallo Roman, Inv.No.:2014.0.2278, Box: 89, Upper molar, 23/8/18 |
| R2057 | F | 404 calCE | 535 calCE | 14C | Metz: Sablon, Gallo Roman, Inv.No.: 2014.0.2266 young ind., Box 89, L.petrous (CBD) + cranial fragment, 23/8/18 |
| R2058 | F | 404 calCE | 535 calCE | SITE | Metz: Sablon, Gallo Roman, Inv.No.: 2014.0.2266 older ind., Box:89, L.petrous CBD, 23/8/18 |
| R2065 | F | 1297.5 calCE | 1395 calCE | SITE | Metz: Saint-Pierre-aux-Nannains, Gallo Roman, Ins.No.:2016.0.1224 ind. 3, Box:550, L.petrous CBD + cranial fragment, 24/8/18 |
| R2066 | F | 1297.5 calCE | 1395 calCE | 14C | Metz: Saint-Pierre-aux-Nannains, Gallo Roman, Inv.No.:2016.0.1224 ind. 4, Box:550, L.petrous CBD + cranial fragment, 24/8/18 |

**Literature**:

Bulmé, A. 2001. Les sarcophages gallo-romains en plomb du Musée de Metz. Mémoire de maîtrise sous la direction d'A.-M. Adam, université March-Bloch de Strasbourg, 235 p.

Grapin, C. 1996. Metz : le quartier du Sablon. Dossier scientifique spécifique dactylographié, stage patrimonial n° 1, 53 p.

Grapin, C. 2005. « Topographie funéraire du quartier du Sablon ». In (P. Flotté, ed) Metz. 57/2. Académie des Inscriptions et Belles-Lettres, Paris, 315-327.

Keune, J.-B. 1900a. Bericht über die Erwerbungen des städtischen Museums. Geschäftsjahr 1900 nebst ein Überblicke über die Entwickelung der Sammlungen. In Jahrbuch der Gesellschaft für lothringische Geschichte und Altertumskunde, 346-416.

Keune, J.-B. 1903a. Sablon in römischer Zeit. In Jahrbuch der Gesellschaft für lothringische Geschichte und Altertumskunde, 324-460.

Keune, J.-B. 1904a. Altertumsfunde aus der Flur Sablon oder südlichen Vorgelände von Metz (1903-1905). In Jahrbuch der Gesellschaft für lothringische Geschichte und

Altertumskunde, 316-384.
Keune, J.-B. 1910. Altertumsfunde in Lothringen. Erwerbungen des Museums der Stadt Metz, von 1905 bis 1910. In Jahrbuch der Gesellschaft für lothringische Geschichte und Altertumskunde, p. 487-537.
Toussaint, M. 1948. Metz à l’époque gallo-romaine. Metz, Impr. Paul Even, (coll. Annuaire de la Société d’histoire et d’archéologie de la Lorraine, 222 p.

#### Sarrebourg, France

**Collaborators**: Christèle Baillif-Ducros
**Coordinates**: 48.7327, 7.0526
**Individuals** (n = 12): R11550, R11552, R11553, R11554, R11555, R11556, R11557, R11558, R11559, R11560, R11561, R11563

The Late Antiquity necropolis of the antique city of Sarrebourg, *Pons Saravi,* is situated on the western flank of the “Marxberg” hill. The burial ground, dated to the 4th century, is at the intersection and on the edge of two Roman roads leading to, on the eastern side, the secondary Mediomatrici urban centre. 88 burials, of which 71 are individual, have been documented during the excavation of the site. The buried population (a total of 93 individuals) is split between 72 adults (of which 23 males, 24 females, and 25 of indeterminate sex) and 21 immature individuals. To characterise the parental relationships between these individuals (parents, children and lineages), 72 individuals (of which 51 adults and 21 immature individuals) have been sampled for ancient DNA study.

**Literature**:

Christèle Baillif-Ducros et Nicolas Meyer 2020 : Pilleurs de tombes sur la colline du "Marxberg": études de cas au sein de la nécropole de l'Antiquité tardive de *Pons Saravi* (Sarrebourg, Moselle, France). In A. A. Noterman et M. Cervel (Eds), *Ritualiser, gérer, piller* (pp.67-71). Publication du Gaaf n°9

Christèle Baillif-Ducros et Nicolas Meyer 2017 : *Pilleurs de tombes sur la colline du "Marxberg" : études de cas au sein de la nécropole de l'Antiquité tardive de Pons Saravi (Sarrebourg, Moselle, France)* [Poster], Rencontre autour des réouvertures de tombes et de la manipulation des ossements, 9e rencontre du Gaaf, 10-12 mai 2017, Poitiers (France)

Nicolas Meyer et Christèle Baillif-Ducros 2016 : *La nécropole du "Marxberg" de Pons Saravi/Sarrebourg (Moselle, France)* [Poster], Colloque international ATEG V, 7-9 décembre 2016, Strasbourg (France)

| sample_id | sex (genetic) | date lower (95% CI) | date upper (95% CI) | date inference | burial_id | excluded | notes |
|---|---|---|---|---|---|---|---|
| R11550 | M | 257.5 calCE | 404.5 calCE | 14C | Marxberg Necropolis (Pons Saravi, Sarrebourg)_1088 | | |
| R11552 | M | 131 calCE | 413 calCE | SITE | Marxberg Necropolis (Pons Saravi, Sarrebourg)_1126 | | |
| R11553 | F | 131 calCE | 413 calCE | SITE | Marxberg Necropolis (Pons Saravi, Sarrebourg)_1127 passe1 | | REL_1ST of R11554 |
| R11554 | F | 131 calCE | 413 calCE | SITE | Marxberg Necropolis (Pons Saravi, Sarrebourg)_1127 S1 | Yes (Contam./ Related) | REL_1ST of R11553 |
| R11555 | F | 131 calCE | 413 calCE | SITE | Marxberg Necropolis (Pons Saravi, Sarrebourg)_1127 S2 | Yes (Contam./ Related) | IDENTICAL to R11553 REL_1ST of 11554 |
| R11556 | F | 229.5 calCE | 329.5 calCE | 14C | Marxberg Necropolis (Pons Saravi, Sarrebourg)_1141 S1 | | |
| R11557 | M | 261.5 calCE | 413 calCE | 14C | Marxberg Necropolis (Pons Saravi, Sarrebourg)_1171 | | |
| R11558 | M | 257.5 calCE | 404.5 calCE | 14C | Marxberg Necropolis (Pons Saravi, Sarrebourg)_1173a S1 | | |
| R11559 | F | 131 calCE | 413 calCE | SITE | Marxberg Necropolis (Pons Saravi, Sarrebourg)_1173b S2 | Yes (Contam.) | |
| R11560 | M | 259 calCE | 409.5 calCE | 14C | Marxberg Necropolis (Pons Saravi, Sarrebourg)_1177 | | |
| R11561 | M | 131 calCE | 413 calCE | SITE | Marxberg Necropolis (Pons Saravi, Sarrebourg)_1179 | | |
| R11563 | M | 131 calCE | 243.5 calCE | 14C | Marxberg Necropolis (Pons Saravi, Sarrebourg)_1182 | | |

#### Vendeuil Caply Les Marmousets, France

**Collaborators**: Valérie Kozlowski, Sandra Legrand, Olivia Cheronet
**Coordinates**: 49.611502, 2.297416
**Individuals** (n = 1): R1874

The site of Vendeuil-Caply is spread over 130 hectares and appears to be the location of a large *vicus*. Large-scale construction started in the 1$^{st}$ century CE. Interest in this site dates back to 1574 when the first document reference to it was published by the Prince of Condée. However, modern excavations have taken place on this site following its rediscovery in an aerial photograph in 1955.
The site is flanked by two hills (the Catelet and the Calmont), separated by the Saint-Denis Valley. On the Catelet hill, a Roman camp was discovered as well as a *fanum* and a small theater. The center of the settlement was in the Saint Denis Valley, where numerous houses were uncovered. The valley is also where the large theater was situated.
A more recent excavation in Vendeuil Caply in 2008 revealed the necropolis of Les Marmousets dating from the fourth to the fifth centuries composed of 122 graves. The individuals buried there are thought to be the inhabitants of the Saint-Denis valley on the site of Vendeuil-Caply.
The individual analyzed was buried in tomb 75. This was an intact inhumation in a pit carved in chalk, oriented NW-SE. The individual was laying on its back with the right forearm folded on the chest and the left on the abdomen. The legs were stretched out. No grave goods were associated with this individual. The grave is in a zone of the necropolis where wooden sarcophagi were used suggesting a proto-merovingien date (end of the third to the start of the fourth centuries CE). It is also a zone of the necropolis where the use of Charon's obols occurred in some inhumations.

| sample_id | sex (genetic) | date lower (95% CI) | date upper (95% CI) | date inference | burial_id |
|---|---|---|---|---|---|
| R1874 | F | 255 calCE | 402.5 calCE | 14C | T.75 V.C. 60120 |

**Literature**:

Dufour, Gérard, and Daniel Piton. "Vendeuil-Caply, une agglomération antique, anonyme et disparue." Revue archéologique de Picardie, vol. 3, no. 1, 1984, pp. 283–94, https://doi.org/10.3406/pica.1984.1450.

#### Hassleben, Germany

**Collaborators**: Jan Novacek, Diethard Walter
**Coordinates**: 51.109, 10.9956
**Individuals** (n = 7): R11866, R11867, R11868, R11872, R11873, R11875, R11877

Haßleben Necropolis

The Germanic necropolis of Haßleben (District Sömmerda) has been excavated by Armin Möller, curator of the former City Museum of Prehistory Weimar between 1910 and 1934 during several excavation seasons. The site has been identified due to surface gravel mining southeast of the town. The site is located about 15 km north to Erfurt in the lowland of the river Gera in the middle of the Thuringian Basin. Due to the destruction by the surface mine, the necropolis was not excavated completely. The investigated 24 graves are mainly oriented from north to south with few exceptions to the west and east. The cemetery was founded in the second half of the third century CE (phase C2 according to EGGERS). Although cremation was customary during this time, all the graves were inhumations. Several rich burials – partly equipped with imports from Roman provincial products (especially grave 8 and 21) – show parallels to "Fürstengräber" ("princely tombs") from Leuna, Stráže, Sakrau, and Himlingøje, which are accounted as belonging to Germanic *nobiles*.
On the burial ground of Haßleben, 22 individuals were identified. Among them, two individuals died during or shortly after birth, another two individuals died at less than 5 years of age. Further three individuals, two of them females, died in late juvenile, or early adult age. The individual from the grave 8, the so-called "Princess of Haßleben", was a female of about 20-30 age-at-death. Another two males and four females were adults at their time of death. Further two males and one female died as late adult or early mature age, as well as two males and one female were mature. The last two individuals were males, who died either at late mature or senile age. This distribution indicates a normal population with a rather low fraction of subadults, but high number of juvenile individuals, however, the low total number of individuals prevents any statistically robust conclusion. Dental pathologies, osteoarthritis, as well as cases of orbital cribra and meningeal reactions were the most common pathologies found in the skeletons from Haßleben.

| sample_id | sex (genetic) | date lower (95% CI) | date upper (95% CI) | date inference | burial_id | excluded |
|---|---|---|---|---|---|---|
| R11866 | F | 200 CE | 700 CE | ARCH | Haß 151 (4782), Haßleben, box 12362, 18.11.2019 | |
| R11867 | F | 200 CE | 700 CE | ARCH | Haß 235, Haß 238, Haßleben SÖM 71-1-3, box 12369, Grab 21, 19.11.2019 | |
| R11868 | M | 200 CE | 700 CE | ARCH | Haß 187 (6167), box 12359, Grab 17, 19.11.2019 | |
| R11872 | M | 200 CE | 700 CE | ARCH | Haßleben SÖM 70-4-4, Haß 217 (7233), Grab 20, left petrous, 19.11.2019 | |
| R11873 | F | 200 CE | 700 CE | ARCH | Haßleben SÖM 71-1-2, Grab 9, Haß 145 (4748), boc 12365, 19.11.2019 | |
| R11875 | M | 200 CE | 700 CE | ARCH | Haß 129 (4257), Haßleben, box 12361, 18.11.2019 | Yes (Contam.) |
| R11877 | F | 200 CE | 700 CE | ARCH | Fürstengrab von Haßleben, Grab 8, RI2; 19.11.2019 | |

**Literature**:

Behm-Blancke, G. (1973): Gesellschaft und Kunst der Germanen. Berlin 1973.

Birkenbeil, S.; Finke, L. (2004): Das Gräberfeld um die “Fürstin” von Hassleben (3. Jahrhundert). Unpublished anthropological report, TLDA Weimar.

Dušek, S. (1999): Haßleben. In: Reallexikon der germanischen Altertumskunde. Berlin/New York 1999, 41-43.

Schulz, W. (1933): Das Fürstengrab von Hassleben. Berlin 1933.

Weidenreich, F. (1933): Das Fürstengrab und das Grabfeld von Hassleben. In: Schulz, W., Das Fürstengrab von Hassleben, de Gruyter, Berlin.

#### Isola Sacra, Italy

**Collaborators**: Luca Bondioli, Alessia Nava, Alessandra Sperduti
**Coordinates**: 41.7621, 12.2491
**Individuals** (n = 11): R11109, R11111, R11112, R11113, R11115, R11116, R11117, R11118, R11119, R11120, R11121

The necropolis of Isola Sacra, located approximately 23 km southwest of Rome, was used by the inhabitants of *Portus Romae.* Developed during the Roman imperial time, *Portus* was a key trading center for Rome; through its docks, any kinds of goods from the farthest provinces of the Empire arrived in the capital. Isola Sacra extends approximately 1.5 km along the road between Ostia and *Portus Romae,* and comprises diverse burial structures, ranging from simple interments in sand to monumental tombs, dated between the 1st to the 3rd centuries AD. The inhabitants of *Portus* were middle-class administrators, traders, merchants, and sea workers often descended from slaves (Garnsey 1999). The excavation campaigns yielded over 2000 individuals that are currently stored at the Museo delle Civiltà in Rome. Most of the remains come from the so-called "fields of poor people" (Olivanti and Spanu 2018) and were extensively studied for their demographic profile (Sperduti 1995; Sperduti et al. 2012), aDNA (Antonio et al. 2019), migration (Prowse et al. 2007; Stark 2016), paleodiet (Prowse et al 2004, 2005, 2008), occupational markers (Crowe et al. 2010), metabolic stress of the nonadult segment (FitzGerald et al 2006; Nava et al. 2019, Rossi et al. 1999), and paleopathology (Bondioli et al. 2016, Hoover et al. 2005, Lockau et al. 2019).
As for the individuals analyzed in this study, the sample includes: **11117-SCR 136** a 6-12-month-old child buried in amphora; **11118-SCR 599** a 3-4-year-old child from a monumental tomb. Based on oxygen isotopic analyses (Prowse et al. 2007; Stark 2016), Three adult males (**11109-SCR 617**, **11116-SCR 174, 11119-SCR 303)** and one 16-18-year-old male (**11112-SCR 138**) are likely born elsewhere and can be considered immigrants. Four individuals, two adult males (**11111-SCR 448, 11121-SCR 325**) and two adult females (**1113-SCR 68**, **R11115-SCR 426**) are likely born locally. As for the other individuals (**11117-SCR 136, 11118-SCR 599, R11120-SCR 323**), oxygen isotopic data are not available.

| sample_id | sex (genetic) | date lower (95% CI) | date upper (95% CI) | date inference | burial_id | excluded |
|---|---|---|---|---|---|---|
| R11117 | M | 1 CE | 400 CE | ARCH | Isola Sacra_SCR136 | |
| R11118 | F | 1 CE | 400 CE | ARCH | Isola Sacra_SCR599 | |
| R11109 | M | 1 CE | 400 CE | ARCH | Isola Sacra_SCR617 | |
| R11116 | F | 1 CE | 400 CE | ARCH | Isola Sacra_SCR174 | |
| R11119 | M | 1 CE | 400 CE | ARCH | Isola Sacra_SCR303 | |

| | | | | | | |
|---|---|---|---|---|---|---|
| R11112 | M | 1 CE | 400 CE | ARCH | Isola Sacra_SCR138 | |
| R11111 | M | 1 CE | 400 CE | ARCH | Isola Sacra_SCR448 | |
| R11121 | M | 1 CE | 400 CE | ARCH | Isola Sacra_SCR325 | |
| R11113 | F | 1 CE | 400 CE | ARCH | Isola Sacra_SCR68 | |
| R11115 | F | 1 CE | 400 CE | ARCH | Isola Sacra_SCR426 | Yes |
| R11120 | F | 1 CE | 400 CE | ARCH | Isola Sacra_SCR323 | |

**Literature**:

Antonio M.L., Gao Z., Moots H.M., Lucci M., Candilio F., Sawyer S., Oberreiter V., Calderon D., Devitofranceschi K., Aikens R.C., Aneli S., Bartoli F., Bedini A., Cheronet O., Cotter D.J., Fernandes D.M., Gasperetti G., Grifoni R., Guidi A., La Pastina F., Loreti E., Manacorda D., Matullo G., Morretta S., Nava A., Fiocchi Nicolai V., Nomi F., Pavolini C., Pentiricci M., Pergola P., Piranomonte M., Schmidt R., Spinola G., Sperduti A., Rubini M., Bondioli L., Coppa A., Pinhasi R. & Pritchard J.K. 2019. Ancient Rome: A genetic crossroads of Europe and the Mediterranean. Science 366: 708-714.

Bondioli, L., Nava, A., Rossi, P.F. & Sperduti, A. 2016. Diet and health in Central-Southern Italy during the Roman Imperial time. ACTA IMEKO, 5(2), 19-25.

Crowe, F., Sperduti, A., O'Connell, T. C., Craig, O. E., Kirsanow, K., Germoni, P., Macchiarelli R., Garnsey P. & Bondioli, L. 2010. Water--related occupations and diet in two Roman coastal communities (Italy, first to third century AD): Correlation between stable carbon and nitrogen isotope values and auricular exostosis prevalence. Am. J. Phys. Anthropol., 142(3), 355-366.

FitzGerald, C.M., Saunders, S., Bondioli, L. & Macchiarelli, R. 2006. Health of infants in an Imperial Roman skeletal sample: perspective from dental microstructure. Am. J. Phys. Anthropol., 130(2), 179-189.

Garnsey, P. 1999. The people of Isola Sacra. In L. Bondioli & R. Macchiarelli (Eds.). Digital archives of human paleobiology. Osteodental biology of the people of Portus Romae (Necropolis of Isola Sacra, 2nd–3rd Cent. AD). I. Enamel microstructure and developmental defects of the primary dentition. Rome: National Prehistoric Ethnographic ‘‘L. Pigorini’’ Museum. Published on CD-ROM

Hoover, K. C., Corruccini, R. S., Bondioli, L. & Macchiarelli, R. 2005. Exploring the relationship between hypoplasia and odontometric asymmetry in Isola Sacra, an Imperial Roman necropolis. Am. J. Hum. Biol., 17(6), 752-764.

Lockau, L., Atkinson, S., Mays, S., Prowse, T., George, M., Sperduti, A., Bondioli L., Wood C., Ledger M. & Brickley, M.B. 2019. Vitamin D deficiency and the ancient city: Skeletal evidence across the life course from the Roman period site of Isola Sacra, Italy. Journal of Anthropological Archaeology, 55, 101069. doi:https://doi.org/10.1016/j.jaa.2019.101069

Olivanti, P. & Spanu, M. 2018. Necropoli dell'Isola Sacra, scavo 1988-1989: alcune riflessioni su occupazione degli spazi, cronologia delle sepolture, corredi. In M. Cébeillac-Gervasoni, N. Laubry, F. Zevi (a cura di) Ricerche su Ostia e il suo territorio. Atti del Terzo Seminario Ostiense, Collection de l'École française de Rome, pp. 67-77.

Nava, A., Frayer, D.W. & Bondioli, L. 2019 Longitudinal analysis of the microscopic dental enamel defects of children in the Imperial Roman community of Portus Romae (necropolis of Isola Sacra,2nd to 4th century CE, Italy). J. Archaeol. Sci Rep. 23, 406-415

Prowse, T., Schwarcz, H.P., Saunders, S., Macchiarelli, R. & Bondioli, L. 2004. Isotopic paleodiet studies of skeletons from the imperial Roman-age cemetery of Isola Sacra, Rome, Italy. J. Archaeol. Sci., 31(3), 259-272

Prowse, T.L., Saunders, S. R., Schwarcz, H.P., Garnsey, P., Macchiarelli, R. & Bondioli, L. 2008. Isotopic and dental evidence for infant and young child feeding practices in an Imperial Roman skeletal sample. Am. J. Phys. Anthropol., 137(3), 294-308

Prowse, T.L., Schwarcz, H.P., Garnsey, P., Knyf, M., Macchiarelli, R. & Bondioli L. 2007. Isotopic evidence for age--related immigration to imperial Rome. Am. J. Phys. Anthropol., 132(4), 510-519.

Prowse, T.L., Schwarcz, H.P., Saunders, S.R., Macchiarelli, R. & Bondioli, L. 2005. Isotopic evidence for age--related variation in diet from Isola Sacra, Italy. Am. J. Phys. Anthropol., 128(1), 2-13.

Rossi, P., Bondioli, L., Geusa, G. & Macchiarelli, R. 1999. Digital Archives of Human Paleobiology. Osteodental biology of the people of *Portus Romae* (Necropolis of Isola Sacra, 2nd–3rd Cent. AD). I. Enamel microstructure and developmental defects of the primary dentition. In: Bondioli, L., and Macchiarelli, R., editors. Digital Archives of Human Paleobiology. Rome: National Prehistoric Ethnographic ‘‘L. Pigorini’’ Museum.

Sperduti, A. 1995. I resti scheletrici umani della necropoli di età romano-imperiale di Isola Sacra (I-III sec. d. C.): analisi paleodemografica (Unpublished doctoral dissertation). Sapienza University of Rome, Rome

Sperduti, A., Bondioli, L. & Garnsey, P. 2012. Skeletal evidence for occupational structure at the coastal towns of Portus and Velia (1st–3rd c. AD). J. Roman Archaeol., S.91, 53-70

Stark, R.J. 2016. Ancient lives in motion: a bioarchaeological examination of stable isotopes, nonmetric traits, and human mobility in an imperial roman context (1st-3rd c. Ce). Doctoral dissertation. McMaster University, Hamilton

#### Palazzo della Cancelleria, Italy

**Collaborators**: Alfredo Coppa, Michaela Lucci
**Coordinates**: 41.8965, 12.4719
**Individuals** (n = 5): R1223, R1225, R1291, R1292, R1294

Cancelleria - The Basilica of San Lorenzo in Damaso

The Basilica of San Lorenzo was erected by Pope Damaso (366-384 CE) in south-western Campo Marzio, reusing part of an architectural complex in which it is possible to recognize the buildings of the *factio prasina*, one of the four factions of the circus. The Basilica, with three naves, occupied a large area largely coinciding with that of the courtyard of the Palazzo della Cancelleria, in one of the most central areas of Rome, halfway between Piazza Farnese and Piazza Navona.
Probably as early as the sixth century CE there are numerous burials (subsequently reworked several times) which were carried out in the area of the church, in particular in a vast environment located close to the south side of the building.
A radical transformation of the Basilica is recorded in the second quarter of the 11th century CE following a fire, of which extensive traces have been found. In addition to conspicuous transformations of a structural nature, the floor of all the sections of the Basilica was raised by about 1 m. In the church, starting from this date until its destruction, numerous burials were built including several masonry ossuaries. New changes to the structure of the church were made during the second half of the fifteenth century. The numismatic artifacts found have allowed us to date, at the beginning of the last quarter of the fifteenth century, a large mass grave in which hundreds of burials were deposited (SU17, SU 30 and SU 471). In the way of organizing the burials it is likely to recognize the effects of a plague epidemic which we know to have struck the city between 1476 and 1479 CE, a hypothesis that would also be confirmed by the study of skeletal remains. In 1489 CE the building of the Palazzo della Cancelleria began and the church was totally destroyed. The population of this necropolis covers most of the Middle Ages and is representative of the population of Rome of this period.

| sample_id | sex (genetic) | date lower (95% CI) | date upper (95% CI) | date inference | burial_id |
|---|---|---|---|---|---|
| R1223 | M | 1480 CE | 1490 CE | ARCH | Cancelleria US 17 17 |
| R1225 | M | 1480 CE | 1490 CE | ARCH | Cancelleria US 17 37 |
| R1291 | M | 500 CE | 1400 CE | ARCH | Palazzo della Cancelleria US 530 |
| R1292 | M | 500 CE | 1400 CE | ARCH | Palazzo della Cancelleria US 535 |
| R1294 | M | 500 CE | 1400 CE | ARCH | Palazzo della Cancelleria US 626 |

**Literature**:

Coppa A., Cucina A., Mancinelli D., Lucci M., Vargiu R., Aspetti demografici e nutrizionali nella Roma della seconda metà del ‘400: le sepolture della chiesa di S. Lorenzo in Damaso al Palazzo della Cancelleria. In Sonnino E. (Ed.): Popolazione e Società a Roma dal Medioevo dell’Età Contemporanea, 1998, Editrice il Calamo, Roma, pp. 435-481.

Frommel Ch. L., Pentiricci M. (Eds.), L’antica basilica di San Lorenzo in Damaso, I (Gli scavi), II (I materiali), Roma 2009.

Pentiricci M., S. Lorenzo in Damaso, in M. Cecchelli (Ed.), Materiali e tecniche dell’edilizia paleocristiana a Roma dal IV al VII secolo d.C., Roma 2001.

Salamon M., Coppa A., Mccormick M., Rubini M., Vargiu R., Tuross N., The consilience of historical and isotopic approaches in reconstructing the medieval Mediterranean diet. Journal of Archaeological Science, 2008, 35:1667-1672

Vargiu R., Cucina A., Mancinelli M., Lucci M., Coppa A, Condizioni di vita e paleonutrizione a Roma nella seconda metà del ‘400. In: Gli Scavi di San Lorenzo in Damaso Volume II – I materiali, C.L. Frommel e M. Pentiricci Eds., De Luca Editore, Roma, 2009, pp. 547-571. ISBN 978-88-8016-925-3.

### Urbino Bivio, Italy

**Collaborators**: Alfredo Coppa, Michaela Lucci
**Coordinates**: 43.7237164, 12.6346536
**Individuals** (n = 4): R1554, R1555, R1556, R1557

Bivio della Croce dei Missionari, Urbino (Pesaro): Roman necropolis
Francesca Candilio

The Roman necropolis of Urbino “Bivio della Croce dei Missionari” was discovered in 1972 during road works. The excavations uncovered a very densely used funerary area in which many of the graves overlapped or cut into previously dug ones. Inhumations and cremations were present in almost equal proportions and included simple pits or earth graves as well as graves lined in tiles or delimited by walls. 92 of the graves (48 inhumations and 44 cremations) were described, on the basis of dig reports, by Mercando et al in 1982. A health assessment was conducted in 2009 by Paine et al. pooling the individuals with those of a coeval necropolis discovered approximately half a kilometer away that are considered to be the same community and cultural period (Mercando et al. 1982; Paine et al. 2009).
The individuals, dated archaeologically to the 1st-3rd century A.D. (Mercando et al. 1982), appear to have been common people that had, on average, chronic ailments and a short life span (Paine et al. 2009).

**T. 68** is described (Mercando et al. 1982) as a single earthen grave with an E-W orientation. The disposition of the bones in the image provided and the indication of the presence of iron nails along the margins of the grave suggest the presence of a perishable, possibly wooden, casket. Grave goods included a bronze coin and pottery fragments. Archaeological indicators place the burials to the last quarter of the 1st century BC.

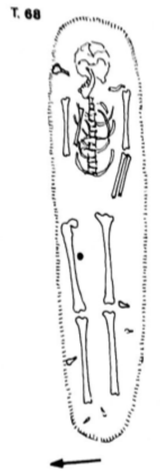

**T. 73** held a single, badly preserved individual placed in an E-W orientation in a simple earthen grave. Grave goods included a bronze coin positioned near the individual's right forearm and iron nails along the tomb's margins. The individual's arms were flexed above the pelvis with the left positioned above the right (Mercando et al. 1982).

**T. 86** Inhumation oriented NE-SW. Grave goods included portions of four glass unguentariums and five iron nails. Archaeological indicators date the individual to the 2nd century AD. (Mercando et al. 1982).

| sample_id | sex (genetic) | date lower (95% CI) | date upper (95% CI) | date inference | burial_id |
|---|---|---|---|---|---|
| R1557 | M | 81.5 calCE | 220.5 calCE | SITE | Urbino T.68 R. petrous 3.5.18 |
| R1554 | M | 125.5 calCE | 220.5 calCE | 14C | Urbino Bivio T.73 R. petrous 3.5.18 |
| R1555 | F | 81.5 calCE | 210 calCE | 14C | Urbino Bivio T. 86 R.petrous 3.5.18 |
| R1556 | M | 81.5 calCE | 220.5 calCE | SITE | Urbino T.100 L. petrous 3.5.18 |

**Literature**:
Mercando, L., Luni, M., Taborelli, L., Sorda, S., Corrain, C., Erspamer, G., Malgeri, G. 1982. Urbino (Pesaro). Necropoli romana. Tombe al Bivio della Croce dei Missionari e a S. Donato. *Notizie degli Scavi di Antichità*, 36:109-420.
Paine, R.R., Vargiu, R., Signoretti, C., Coppa, A. 2009. *A health assessment for Imperial Roman burials recovered from the necropolis of San Donato and Bivio CH, Urbino, Italy*. *Journal of Anthropological Sciences*, 87: 193-210

#### Chhim, Lebanon

**Collaborators**: Tomasz Waliszewski, Arkadiusz Sołtysiak
**Coordinates**: 33.622519, 35.499985
**Individuals** (n = 5): R3473, R3476, R3477, R9818, R9823

Chhim (central Phoenicia) : Burial B.V.3

The ruins of a village dated to the 1st – 7th century AD are situated approx. 35 km south of Beirut in Lebanon and 14 km north of Saida, the famous Phoenician metropolis of Sidon. The village located in the mountains ca. 8 km from the shore of the Mediterranean Sea consisted of a sanctuary active between the 1st and the 4th century, a church built in 498 AD, houses, and at least five large oileries. Subsistence of the village was based on agriculture (especially olive oil production) and animal husbandry (especially sheep and goat). The population of the village is estimated at around 300-400 individuals (Waliszewski & Wicenciak, 2015). The project at Chhim was conducted between 1996 and 2016 by a team from the Polish Center of Mediterranean Archaeology of the University of Warsaw.
The tomb B.V.3 was uncovered during the 1998 season in the southern end of the narthex of a Christian basilica, next to the entrance to the southern aisle and the dedicatory Greek inscription from 498 AD. The flat stone slabs closing it were lifted to reveal a rectangular outline (ca. 1.90 x 0.60 m (west) x 0.42 m (east)) narrowing to the east, lined with stone slabs set up on end (depth of the tomb c. 0.64 – 0.68 m).
The tomb was cut into the well preserved, repeatedly renovated lime mortar floor of the narthex and thus belongs to its late phase. Inside, there were several human skulls, painstakingly arranged along the west side, and the other bones also arranged according to their anatomical type. On the very bottom there was a rather complete skeleton, perhaps laid to rest at an earlier date. The modest furnishing included a few rings, pendants, an earring and two gold coins of emperor Phocas (gold semissis minted at Constantinople, dated to AD 607-610), both found on the floor of the grave near the pelvis of the complete skeleton. The coins might be evidence for the dating of the last phase before the secondary mass burial. The burial seems to be dated generally to the 7th c. AD, probably its first half (Waliszewski, 1999).
The minimum number of individuals buried in the tomb B.V.3 is 31 (based on right femora); 15 adult and 16 subadult individuals. Any reliable sex assessment was possible for eight individuals and all of them were males. Subadult individuals represented different age-at-death categories from infants to adolescents. Most elements were very well preserved and also minor bones (as carpals and phalanges) are present, which suggests that these skeletons were very carefully transported from the original deposition place to the final burial in the basilica (Sołtysiak & Waliszewski, 2021).

| sample_id | sex (genetic) | date lower (95% CI) | date upper (95% CI) | date inference | burial_id | excluded | notes |
|---|---|---|---|---|---|---|---|
| R3473 | F | 650.5 calCE | 759.5 calCE | 14C | CHHIM 1998; B-Nanitx (?); (58) | | |
| R3476 | M | 435.5 calCE | 575.0 calCE | 14C | CHHIM 1998; B-Nanitx; TEETH (3) | | IDENTICAL to R3477 |
| R3477 | M | 434.5 calCE | 565 calCE | 14C | CHHIM 1998; B-Nanitx; SKULL A | Yes (Related) | IDENTICAL to R3476 |
| R9818 | M | 577 calCE | 642.5 calCE | 14C | Chhim, B 7.(10) | Yes (Contam./ Related) | IDENTICAL to R9823 |
| R9823 | M | 599.5 calCE | 648 calCE | 14C | Chhim, B teeth (2) | | IDENTICAL to R9818 |

<u>**Literature**</u>:

Sołtysiak, A., & Waliszewski, T. (2021), Human remains from Chhîm, Lebanon, 1998–2009. Bioarchaeology of the Near East, 15, 104-110.

Waliszewski, T. (1999). Chhim explorations, 1998. Polish Archaeology in the Mediterranean, 10, 177–185.

Waliszewski, T., & Wicenciak, U. (2015). Chhim, Lebanon. A Roman and Late Antique village in the Sidon hinterland. Journal of Eastern Mediterranean Archaeology and Heritage Studies, 3(4), 372–395.

### Ej-Jaouze, Lebanon

**Collaborators**: Lina Nacouzi, Marie Laguardia
**Coordinates**: 33.8522, 35.8292
**Individuals** (n = 4): R12229, R12233, R12245, R12246

The Tombs of Ej-Jaouze, Lebanon

The Ej-Jaouze site is located east of Beirut, on Mount Lebanon, at an altitude of 1400m. It is an isolated site and difficult to access, but the natural resources of the region and its strategic position along an old road linking Beirut to Damascus have contributed to its location. Poorly known, the Lebanese mountain is often associated with a deserted region, reserved only for cult, attested by the presence of many Roman temples. The archaeological excavations carried out at Ej-Jaouze have revealed a village composed of a hill housing dwellings and a lower town with monumental buildings. The site presents a long period of occupation, the first traces of which date back to the early Bronze Age and extend to the medieval period (12th-14th century) with a hiatus from the 7th to the 12th century.

During excavations, four tombs were discovered inside buildings. Three of them were excavated as collective graves containing a total of 57 individuals, of both sexes and all ages. The ceramics and grave goods found inside have been used to date the tombs between the 5th and 7th centuries.
The discovery of these tombs shows the particularity of the funerary practices of Ej-Jaouze and testifies to the interest of the studies carried out in the Lebanese mountains.
The bones, currently stored at the French Institute of Near East in Beirut, were the subject of a preliminary study published in Nacouzi et al. 2018.

| sample_id | sex (genetic) | date lower (95% CI) | date upper (95% CI) | date inference | burial_id | excluded |
|---|---|---|---|---|---|---|
| R12229 | M | 431 calCE | 545 calCE | 14C | Ej Jaouze _K3t1_DH10 | |
| R12233 | F | 432.5 calCE | 547.5 calCE | 14C | Ej Jaouze _F2t1_#2 | |
| R12245 | F | 592.5 calCE | 647 calCE | 14C | Ej Jaouze _A2t1_#2 | |
| R12246 | F | 558.5 calCE | 639 calCE | 14C | Ej Jaouze _A2t1_#3 | Yes (Contam.) |

**Literature**:

Nacouzi L. 2004. Ej-Jaouze (MEtn, Liban). Mission de 2003. *Bulletin d'Archéologie et d'Architecture Libanaises 8*, p. 211-247.

Nacouzi L. et al. 2016. Ej-Jaouze (Metn). Rapport des travaux menés en 2012-2013. *Bulletin d'Archéologie et d'Architecture Libanaises 16*, p.131-192.
Nacouzi L. et al. 2018. Ej-Jaouze (Metn). Rapport sur les travaux menés en 2014, 2015 et 2016. *Bulletin d'Archéologie et d'Architecture Libanaises 18,* p. 79-200.

#### Bailuliai, Lithuania

**Collaborator**: Rimantas Jankauskas
**Coordinates**: 54.687157, 25.279652
**Individuals** (n = 1): R10840

The site was excavated in 1999 and 2000. Archaeologists dated it according to artefacts 5th-6th c. AD (400-600 AD), East Lithuanian Barrow culture. The site consists of 15 barrows, containing 4 inhumations (1 barrow – 1 inhumation) and 11 barrows with cremations (from 1 to 3 in each).
The site was most likely used only briefly. The studied grave was one of the poorest in artefacts (bronze and iron, some glass beads), but there are quite clear connections according to artefacts with other very rich burials, having non-local features. Such a mixture of burial customs (inhumations and cremations) suggests transition from one ideology to another.
East Lithuanian Barrow culture – local development: As Eastern Lithuania has poor soils, covered with mostly pine forest vegetation, it was relatively sparsely populated, with an economy based on slash-and-burn, and mobile small settlements that were periodically moved to another site after soil exploitation. It seems the site was rather isolated. Sites such as Marvelė could be of interest to scouts or merchants from Roman provinces for being close to two large rivers (waterways) and for the deep hinterland on the Amber route. Areas east from there (East Lithuanian Barrow culture developed from earlier Brushed pottery culture) were of absolutely no interest for them – these areas could offer nothing valuable.

| sample_id | sex (genetic) | date lower (95% CI) | date upper (95% CI) | date inference | burial_id |
|---|---|---|---|---|---|
| R10840 | M | 262 calCE | 415 calCE | 14C | BAILULIAI_10/1_I391A |

#### Marvelė, Lithuania

**Collaborator**: Rimantas Jankauskas
**Coordinates**: 54.8985, 23.9036
**Individuals** (n = 4): R10830, R10832, R10836, R10838

The Marvelė necropolis

In the years 1991-2011, a cemetery area of nearly 3 ha was excavated, where ca. 1590 inhumation and cremation human burials with abundant grave goods were found and examined. About 300 horse skeletal remains with numerous artifacts were also uncovered.

The burial ground is located on the left bank of the Nemunas River within the current borders of Kaunas City. It was an open place surrounded by water from three sides. The Marvelė burial ground consists of three sites: (A) the larger southeastern part with skeletal and cremation human burials and horse burials dated to the Roman Period and Viking Age (2nd-13th cent.); (B) the southwestern part is located 120 m away from the main part (dozens of skeletal human burials, multiple cremation human and horse burials) dated to the Migration Period and Viking Age (6-12th cent.); (C) a small group of cremation graves dated to the Migration Period (5-7th cent.) was investigated on the northern edge of the necropolis. There are no cases of later burials destroying earlier graves. This fact confirms the idea that there should have been clear and enduring memory signs in the area of the necropolis. All Marvelė samples studied here were dated to the years 350-450.

Leading archeologists were Audrius Astrauskas and Mindaugas Bertašius. They authored the majority of bibliographic studies on the Marvelė necropolis. A part of these studies are unpublished reports of excavations. Many others are very brief information texts on field research published earlier biannually and now annually. A number of articles deal with burial customs, change of cultural orientation, cremation burials, burials of horses, numismatic questions, social structure of community, and multidimensional analysis of iron artefacts. Several monographs appeared as well. They cover issues, such as: social and cultural development of prehistoric Marvelė community, overview of Central Lithuania in the $8^{th}$-$12^{th}$ cent.

Few other researchers published articles on this necropolis, including work on bioarcheology (Jankauskas 2005, 2009; Jankauskas, Barkus 1994), and zooarcheology (Daugnora 1994; Bertašius, Daugnora 1997, 2001).

| sample_id | sex (genetic) | date lower (95% CI) | date upper (95% CI) | date inference | burial_id |
|---|---|---|---|---|---|
| R10830 | F | 402.5 calCE | 533.5 calCE | 14C | MARVELE_316_I128 |
| R10832 | F | 402.5 calCE | 533.5 calCE | SITE | MARVELE_325_164 |
| R10836 | M | 402.5 calCE | 533.5 calCE | SITE | MARVELE_312_177 |
| R10838 | M | 402.5 calCE | 533.5 calCE | SITE | MARVELE_194_I133 |

**Literature**:


Astrauskas, A. 1994a. Marvelės kapinyno (II-VII a. kapai) tyrinėjimai 1992 ir 1993 metais. Archeologiniai tyrinėjimai Lietuvoje 1992 ir 1993 metais, 120-124

Astrauskas, A. 1994b. Laidojimo papročiai Marvelės kapinyne. In: A. Astrauskas (Ed.). Vidurio Lietuvos archeologija: konferencijos medžiaga. Vilnius: Žalioji Lietuva, 21-27

Astrauskas, A. 1995. Marvelės pilkapiai. In: M. Michelbertas (Ed.). Baltų archeologija: naujausių tyrimų rezultatai. Vilnius: VU leidykla, 30-34

Astrauskas, A. 1996a. Marvelės kapinyno senojo ir viduriniojo geležies amžių kapų tyrinėjimai 1994 ir 1995 metai. Archeologiniai tyrinėjimai Lietuvoje 1994 ir 1995 metais, 93-95

Astrauskas, A. 1996b. Vidurio Lietuvos gyventojų kultūrinės orientacijos kaita SGA-VIGA. In: A. Astrauskas, M. Bertašius (Eds.). Vidurio Lietuvos archeologija. Etnokultūriniai ryšiai: konferencijos medžiaga. Vilnius: Žalioji Lietuva, 4-10

Astrauskas, A. 1998a. Marvelės bendruomenė II a. pabaigoje – V a. Vilnius: Vilnius University. Unpublished doctoral dissertation

Astrauskas, A. 1998b. Marvelės (Kauno miestas) kapinyno senojo ir viduriniojo geležies amžių kapų tyrinėjimai 1996 ir 1997 metai. Archeologiniai tyrinėjimai Lietuvoje 1996 ir 1997 metais, 173-175

Astrauskas, A., Bertašius, M. 1992a. Marvelės kapinyno 1991 metų tyrinėjimų ataskaita. I dalis. Lietuvos istorijos instituto rankraštynas, f. 1, b. 1831. Vilnius. Unpublished report

Astrauskas, A., Bertašius, M. 1992b. Marvelės kapinyno 1991 metų tyrinėjimų ataskaita. II dalis. Lietuvos istorijos instituto rankraštynas, f. 1, b. 1831a. Vilnius. Unpublished report

Astrauskas, A., Bertašius, M. 1992c. Marvelės kapinyno 1991 metų tyrinėjimų ataskaita. III dalis. Lietuvos istorijos instituto rankraštynas, f. 1, b. 1832b. Vilnius. Unpublished report

Astrauskas, A., Bertašius, M. 1992d. Marvelės kapinyno tyrinėjimai 1991 m. Archeologiniai tyrinėjimai Lietuvoje 1990 ir 1991 metais, 1, 90-94

Astrauskas, A., Bertašius, M. 1993a. Marvelės kapinyno 1992 metų tyrinėjimo ataskaita. I dalis. Lietuvos istorijos instituto rankraštynas, f. 1, b. 1963. Vilnius. Unpublished report

Astrauskas, A., Bertašius, M. 1993b. Marvelės kapinyno 1992 metų tyrinėjimo ataskaita. II dalis (plotai Nr.24-69) /brėžiniai/. Lietuvos istorijos instituto rankraštynas, f. 1, b. 1964. Vilnius. Unpublished report

Astrauskas, A., Bertašius, M. 1993c. Marvelės kapinyno 1992 metų tyrinėjimo ataskaita. III dalis (plotai Nr.70-114) /brėžiniai/. Lietuvos istorijos instituto rankraštynas, f. 1, b. 1965. Vilnius. Unpublished report
Astrauskas, A., Bertašius, M. 1993d. Marvelės kapinyno 1992 metų tyrinėjimo ataskaita. IV dalis (k. Nr. 68-215 įkapės) /piešiniai/. Lietuvos istorijos instituto rankraštynas, f. 1, b. 1966. Vilnius. Unpublished report
Astrauskas, A., Bertašius, M. 1993e. Marvelės kapinyno 1992 metų tyrinėjimo ataskaita. V dalis (k. Nr. 218-348) /piešiniai/. Lietuvos istorijos instituto rankraštynas, f. 1, b. 1967. Vilnius. Unpublished report
Astrauskas, A., Bertašius, M. 1993f. Marvelės kapinyno 1992 metų tyrinėjimo ataskaita. VI dalis (fotonuotraukos). Lietuvos istorijos instituto rankraštynas, f. 1, b. 1968. Vilnius. Unpublished report
Astrauskas, A., Bertašius, M. 1993g. Marvelės kapinyno /Kaunas/ 1992 metų tyrinėjimų ataskaita IX dalis (Marvelės “žirgyno” medžiaga). Lietuvos istorijos instituto rankraštynas, f. 1, b. 2506. Vilnius. Unpublished report
Astrauskas, A., Bertašius, M. 1994a. Marvelės kapinyno 1993 m. tyrinėjimo ataskaita. I dalis. Lietuvos istorijos instituto rankraštynas, f. 1, b. 2213. Vilnius. Unpublished report
Astrauskas, A., Bertašius, M. 1994b. Marvelės kapinyno 1993 m. tyrinėjimo ataskaita. II dalis (fiksacijos brėžiniai). Lietuvos istorijos instituto rankraštynas, f. 1, b. 2215. Vilnius. Unpublished report
Astrauskas, A., Bertašius, M. 1994c. Marvelės kapinyno 1993 m. tyrinėjimo ataskaita. III dalis (radinių piešiniai). Lietuvos istorijos instituto rankraštynas, f. 1, b. 2214. Vilnius. Unpublished report
Astrauskas, A., Bertašius, M. 1994d. Marvelės kapinyno 1993 m. tyrinėjimo ataskaita. IV dalis. Lietuvos istorijos instituto rankraštynas, f. 1, b. 2216. Vilnius. Unpublished report
Astrauskas, A., Bertašius, M. 1995a. Marvelės kapinyno /Kaunas/ 1994 m. tyrinėjimų ataskaita. Lietuvos istorijos instituto rankraštynas, f. 1, b. 2413. Vilnius. Unpublished report
Astrauskas, A., Bertašius, M. 1995b. Marvelės kapinyno /Kaunas/ 1994 m. tyrinėjimų ataskaita. II dalis. Lietuvos istorijos instituto rankraštynas, f. 1, b. 2414. Vilnius. Unpublished report
Astrauskas, A., Bertašius, M. 1995c. Marvelės kapinyno /Kaunas/ 1994 m. tyrinėjimų ataskaita. III dalis. Lietuvos istorijos instituto rankraštynas, f. 1, b. 2415. Vilnius. Unpublished report
Astrauskas, A., Bertašius, M. 1995d. Marvelės kapinyno /Kaunas/ 1994 m. tyrinėjimų ataskaita. IV dalis. Lietuvos istorijos instituto rankraštynas, f. 1, b. 2416. Vilnius. Unpublished report
Astrauskas, A., Bertašius, M. 1996a. Marvelės kapinyno /Kaunas/ 1995 m. tyrinėjimo ataskaita. I dalis. Lietuvos istorijos instituto rankraštynas, f. 1, b. 2569. Vilnius. Unpublished report

Astrauskas, A., Bertašius, M. 1996b. Marvelės kapinyno /Kaunas/ 1995 m. tyrinėjimo ataskaita. II dalis. Lietuvos istorijos instituto rankraštynas, f. 1, b. 2570. Vilnius. Unpublished report

Astrauskas, A., Bertašius, M. 1997. Marvelės kapinyno /Kauno m./ 1996 m. tyrinėjimų ataskaita. Lietuvos istorijos instituto rankraštynas, f. 1, b. 2754. Vilnius. Unpublished report

Astrauskas, A., Bertašius, M. 1998a. Marvelės kapinyno 1997 m. tyrinėjimų ataskaita, 1 dalis /tekstas, plotų fiksacijos brėžiniai/. Lietuvos istorijos instituto rankraštynas, f. 1, b. 2904. Vilnius. Unpublished report

Astrauskas, A., Bertašius, M. 1998b. Marvelės kapinyno 1997 m. tyrinėjimų ataskaita, 2 dalis /kapų, radinių piešiniai, foto nuotraukos/. Lietuvos istorijos instituto rankraštynas, f. 1, b. 2905. Vilnius. Unpublished report

Astrauskas, A., Bertašius, M. 2010. Marvelės kapinyno 1998 m. tyrinėjimų ataskaita. Lietuvos istorijos instituto rankraštynas, b. 5265. Vilnius. Unpublished report

Bertašius, M. 1992. Kauno m. vandenvalos įrengimų teritorijos žvalgomųjų archeologinių tyrimų ataskaita /1991 m., Marvelės kapinyno vietoje/. Lietuvos istorijos instituto rankraštynas, f. 1, b. 1890. Kaunas. Unpublished report

Bertašius, M. 1994a. Marvelės degintiniai kapai. In: A. Astrauskas (Ed.). Vidurio Lietuvos archeologija: konferencijos medžiaga. Vilnius: Žalioji Lietuva, 56-69

Bertašius, M. 1994b. Vėlyvojo geležies amžiaus kapai Marvelės kapinyne. Archeologiniai tyrinėjimai Lietuvoje 1992 ir 1993 metais, 128-132

Bertašius, M. 1995. Marvelės žirgų kapai. In: M. Michelbertas (Ed.). Baltų archeologija: naujausių tyrimų rezultatai. Vilnius: VU leidykla, 30-34

Bertašius, M. 1996. 1994-1995 metais tyrinėti Marvelės degintiniai kapai. Archeologiniai tyrinėjimai Lietuvoje 1994 ir 1995 metais, 99-101

Bertašius, M. 1998a. Marvelės kapinyno VIII-XII amžiaus kapų tyrinėjimai 1996-1997 metais. Archeologiniai tyrinėjimai Lietuvoje 1996 ir 1997 metais, 184-187

Bertašius, M. 1998b. Žirgų kapai – vienas Kauno proistorės klausimas. Kauno istorijos metraštis, 1, 12-24

Bertašius, M. 1998c. The Ritual of Horse Offer in Middle Lithuania. Abstract Book. 4th Annual Meeting. Göteborg. September 23-27, 170

Bertašius, M. 1999. Marvelės kapinynas. 1999 metų tyrinėjimų ataskaita. Lietuvos istorijos instituto rankraštynas, f. 1, b. 3401. Kaunas. Unpublished report

Bertašius, M. 2000a. Marvelės kapinyno tyrinėjimai. Archeologiniai tyrinėjimai Lietuvoje 1998 ir 1999 metais, 248-251

Bertašius, M. 2000b. Kauno apylinkių teritorinių struktūrų formavimosi ypatybės. Kauno istorijos metraštis, 2, 5-29

Bertašius, M. 2001a. Marvelės kapinynas. 2000 metų tyrinėjimų ataskaita. Lietuvos istorijos instituto rankraštynas, f. 1, b. 3617. Kaunas. Unpublished report

Bertašius, M. 2001b. Vikingiškosios tarpregioninės kultūros atspindžiai Marvelės kapinyne Kaune. Lietuvos archeologija, 21, 193–202
Bertašius, M. 2002a. Marvelės kapinynas. 2001 metų tyrinėjimų ataskaita. Lietuvos istorijos instituto rankraštynas, f. 1, b. 3811. Kaunas. Unpublished report
Bertašius, M. 2002b. Marvelės kapinynas. Archeologiniai tyrinėjimai Lietuvoje 2000 metais, 91-95
Bertašius, M. 2002c. Marvelės kapinynas. Archeologiniai tyrinėjimai Lietuvoje 2001 metais, 114-116
Bertašius, M. 2002d. Vidurio Lietuva VIII-XII a. Kaunas: VDU leidykla
Bertašius, M. 2003. Marvelės kapinynas. 2002 metų tyrinėjimų ataskaita. Lietuvos istorijos instituto rankraštynas, f. 1, b. 3982. Kaunas. Unpublished report
Bertašius, M. 2004a. Marvelės kapinynas. 2003 metų tyrinėjimų ataskaita. Lietuvos istorijos instituto rankraštynas, f. 1, b. 4257. Kaunas. Unpublished report
Bertašius, M. 2004b. Vakarų Europos monetos Marvelės kapuose. Numizmatika, metraštis 2001-2002, 2-3, 169-174
Bertašius, M. 2005a. Marvelės kapinynas. Archeologiniai tyrinėjimai Lietuvoje 2002 metais, 92-93
Bertašius, M. 2005b. Marvelės kapinynas. Archeologiniai tyrinėjimai Lietuvoje 2003 metais, 113-114
Bertašius, M. 2005c. Marvelė. Ein Gräberfeld Mittellitauens. I band. Vidurio Lietuvos aukštaičių II-XII a. kapinynas: monografija. Kaunas: Kauno technologijos universitetas Humanitarinių mokslų fakultetas
Bertašius, M. 2006a. Marvelės kapinynas. 2005 metų tyrinėjimų ataskaita. Lietuvos istorijos instituto rankraštynas, f. 1, b. 4453. Kaunas. Unpublished report
Bertašius, M. 2006b. Marvelės kapinynas. Archeologiniai tyrinėjimai Lietuvoje 2005 metais, 109-110
Bertašius, M. 2007a. Marvelės kapinynas. 2006 m. tyrinėjimų ataskaita. Lietuvos istorijos instituto rankraštynas, f. 1, b. 4741. Kaunas. Unpublished report
Bertašius, M. 2007b. Marvelės kapinynas. Archeologiniai tyrinėjimai Lietuvoje 2006 metais, 141-144
Bertašius, M. 2007c. Uwagi na temat kontaktów ludności kultury bogaczewskiej i mieszkanców środkowej Litwy na podstawie materiałów z cmentarzyska w Marvelė. In: A. Bitner-Wróblewska (Ed.). Kultura bogaczewska w 20 lat później. Materiały z konferencji. Warszawa: PMA, 251-260
Bertašius, M. 2008a. Marvelės kapinynas. 2007 m. tyrinėjimų ataskaita. Lietuvos istorijos instituto rankraštynas, f. 1, b. 4922. Kaunas. Unpublished report
Bertašius, M. 2008b. Marvelės kapinynas. Archeologiniai tyrinėjimai Lietuvoje 2007 metais, 175-178

Bertašius, M. 2009a. Marvelė. Ein Bestattungsplatz mit Mittellitauischer Pferdegräber. II Band. Marvelės žirgų kapinynas. Kaunas: Kauno technologijos universitetas Humanitarinių mokslų fakultetas

Bertašius, M. 2009b. Goth's trace – glass bead from Marvelė. In: A. Bitner-Wróblewska, G. Iwanowska (Eds.). Baltowie i ich sąsiedzi, Marian Kaczyński in memoriam. Warszawa: PMA, 239-244

Bertašius, M. 2009c. Marvelės kapinynas, 2008 metų III etapo tyrimų studija, ataskaita. Mokslo studija. Mašinraštis. Kaunas: Kauno technologijos universiteto Filosofijos ir kultūrologijos katedros archyvas

Bertašius, M. 2012. Marvelės kapinyno 2011 m. tyrinėjimų ataskaita. Lietuvos istorijos instituto rankraštynas, b. 5692. Kaunas. Unpublished report

Bertašius, M. 2018. Priešistorinė Marvelės bendruomenė: kultūrinės ir socialinės raidos aspektai. Kaunas: Technologija.

Bertašius, M., Daugnora, L. 1997. Kauno apylinkių žirgai. Veterinarija ir zootechnika, 4(26), 7-15

Bertašius, M., Daugnora, L. 2001. Viking Age horse graves from Kaunas region (Middle Lithuania). International Journal of Osteoarchaeology, 11, 387-399

Bertašius, M., Navasaitis, J., Selskienė, A. & Žaladarys, G. 2010. Marvelės kapinyno geležies dirbinių metalografiniai, mechaninių savybių ir elementinės sudėties tyrimai. Lietuvos archeologija, 36, 153-182

Daugnora, L. 1994. Marvelės kapinyno žirgai. In: A. Astrauskas (Ed.). Vidurio Lietuvos archeologija: konferencijos medžiaga. Vilnius: Žalioji Lietuva, 86-90

Jankauskas, R. 2005. Skeletal Inventory, Age, Sex and Pathology of Marvelė sample. In: M. Bertašius. Marvelė. Ein Gräberfeld Mittellitauens. I band. Kaunas: Kauno technologijos universitetas Humanitarinių mokslų fakultetas, 95-102

Jankauskas, R. 2009. Skeletal Inventory, Age, Sex and Pathologies of Marvelė Sample. In: M. Bertašius. Marvelė. Ein Bestattungsplatz mit Mittellitauischer Pferdegräber. II Band. Marvelės žirgų kapinynas. Kaunas: Kauno technologijos universitetas Humanitarinių mokslų fakultetas, 93-98

Jankauskas, R., Barkus, A. 1994. Marvelės senkapio (I-VII m.e.a.) antropologija (1991-1992 m. medžiaga). In: A. Astrauskas (Ed.). Vidurio Lietuvos archeologija: konferencijos medžiaga. Vilnius: Žalioji Lietuva, 79-85

Jankauskas, R., Čepliauskaitė, I. 2010. Senojo geležies amžiaus bendruomenės socialinės struktūros klausimu (Marvelės kapinyno duomenimis). In: G. Blažienė, S. Grigaravičiūtė, A. Ragauskas (Eds.). Florilegium Lithuanum: in honorem eximii professoris atque academici Lithuani domini Eugenii Jovaiša anniversarii sexagesimi causa dicatum. Vilnius: VPU leidykla, 92-103

### Doclea Bjelovine, Montenegro

**Collaborator**: Dusan Boric, Milos Zivanovic
**Coordinates**: 42.466869, 19.266164
**Individuals** (n = 6): R3478, R3481, R3482, R9918, R9919, R9920

Doclea necropolises

Doclea (http://www.antickadukljacg.com/en) is an ancient city situated in the hinterland of the eastern Adriatic coast. Until the 4th century it was part of the Roman province of Dalmatia and from the end of the 4th century it belonged to the province of Prevaletania (Sticotti 1913). The city was built at the beginning of the 1st century AD, in the center of the previous Illyrian territories that belonged to the Docleati tribes. In Late Antiquity, the city flourished the most. This situation can probably be linked to the sentiment of the Roman emperor Diocletian, who was born in the vicinity of the city. Relatively rapidly, already in the 7th century AD, the city was abandoned and later, during medieval times, never revived. The reason for this lies in the fact that it was not located on the main road of Via Egnatia, connecting Rome and Constantinople.

The remains of the dead from the city were interred in the southeastern necropolis that extended to Zagorič (Cermanović et al. 1975), the northeastern part that was next to an ancillary road (Medenica 2011, Gončarova 2012) (these are Samples #1-3), and at the western necropolises of Lovišta and Bjelovine. At the latter necropolis, of particular importance is the burial of the aristocratic Christian social stratum where an exceptional artefact known as the Podgorica Glass also originates (Živanović 2015; http://balkans.aljazeera.net/vijesti/podgoricka-casa-preko-sarajeva-zavrsila-u-ermitazu). Exceptionally, in Doclea, there is also evidence of human remains within the city (Baković 2011), with only several burials, which testify of turbulent periods of the 4th and 5th centuries.

Burials 17 and 25 from Bjelovina have been studied anthropologically. These burials should be younger than the 4$^{th}$ century AD, i.e. could belong to the 6th-7th centuries, which is significant since this period is poorly represented among the finds at Doclea.

Burial 1/2010 is found within the city limits. The individual was buried in a crouched position with a peculiar knife, which may suggest that this individual was a newcomer from another region. This burial in the city is an exception to the general rule. A preliminary anthropological analysis suggested this was a female.

| sample_id | sex (genetic) | date lower (95% CI) | date upper (95% CI) | date inference | burial_id |
|---|---|---|---|---|---|
| R3482 | F | 1025.5 calCE | 1152.5 calCE | 14C | smp6; Doclea-Bjelovine; Burial 17 |
| R3478 | F | 709.5 calCE | 880.5 calCE | 14C | Smp7; Doclea-Bjelovine; Burial 25 |
| R3481 | M | 211 calCE | 320.5 calCE | 14C | smp4; Doclea 2010; Burial 1 |
| R9918 | M | 996.5 calCE | 1150.5 calCE | 14C | Doclea, Bjelovine, B. 9 |
| R9919 | F | 900.5 calCE | 1024.5 calCE | 14C | Doclea, Bjelovine, B. 52 |
| R9920 | F | 1041 calCE | 1162.5 calCE | 14C | Doclea, Bjelovine, B. 55 |

**Literature**:

Baković, M., 2011. Preliminarni razultati istraživanja na prostoru kapitolnog hrama lokaliteta Doclea, *Nova antička Duklja* II, 9–26.

Gončarova, N., 2013. Анализ антропологического материала из гробницы Грудице около Дукли, *Nova antička Duklja* III, 95–104.

Sticotti, P. 1913. *Die Romische Stadt Doclea in Montenegr*o, Wien.

Cermanović-Kuzmanović, A., Velimirović-Žižić, O., Srejović, D. 1975. *Antička Duklja, nekropole,* Cetinje.

Medenica, I. 2011. Rimska grobnica na lokalitetu Grudice kod Duklje, *Nova antička Duklja* II, 113–136.

Živanović, M., 2015. Preispitivanje čuvene podgoričke čaše, *Nova antička Duklja* V, Podgorica, 77–108.

#### Weklice, Poland

**Collaborators**: Paulina Borówka, Iwona Teul, Magdalena Natuniewicz-Sekuła, Wiesław Lorkiewicz
**Coordinates**: 54.1176, 19.5742
**Individuals** (n = 9): R10618, R10620, R10625, R10626, R10631, R10633, R10634, R10636, R11391

The cemetery at Weklice is one of the most important and best studied archaeological sites of Wielbark Culture, which is connected to the Goths and Gepids, on territory of today's Poland. The cemetery was used from the end of the 1st century to the beginning of the 4th century A.D. The site is situated on Elbląg Heights, 12 km east of Elbląg (Warmian-Masurian Voivodeship), in northern Poland (54˚06´55˝N and 19˚34´23˝E). In the Roman Period it was the center of an amber route to the coast of the Baltic Sea (Vistula estuary), which predisposed it to being a strategic trading and cultural exchange center, linking the main trade routes in *European Barbaricum*. Reconstruction of the hydrological view of the Vistula delta during the Roman Period indicates that the Weklice site was an early port of trade settlement type, facilitating shipping connections in the Baltic Sea. Until now over 600 graves and other features have been discovered. Finds from the necropolis are currently stored at the Institute of Archaeology and Ethnology Polish Academy of Sciences in Warsaw, Poland. The graveyard is bi-ritual, both cremation (approx. 220) and inhumation (approx. 380). Several are significantly distinguished by a wealth of equipment and they are associated with the local elite of the Barbarian population.
Burial inventories up to now constitute about 4600 artifacts made of gold, silver, copper alloys, glass, amber, and clay. These are primarily ornaments and costume elements, both local manufacturing and imported from other areas of Barbaricum. There are also numerous Roman imports (bronze and glass vessels, beads and jewelry) testifying to transregional contacts of the population living here. An unusual wealth and comprehensiveness of discovered grave goods testifies to the broad connections of this population with the Roman Empire, Scandinavia and Black Sea territories.
The multifaceted analysis of materials from Weklice allowed for a significant increase in anthropological and archeological knowledge about settlement of Wielbark culture on Polish lands during the Roman Period.

| sample_id | sex (genetic) | date lower (95% CI) | date upper (95% CI) | date inference | burial_id |
|---|---|---|---|---|---|
| R10618 | F | 88.5 calCE | 212 calCE | 14C | Weklice_WEK_84 |
| R10620 | M | 41.5 calBCE | 64.5 calCE | 14C | Weklice_WEK_299 |
| R10625 | F | 41.5 calBCE | 212 calCE | SITE | Weklice_WEK_303 |
| R10626 | M | 41.5 calBCE | 212 calCE | SITE | Weklice_WEK_22 |
| R10631 | F | 41.5 calBCE | 212 calCE | SITE | Weklice_WEK_334 |
| R10633 | F | 41.5 calBCE | 212 calCE | SITE | Weklice_WEK_265 |
| R10634 | F | 41.5 calBCE | 212 calCE | SITE | Weklice_WEK_99/03/nr1 |
| R10636 | M | 41.5 calBCE | 212 calCE | SITE | Weklice_WEK_527 |
| R11391 | M | 41.5 calBCE | 212 calCE | SITE | WEK_50 |

**Literature**:


Natuniewicz-Sekuła M. 2008. Wheel-made vessels from Wielbark culture cemetery at Weklice nearby Elbląg. The contribution to research on pottery workshop of wielbark culture. IN: A. Błażejewski (ed.) *Ceramika warsztatowa w środkowoeuropejskim Barbaricum.* Instytut Archeologii Uniwersytetu Wrocławskiego, Wrocław, 47–66.

Natuniewicz-Sekuła M. 2017. The Craft of the Goldsmith in Wielbark Culture in the Light of the Finds from the Cemetery at Weklice, Elbląg Commune and Other Necropolis of Roman Period from Elbląg Heights. Technological Studies of Selected Aspects. *Sprawozdania Archeologiczne* 69, 185–233.

Natuniewicz-Sekuła M., Okulicz-Kozaryn J. 2006. Weklice. IN: *Reallexikon der Germanischen Altertumskunde* 33, 440–444.

Natuniewicz-Sekuła M., Okulicz-Kozaryn J. 2008. Two richly furnished graves with Roman imports from the cemetery at Weklice, site 7, Elbląg commune (Poland). *Germania* 86, 227–269.

Natuniewicz-Sekuła M., Okulicz-Kozaryn J. 2011. *Weklice. A Cemetery of Wielbark Culture on the Eastern Margin of Vistula Delta (excavations 1984-2004).* Monumenta Archaeologica Barbarica 17. Fundacja Monumenta Archaeologica Barbarica – Instytut Archeologii i Etnologii Polskiej Akademii Nauk – Państwowe Muzeum Archeologiczne w Warszawie, Warszawa.

Natuniewicz-Sekuła M., Rein Seehusen Ch. 2010. Baltic connections. Some remarks about studies of boat-graves from Roman Iron Age. Finds from the Slusegård and Weklice cemeteries. IN:  U. Lund Hansen, A. Bitner-Wróblewska (eds.), *Worlds Apart? Contacts across the Baltic Sea in the Iron Age. Network Denmark-Poland, 2005–2008.* Nordiske Fortidsminder C/7. Det Kongelige Nordiske Oldskriftselskab – Państwowe Muzeum Archeologiczne w Warszawie, København-Warszawa, 287–313.

Natuniewicz-Sekuła M., Skóra K. 2016. Selected children's burials from the Wielbark culture cemetery at Weklice, site 7, Elbląg comunne, warmińsko-mazurskie voivodeship *Archaeologia Polona* 51-52, 43–79.

Natuniewicz-Sekuła M. Werra D. H. 2016. Flint artefacts from the Wielbark culture cemetery at Weklice, Site 7, Elbląg county. *Journal of Lithic Studies* 3/2, http://journals.ed.ac.uk/lithicstudies/

#### Conímbriga, Portugal

**Collaborator**: Ana Maria Silva
**Coordinates**: 40.099444, -8.490556
**Individuals** (n = 2): R10487, R10488

The archaeological site of Conímbriga (Condeixa-à-Velha, Coimbra, Portugal) was excavated over several campaigns beginning at the end of the XIX century. These excavations confirmed human occupations from the Iron age until the Medieval period. The archaeological intervention took place in several areas, some of them revealing human burials. The eight graves recovered from area B are assigned to the Roman period. Although no direct radiocarbon dating on the human bones is available, coins from the IV$^{th}$ and V$^{th}$ centuries were recovered in one of these tombs. The roman occupation of Conímbriga is possibly linked to the Pacification campaign of Lusitania by the troops of Decímo Júnio Bruto during the second half of the 2$^{nd}$ century BC (Gameiro, 1998).
Petrous bones of two individuals (skeletons 6 and 8) were sampled. The former belongs to an adult male, aged over 30 years old at death. Remodeled infection lesions were observed in the distal portion of the right tibia. Skeleton 8 belongs to an adult female, also aged over 30 years old at death. Several pathologies were observed, including *antemortem* loss of 4 teeth, degenerative lesions in the column and right clavicle, entheseal changes on the right patella and both Achilles tendons, and a possible fracture of the left ulna (Gameiro, 1998).

| sample_id | sex (genetic) | date lower (95% CI) | date upper (95% CI) | date inference | burial_id |
|---|---|---|---|---|---|
| R10487 | F | 261.5 calCE | 413 calCE | 14C | Conímbriga, Sep. 6 |
| R10488 | M | 261.5 calCE | 413 calCE | SITE | Conímbriga, Sep. 2 |

**Literature**:
Gameiro, AL. (1998). Troia Romana: Paleobiologia de uma população romana da Necrópole de Tróia. Dissertação de Mestrado em Evolução Humana do Departamento de Antropologia, Faculdade de Ciências e Tecnologia da Universidade de Coimbra. Coimbra, Portugal.Unpublished Master thesis.

#### Miroiço, Portugal

**Collaborator**: Ana Maria Silva
**Coordinates**: 38.7261, -9.3653
**Individuals** (n = 8): R10499, R10500, R10501, R10502, R10503, R10506, R10507, R10508

The Late Roman cemetery of Miroiço (Cascais, Portugal), although not directly dated, is suggestive of being used during the 2nd–4th centuries AD. Field intervention took place between the years of 1999 and 2001, and 15 individual graves, 12 reused graves, 3 ossuary graves and two undetermined graves (due to the absence of human remains) were identified. The burials are primarily oriented East and North-South, delimited by lateral pillars and covered by limestone slabs. In addition to 32 inhumations, one incineration was recovered (Cardoso, 2018).

The anthropological analysis of the inhumations confirmed the presence of at least 77 individuals: 49 adults (both sexes) and 28 non adults. Low frequencies of oral, traumatic and degenerative pathologies, as well as, stress indicators (linear enamel hypoplasia) were registered (Silva, 2003; Silva and Silva, 2010).
Ten petrous bones were used for DNA analysis, including individuals of both sexes.

| sample_id | sex (genetic) | date lower (95% CI) | date upper (95% CI) | date inference | burial_id | excluded | notes |
|---|---|---|---|---|---|---|---|
| R10499 | M | 257.5 calCE | 404.5 calCE | 14C | Miroiço, Sep. 5 | | |
| R10500 | F | 257.5 calCE | 593.5 calCE | SITE | Miroiço, Sep. 8 | | |
| R10501 | M | 257.5 calCE | 593.5 calCE | SITE | Moiroiço, Sep. 11 | Yes (Related) | REL_2ND of R10508 |
| R10502 | F | 257.5 calCE | 593.5 calCE | SITE | Miroiço, Sep. 14 | | |
| R10503 | M | 441.5 calCE | 593.5 calCE | 14C | Miroiço, Sep. 17 | | |
| R10506 | F | 257.5 calCE | 593.5 calCE | SITE | Miroiço, Sep. 20 | | |
| R10507 | F | 257.5 calCE | 593.5 calCE | SITE | Miroiço, Sep. 21 | | |
| R10508 | F | 261.5 calCE | 413 calCE | 14C | Miroiço, Sep. 23 | | REL_2ND of R10501 |

<u>**Literature**</u>:

Cardoso, G. (2018). As Necrópoles romanas/visigóticas de Miroiço e Alcoitão (Cascais). *Conimbriga 57*: 169-216.

Silva, AL. (2003). *Registo de um quotidiano na Villa romana de Miroiço. Análise paleobiológica de uma amostra de esqueletos exumados da Necrópole de Miroiço.* Tese de Licenciatura em Antropologia. Departamento de Antropologia, Faculdade de Ciências e Tecnologia da Universidade de Coimbra. Unpublished Graduation thesis.

Silva, AM.; Silva AL. (2010). Unilateral non-osseous calcaneonavicular coalition: report of a Portuguese archaeological case. *Anthropological Science* 118(1): 61 - 64. (doi:10.1537/ase.090429).

#### Monte da Nora, Portugal

**Collaborator**: Ana Maria Silva
**Coordinates**: 38.8803, -7.1637
**Individuals** (n = 3): R10491, R10494, R10496

The site of Monte da Nora was discovered during construction work of the A6 highway (the Borba /Elvas sub-stretch, Portugal). This archaeological site revealed several occupations, dating from the Iron Age II to the late Roman period (V$^{th}$-VI$^{th}$ centuries). This place was initially an Iron Age fortified settlement and later occupied between the 1$^{st}$ century BC and the 6$^{th}$ century BC successively by Roman housing, production and storage structures. The results of the archaeological intervention points to a possible Roman *Vicus*. The site seems to have been abandoned in the 5$^{th}$ to 6$^{th}$ centuries and later occupied by a Necropolis.
In December of 1998 and February of 1999, a team from the Department of Anthropology of the University of Coimbra (currently, the Department of Life Sciences) excavated 20 graves. These are grouped into three nuclei: nucleus 1 (graves A and B), nucleus 2 (grave F) and nucleus 3 (remaining graves). The graves were dug into the rock (cystic nature), and in most cases covered by slate slabs. The burials are individual and there are no signs of reuse, although traces of old perturbations were observed especially for nuclei 2 and 3.
The individuals were deposited in supine position, oriented W - E, with variability in the position of the upper limbs. The sample consists of 19 adults and 1 non-adult (10/11 years). Adults include 7 females, 6 males and 6 individuals of unknown sex. Of the adults, according to the available age indicators, at least 7 were aged more than 30 years old at death.
Only three of the ten analysed petrous bone preserved DNA: graves G, J and O, all belonging to nucleus 3. The skeleton of grave G belongs to a female of more than 45 years old at death. The estimated stature is 152 ± 5.92 cm. Degenerative changes were observed in the articulations of the shoulder, elbow, knee and vertebral column.
The female unearthed from Grave J was more than 40 years old at death. 156 ± 5.92 cm was the estimated height. The preserved teeth display moderate dental wear, small deposits of calculus and a large cariogenic lesion was observed on the first upper molar. Articular degenerative lesions were only observed in the left mandibular condyle and in the corresponding temporomandibular articulation. Modest entheseal changes were observed in the thoracic vertebrae. Two enamel hypoplasia, a non-specific stress indicator, were observed in the left lower canine.
Grave J belongs to a female aged between 30 - 40 years at death. This skeleton stands out in terms of their extensive *antemortem* tooth loss: 7 upper (n=14) and 7 lower teeth (n=14). The upper left first premolar displays a moderate cariogenic lesion. In the postcranial skeleton, evidence of a possible dislocation was observed in the left elbow.

Two small “button” tumours (osteomas) were detected in the right parietal and temporal bones.

| sample_id | sex (genetic) | date lower (95% CI) | date upper (95% CI) | date inference | burial_id |
|---|---|---|---|---|---|
| R10491 | F | 782.5 calCE | 881.5 calCE | 14C | Monte da Nora, MN. Sep. G |
| R10494 | F | 682.5 calCE | 878.5 calCE | 14C | Monte da Nora, MN. Sep. J |
| R10496 | F | 774.5 calCE | 884 calCE | 14C | Monte da Nora, MN. Sep. O |

**Literature**:

Silva, AM. (1999). *Relatório antropológico dos restos ósseos humanos exumados do Monte da Nora (Elvas).* Coimbra, Departamento de Antropologia da Universidade de Coimbra. Technical-Scientific Report.

#### Sant’ Imbenia, Sardinia

**Collaborators**: Gabriella Gasperetti, Michaela Lucci, Alfredo Coppa, Francesca Candilio
**Coordinates**: 40.5938, 8.2045
**Individuals** (n = 7): R11828, R11829, R11832, R11833, R11834, R11835, R11836

The archaeological site of Sant’Imbenia lies in Porto Conte bay, a natural harbour called *Portus Nympharum* by the geographer Ptolemy in the second century AD, 10 km to the WNW of Alghero (SS). During the fourteenth century BCE a single-tower nuraghe was built with some huts around within the current coastline, subsequently surrounded by a bastion (Bafico *et al.* 1985; Bafico 1998). The nuraghe was in use until the tenth century BCE and the village evolved towards an "urban" organization at the end of the ninth century BCE, around an open space interpreted as a market square. This development is related to seafaring travels around the Western Mediterranean Sea by Phoenicians, Etruscans and Greeks. The site remained a very active center at least until the seventh century BCE, an era of transformations in the indigenous world, which saw the establishment of the Phoenician colonies in SW of Sardinia (Garau, Rendeli 2012; Rendeli 2018; Clemenza *et al.*).

About 900 m SW of the nuragic village, between the first and third centuries AD was built and inhabited the seaside villa of Sant'Imbenia, a large private dwelling dedicated to holidaying and leisure, covering a total area of over 3500 m2, decorated with precious elements, mosaics and refined stucco work on the walls and ceiling, that testify to an opulent lifestyle, found only in Lazio, in Rome, and in the Campi Flegrei area in Campania. Archaeological stratigraphies show that it was still inhabited at least until the 7th century AD, no longer as a luxurious private residence, but sporadically by small social groups, simple pastoral and farming communities (Manconi 1999; Rovina 2015).

An area used as a necropolis, dating back to I-III century AD, was identified and partially excavated in the 1960s near the road connecting the hamlet of Santa Maria la Palma to Porto Conte. Some burials reused limestone steles with schematically engraved human faces dated to Punic-roman age (Maetzke, 1962). Later, in the 1990s, an emergency archaeological excavation due to the installation of a pipeline was directed by the Superintendence of Archaeological Heritage for the Provinces of Sassari and Nuoro, with the excavation of other burials, mainly human graves, often within amphorae, and sometimes used for multiple burials.

The samples being analyzed come from a stratigraphy damaged by previous land drainage works, plowing and stone removal carried out during the twentieth century. For these reasons, for example, tombs 1, 2, 3, 4 were upset. The necropolis, extremely poor in dating materials, has been referred to the Roman imperial age; however we cannot exclude the presence of remains from a previous period, possibly in a secondary position, due to the conditions of the excavation.

| sample_id | sex (genetic) | date lower (95% CI) | date upper (95% CI) | date inference | burial_id | excluded | notes |
|---|---|---|---|---|---|---|---|
| R11828 | M | 1115 calBCE | 984.5 calBCE | 14C | S. Imbenia_7489/T1 | | |
| R11829 | F | 1004.5 calBCE | 909.5 calBCE | 14C | S. Imbenia_7489/T2 | | |
| R11832 | F | 242 calCE | 346 calCE | 14C | S. Imbenia_7490 B | | |
| R11833 | F | 248.5 calCE | 378 calCE | 14C | S. Imbenia_7491/S2 | | |
| R11834 | F | 129.5 calCE | 234.5 calCE | 14C | S. Imbenia_7492 A | | REL_2ND of R11836 |
| R11835 | M | 800.5 calBCE | 774.5 calBCE | 14C | S. Imbenia_7494/T3 | | |
| R11836 | F | 130 calBCE | 378 calCE | SITE | S. Imbenia_7497F16 | Yes (Related) | REL_2ND of R11834 |


**Literature**:

Bafico, Susanna. 1998. *Nuraghe e villaggio Sant'Imbenia, Alghero*.Viterbo: Betagamma editor.

Bafico, Susanna, D'Oriano, Rubens, Lo Schiavo, Fulvia. 1985. Il villaggio nuragico di Sant'Imbenia (SS). Nota preliminare. *Actes du IIIe Congres International des Etudes pheniciennes et puniques*, Tunis: 87–94.

Clemenza, Massimiliano *et alii* 2021. Sant'Imbenia (Alghero): further archaeometric evidence for an Iron Age market square, *Archaeological and Anthropological Sciences* 13: 181. https://doi.org/10.1007/s12520-021-01425-x

Garau, Eliabetta, Rendeli, Marco. 2012. From huts to houses. Urbanistica a Sant'Imbenia", *Atti della XLIV Riunione Scientifica dell'Istituto Italiano di Preistoria e Protostoria, La preistoria e la protostoria della Sardegna, Cagliari-Barumini-Sassari 23-28 novembre 2009*, 2-3. Firenze: 893-898.

Maetzke, Guglielmo. 1962. Porto Conte (Alghero), Scavi e scoperte nelle province di Sassari e Nuoro. *Studi Sardi XVII*, 1959-61: 656-689.

Manconi, Francesca. 1999. *Villa romana di Sant'Imbenia a Porto Conte, Alghero.* Viterbo: Betagamma editor.

Rendeli, Marco. 2018. Sant'Imbenia and the Topic of the Emporia in Sardinia, Gailledrat E., Dietler M., Plana-Mallart R. (edd) *The Emporion in the Ancient Western*

*Mediterranean. Trade and Colonial Encounters from the Archaic to the Hellenistic Period*. Montpellier: Presses Universitaires de la Mediterranee: 191–204.

Rovina, Daniela. 2015. La villa romana di Sant'Imbenia, in Costanzi Cobau, A., Nardi, R. *Una villa in una stanza. La villa romana di Sant'Imbenia. documentazione, conservazione e musealizzazione dei reperti rinvenuti nel corso degli scavi 1994-2005*. Tivoli: CCA Editore: 13-23.

#### Beška-Brest, Serbia

**Collaborators**: Dusan Boric, Tijana Pesterac
**Coordinates**: 45.129863, 20.069845
**Individuals** (n = 2): R6681, R6688

The archaeological site of Brest is situated in the village of Beška. The necropolis was found on the slopes of the Fruška Gora mountain, 5 km from the bank of the Danube River. According to the Roman map of the border zone and the *Limes* area between Cusum (present-day Petrovaradin) and Acumincum (present-day Stari Slankamen), the necropolis is located in the very vicinity of the Roman road by the Danube. The remains of the nearest *castrum* can be seen in the woods of Čortanovci, 11 km away. During the excavations of the necropolis, field surveys have also been conducted in order to obtain information about a possible settlement in which the people buried here might have lived. Excavated in its entirety, this graveyard shows valuable and well preserved inventories, especially notable for the famous tomb with frescoes. Biritual burial customs have been revealed here: from 111 graves in total, 22 are cremation finds, 81 inhumations and 8 cenotaphs. At the eastern and western end of the necropolis, ovens and graves as cult pits have also been found. The excavations of this important site were conducted by the Museum of Vojvodina between 1967 and 1974. The results were published in 1987 by Mirjana Marijanski-Manojlović under the title *Rimska nekropola kod Beške u Sremu*, with detailed description of the graves, burial rites, and artefacts. The results of the anthropological analysis are also published and show the gender determination of the skeletal remains. The cemetery might have been used over a long period of time with the datable grave goods falling into the span from the dynasty of Severi (AD 222–235) to Valentian and Valens (AD 364–378). The material is stored in the Museum of Vojvodina in Novi Sad.

**Burial 56** contains the remains of an adult individual placed in extended supine position and oriented NE-SW. A bronze hairpin was found on the right side of the forehead (Marijanski-Manojlović 1987: 53, T. 35).

**Burial 74** contains the remains of an adult individual oriented W-E. Next to the right leg there was a ceramic jug (Marijanski-Manojlović 1987: 57, T. 41).

| sample_id | sex (genetic) | date lower (95% CI) | date upper (95% CI) | date inference | burial_id |
|---|---|---|---|---|---|
| R6688 | M | 257.5 calCE | 404.5 calCE | SITE | Beška-Brest; Burial 56 |
| R6681 | F | 257.5 calCE | 404.5 calCE | 14C | Beška-Brest; Burial 74 |

**Literature**:

Marijanski-Manojlović, M., 1987. Rimska nekropola kod Beške u Sremu. Vojvođanski muzej, Novi Sad.

### Naissus, Serbia

**Collaborators**: Nataša Miladinović-Radmilović (anthropologist), Dragana Vulović (anthropologist) and Gordana Jeremić (archaeologist)
**Coordinates**: 43.326684, 21.896187
**Individuals** (n = 2): R6764, R6769

Late antique city necropolis of Naissus (Niš, SRB)

The largest city necropolis of Naissus (Niš) in the Late Antiquity period was formed on the loess plateau on the right bank of the Nišava, east of the fortress, along the road to *Ratiaria*. Since it is located in today's city quarter of Jagodin Mala, it became known by that name in the literature. In Late Antiquity, this area had been used intensively for burials, from the mid of the 4$^{th}$ century to the 7$^{th}$ century.
Systematic investigation of the necropolis in Jagodin Mala has been conducted intermittently, for no less than eight decades. Its existence was discovered at the end of the 19$^{th}$ century. The assumed length of the necropolis in an east-west direction was about 2 kilometres, while the width in a north-south direction was about 600 metres. During the systematic exploration of this area, five church edifices, a crypt, over 65 brick vaulted tombs, a large number of brick lined graves and around 400 graves of the deceased, interred in the ground, have been identified. Between the tombs and in the free spaces, the deceased were buried in simple grave holes or in wooden boxes. The area has revealed different types of sepulchral structures, building methods, and materials that were used, as well as the funeral practices of the inhabitants of *Naissus* in Late Antiquity, during Early Christianity. The wealth of grave goods and inventory suggests that the city population lived in a time of social and economic prosperity. According to the results of the archaeological excavations, the central part of the necropolis was located between Čegarska Street and the Benetton factory, where densely lined and regular rows of graves were discovered, from the mid of the 4$^{th}$ century to the mid of the 5$^{th}$ century. The proposed grave units for analyzing belong to the period of the mid – second half of the 4$^{th}$ century (graves 2, 5, 8, 11, 12, 13, 14 and 63 from the excavations in 2012), encompassing the part of the middle class of the citizens of the late Antique Naissus.

| sample_id | sex (genetic) | date lower (95% CI) | date upper (95% CI) | date inference | burial_id |
|---|---|---|---|---|---|
| R6764 | M | 255.5 calCE | 405.0 calCE | 14C | Naissus, Jagodin Mala-Benetton; G-2 (2) |
| R6769 | F | 26.0 calCE | 126.0 calCE | 14C | Naissus, Jagodin Mala-Benetton; G-8 (3) |

**Literature**:

*Late antique necropolis Jagodin mala*, Introductory text G. Jeremić, authors of catalogue entries S. Drča, G. Jeremić, V. Crnoglavac, National Museum in Niš, Niš 2014.

G. Jeremić, Burials in *Naissus* in Late Antiquity – case study of the necropolis in Jagodin Mala, in: *Constantine the Great and the Edict of Milan 313. The birth of christianity in the Roman provinces on the soil of Serbia*, ed. I. Popović, B. Borić-Brešković, Archaeological Monographs 22, National Museum in Belgrade, Belgrade 2013, 126-135.

G. Jeremić, T. Čerškov, Requescit in pace – neka zapažanja o poremećajima "večnog počinka" na primeru jugoistočnog dela nekropole u Jagodin Mali (Naissus) (summary: Requescit in pace – some remarks on disturbances of "etrnal rest" on the example of the south-eastern part of the necropolis of Jagodin Mala (Naissus) ), u/in: *Bioarheologija na Balkanu. Metodološke, komparativne i rekonstruktivne studije života u prošlosti / Bioarchaeology in the Balkans. Methodological, comparative and reconstructive studies of lifes in past*, ur./ed. N. Miladinović-Radmilović, S. Vitezović, Beograd, Sremska Mitrovica 2016, 127–145.

Г. Јеремић, Шестар занатлије из касноантичке некрополе у Јагодин мали у Нишу (Naissus) (summary: An artisan`s compass – divider from the late antiquity necropolis in Jagodin mala, Nis (Naissus), *Зборник Народног музеја (Ниш)* / *Papers of the National Museum of Niš* 22, 2013, 29-46.

Т. Чершков, Г. Јеремић, А. Алексић, Заштитна археолошка истраживања на локалитету „Бенетон Србија“ у Јагодин мали у Нишу, *Гласник Друштва конзерватора Србије* 38, 2014, 53–60.

Т. Чершков, Г. Јеремић, Д. Вуловић, О једној занимљивој гробној целини из Јагодин мале, Ниш (*Naissus*) (About one interesting masonry grave in Jagodin Mala, Niš (Naissus)), *Зборник Народног музеја (Ниш)* / *Papers of the National Museum of Niš* 23, 2014, 35–64.

Г. Јеремић, Т. Чершков, Д. Вуловић, Гроб богате грађанке из касноантичког Наиса (Naissus) (summary: Grave of a wealthy female citizen from the Late Roman Naissus), *Гласник Српског археолошког друштва* / *Journal of Serbian Archaeological Society*, 30, Београд/Belgrade 2014, 83–108.

G. Jeremić, S. Golubović, S. Drča, Unpublished glass findings from the eastern necropolis of Naissus (Jagodin Mala, Niš), *Starinar* LXVII, Belgrade 2017, 109–130.

G. Jeremić, A. Filipović, Traces of Early Christianity in *Naissus*, in: *Acta XVI Congresvs internationalis archaeologiae christianae, Romae (22-29.9.2013), Constantino e i Constantinidi, l'innovazione costantiniana, le sue radici e i suoi sviluppi*, Pars II, eds. O. Brandt, V. Fiocchi Nicolai, Studi antichità cristiana, Pontificio istitituto di archeologia cristiana vol. LXVI, Città dell Vaticano 2016, 1743–1758.

#### Sirmium, Serbia

**Collaborators**: Nataša Miladinović-Radmilović (anthropologist), Dragana Vulović (anthropologist) and Stefan Pop-Lazić (archaeologist)
**Coordinates**: 44.966801, 19.609892
**Individuals** (n = 5): R3906, R3918, R6730, R6737, R9662

Basic description of sites 55, 76, 77 – Sirmium
Dr. Stefan Pop-Lazić

All three sites are positioned near the same modern street „Arsenija Čarnojevića" which is the historical main street to the east, and also part of the eastern necropolis.
Site 55 (N: 4980803 E: 391714) or *Irineus basilica* was excavated in two campaigns with more than 90 graves found inside and close to the early christian basilica. Based on the position of the inhumations Nr. 1, 5, 15 close to the apsys within built grave structures they could be of high rank within the early christian community. Their position within Roman society is indiscernible. The same applies for graves Nr. 12 and 38.
Graves found in the sites 76 (N: 4980686 E: 391109) and 77 (N: 4980698 E: 390820) belong to the same necropolis. Dating is problematic since the burials do not have any kind of grave goods buried with them. Based on the fact that all of the grave pits were buried in the second half of the IV century horizon, they should be dated to after the IV century. It is unlikely that they belong to the V-VI century period, based on the fact that there are no grave goods and built grave constructions. In a few graves we have found nails which point to the fact that bodies were buried in wooden coffins, which raises the possibility that these graves belong to the population of the XV-XVII century period. This could be the population living in the eastern suburbs of Sremska Mitrovica in that period. We have one necropolis of that period in the center of Sremska Mitrovica, so I think that this necropolis marks the area where another church should be looked for. Based on grave types and non-existent grave goods, one can not say anything about the social position of the deceased. Taking into account that they were buried in the town necropolis, they are from the urban population.

| sample_id | sex (genetic) | date lower (95% CI) | date upper (95% CI) | date inference | burial_id | excluded |
|---|---|---|---|---|---|---|
| R3906 | M | 1479 calCE | 1634 calCE | 14C | Sirmium, Site 77; Gr. 2 (I) | |
| R3918 | M | 993.5 calCE | 1149 calCE | 14C | Sirmium, Site No. 55; Gr. 5 (I) | |
| R6730 | M | 266 calCE | 430 calCE | 14C | SIRMIUM; SITE No. 55; GRAVE no.1 (I) | Yes (Contam.) |
| R6737 | F | 1481.5 calCE | 1635.0 calCE | 14C | SIRMIUM; SITE No. 76; GRAVE No. 12 | |
| R9662 | F | 1461 calCE | 1630.5 calCE | 14C | Sirmium, Site 77; Gr. 19 (I) | |

**Literature - Archaeology - sites 55, 76 and 77:**

Јесретић, М. 2007 Преглед археолошких истраживања Сирмијума од 1991. до 2006. године. *Зборник Музеја Срема* 7: 29–40.

Milošević, P. 1979 Три ранохришћанска гробна објекта у Срему. Рад војвођанских музеја 25: 115–120.

Milošević, P. 1994 *Топографија Сирмијума*. Археолошка грађа Србије I/3. Грађа за археолошку карту Војводине 1. Нови Сад: Српска академија наука и уметности, огранак у Новом Саду.

Mirković, M. 1971 Sirmium – its History from the I Century A. D. to 582 A. D. *Sirmium* I: 5–94.

Mirković, M. 2008 *Sirmium, istorija rimskog grada od I do kraja VI veka*. Sremska Mitrovica: Blago Sirmijuma, drugo izdanje.

Popović, V. 2003 *Град царева и мученика*. Сремска Митровица: Благо Сирмијума.

Прица, Р. 2001 Хришћански мученици у Сирмијуму. *Историјски архив „Срем“* 1946–2001: 170–173.

Василић, Б. 1952 Топографска истраживања Сирмијума. *Зборник Матице Српске – серија друштвених наука* 3: 165–168. Нови Сад.

**Literature - Anthropology - sites 55, 76, 77 & all osteological material (Sirmium):**

Miladinović, N. 2005. *Paleodemografska struktura i problematika srednjovekovne nekropole u Sremskoj Mitrovici*, Magistarska teza, Filozofski fakultet, Univerzitet u Beogradu.

Миладиновић, Н. 2006. Физичко-антрополошка анализа остеолошког материјала из германских гробова са локалитета 85 у Сремској Митровици. *ГСАД* 22: 409–434.

Миладиновић-Радмиловић, Н. 2008. Полна и старосна структура дечијих индивидуа сахрањених на средњовековној некрополи на локалитету 85 у Сремској Митровици. *ГСАД* 24: 445–456.
Миладиновић-Радмиловић, Н. 2008. Феномен трепанације на примеру лобање из Сремске Митровице. *Рад музеја Војводине* 50: 175–186.
Миладиновић-Радмиловић, Н. 2008. Болести зглобова као најчешћа обољења зглобова на скелетима са археолошких локалитета у Србији. *Саопштења* 40: 151–162.
Миладиновић-Радмиловић, Н. 2009. Лобања из саркофага пронађеног у североисточном делу Сремске Митровице. *Рад музеја Војводине* 51: 137–145.
Миладиновић-Радмиловић, Н. 2009. Аварски коњанички гроб пронађен 1968. на локалитету Стара циглана – антрополошка анализа. *Зборник Музеја Срема* 8: 122–136.
Miladinović-Radmilović, N. 2010. *Antropološka struktura stanovništva Sirmijuma/Sremske Mitrovice kroz istorijske periode*, Doktorska disertacija, Filozofski fakultet, Univerzitet u Beogradu.
Miladinović-Radmilović, N. 2010. Exostoses of the External Auditory canal. *Старинар* LX/2010: 137–146.
Miladinović-Radmilović, N. 2011. Sirmium – Necropolis. Beograd: Arheološki institut, Sremska Mitrovica: Blago Sirmijuma.
Brown M. and Ortner D. J. 2011. Childhood Scurvy in a Medieval Burial from Mačvanska Mitrovica, Serbia. *International Journal of Osteoarchaeology* 21: 197–207.
Miladinović-Radmilović, N. 2012. Analysis of human osteological material from the eastern part of site No. 37 in Sremska Mitrovica. *Старинар* LXII/2012: 181–204.
Miladinović-Radmilović, N. 2012. Artificial cranial deformation. *ГСАД* 28: 303–314.
Миладиновић-Радмиловић, Н. и Димовски, Н. 2012. Стафнеов дефект на налазима мандибула из римског периода и позног средњег века са два локалитета у Војводини. *Рад Музеја Војводине* 54: 97–103.
Миладиновић-Радмиловић, Н. 2012. Учесталост и дистрибуција *cribrae orbitaliae* у Сирмијуму. *Саопштења* XLIV: 229–236.
Miladinović-Radmilović, N. 2012. Antropološka istraživanja u 2011. i 2012. godini, u V. Bikić, S. Golubović, D. Antonović (ur.) *Arheologija u Srbiji: projekti Arheološkog instituta u 2011. godini*, Beograd, Arheološki institut: 86–89.
Miladinović-Radmilović, N. 2013. *Caring for the human skeleton remains in Sremska Mitrovica (Sirmium).* Strategie e Programmazione della Conservazione e Trasmissibilità del Patrimonio Culturale, a cura di A. Filipović, Williams Troiano. Edizioni Scientifiche Fidei Signa. Rome: Italian Heritage Awards 2013, 282–291.
Миладиновић-Радмиловић, Н. и Вуловић, Д. 2013. Хернијација интервертебралног диска – Шморлов дефект на скелетним остацима са археолошких налазишта. *ГСАД* 29: 369–386.

Миладиновић-Радмиловић, Н., и Вуловић, Д. 2015. *Osteochondritis dissecans* – учесталост и дистрибуција код становника античког Сирмијума, у: *Српско археолошко друштво, XXXVIII годишњи скуп, Пирот 4–6. јун 2015*, ур. Д. Антоновић, В. Филиповић, Пирот 2015, 82–83.
Вуловић, Д. и Миладиновић-Радмиловић, Н. 2015. Случајеви *patella bipartita* и *os acromiale* из Сремске Митровице, у: *Српско археолошко друштво, XXXVIII годишњи скуп, Пирот 4–6. јун 2015*, ур. Д. Антоновић, В. Филиповић, Пирот 2015, 84.
Miladinović-Radmilović, N., Vulović, D. i Đukić, K. 2016. Health status of children in the ancient Sirmium. *Starinar* LXVI/2016: 65–80.
Миладиновић-Радмиловић, Н., Бизјак, Д. и Вуловић, Д. 2016 Учесталост и дистрибуција *suturae metopicae* код становника античког Сирмијума, у: *Српско археолошко друштво, XXXIX Скупштина и годишњи скуп, Вршац 2– 4. јун 2016,* ур. А. Црнобрња, В. Филиповић, Вршац 2016, 74.
Miladinović-Radmilović, N. 2017 Matériel ostéologique humain des tombeaux du V$^{e}$ et V$^{e}$ siècle, in: Sirmium à l'époque des Grandes migrations (eds. I. Popović, M. Kazanski, V. Ivanišević). Leuven-Paris-Bristol, CT: Peeters (Centre de recherche d'histoire et civilisation de Byzance, monographies 53); Belgrade: Institut archéologique de Belgrade (monographies 60), 2017, Chapitre IV : 93–124.
Miladinović-Radmilović, N. 2017 *Preparation of final documentation and provision of permanent and safe storage of osteological material from earlier anthropological research in Sirmium*, in: *Association for Biological Anthropology and Osteoarchaeology Annual Review 2016* (ed. Ronika K. Power, Macquarie University). British Association for Biological Anthropology and Osteoarchaeology, 2017, Issue 18, 18–19.
Vulović, D. i Miladinović-Radmilović, N. 2017 Slučajevi *patellae bipartitae* iz Sremske Mitrovice (Sirmium), u: *Bioarheologija na Balkanu. Markeri okupacionog stresa i druge studije. Radovi Bioarheološke sekcije Srpskog arheološkog društva* (*Bioarchaeology in the Balkans. Markers of occupational stress and other studies. Papers of the Bioarchaeological section of The Serbian Archaeological Society*) (ur. N. Miladinović-Radmilović i K. Đukić). Srpsko arheološko društvo, Blago Sirmijuma, Beograd-Sremska Mitrovica 2017, 45–56.
Miladinović-Radmilović, N., Đukić, K. i Vulović, D. 2017 Formiranje antičke antropološke zbirke u Muzeju Srema u Sremskoj Mitrovici, u: *Bioarheologija na Balkanu. Markeri okupacionog stresa i druge studije. Radovi Bioarheološke sekcije Srpskog arheološkog društva* (*Bioarchaeology in the Balkans. Markers of occupational stress and other studies. Papers of the Bioarchaeological section of The Serbian Archaeological Society*) (ur. N. Miladinović-Radmilović i K. Đukić). Srpsko arheološko društvo, Blago Sirmijuma, Beograd-Sremska Mitrovica 2017, 177–202.
Миладиновић-Радмиловић, Н. 2017 Антички Сирмијум – трагови једне неуспеле политике, у: *Српско археолошко друштво, XL Скупштина и годишњи скуп и*

*прослава 70 година Археолошког института (Mnemosynon Firmitatis), Београд 5–7. јун 2017,* ур. А. Црнобрња, В. Филиповић, Београд 2017, 64–65.
Miladinović-Radmilović, N. 2018 Population of Ancient Sirmium, in: *Vivere militare est. From Populus to Emperors – Living on the Frontier* (eds. S. Golubović and N. Mrđić). Volume II. Belgrade: Institute of Archaeology, Monographies No. 68/2, 2018, 7–39.
Djukic, K., Miladinović-Radmilović, N., Draskovic, M. and Djuric, M. 2018 Morphological appearance of muscle attachment sites on lower limbs: horse riders versus agricultural population. *International Journal of Osteoarchaeology* 28 (6), 656–668.
Miladinović-Radmilović, N., Vulović, D. i Đukić, K. 2018 Sirmijum – Rezultati antropološkog projekta u 2016. godini, u: *Arheologija u Srbiji: projekti Arheološkog instituta u 2016. godini*, I. Bugarski, N. Gavrilović Vitas, V. Filipović (ur.) Beograd, Arheološki institut: 148–153.
Миладиновић-Радмиловић, Н. 2017. године Трагови Пробове неуспеле политике у античком Сирмијуму, Музеј Срема, Сремска Митровица, свечана сала, предавање поводом манифестације Музеји Србије од 10 до 10, 18. 05. 2017. године.
Миладиновић-Радмиловић, Н. 2017 Становништво античког Сирмијума, у: *Први симпозијум о археологији у Сирмијуму, Музеј Срема, Сремска Митровица, 23–29. новембра 2017,* Сремска Митровица 2017.

#### Sviloš-Kruševlje, Serbia

**Collaborators**: Dusan Boric, Tijana Pesterac
**Coordinates**: 45.912477, 19.174329
**Individuals** (n = 2): R6693, R6701

The Sviloš-Kruševlje necropolis

Another Late Roman necropolis, excavated by the experts of the Museum of Vojvodina, is situated in the village of Sviloš, on a small hill of Kruševlje. The village can be found on the present-day road between the Fruška Gora mountain and the Danube River. This area had a good geographic position as the first zone of defense between the Limes and the plains of the Syrmia district, connecting the Danube with Sirmium and Bassianae in the Late Roman period. Previously conducted surveys revealed the continuity of habitation from the early Roman settlement until the necropolis of the 3rd and 4th centuries AD, as well as medieval cemeteries dating to the 14th and 15th centuries. Systematic rescue excavations were conducted from 1976 to 1983 and covered a large area. It is possible that the necropolis covered up to 3,484 $m^2$. There were investigated graves with the deceased buried in burial pits (n=34) or in-built tombs (n=11). Most of the tombs are devastated and skeletons are dislocated. Referring to the finds of coins in the graves, it is possible to establish the chronological span from the middle of the 3rd to the end of the 4th centuries AD, the earliest being antoniniani of Galienus, Probus, Tacitus, Aurelianus and Claudius II and the latest being the coin of Constantius II minted in Siscia from AD 355 to 361. The osteological remains of 63 individuals are preserved, so anthropological analyses have also been conducted, aiding our knowledge about bioarchaeological characteristics of the population buried here. According to the osteological analyses made by Zs. Zoffmann, in the investigated part of the necropolis, there were buried 28 men, 18 women, and 16 children. Together with some contemporary necropolises, such as the one in Beška and others found in present-day Hungary, this place shows the unity of material and spiritual culture, although likely an ethnically heterogeneous population in the late 4th century. It shows the same religious, social, and economic conditions, which resulted in the uniformity of the cultural space.

**Burial 40** contains the remains of a 1.5-2-year-old child placed in an extended supine position with E-W orientation and the face turned to the East. Five Galienus coins were found next to the left side of the pelvis. There was also an iron knife next to the left knee, and a ceramic vessel next to the left foot (Dautova-Ruševljan 2003: 19–20, 50).

**Burial 45** contains the remains of a 9-10-year-old child placed in an extended supine position with E-W orientation and the face turned to the East. One Constantius II coin was found next to the right leg, two coins were found next to the left side of the skull. Beneath the skull was an iron buckle, and an iron knife was found next to the left leg. Next to the right foot there was a ceramic jug and a glass (Dautova-Ruševljan 2003: 22, 57).

| sample_id | sex (genetic) | date lower (95% CI) | date upper (95% CI) | date inference | burial_id |
|---|---|---|---|---|---|
| R6693 | M | 236.0 calCE | 331.5 calCE | SITE | Site: Sviloš-Kruševlje; 1980; Burial 40 |
| R6701 | M | 236.0 calCE | 331.5 calCE | 14C | Site: Sviloš-Kruševlje; 1980; Burial 45 |

**<u>Literature</u>**:

Dautova-Ruševljan, V., 2003. Kasnoantička nekropola u Sremu (Late Roman Necropolis near Sviloš in Srem). Muzej Vojvodine, Novi Sad.

Zoffmann, Zs. 2003. Antropološki pregled rimske nekropole sa lokaliteta Sviloš-Kruševlje, in V. Dautova-Ruševljan, Kasnoantička nekropola u Sremu: 175–191. Muzej Vojvodine, Novi Sad.

#### Viminacium, Serbia

**Collaborators**: Ilija Mikić (anthropologist), Miomir Korać (archaeologist), Snežana Golubović (archaeologist)
**Coordinates**: 44.7167, 21.1667
**Individuals** (n = 7): R3931, R6750, R6756, R6759, R9669, R9673, R9674

KOD KORABA

The site named “Kod Koraba” is located in the broader zone of Viminacium, approximately 650 m to the South-East from a military camp. The whole area of the locality was endangered by the expanding “Drmno” surface coal mine. Rescue investigations lasted from 2005 until 2008 and they included geophysical survey and archaeological excavation.
On the “Kod Koraba” site 210 graves were found. Both types of burials were detected – there were 132 cremations and 78 inhumations. According to the types of graves and the goods found in them, the necropolis dates back to the period between the second part of the first century A.D. and the second part of the third century, with the exception of a single fourth-century grave.

**G-36**, trench 40, freely buried, at depth 1,60 m
Orientation: N-S with dev. 8º to W
Inventory: bronze coin, iron shunails, ceramic pot, ceramic beaker

| sample_id | sex (genetic) | date lower (95% CI) | date upper (95% CI) | date inference | burial_id |
|---|---|---|---|---|---|
| R6759 | M | 88.5 calCE | 212.0 calCE | 14C | VIMINACIUM; KOD KORABA; G-36 |

**Literature**:
Bogdanović, I. 2009 Rezultati arheološko-geofizičkih istraživanja na lokalitetu “Kod Koraba” (istočna nekropola Viminacijuma) (Results of archaeological and geophysical research at “Kod Koraba” (eastern necropolis of Viminacium)). *Archaeology and Science* 5: 83–100.

PIRIVOJ

At this site east from the military fort the necropolis with 436 inhumations and 76 cremations from the 2nd to 4th century was investigated. Pirivoj is noteworthy because of a mausoleum complex measuring 20 x 20 m, which was investigated in the year 2003. The mausoleum is unique and thus of particular importance because it is surrounded by a square wall. The complex was constructed with stone blocks and ashlars decorated with columns. In its central part, a main building measuring 5 x 5 m was built from green schist, bonded by plaster, and contains a tomb. This form of burial, known as *bustum*, is generally very rare, and it was even exceptional during that time. The buried individual was cremated, and must have been a person of great distinction in the Roman hierarchy. Other graves surrounding this central burial had conspicuous inventories, among them twenty gold objects and a gilded fibula (Korać et al. 2009: 99). The skeletons and cremations were located outside the square wall surrounding the mausoleum. Among these, a grave with tile structures and fresco painting dating to the early 4th century was detected, which is now open to the public. According to the depictions, it was a pagan tomb. However, both pagans and Christians were buried at Pirivoj, since in another grave, a silver fingerring with a carved Christogram was found. Another conspicuous feature of the Pirivoj site is a trench at its very east with more than 100 graves containing both uncremated and cremated individuals. Here, the deceased had been buried within a rather short time span. At the deepest level, two stepped graves with a human inhumation and a horse skeleton were found.

**G-349** freely buried, at depth 2,05 m
Orientation: S-N with dev. from 380˚ S partly to W
Inventory: An iron fibula (type: Hinge-like fibulae similar to Aucisa fibula; II-middle III century)

**G-354A** freely buried, at depth 1,65 m
Orientation: S-N with dev. 32˚ S to W

**G-360** freely buried, at depth 2,1 m
Orientation: N-S with dev. 320˚ S to W
Inventory: Coin, shuenails

**G-385** funeral in a wooden coffin, at depth 3,2 m
Orientation: N-S with dev. 290˚ S to E
Inventory: Ceramic vessel

**G-386** freely buried, at depth 3,5 m
Orientation: E-W with dev. 330˚ E to N

**G-411** freely buried
Orientation: W-E with dev. 5 ° W to S
Inventory: Bronze coin, Iron shuenails, ceramic cup

| sample_id | sex (genetic) | date lower (95% CI) | date upper (95% CI) | date inference | burial_id |
|---|---|---|---|---|---|
| R6756 | M | 129.5 calCE | 230.5 calCE | 14C | VIMINACIUM; PIRIVOJ; G-349 |
| R9669 | M | 129.5 calCE | 310.5 calCE | 14C | Viminacium, Pirivoj; Gr. 354A |
| R6750 | M | 80.5 calCE | 215.0 calCE | 14C | VIMINACIUM; PIRIVOJ; G-360 |
| R9673 | M | 79 calCE | 213 calCE | 14C | Viminacium, Pirivoj; Gr. 385 |
| R9674 | F | 63 calCE | 204 calCE | 14C | Viminacium, Pirivoj; Gr. 386 |
| R3931 | M | 129.5 calCE | 230.5 calCE | 14C | Viminacium, Pirivoj; Gr. 411 |

**Literature**:

Speal, C. S. 2015 A paleodemographic/mortuary study of graves from the eastern necropolis at Roman Viminacium. *Archaeology and Science* 11: 167–186.

#### Bytča-Hrabové, Slovakia

**Collaborators:** Karol Pieta, Mária Krošláková, Zuzana Hukeľová
**Coordinates:** 49.2056, 18.5662
**Individuals** (n = 2): R2200, R2201

Bytča-Hrabové, Bytča district

Dating: 0 – 50 AD; turn of the La Tène and Roman Periods
Head of excavation: Karol Pieta, Institute of Archaeology, Slovak Academy of Sciences

In 2008, remains of two skeletons lying close to each other were revealed at the site. Both skeletons, with their heads oriented to the south, were found in the western part of the excavated fortification, in the soil coming from the rampart destruction.
Burial customs at the end of the La Tène Period remain unknown in the entire Western Carpathian region. Human remains dated to this period are represented only by single bones found within settlement layers or at sacrificial sites which started to appear at the end of the early Middle La Tène Period (Pieta 2010, 317 – 324; 2018). The discovery of skeletal graves dated to the Púchov culture in the Middle Váh River (north of the Danube) basin was therefore really surprising.
According to archaeological sources, the structure of the settlements in the areas associated with the Púchov culture substantially changed at the beginning of the Roman Period. The whole process started with the destruction and abandonment of the fortifications. Such a major transition is probably related to political turbulence resulting from massive migrations. The background of this process is also mentioned in written sources. Western parts of the Púchov culture were then placed between two expanding formations – the Quadi realm ruled by Vannius, and their enemies, the Lugii, who settled on the other side of the Carpathian crest. Conflicts between these two groups in this period were mentioned, for instance, by Tacitus in his Annals (XII, 29, 30).

| sample_id | sex (genetic) | date lower (95% CI) | date upper (95% CI) | date inference | burial_id |
|---|---|---|---|---|---|
| R2200 | F | 48 calBCE | 56 calCE | 14C | Nr. 1, Obj. 6 |
| R2201 | F | 48 calBCE | 56 calCE | SITE | Bytča-Hrabové, Nr. 2, Obj. 1 |

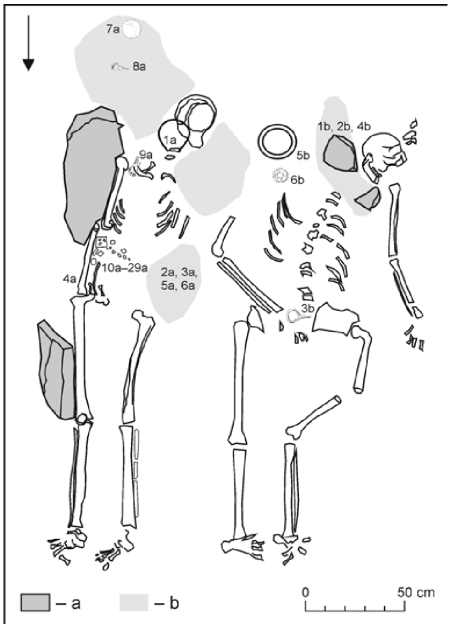

Position of the skeletons from Bytča-Hrabové, Slovakia.

**Literature**:

JAKAB, Július. *Bytča - Hrabové: VS 16 765.* Nitra: AÚ SAV, 2008.

JAKAB, Július. Kostry zo ženského dvojhrobu púchovskej kultúry v Bytči. In *Archeologické výskumy a nálezy na Slovensku v roku 2008*, 2011, 116-118. ISSN 0231-925X.

PIETA, Karol. Early Roman Period Burials of Púchov Culture: Buried Natives or Offered Foreigners?. In *Slovenská archeológia*, vol. LXVII-2, 2019, 241-286. DOI: https://doi.org/10.31577/slovarch.2019.67.8

#### Mikušovce, Slovakia

**Collaborators**: Karol Pieta, Mária Krošláková, Zuzana Hukeľová
**Coordinates**: 49.065963, 18.205994
**Individuals** (n = 1): R2202

Mikušovce, Ilava district

Dating: 0 – 50 AD; turn of the La Tène and Roman Periods
Head of excavation: Karol Pieta, Institute of Archaeology, Slovak Academy of Sciences

**Grave 1**
The grave was oriented in the N/NW – S/SE direction, with the skeleton's head pointing north. In its lowest part, the grave pit was about 50 – 55 cm deep, its upper part was only 15 – 20 cm deep. The skeleton lay in supine position, with the head slightly tilted to the west. Its left hand was stretched along the body, the area around the right hand and the torso was disturbed by an amateur survey. The entire right side of the skeleton was slightly dislocated due to soil erosion, confirmed also by the dislocation of the right femur which had shifted down the slope. In this disturbed part of the grave, fragments of another skull have been found. There were also animal bones (sheep/goat metatarsal bones; analyzed by Z. Bielichová) whose original position in the grave is unknown.
As suggested by anthropological analysis (by J. Jakab), the skeleton belonged to an adult individual (adultus I, 20 – 30 years), with no marked muscle attachment sites. The body height is unknown. The remains of the additional skull belonged to an infant (infans I).

**Grave 2**
The deceased was buried in the layer of black humus, lying in supine position, with legs placed along each other. The body was oriented in N/NE - S/SW direction, with the head towards N/NE. The left arm was placed along the body, while the right (heavily damaged) hand was located within the pelvic area. The skull lay on the left side. Severely damaged facial bones were partially covered with a long animal bone.
According to anthropological analysis (by J. Jakab), the skeleton belonged to a 148 cm tall adult female (adultus I, 20 – 30 years).

| sample_id | sex (genetic) | date lower (95% CI) | date upper (95% CI) | date inference | burial_id |
|---|---|---|---|---|---|
| R2202 | F | 0 CE | 50 CE | ARCH | Mikušovce, Nr. 2, Slovakia, 19.09.18, R. petrous CBD, 0-50AD |

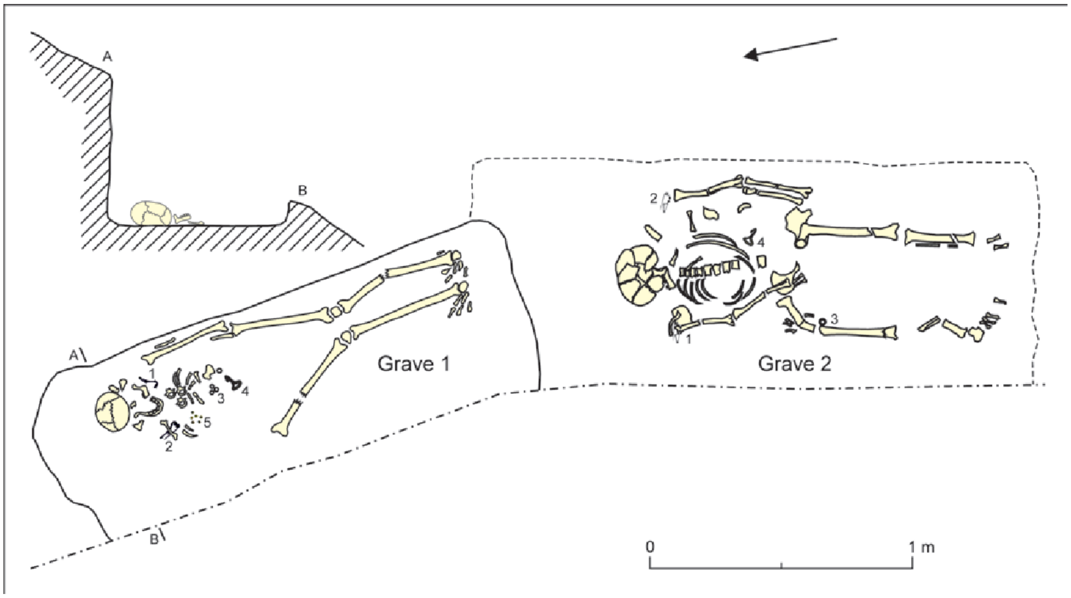

Position of the skeletons from Mikušovce, Slovakia.

**Literature**:
JAKAB, Július. *Mikušovce - Skalice: VS 17428.* Nitra: AÚ SAV, 2011.
PIETA, Karol. Early Roman Period Burials of Púchov Culture: Buried Natives or Offered Foreigners? *Slovenská Archeológia LXVII-2,* 2019, 241-286. DOI: https://doi.org/10.31577/slovarch.2019.67.8

### Tesárske Mlyňany, Slovakia

**Collaborators**: Matej Ruttkay, Mária Krošláková, Zuzana Hukeľová
**Coordinates**: 48.337073, 18.356218
**Individuals** (n = 6): R2206, R2207, R2208, R2209, R2210, R2211

<u>Tesárske Mlyňany, Zlaté Moravce district</u>

Dating: $5^{th}$ – first half of $6^{th}$ century AD; the Migration Period
Head of excavation: Matej Ruttkay, Institute of Archaeology, Slovak Academy of Sciences

The burial ground from the Migration Period was partially excavated in 2002-2003 and 2009. The site is located about 65 km north of the Danube River. A total of 98 graves were examined. It represents the biggest cemetery from the Migration Period in Slovakia, north of the Danube River. Most of the graves were secondarily violated shortly after the burial of the remains (several years after?). Most of the graves were oriented, more or less, in the W-E direction: the deceased were buried with the head to the west, bodies in supine positions, with arms being placed along the body. In some cases, there were two skulls in the burial pit. Although we were unable to unambiguously identify the traces of robberies, some contexts suggest that the graves had been robbed. The violations varied in intensity. Generally, the western parts of graves — where the upper parts of the bodies lay — attracted more attention. Several contexts indicate that when the robberies took place, the soft tissues had not yet fully decomposed.
Because all graves were secondarily violated, the finds were rather modest — beads, metal arm rings, bronze and iron buckles, firesteels, flints, bone combs, etc. Many graves were particularly interesting because they contained skulls with clear signs of artificial deformations. Exceptionally, there were traces of coffins, planks lining the graves or wooden covers. The relatively large number of examined graves indicate that the burial ground belonged to a settlement that had existed for at least several decades or for about one hundred years.
The burial ground is roughly dated back to the 5th century – first half of the 6th century. Historically, this was a time when the area — originally a Germanic settlement — was under Hunnic control for a short time, until the advance of the first Slavic groups. Due to large-scale population movements at this time, ethnic classification is still complicated (assuming Germanic population?).

**List of analyzed individuals:**
Grave 23, male (?), adultus I,
Grave 25, female, adultus I (20-25 years)
Grave 75, female, adultus I
Grave 83, male, adultus II
Grave 88, child, infans II (3-5 years)
Grave 92B, male (?), maturus II

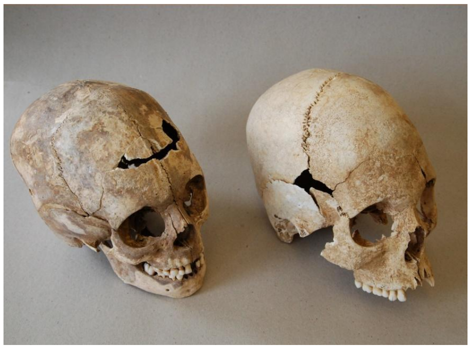

Artificial deformation on the skulls from Tesárske Mlyňany, Slovakia.

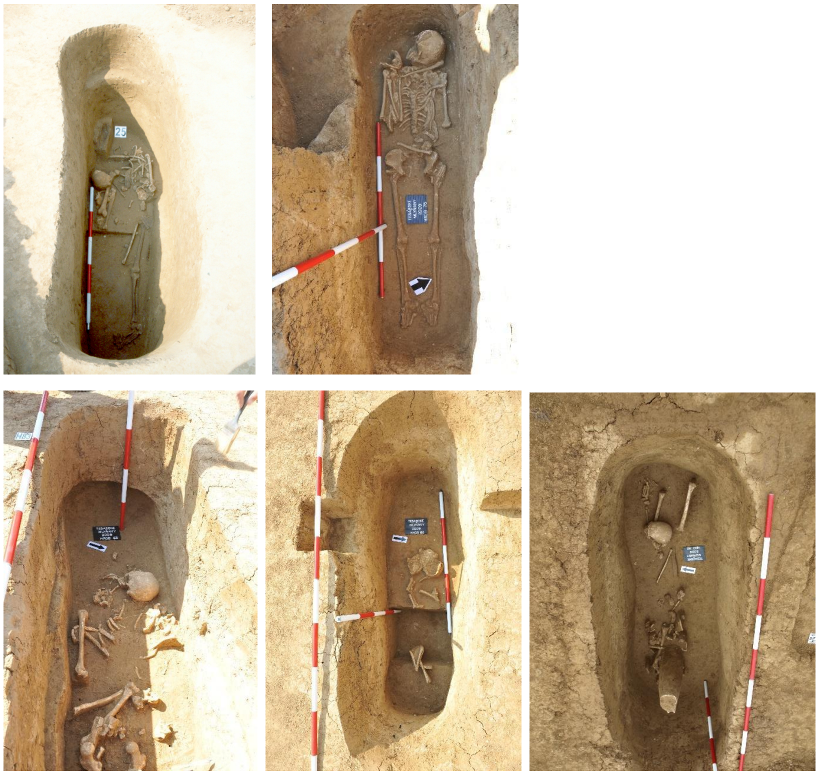

Position of the skeletons from Tesárske Mlyňany, Slovakia.

| sample_id | sex (genetic) | date lower (95% CI) | date upper (95% CI) | date inference | burial_id |
|---|---|---|---|---|---|
| R2208 | F | 200 CE | 400 CE | ARCH | Tes Mlynany, Gr. 23, Nr. 179 |
| R2207 | F | 300 CE | 600 CE | ARCH | Tes Mlynany, Gr. 25, 5th-6th century, Migration Period, L.petrous CBD, 18.9.18 |
| R2209 | F | 200 CE | 600 CE | ARCH | Tes Mlynany, Gr. 75, Nr. 20 |
| R2206 | M | 300 CE | 600 CE | ARCH | Tes Mlynany, Gr. 83, Migration Period, L.petrous CBD, 18.09.2018 |
| R2211 | M | 200 CE | 600 CE | ARCH | Tes Mlynany, Gr. 88, Nr. 48 |
| R2210 | M | 200 CE | 600 CE | ARCH | Tes Mlynany, Gr. 92/B, Nr. 68 |

**Literature**:

JAKAB, Július. *Tesárske Mlyňany - č. Tesáre nad Žitavou - Gočol: VS 18134.* Nitra: AÚ SAV, 2013.

RUTTKAY, Matej. Pohrebisko z doby sťahovania národov v Tesárskych Mlyňanoch. In *Archeologické výskumy a nálezy na Slovensku v roku 2002.* - Nitra: Archeologický ústav SAV, 2003, 112-113. ISSN 0231-925-X.

RUTTKAY, Matej. Druhá sezóna výskumu pohrebiska z doby sťahovania národov v Tesárskych Mlyňanoch. In *Archeologické výskumy a nálezy na Slovensku v roku 2003*, 2004,157-160. ISSN 0231-925X.

RUTTKAY, Matej. Das völkerwanderungszeitliche Gräberfeld in Tesárske Mlyňany, Bez. Zlaté Moravce. In *Barbaren im Wandel : Beiträge zur Kultur- und Identitätsumbildung in der Völkerwanderungszeit. Herausgegeben Jaroslav Tejral ; rezensent Herwig Friesinger.* - Brno: Archeologický ústav AV ČR, 2007, 321-338. ISBN 80-86023-87-7.

RUTTKAY, Matej. The North of the Carpathian Basin in the 5th and 6th Centuries AD. In *Foreigners in Early Medieval Europe: Thirteen International Studies on Early Medieval Mobility.* - Mainz: Römisch-Germanisches Zentralmuseums, 2009, 273-293. ISBN 978-3-88467-131-3.

### Zohor, Slovakia

**Collaborators:** Kristián Elschek, Mária Krošláková, Zuzana Hukeľová
**Coordinates**: 48.3175, 16.9829
**Individuals** (n = 1): R2204

Zohor, Malacky district

Dating: second half of the 1st century AD
Head of excavation: Kristián Elschek, Institute of Archaeology, Slovak Academy of Sciences

During the rescue excavation in 2010, a single-body grave (Nr. 230) at the edge of the humus and topsoil layers was excavated. The grave was located within the area of the Germanic settlement area from the 1st century AD. The grave of a Germanic noble was examined in the northern part of the area in 2010. North of the grave, a central Germanic settlement, dated to the 1st-4th centuries AD, was discovered.

**Grave 230**
The deceased lay stretched in supine position. There was an 'Almgren 68' pin on each shoulder, two small ceramic vessels, an iron knife and a bronze sewing needle located at the feet. According to anthropological analysis (by J. Jakab), the individual was an elderly female, about 50-60 years of age, and her height was approximately 161-164 cm. The woman was buried during the 2nd half of the 1st century AD and represents a new Germanic migration wave within the Middle Danube area at the border of the Roman Empire.

| sample_id | sex (genetic) | date lower (95% CI) | date upper (95% CI) | date inference | burial_id |
|---|---|---|---|---|---|
| R2204 | F | 100 CE | 200 CE | ARCH | Zohor, Gr. 2, Obj. 230, R. petrous CBD, 19.09.2018 |

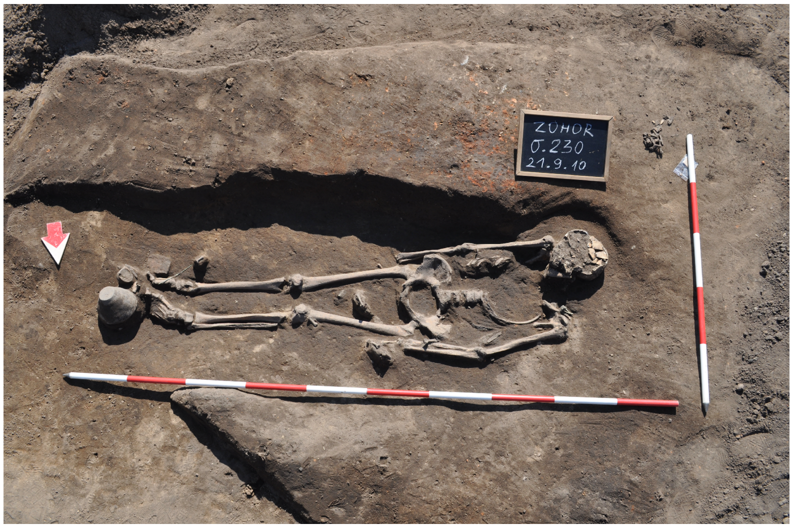

Position of the skeleton from Zohor, Slovakia.

**Literature**:

Elschek, K.: ZOHOR – Ein neues Fürstengrab der „Lübsow- Gruppe“ und Brandgräber mit Edelmetallbeigaben aus Zohor (Westslowakei). In: *Grundprobleme. Thema: Macht des Goldes - Gold der Macht. Forschungen zu Spätantike und Mittelalter 2. M. Hardt, O. Heinrich-Tamáska (Hrsg.).* Weinstadt 2013, 91-123. ISBN 978-3-86705-071-5.

Elschek, K.: Nové žiarové hroby z 2. polovice 2. storočia zo Zohora na západnom Slovensku (Zusammenfassung: Neue Brandgräber aus der 2. Hälfte des 2. Jahrhunderts von Zohor, Westslowakei). In: *Zborník SNM 2014 – Supplementum 8, Column of Marcus Aurelius and the Middle Danube area.* Bratislava 2014a, 41-50. ISBN 978-80-8060-335-9.

Elschek, K.: Zohor v dobe rímskej. Nový germánsky kniežací hrob a žiarové pohrebiská na Záhorí (Zusammenfassung: Zohor während der römischen Kaiserzeit, ein neues germanisches Fürstengrab und Brandgräberfelder im Záhorie-Gebiet). In: B. Komoróczy (Ed.) *Archeológie Barbarů-Archäologie der Barbaren 2011. Spisy AÚ AV ČR Brno 44.* Brno 2014b, 113-131. ISBN 978-80-86023-25-0.

JAKAB, Július. Dve kostry zo staršej doby rímskej v Zohore = Zwei Skelette aus der älteren Römerzeit in Zohor. In *Archeologické výskumy a nálezy na Slovensku v roku 2010*, 2015, 119-120. ISSN 0231-925X.

JAKAB, Július. Z*ohor - ASA: VS 17429.* Nitra: AÚ SAV, 2011.

Kolník, T.: Germánske hroby zo staršej doby rímskej zo Zohora, Žlkoviec a Kostolnej pri Dunaji (Germanische Gräber aus der älteren römischen Kaiserzeit in Zohor, Žlkovce und Kostolná pri Dunaji). *Slovenská Archeológia 7*, 1959, 144-162.

Kraskovská, Ľ. : Hroby z doby rímskej v Zohore (Gräber aus der römischen Zeit in Zohor). *Slovenská. Archeológia* 7, 1959, 99-143.

### Emona, Slovenia

**Collaborators**: Andrej Gaspari, Rene Masaryk, Tamara Leskovar
**Coordinates**: 46.0569, 14.5058
**Individuals** (n = 7): R10467, R10469, R10471, R10473, R10474, R10477, R10478

<u>Late Roman cemetery at Gosposvetska road in Ljubljana</u>

*Colonia Iulia Emona* (present-day Ljubljana, the capital of Slovenia) was an autonomous town in the northeastern part of Roman Italy (regio X). Its importance is closely linked with the favourable position at the crossroad of land and water routes connecting Italy with continental provinces and Eastern regions of the Empire. The city flourished between Augustan deduction and the 5th century, afterward rapidly falling into decline. Settled by colonists (veterans, merchants, entrepreneurs) of north and central Italian origin with a certain number of natives, belonging to the North Adriatic group of pre-Roman peoples with Celtic elements. During the 2nd c. and later periods, a significant influx of migrants from the East is hypothesized from the archaeological and partly onomastic evidence. The existence of the Greek-speaking Early Christian community in Emona is attested both in written sources and archaeologically.
The town had three major necropolises, situated along road approaches to the city from the South-west (direction towards Aquileia), East (Siscia, Pannonia), and the North (Poetovio, Pannonia). The total number of documented graves reaches between 3500 and 4500. The northern necropolis, extending 2.5 km from the North city gates, was relatively well researched before different spatial interventions from 1907 onwards, but especially extensively during construction works in the 1960s and after 2000. During the early and middle Imperial periods, it consisted of family grave groups within built enclosures as well as of numerous individual graves along both sides of via publica Emona-Celeia. In the Late Roman period, when inhumation completely prevailed over cremation, interment occurred both in small groups of burials in particular areas of old necropolises, often with grave pits cut into the rubble of abandoned buildings outside the city.
Excavations of the Gosposvetska road, conducted in 2017/2018, supplemented previous information on this particular area of Northern necropolis of Emona and revealed a cemeterial complex, spatially separated from the neighboring burial ground. The internment at this location began around the mid 3rd century and lasted until the first decades of the 5th century. The complex is characterized by multi-phased masonry buildings south of the *via publica* with its long axis aligned perpendicular at the road. The initial phase of the cemetery is marked by square cella, hosting the earliest burials. After approx. 30-50 years the walls of the cella were leveled to the ground, followed by the construction of a larger rectangular, hall-like building with subsequent annexes. The central building had mortar pavements and plastered walls, the western annexe,

interpreted as a mausoleum, and also poorly preserved mosaic floor and arcosolia. The buildings, most probably with tile roofing, contained interred sarcophagi, masonry tombs, stone and lead coffins as well as burials in plain pits. Altogether (with previous research) more than 400 skeletal graves were documented on the site, and the projected total number could reach up to some 500 buried individuals, based on the assumptions about the size of the un-researched areas of the complex. The epigraphic evidence from two slabs, covering the children's graves in the hall of the central building, supports the assumption that the cemetery appeals to the segregated and highly organized community, most probably of religious background.

Most recent excavations of the Gosposvetska road (2017/2018) yielded 327 graves in which 388 individuals were buried. Osteological and palaeopathological analyses were performed on all the 388 individuals, 18 individuals were sampled for aDNA and 28 for isotopic (C, N, O, Sr) analyses. As the analyses are still in progress, the results are not published yet.

In sum, most of the graves were singular (266), while occasionally two to eight individuals were buried together. 33% of the analysed individuals were juveniles and 66% were adults. 1% of the remains was too poorly preserved for the age assessment. Though sex of juveniles was occasionally assessed (25 % females and 18 % males), the reliability of macroscopically assessed sex in juveniles is low and needs to be supported by additional analyses (e.g. aDNA or enamel peptides). Among adult individuals, 37% were male or probably male, 46% were female or probably female, while 17% remain undetermined due to poor preservation. Average female height was 159 ± 4 cm and an average male height 169 ± 4 cm.

Among children, most of the observed pathological changes were related to various infections and/or signs of malnutrition/metabolic diseases. Rarely other pathological changes were observed, such as trauma and neoplasms. Among adults, there were a lot of pathological changes related to dentition (e.g. antemortem tooth loss, caries, infections, cysts, periodontal disease) and joints (e.g. DISH, AS, osteoarthritis). Occasionally, indications of malnutrition and/or metabolic disease, myositis, trauma, neoplasms and congenital defects were observed.

Isotopic analyses highlighted some possible outliers based on diet (low protein consumption) and origin (outstanding Sr and/or O values). However, none of the outliers had successful aDNA sequencing.

All the remains are stored in the Museum and Galleries of Ljubljana.

| sample_id | sex (genetic) | date lower (95% CI) | date upper (95% CI) | date inference | burial_id |
|---|---|---|---|---|---|
| R10467 | F | 252 calCE | 402 calCE | 14C | Emona 2 |
| R10469 | F | 242.5 calCE | 350 calCE | 14C | Emona 4 |
| R10471 | F | 241 calCE | 341 calCE | 14C | Emona 6 |
| R10473 | F | 248.5 calCE | 378 calCE | 14C | Emona 8 |
| R10474 | F | 257.5 calCE | 404.5 calCE | 14C | Emona 9 |
| R10477 | F | 238 calCE | 333.5 calCE | 14C | Emona 12 |
| R10478 | M | 255 calCE | 402.5 calCE | 14C | Emona 13 |

#### Tell Masaikh, Syria

**Collaborator**: Arkadiusz Sołtysiak
**Coordinates**: 34.973503, 40.555681
**Individuals** (n = 2): R3359, R3361

Tell Masaikh is located in eastern Syria, on the left bank of the Euphrates, c. 22 km downstream of Khabur confluence (34°58'23"N, 40°33'13"E). It has been inhabited during the Chalcolithic (Halaf culture, 6th millennium BCE), then in the Middle Bronze Age (first half of the 2nd millennium BCE) and finally in the Neo-Assyrian period (9th-7th centuries BCE) when a regular rectangular town covering ca. 25 ha was established next to a palace that was likely the seat of the governor of the Assyrian Rasappa province. After the fall of the Neo-Assyrian empire it has been largely abandoned and only the remains of a hamlet are dated to later periods. However, the ruins of the Assyrian town were used for many centuries as a cemetery, with the majority of graves dated tentatively to the Late Roman and Early Islamic (i.e., Ubayyad and Abbasid) periods.

Regular excavations at the site were conducted between 1996-2011 by a French-Syrian team directed by Maria Grazia Masetti-Rouault (École Pratique des Hautes Études – Sorbonne, Paris, France) (Masetti-Rouault & Salmon, 2010). In total, 404 skeletons were unearthed at Tell Masaikh between 1998-2007. Among them, 56 (mainly subadults) were dated to the Late Roman period, and 116 to the Early Islamic period, with almost 200 skeletons dated only broadly to the time between the fall of the Neo-Assyrian empire and major depopulation of the region due to the Mongolian invasion during the 13th century CE (Sołtysiak & Tomczyk, 2008). As most of them were buried in simple pit graves with no artifacts, sometimes the only chronological indicator was grave orientation (Frank, 2006).

Tell Masaikh and nearby Tell Ashara, a Bronze Age site located a few kilometers south to Tell Masaikh with the total number of skeletons above 650, represented several periods of occupation of the region from c. 2700 BCE to c. 1950 CE, and provided very good samples to study changes in economy and living conditions in the local population. There is evidence of subsistence continuity between the Neo-Assyrian and Early Islamic period (Sołtysiak & Schutkowski, 2018), although higher prevalence of linear enamel hypoplasia suggests decline in living conditions during the Late Roman period (Tomczyk et al., 2007). Research on dental non-metric traits has suggested population continuity until the Mongolian invasion when the region was largely deserted and then re-populated by Bedouin tribes from the Arabian Peninsula (Sołtysiak & Bialon, 2013).

| sample_id | sex (genetic) | date lower (95% CI) | date upper (95% CI) | date inference | burial_id |
|---|---|---|---|---|---|
| R3359 | M | 680 calCE | 827.5 calCE | 14C | Tell Masaikh, 7-E-58 |
| R3361 | F | 1176 calCE | 1264 calCE | 14C | Tell Masaikh, 7-F-77 |

**Literature**:

Frank, C. A. (2006). Funeral practices at Tell Masaikh (Syria): Late Roman and Islamic graves. Studies in Historical Anthropology, 3, 93–120.

Masetti-Rouault, M. G., & Salmon, S. (2010). The Neo-Assyrian colony of Tell Masaikh in the region of the Syrian lower middle Euphrates valley: Report on the latest excavations. In P.

Matthiae, F. Pinnock, L. Nigro, & N. Marchetti (Eds.), Proceedings of the 6th International Congress of the Archaeology of the Ancient Near East (Vol. 2, pp. 385–396). Wiesbaden: Harrassowitz Verlag.

Sołtysiak, A., & Bialon, M. (2013). Population history of the middle Euphrates valley: Dental non-metric traits at Tell Ashara, Tell Masaikh and Jebel Mashtale, Syria. HOMO - Journal of Comparative Human Biology, 64(5), 341–356. https://doi.org/10.1016/j.jchb.2013.04.005

Sołtysiak, A., & Schutkowski, H. (2018). Stable isotopic evidence for land use patterns in the Middle Euphrates Valley, Syria. American Journal of Physical Anthropology, 166(4), 861–874.

https://doi.org/10.1002/ajpa.23480

Sołtysiak, A., & Tomczyk, J. (2008). Short fieldwork report. Tell Masaikh (Syria), seasons 1998–2007. Bioarchaeology of the Near East, 2, 94–97.

Tomczyk, J., Sołtysiak, A., & Tomczyk-Gruca, M. (2007). Temporal changes in frequency of enamel hypoplasia in the middle Euphrates valley (Syria). In E. B. Bodzár & A. Zsákai (Eds.), Human Diversity and Biocultural Researches. Selected papers of the 15th Congress of the European Anthropological Association (pp. 87–97). Budapest: Department of Biological Anthropology.
