## Appendix 4 - Clustering results for "Stable population structure in Europe since the Iron Age, despite high mobility"

For each region, we show the results from pairwise individual-based qpAdm analysis in the form of heatmaps, annotated by dendrograms from the hierarchical clustering analysis. The heatmap color scheme is chosen to signify non-rejected hypotheses (above 0.05) in shades of red and orange, and rejected hypotheses in shades of gray and yellow.
Columns of the heatmaps are annotated by time period: Copper Age (CA), Bronze Age (BA), Iron Age (IA), Imperial Rome & Late Antiquity (IRLA), Middle Ages & Early Modern (MAEM).
Rows of the heatmaps are annotated by cluster assignment based on the hierarchical clustering (see Methods). Clusters identified as outliers through our pipeline downstream are also annotated on the rows. Please note that outlier detection was only performed for individuals from historical periods (IA, IRLA, MAEM).

#### Armenia:

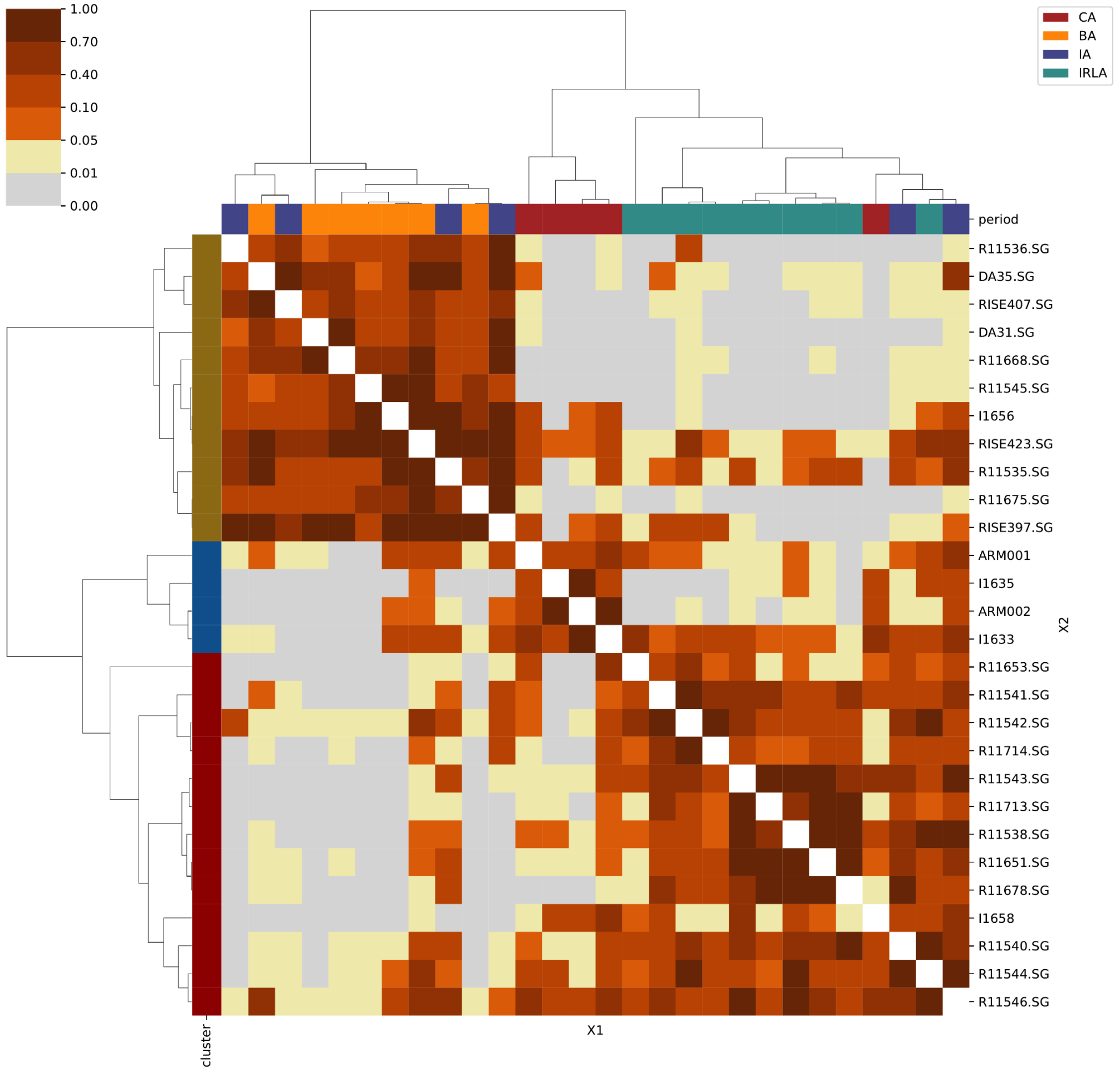

### Mainland Italy:

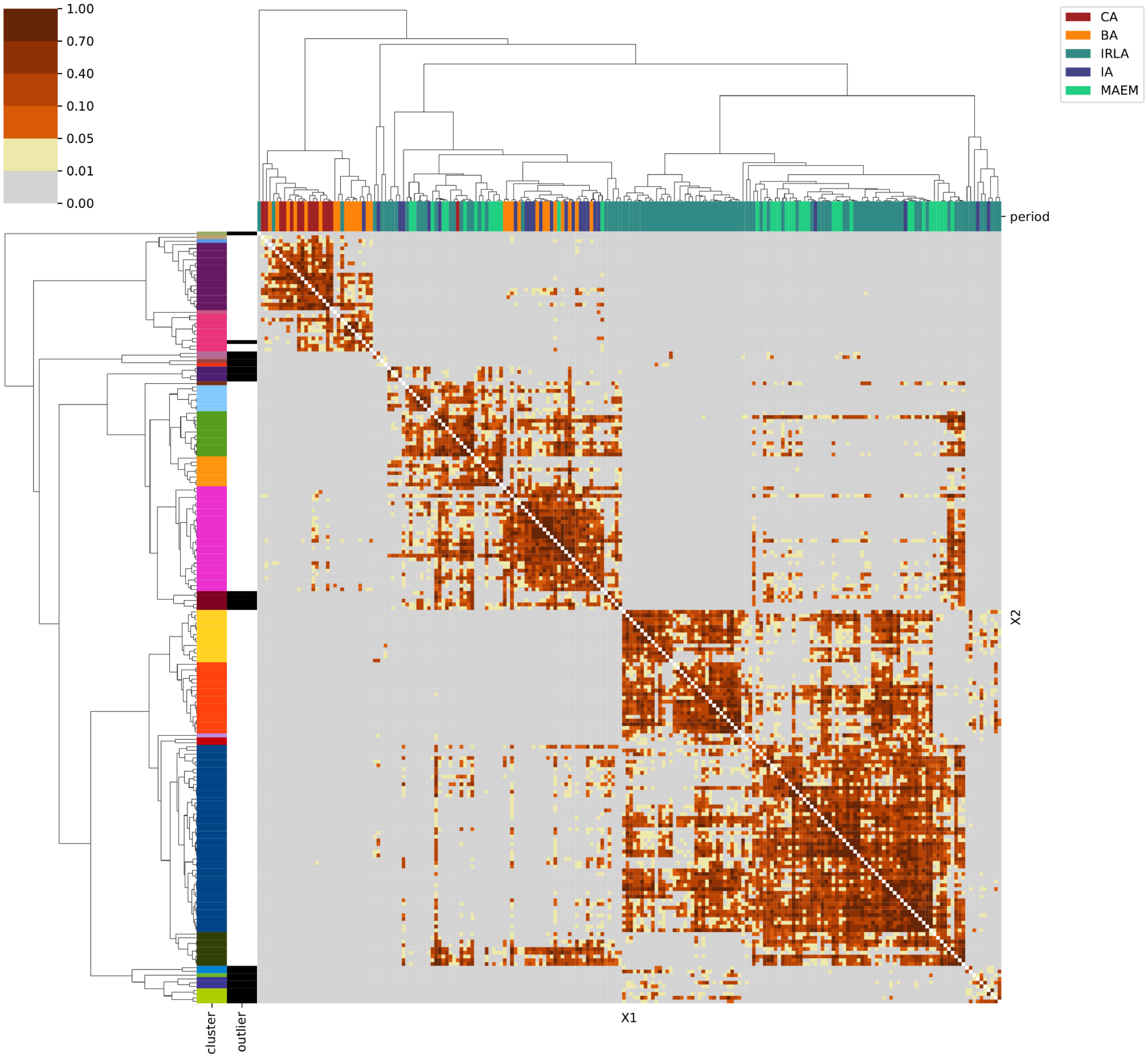

### Sardinia:

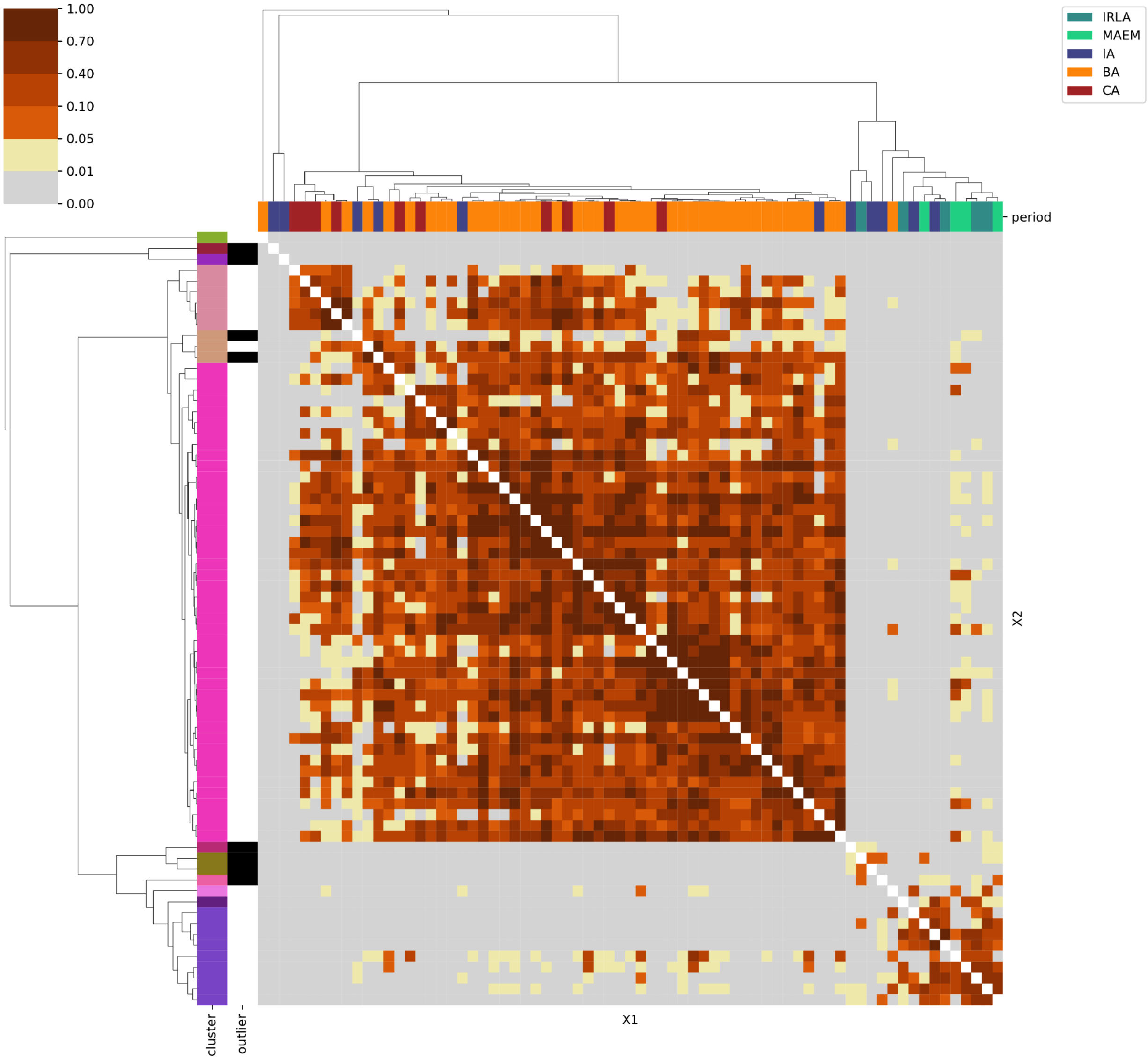
1.00
0.70
0.40
0.10
0.05
0.01
0.00
IRLA
MAEM
IA
BA
CA
period
X2
X1
cluster
outlier

### Levant & Egypt:

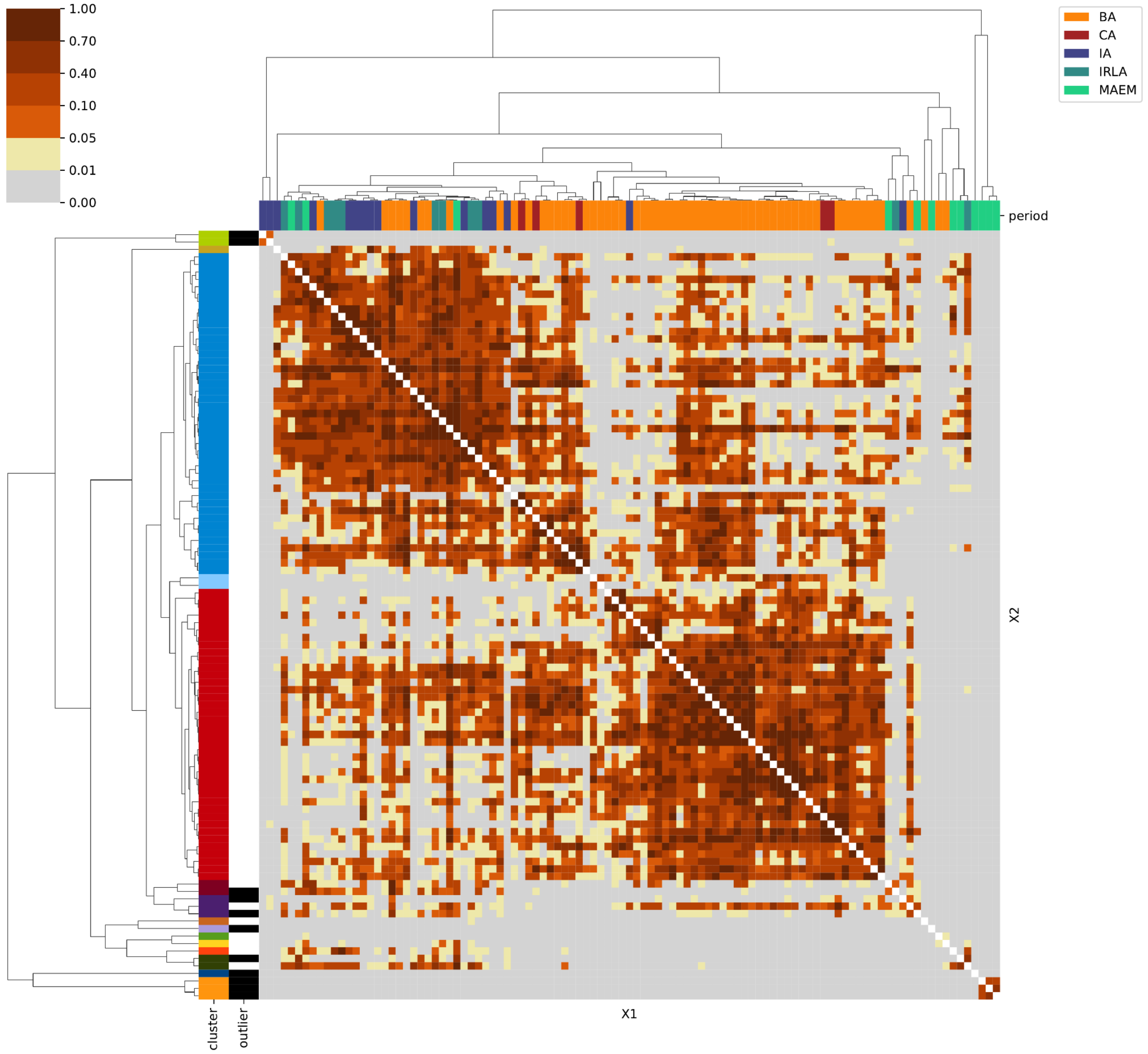

### Southeastern Europe:

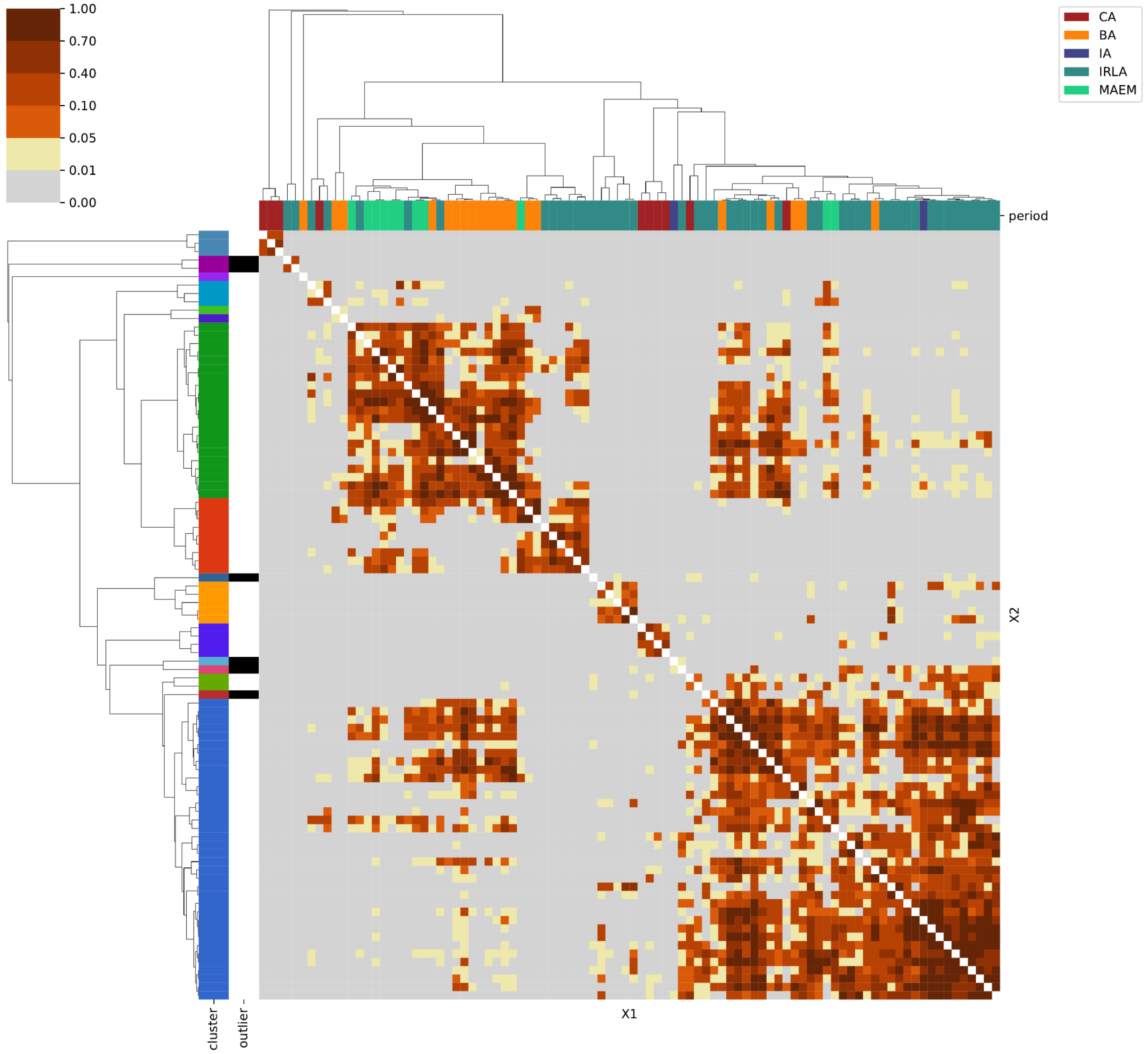

Southwestern Europe:

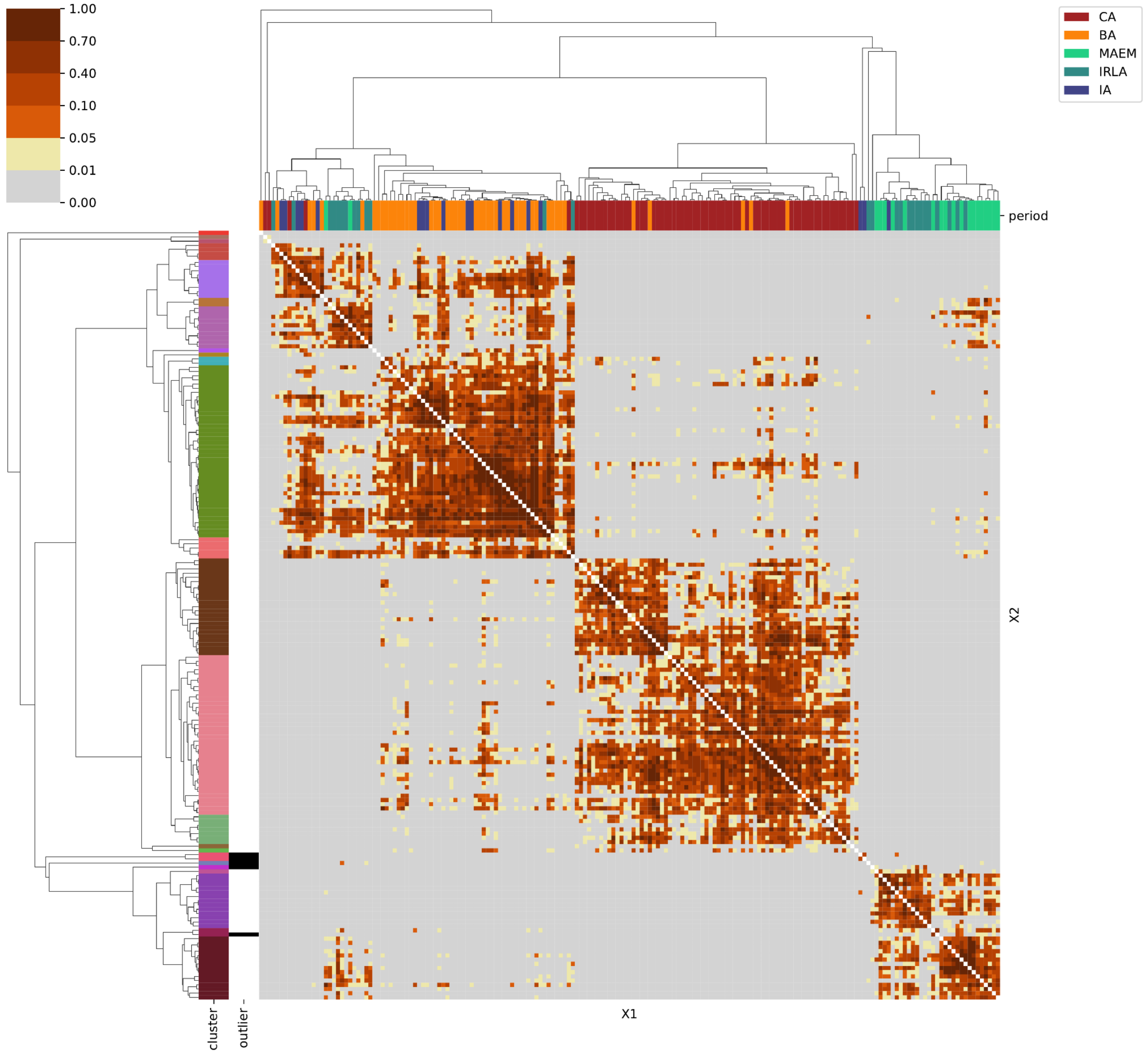

### Western Europe:

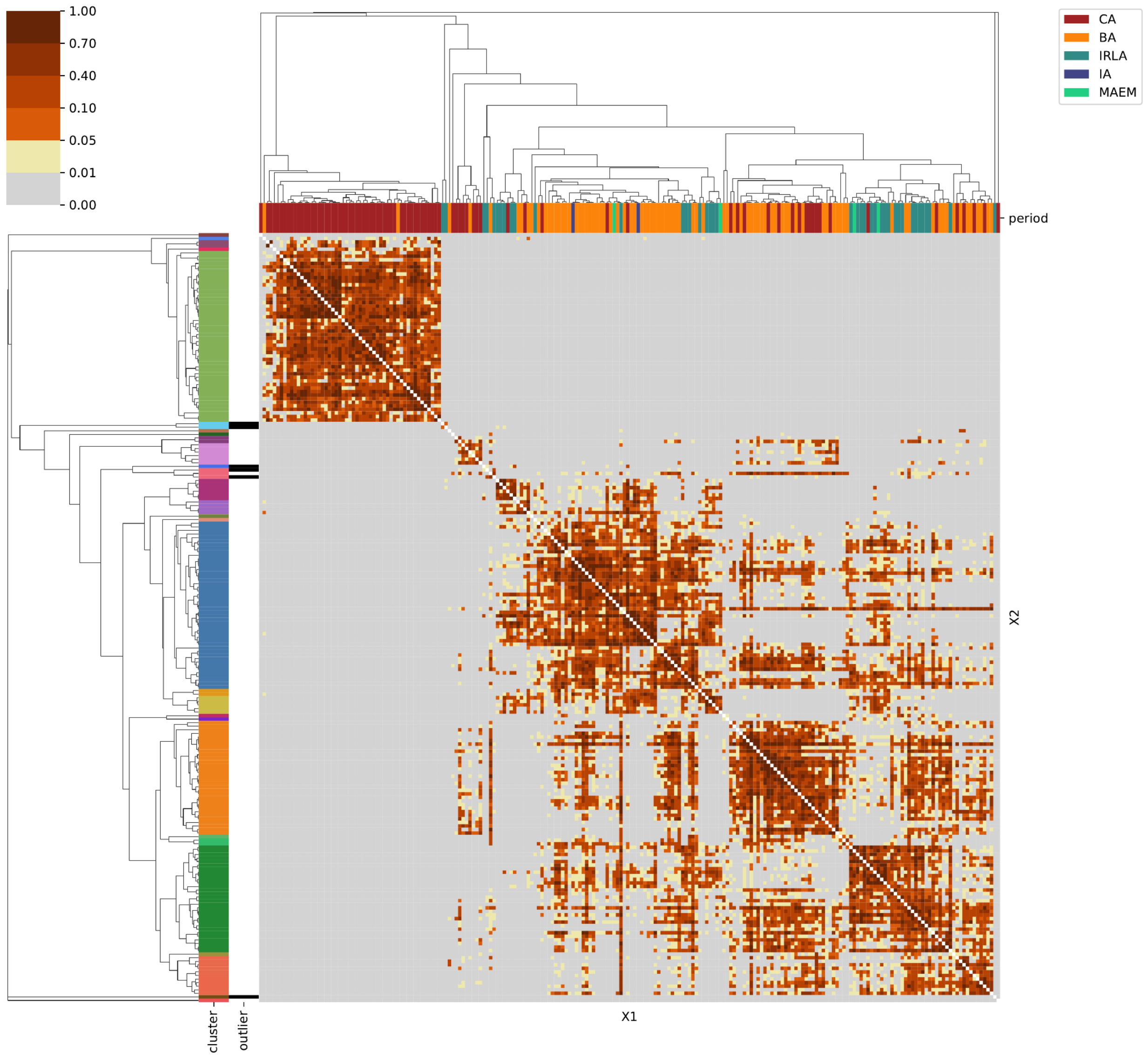

### Northern Europe:

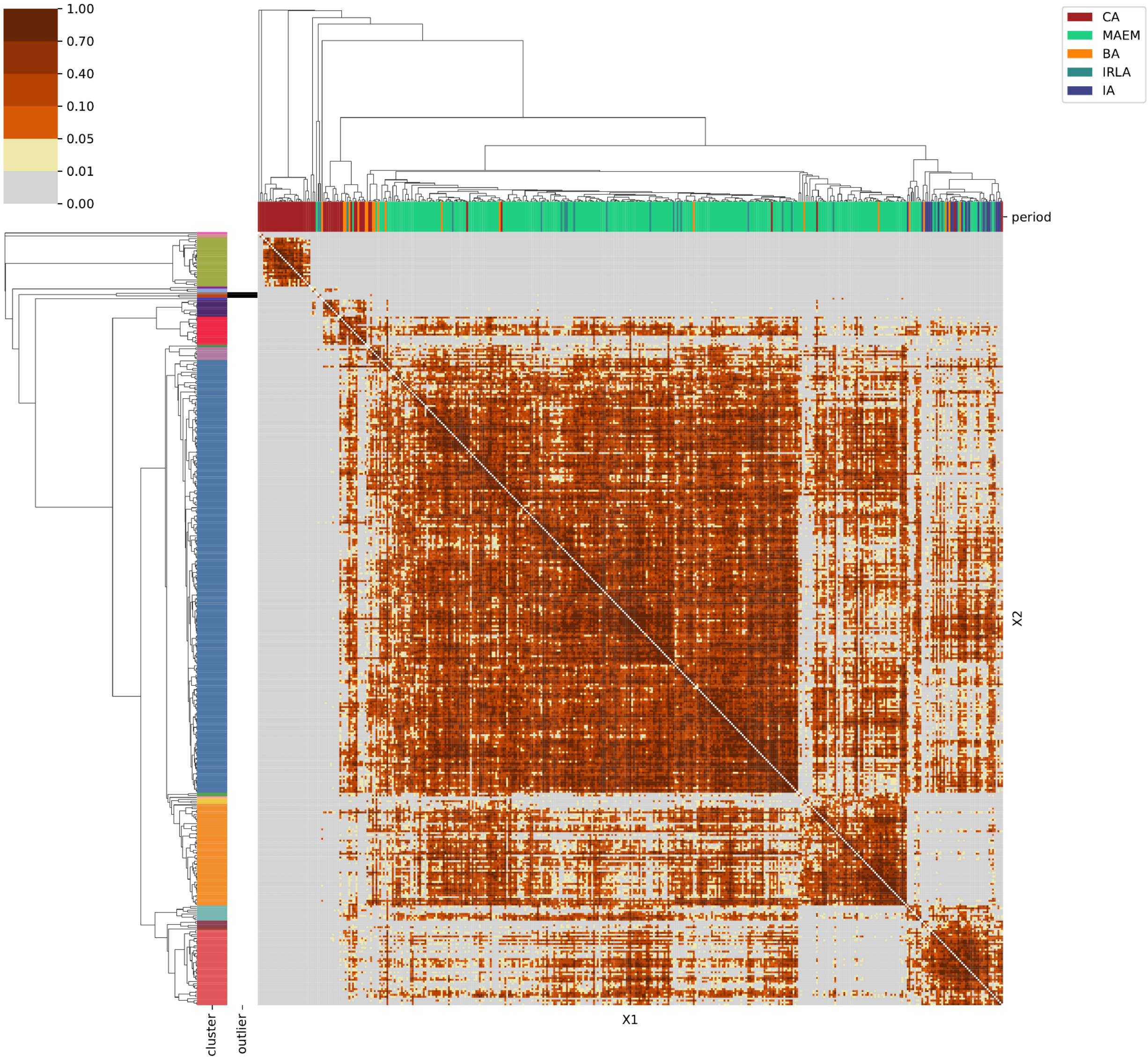

### Great Britain & Ireland:

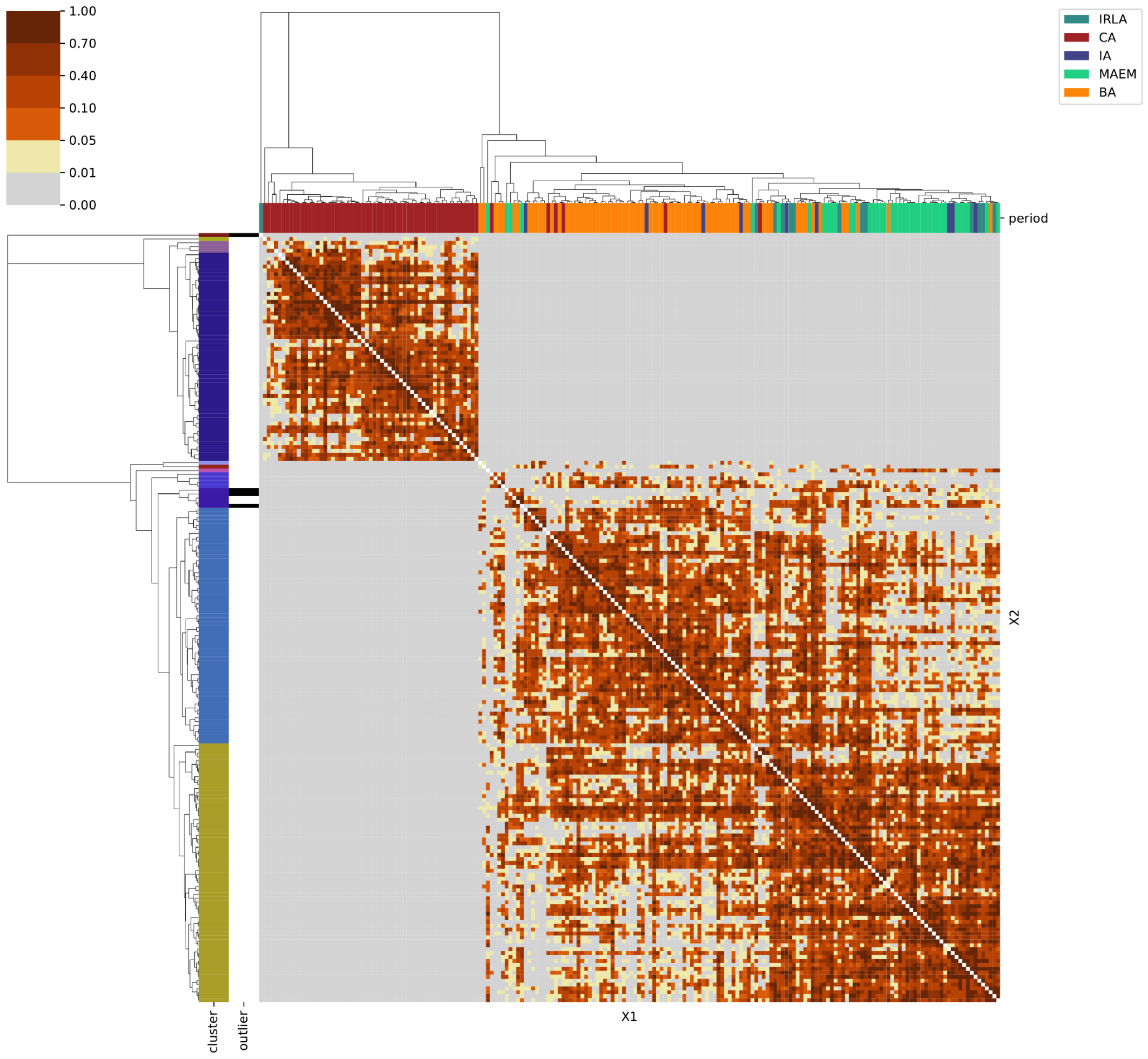

### Eastern Central Europe:

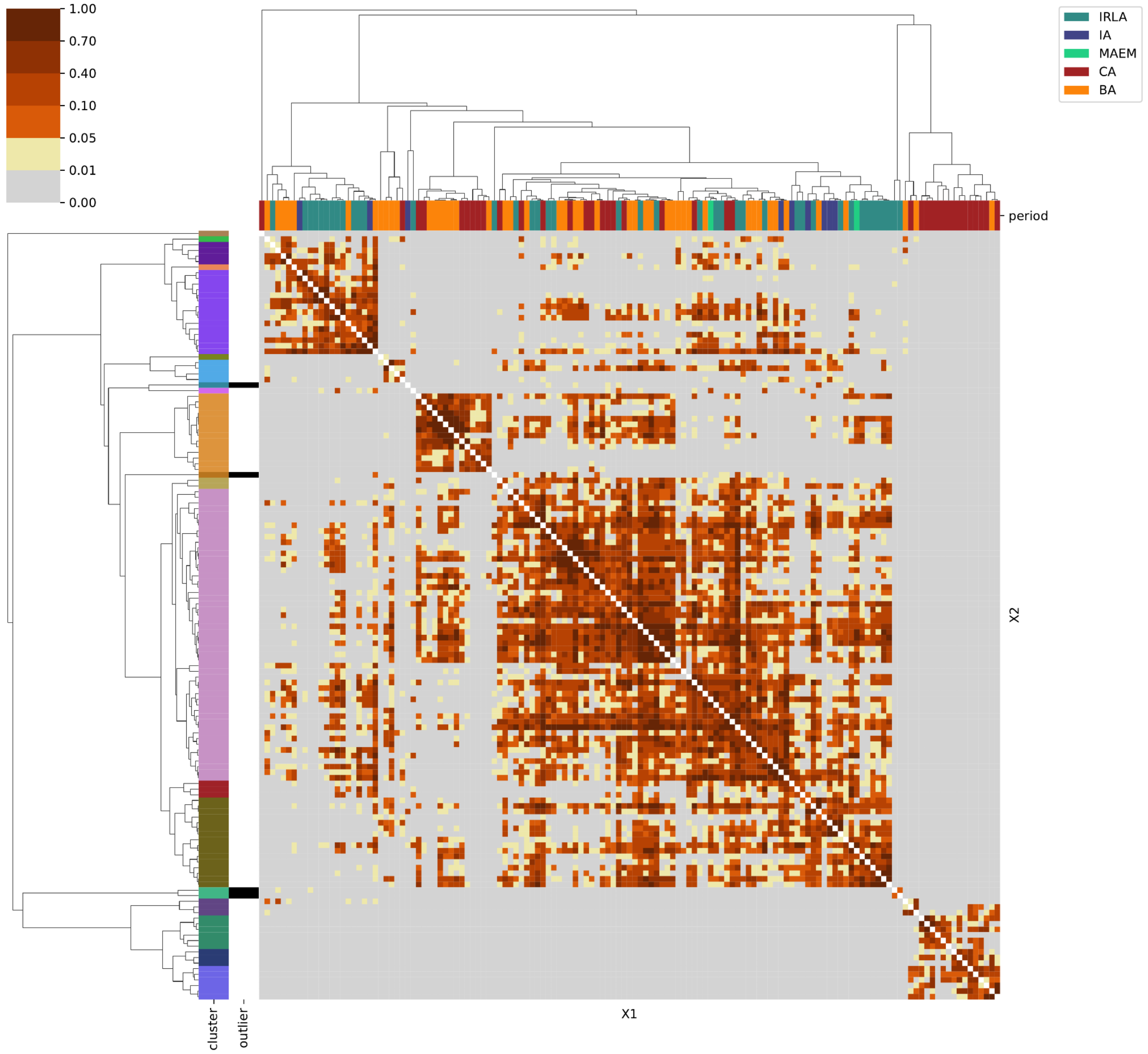

### Eastern Europe & Steppe:

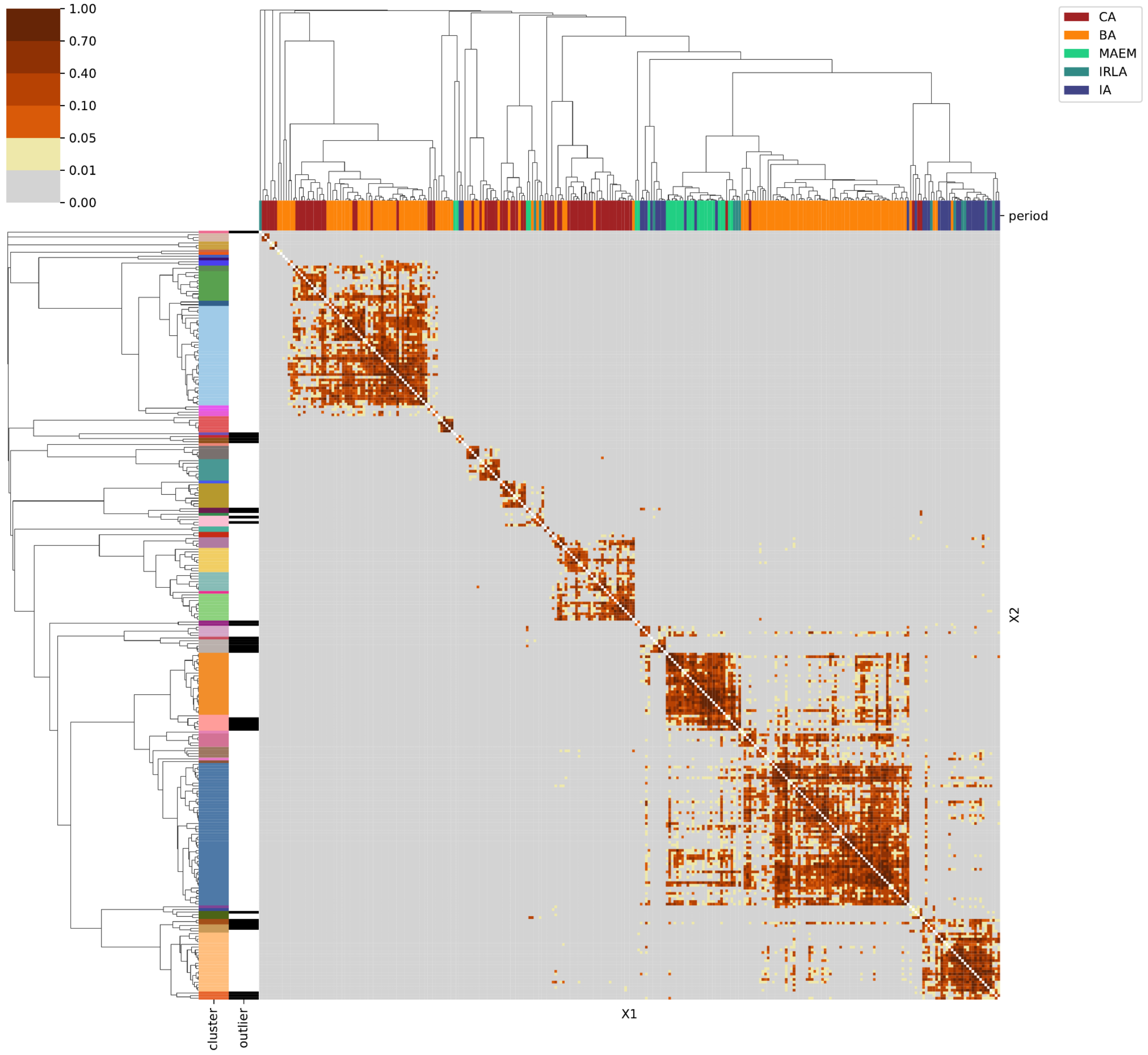

### North Africa:

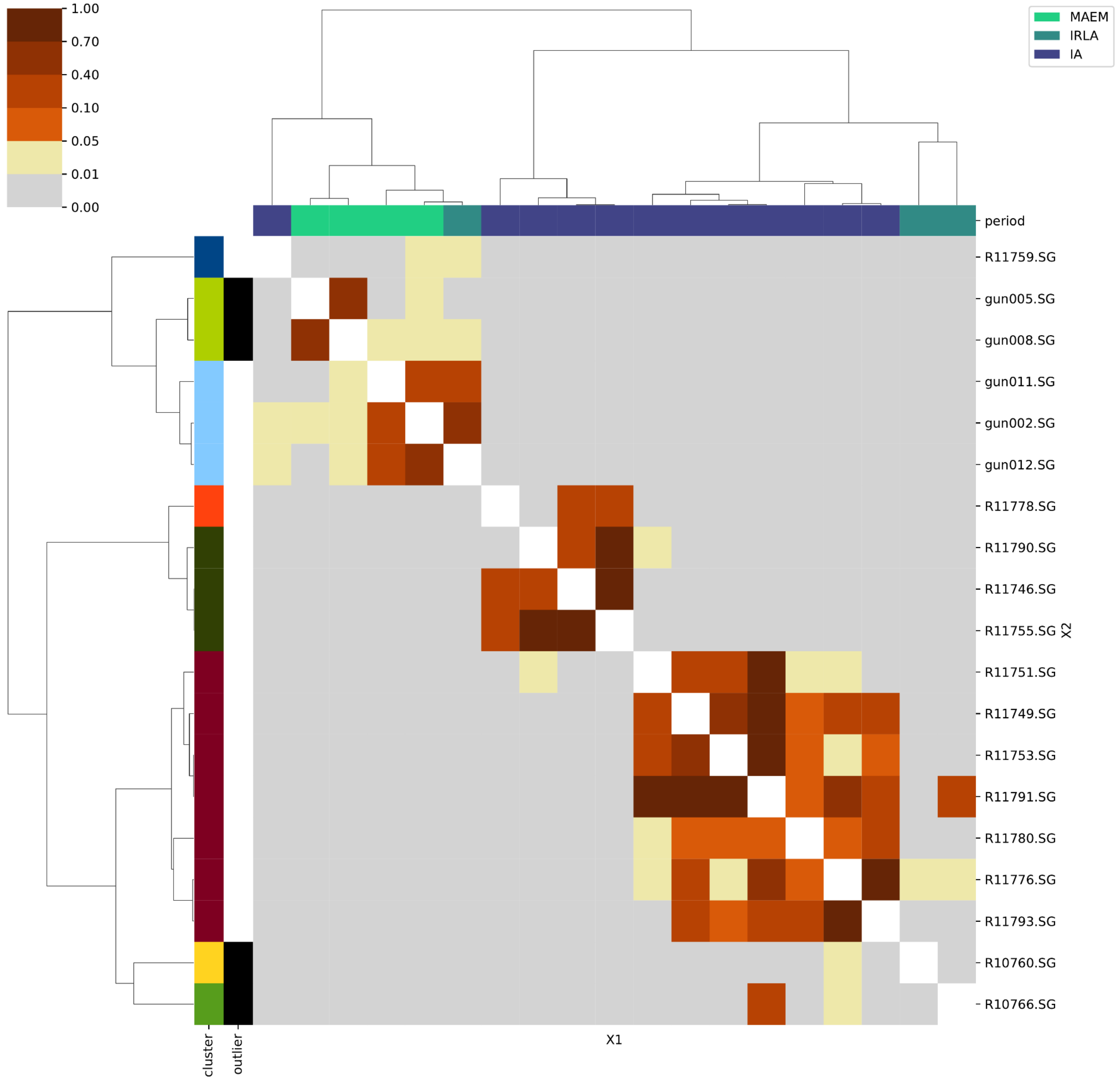
